# Metabolic perturbations to an *E. coli*-based cell-free system reveal a trade-off between transcription and translation

**DOI:** 10.1101/2023.03.22.533877

**Authors:** Manisha Kapasiawala, Richard M. Murray

## Abstract

Cell-free transcription-translation (TX-TL) systems have been used for diverse applications, but their performance and scope are limited by variability and poor predictability. To understand the drivers of this variability, we explored the effects of metabolic perturbations to an *E. coli* Rosetta2 TX-TL system. We targeted three classes of molecules: energy molecules, in the form of nucleotide triphosphates (NTPs); central carbon “fuel” molecules, which regenerate NTPs; and magnesium ions (Mg^2+^). Using malachite green mRNA aptamer (MG aptamer) and destabilized enhanced Green Fluorescent Protein (deGFP) as transcriptional and translational readouts, respectively, we report the presence of a trade-off between optimizing total protein yield and optimizing total mRNA yield, as measured by integrating the area under the curve for mRNA time-course dynamics. We found that a system’s position along the trade-off curve is strongly determined by Mg^2+^ concentration, fuel type and concentration, and cell lysate batch, and that variability can be reduced by modulating these components. Our results further suggest the trade-off arises from limitations in translation regulation and inefficient energy regeneration. This work advances our understanding of the effects of fuel and energy metabolism on TX-TL in cell-free systems and lays a foundation for improving TX-TL performance, lifetime, standardization, and prediction.

**For Table of Contents Use Only:** 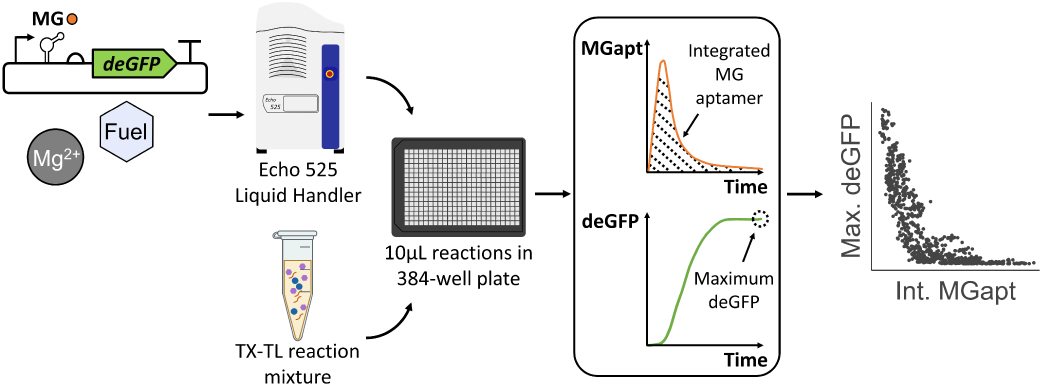

## Introduction

Cell-free synthetic biology is a rapidly growing field that leverages synthetic biology tools and techniques to engineer biological systems outside the traditional context of living cells^1^. The cell-free approach offers many advantages over engineering cellular systems, including greater control over the reaction environment, the ability to produce toxic small molecules or proteins that are difficult to express in living cells, and scalability for biomanufacturing and other applications. In recent years, cell-free systems have been used for protein engineering^1,2^, rapid and high-throughput prototyping of engineered biological components and systems^3–7^, biosensing and diagnostics^8–10^, and the development of synthetic life^11–13^.

Most cell-free applications rely on cell-free protein synthesis, where crude cell lysate or purified proteins are supplemented with DNA and an energy buffer containing small molecules that enable *in vitro* transcription and translation (TX-TL)^14–16^. Although cell lysate-based TX-TL systems have many uses, their performance and scope are limited by issues such as batch-to-batch variability^17^, lack of predictability^18^, and limited lifetime^19^. These issues arise because cell lysate’s proteomic composition largely reflects the cytosolic protein content of living cells. While some of these proteins are harnessed to regenerate energy molecules for TX-TL processes, most of them generate metabolites that are benign or even harmful for TX-TL.

To address these concerns, previous studies have sought to gain insight into TX-TL dynamics through a variety of experimental approaches. Most of these studies have focused on gaining detailed insight into how perturbations to TX-TL reaction conditions – including salts^20,21^, fuel sources^22,23^, crowding agents^20,24^, and other components^15,19,25–28^ – affect TX-TL performance. Some studies have taken a less targeted approach, with the authors opting instead to harness lab automation and machine learning along with experiments to rapidly optimize TX-TL systems^27,29^. While these studies have helped elucidate which reaction components and preparation steps affect TX-TL performance and predictability most significantly, more work is necessary to determine why these factors contribute to variability and limited TX-TL lifetime, as well as the broader implications they have for design principles for TX-TL systems.

Aiming to get a more holistic view of cell-free reaction dynamics, more recent studies have focused on measuring small molecules – including NTPs, amino acids, and central carbon metabolites – that participate in cell-free metabolism and studying or modeling their effects on TX-TL^30–33^. In one such study, the authors found that the majority of *E. coli* metabolic pathways are active during TX-TL^33^, and they subsequently used these insights to form a phenomenological model of TX-TL coupled to cell-free metabolism, with the model containing “fuel,” “energy,” and “waste” species and their effects on TX-TL. By linking cell-free metabolism’s effects on TX-TL dynamics to specific TX-TL reaction components, these studies have suggested that further insight into TX-TL predictability, variability, and lifetime could come from further investigation of the effects of metabolism on TX-TL.

Building off the experiments and insights from these previous studies, to understand the effect of cell-free metabolism on TX-TL variability, we focused our efforts on the chemical composition of the buffer that is used with cell lysate and DNA to form a TX-TL reaction. We targeted two classes of small molecules to modulate: “energy” molecules, specifically nucleotide triphosphates (NTPs), which power TX-TL processes; and “fuel” molecules, which regenerate NTPs via cell lysate metabolism. We also considered the effects of magnesium (Mg^2+^), both because of its biological importance – in stabilizing nucleic acids and ribosomes, acting as an essential cofactor for many enzymes, and making ATP bioactive in the form of Mg-ATP – and because of several previous findings suggesting its role in modulating TX-TL dynamics^21,24,25,28^.

Using mRNA aptamer Malachite Green (MG aptamer) and destabilized enhanced Green Fluorescent Protein (deGFP) as transcriptional and translational readouts, respectively, we report the presence of a trade-off between optimizing total protein yield and optimizing total mRNA yield, as measured by integrating the area under the curve for mRNA time-course dynamics. We found that the trade-off is present across different fuel sources, that a system’s position along the trade-off curve is determined strongly by Mg^2+^ concentration and fuel type, and that the trade-off curve’s location shifts and range becomes larger as DNA concentration is increased. In systems where transcription and translation were decoupled, we found that a distinct regime optimized for translation exists in the translation-only system, but that no distinct regime was optimized for transcription in the transcription-only system, suggesting that the trade-off arises at the translational level. Finally, in systems where additional energy is supplied and where a fuel source is absent, the trade-off is absent. Overall, our results suggest the trade-off arises from limitations in translation regulation and inefficient energy regeneration. By improving our understanding of the effects of fuel and energy metabolism on TX-TL in cell-free systems, this work provides insight into design considerations for future studies aimed at improving TX-TL performance, lifetime, standardization, and prediction.

## Results and Discussion

### E. coli Rosetta2 cell-free systems exhibit a trade-off between transcription and translation across fuel sources and concentrations

As most previous studies have utilized either a design-of-experiments approach (e.g., randomly surveying many reaction compositions) or have modulated components one at a time, we chose to simultaneously vary fuel and Mg^2+^ concentrations. We performed TX-TL reactions consisting of fuel versus Mg^2+^ panels, where the former was varied between 0-45 mM, the latter was simultaneously varied between 0-10 mM, and all other reaction components were kept the same^14^ (see **Materials and Methods** for more details) (**Figure 1A**). As the preparation of cell lysate varies significantly from lab to lab and is difficult to keep consistent^18^, we performed experiments in two cell lysate batches – prepared by different lab members and using different lysis methods, among other differences – to see if these chemical factors acted consistently.

**Figure 1:**
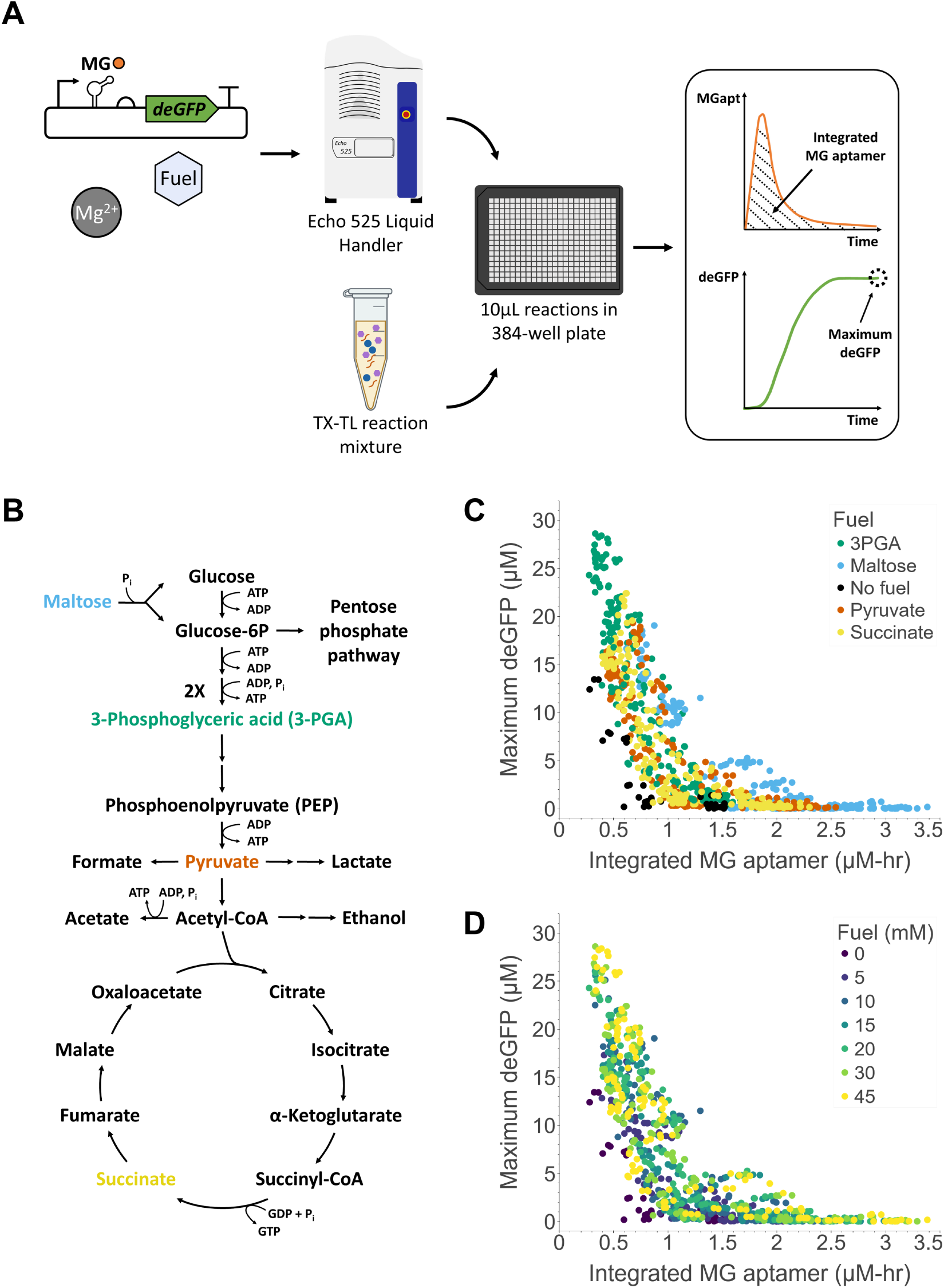
A TX-TL trade-off across fuel sources and concentrations. **(A)** Overview of experimental workflow. **(B)** Simplified map of *E. coli* core metabolism with NTP consumption and regeneration. A simplified glycolysis, TCA cycle, and interacting branch of the Pentose Phosphate Pathway are shown. Consumption or regeneration of other cofactors – such as NADH, FADH, QH2, H_2_O, and CO_2_ – is omitted for simplicity. **(C)** and **(D)** show maximum deGFP values versus integrated MG aptamer values, colored by either (C) fuel type or (D) fuel concentration. Each point represents one of three replicates of a particular set of fuel and Mg^2+^ concentrations. The data shown reflect experiments performed in two batches of cell lysate, Batch 1 and Batch 2 (see Methods and Materials for details), and all experiments were performed using 5 nM DNA.

For fuel molecules, we focused on four fuel metabolites with varying degrees of energy regeneration and waste generation capacity that spanned central carbon metabolism to gain insight into how different fuel molecules affect TX-TL through their effect on metabolism. We chose four central carbon metabolites: 3-phosphoglyceric acid (3PGA), maltose, pyruvate, and succinate (**Figure 1B**). We targeted 3PGA – whose conversion to pyruvate regenerates ATP during glycolysis – because it is one of the standard molecules^14^ used for energy regeneration in cell lysate-based TX-TL systems (the others being phosphoenolpyruvate (PEP) and pyruvate). We chose pyruvate because it is located downstream of 3PGA in the glycolysis pathway and can also participate in energy regeneration, albeit without contributing potential waste molecule inorganic phosphate (Pᵢ) to the system^21^. We selected maltose, which consumes Pᵢ and outputs a glucose molecule and glucose-6-phosphate (G6P) molecule, because it has been used previously to improve energy regeneration by recycling Pᵢ ^34^. However, as one maltose molecule can be used to generate two G6P molecules, and thus four 3PGA molecules, maltose produces more candidate waste molecules per molecule of fuel compared to the other three fuels. These candidate waste molecules include acetate, ethanol, lactate, and formate (and their conjugate organic acids), which may inhibit TX-TL either directly or indirectly, by reducing pH too low for optimal enzyme activity^35–37^. Finally, we considered succinate, whose conversion from succinyl-CoA regenerates GTP. As the succinyl-CoA to succinate reaction is reversible, we believed the addition of succinate could slow down GTP regeneration by favoring the reverse reaction, thereby reducing the overall rate of energy regeneration to a level where energy could be used more efficiently by TX-TL.

TX-TL reactions were supplied with one of two different batches of cell lysate along with plasmid DNA encoding the sequence for the expression of MG aptamer, which fluoresces upon binding to malachite green dye, and deGFP, under a P_OR1OR2_ promoter. For each reaction condition, the total integrated area under the MG aptamer fluorescence curve and maximum deGFP were calculated as measurements of transcription and translation, respectively. The results of these experiments are shown in **Figure 1**.

By plotting maximum deGFP versus integrated MG aptamer for all fuel sources across different fuel and Mg^2+^ concentrations, we observed a trade-off between optimizing maximum deGFP and optimizing integrated MG aptamer (**Figure 1C, D**). The same trade-off curve was observed across all fuel sources, and points corresponding to systems with no added fuel also fell along the same curve (**Figure 1C**). Furthermore, when all fuels are treated equally, a system’s position along the trade-off curve was mostly unaffected by the concentration of fuel (**Figure 1D**, bootstrapped Spearman’s rho correlation of 0.22 ± 0.05). Control experiments indicated that the fluorescence of deGFP was robust across different the 3PGA and Mg^2+^ concentrations used in these experiments, and that MG aptamer fluorescence was robust to changes in Mg^2+^ and pH (**Figure S1A, C**). While deGFP fluorescence was sensitive to pH, it was robust to Mg^2+^ in the relevant range of 0-10 mM and to pH across the estimated range of 6.5-7^22,23^ (**Figure S1B**). As there was no observed trend of decreasing deGFP (or increasing MG aptamer) fluorescence with increasing fuel concentration, there did not appear to be a fuel concentration-dependent effect on fluorescence, including but not limited to pH. This suggested that expression of MG aptamer and deGFP, rather than their fluorescence alone, were changing across the various fuel and Mg^2+^ concentrations. Overall, these results suggest the presence of a trade-off between transcription and translation for the fuels considered here.

The results of this initial experiment were surprising, because all points fell along the same trade-off curve, regardless of fuel type or lysate batch. These results seem to suggest that while these TX-TL systems with different fuel types and concentrations exhibit variability in transcriptional and translational performance, their performance is constrained to a single trade-off curve that could be traversed for a given fuel type and desired performance.

### Variability along the trade-off curve is controlled by Mg^2+^ concentration and lysate batch

To investigate which factors controlled a system’s position along the trade-off curve, we colored the data in **Figure 1** by Mg^2+^ concentration and cell lysate batch, the results of which are shown in **Figure 2**. We found that when considering all fuels equally, as the concentration of added Mg^2+^ increases, a TX-TL system generally shifts from a translation-optimized regime to a transcription-optimized regime (**Figure 2A**, bootstrapped Spearman’s rho correlation of 0.63 ± 0.02). We also found that while data for the two batches of lysate fell along the same curve, variability along the curve could be further explained by lysate batch (**Figure 2B**).

**Figure 2:**
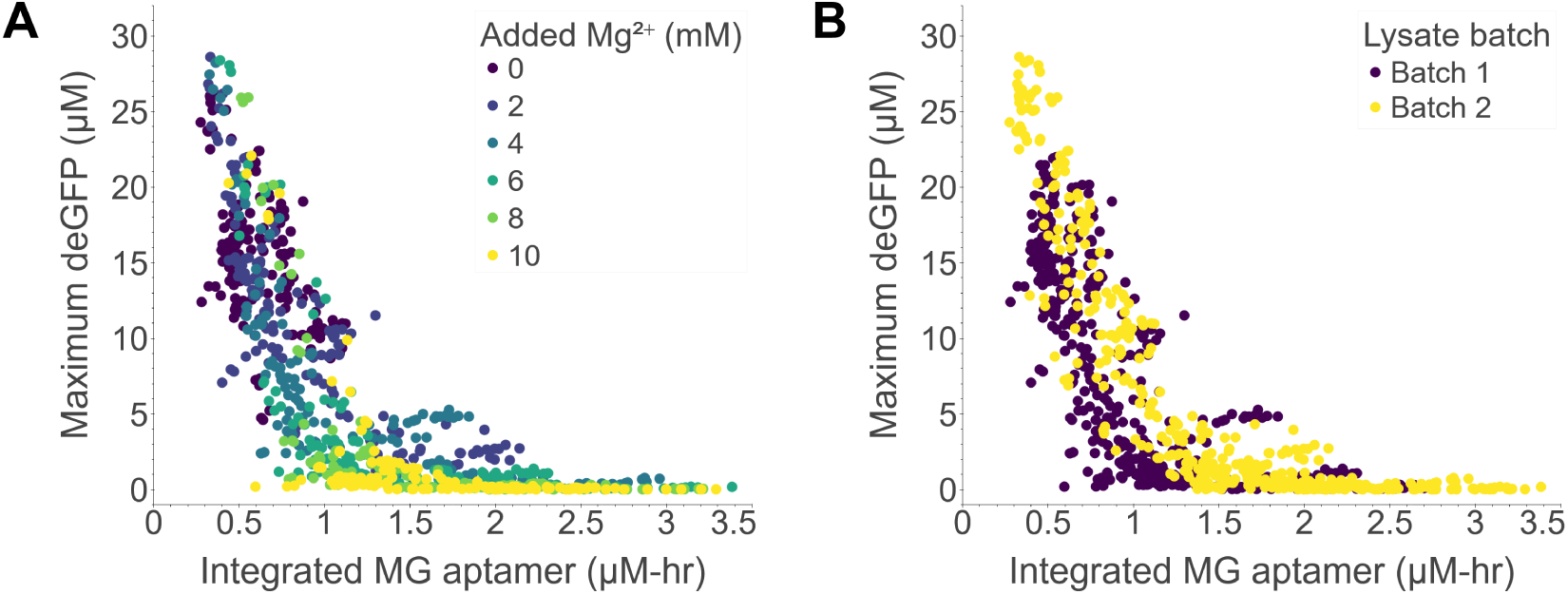
Effects of varying Mg^2+^ concentration and lysate batch on the TX-TL trade-off. These panels show the same data from Figures 1C and 1D colored by **(A)** the concentration of added Mg^2+^ (in addition to the Mg^2+^ that is already in the cell lysate from its preparation in S30 buffer) or **(B)** lysate batch. Each point represents one of three replicates of a particular set of fuel and Mg^2+^ concentrations. The data shown reflect experiments performed in two batches of cell lysate, Batch 1 and Batch 2, in systems using 5 nM DNA.

As the Mg^2+^ trend is present across a variety of fuel sources (**Figure S2**), our results suggest that Mg^2+^ concentration can be tuned for a given set of cell lysate batch, fuel type, and fuel concentration to reduce variability, improve predictability, or design TX-TL systems optimized for performance, including optimizing transcription, translation, or some combination of both. As Mg^2+^ is an essential cofactor for many proteins, these results also suggest the TX-TL trade-off originates in part from Mg^2+^-dependent regulation of TX-TL and/or metabolism machinery.

Inspecting **Figure 2A** more closely revealed that the Mg^2+^ trend was not completely consistent; plotting the trade-off separately for each fuel source revealed that a system’s position along the trade-off curve was less correlated with Mg^2+^ in 3PGA-fueled systems than in systems using other fuels (**Figure S2A**, see figure caption for Spearman’s rho correlation values). Specifically, while TX-TL systems fueled by pyruvate, succinate, and maltose were able to optimize translation at 0 mM added Mg^2+^ and optimize transcription at 10 mM added Mg^2+^, some 3PGA-fueled systems were able to achieve high deGFP yields at high concentrations of added Mg^2+^.

### In 3PGA-fueled systems, increasing fuel pushes a system from a TX-optimized to a TL-optimized regime

To understand why 3PGA-fueled systems deviated from the Mg^2+^ trend, we plotted the data from **Figure 1D** separately for each fuel type, the results of which are shown in **Figure 3**. While systems fueled by maltose, pyruvate, and succinate displayed a weak trend along the trade-off curve with increasing fuel concentration (**Figure 3B-D**, bootstrapped Spearman’s rho correlations of 0.33 ± 0.09, 0.14 ± 0.07, and 0.27 ± 0.07, respectively), we found that in 3PGA-fueled systems, increasing fuel concentration was more strongly correlated with a system’s position along the trade-off curve (**Figure 3A**, bootstrapped Spearman’s rho correlations of 0.55 ± 0.05). Specifically, increasing 3PGA concentration generally shifted a TX-TL system from a transcription-optimized regime to a translation-optimized regime. This phenomenon was also observed in 3PGA-fueled TX-TL systems where the volume fraction of cell lysate in a TX-TL reaction was increased or decreased by 25% (**Figure S3**), with the trend becoming stronger as the lysate fraction was increased (**Figure S3D**), thus extending the relevance of the trend to TX-TL systems using alternative cell lysate volume fractions^25^.

**Figure 3:**
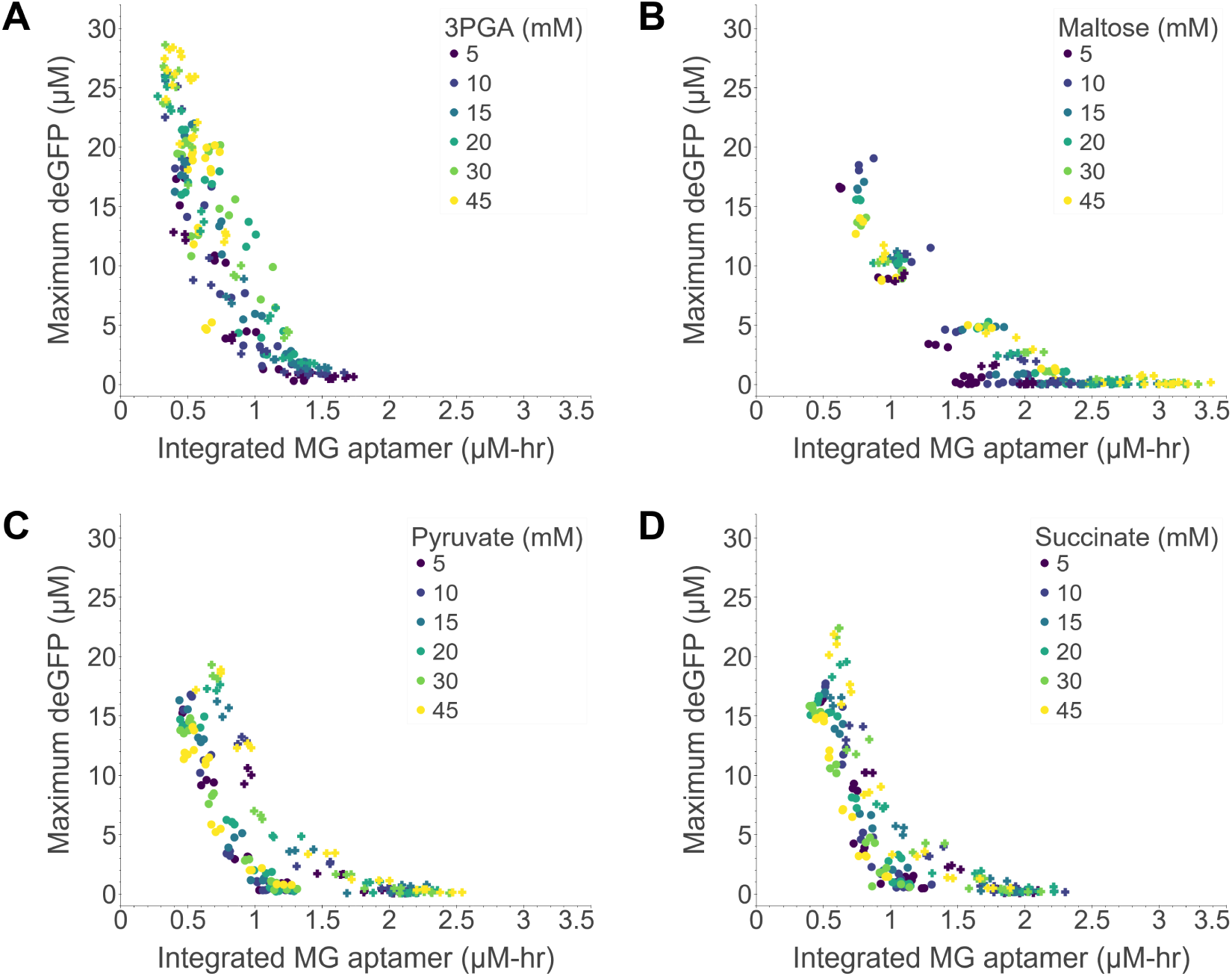
Effects of varying fuel concentration on the TX-TL trade-off by fuel type. The same data are shown as in Figures 1C and 1D with points colored by fuel concentration. Each point is one of three replicates of a set of fuel and Mg^2+^ concentrations, where fuel is either **(A)** 3PGA, **(B)** maltose, **(C)** pyruvate, or **(D)** succinate. Experiments were performed in two batches of cell lysate, Batch 1 (**o**) and Batch 2 (**+**), in systems using 5 nM DNA.

We next sought to confirm whether the TX-TL trade-off and 3PGA trend extended to systems using different reporter plasmids. The results of these 3PGA versus Mg^2+^ panels are shown in **Figure 4**. We first switched the promoter from P_OR1OR2_ to P_T7_, where we found that the TX-TL trade-off and 3PGA trend were present across P_T7_-driven TX-TL systems (**Figure 4B**, bootstrapped Spearman’s rho correlation of 0.58 ± 0.06), including in a system where the order of fluorophores was flipped (**Figure S4**), just as they were in P_OR1OR2_-driven systems (**Figure 4A**, bootstrapped Spearman’s rho correlation of 0.69 ± 0.05). When the promoter was changed to P_Tet_, however, the TX-TL trade-off curve was no longer visible, and the 3PGA trend was weaker (**Figure 4C**, bootstrapped Spearman’s rho correlation of 0.44 ± 0.08). In P_Tet_-driven TX-TL systems, it appeared that only some portion of the data, corresponding to about 15-30 mM 3PGA, fell along the common trade-off curve (**Figure S5**). The data corresponding to 0-5 mM 3PGA did not give the highest MG aptamer yields as in **Figure 4A** and **Figure 4B**, and at high 3PGA concentrations, the left edge of the curve dipped, reflecting decreased deGFP yields at high 3PGA concentrations. To determine if this was a behavior unique to this promoter, we also performed a 3PGA versus Mg^2+^ panel in systems expressing F30-Pepper aptamer and mTurquoise2 under a P_Tet_ promoter (**Figure 4D**). Here the trade-off curve and 3PGA trend were present as before (bootstrapped Spearman’s rho correlation of 0.51 ± 0.06), although as in systems using the P_Tet_-MG aptamer-deGFP plasmid, there was a decrease in protein yields at higher 3PGA concentrations.

**Figure 4:**
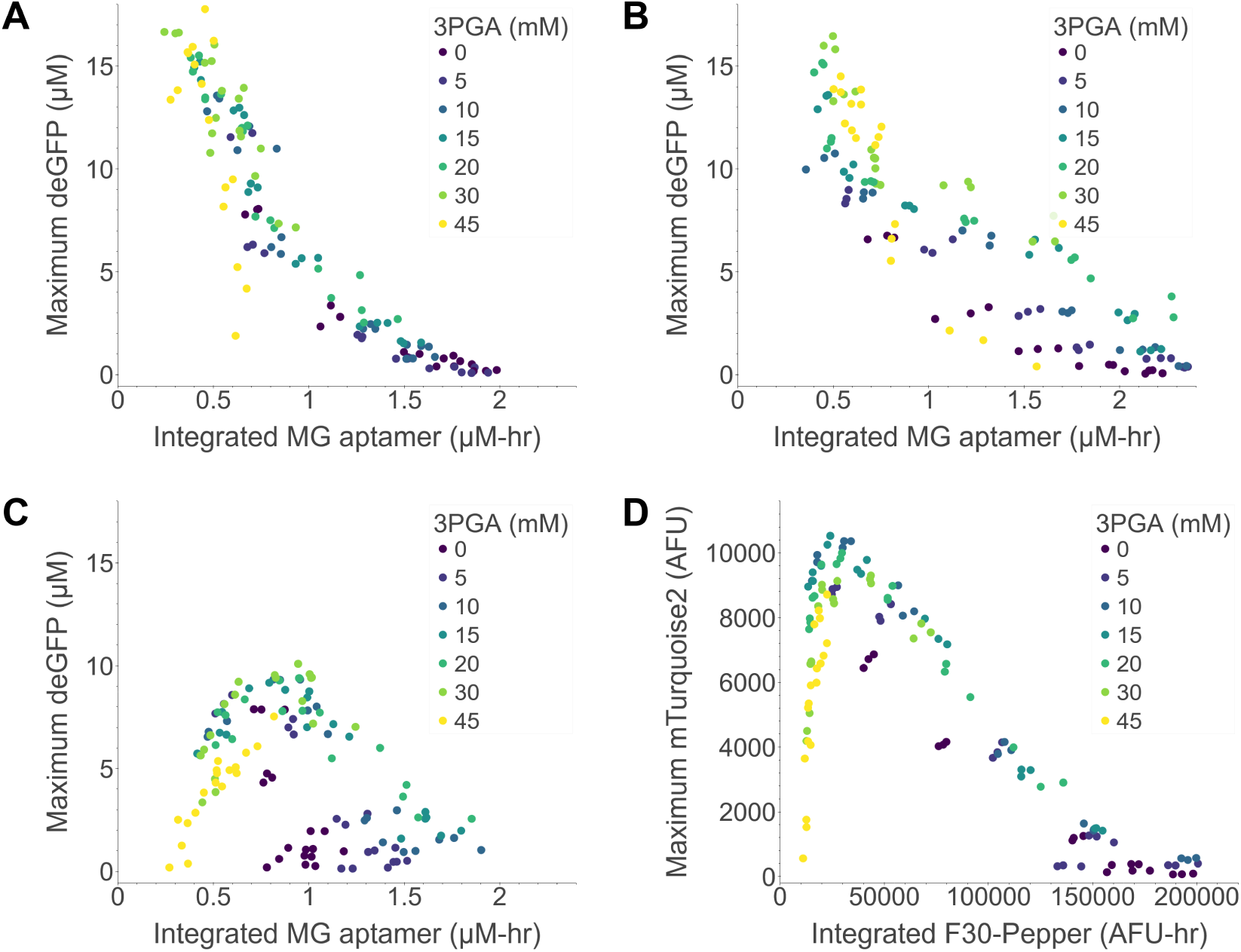
Effects of 3PGA concentration on the TX-TL trade-off across different reporter plasmids. Shown are trade-off curves corresponding to **(A)** P_OR1OR2_-MG aptamer-deGFP, **(B)** P_T7_-MG aptamer-deGFP, **(C)** P_Tet_-MG aptamer-deGFP, and **(D)** P_Tet_-F30 Pepper-mTurquoise2. P_OR1OR2_, a strong promoter and P_Tet_, a medium strength promoter, are constitutive promoters, and P_T7_ is a strong non-*E. coli* promoter expressed in TX-TL reactions supplemented with 10 μM T7 RNA polymerase (see Methods and Materials for sequences). Each point represents one of three replicates of a set of 3PGA and Mg^2+^ concentrations and is colored by 3PGA concentration. The data shown in **(A)**, **(B)**, and **(C)** reflect experiments performed in cell lysate Batch 3, while the data in **(D)** reflect experiments performed in cell lysate Batch 2 (see Methods and Materials for details). All experiments were performed using 5 nM DNA.

While it is not clear what caused this change in the shape of the trade-off curve in P_Tet_-driven systems, this finding is consistent with the hypothesis that the TX-TL trade-off is indicative of a fundamental trade-off in fuel and energy metabolism. In such systems, weaker promoters would not be able to maximize rates of transcription and translation output as effectively, resulting in less deGFP being expressed before energy is dissipated by metabolic processes competing for the same energy pool and/or before translation-inhibiting waste is accumulated, compared to TX-TL under strong promoters. This could explain why deGFP yields tend to drop at high 3PGA concentrations in **Figure 4C** and **Figure 4D**, although this explanation is insufficient to explain the shape of the data in **Figure 4C**. While more promoters need to be tested to determine the generalizability of the TX-TL trade-off to systems using different genetic parts, the presence of the trade-off in P_T7_-driven MG aptamer and deGFP expression was notable, considering that the T7 promoter is widely used in applications aimed at maximizing protein expression.

### In 3PGA-fueled systems, the TX-TL trade-off and fuel trend are consistent across DNA concentrations

Having confirmed that the TX-TL trade-off was present in a wide variety of TX-TL systems, we next sought to confirm whether the trade-off curve scaled with DNA concentration, as previous experiments had all been performed with 5 nM DNA. For this set of experiments, we decided to focus on 3PGA-fueled systems, since 3PGA – and its downstream metabolic product phosphoenolpyruvate (PEP) – are the most widely used metabolites for energy regeneration in TX-TL systems, and gaining insight into the TX-TL trade-off in 3PGA-fueled systems would have greater relevance for studies aimed at characterizing and improving these systems.

We performed 3PGA versus Mg^2+^ panels at DNA concentrations of 1, 2.5, 5, 7.5, and 10 nM, the results of which are shown in **Figure 5**. We found that the TX-TL trade-off curve’s location shifted and range of MG aptamer and deGFP concentrations increased as DNA concentration was increased, with a curve first becoming apparent at 2.5 nM DNA and then becoming more apparent as the DNA concentration was increased (**Figure 5A**).

**Figure 5:**
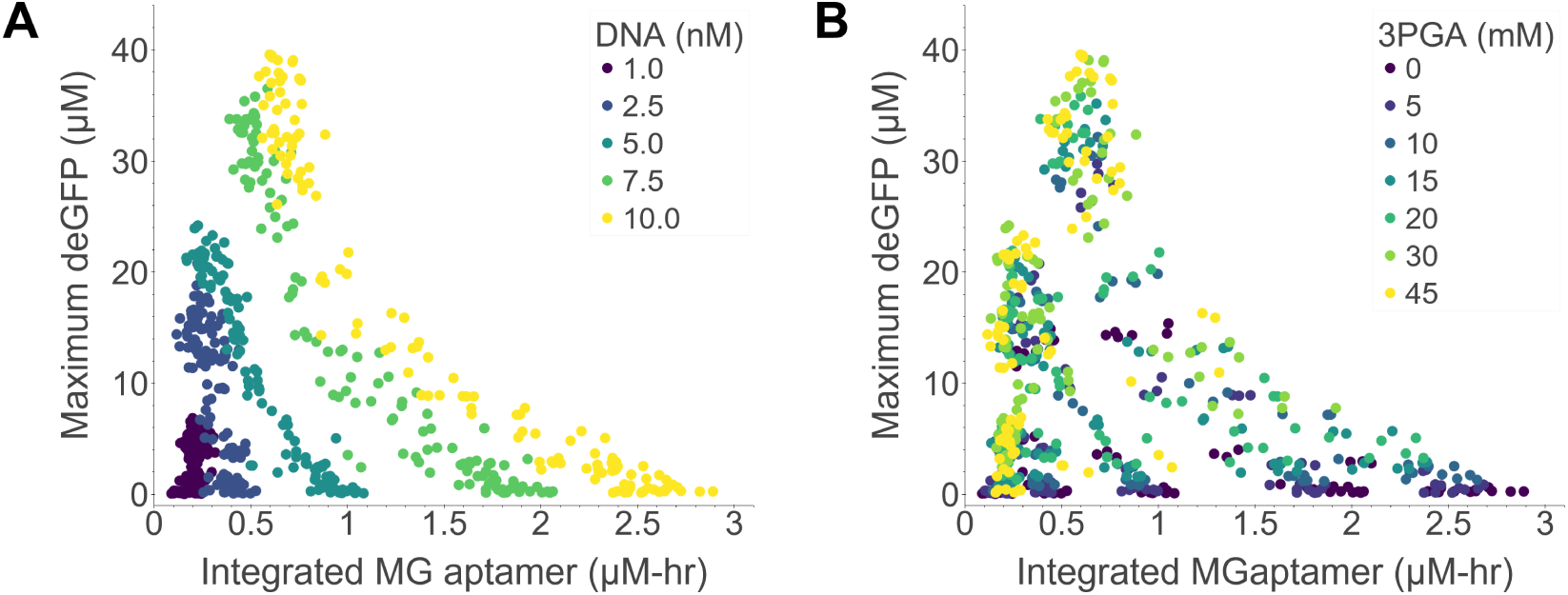
Effects of varying DNA concentration on the TX-TL trade-off. **(A)** Each point represents one of three replicates of a particular set of 3PGA and Mg^2+^ concentrations, where points are colored by DNA concentrations ranging from 1-10 nM. **(B)** The same data are shown as in (A), albeit with the points colored by 3PGA concentration rather than DNA concentration. The data shown reflect experiments performed in cell lysate Batch 2 (see Methods and Materials for details).

As in the 5 nM DNA cases discussed in the previous section, the trend in shifting a system from a transcription-optimized regime to a translation-optimized regime at higher 3PGA concentrations was also consistent in the trade-off curves at DNA concentrations higher than 1 nM (**Figure 5B**, bootstrapped Spearman’s rho correlations of 0.46 ± 0.08, 0.60 ± 0.07, 0.63 ± 0.06, 0.66 ± 0.06, and 0.69 ± 0.06, for DNA concentrations of 1 nM, 2.5 nM, 5 nM, 7.5 nM, and 10 nM, respectively). Preliminary analyses also revealed that transcription scaled linearly with increasing DNA concentration in systems along lines that intersected the trade-off curves (**Figure S6**). This finding is particularly relevant for transcription-only systems, where this linear relationship may make it easier to predict the performance of transcriptional circuits based on the concentrations of DNA species. More experiments are needed to determine whether this linear transcriptional scaling is a general phenomenon in TX-TL systems.

### The trade-off arises at the translational level in 3PGA-fueled systems

Having confirmed that there was a TX-TL trade-off that controlled a system’s transcriptional and translational capacity as a function of Mg^2+^ concentration broadly, and additionally as a function of 3PGA concentration when that fuel source was used, we next sought to gain more insight into the cause of the trade-off in 3PGA-fueled systems. It was not clear from our previous data whether conditions that optimized transcription versus conditions that optimized translation were simply mutually exclusive, in that optimal Mg^2+^ and 3PGA concentrations were simply different for transcription versus translation. An alternative explanation was that either or both of the processes, transcription and translation, had an inhibitory effect on the other process; in the case where only one process inhibited the other, the first process could be inhibited by the reaction environment. A third possible explanation was that transcription and translation were processes competing for the same resources but that one process had a preferred regime for optimization. In this scenario, the energy-dominating process would have an optimal regime, and the other process would only be optimal where the energy-dominating process was strictly not optimal, thereby leaving more resources for the non-dominating process. Combinations of these explanations, or others, could also explain the observed trade-off.

To gain insight into the individual processes of transcription and translation, we decoupled the TX-TL system into a TX-only system and a TL-only system. In the TX-only system, tetracycline was added to the system to block translation via inhibition of translation initiation through its binding to the 30S subunit of the ribosome^38^. In the TL-only system, mRNA was added to the system instead of DNA. The results of these experiments are shown in **Figure 6**.

**Figure 6:**
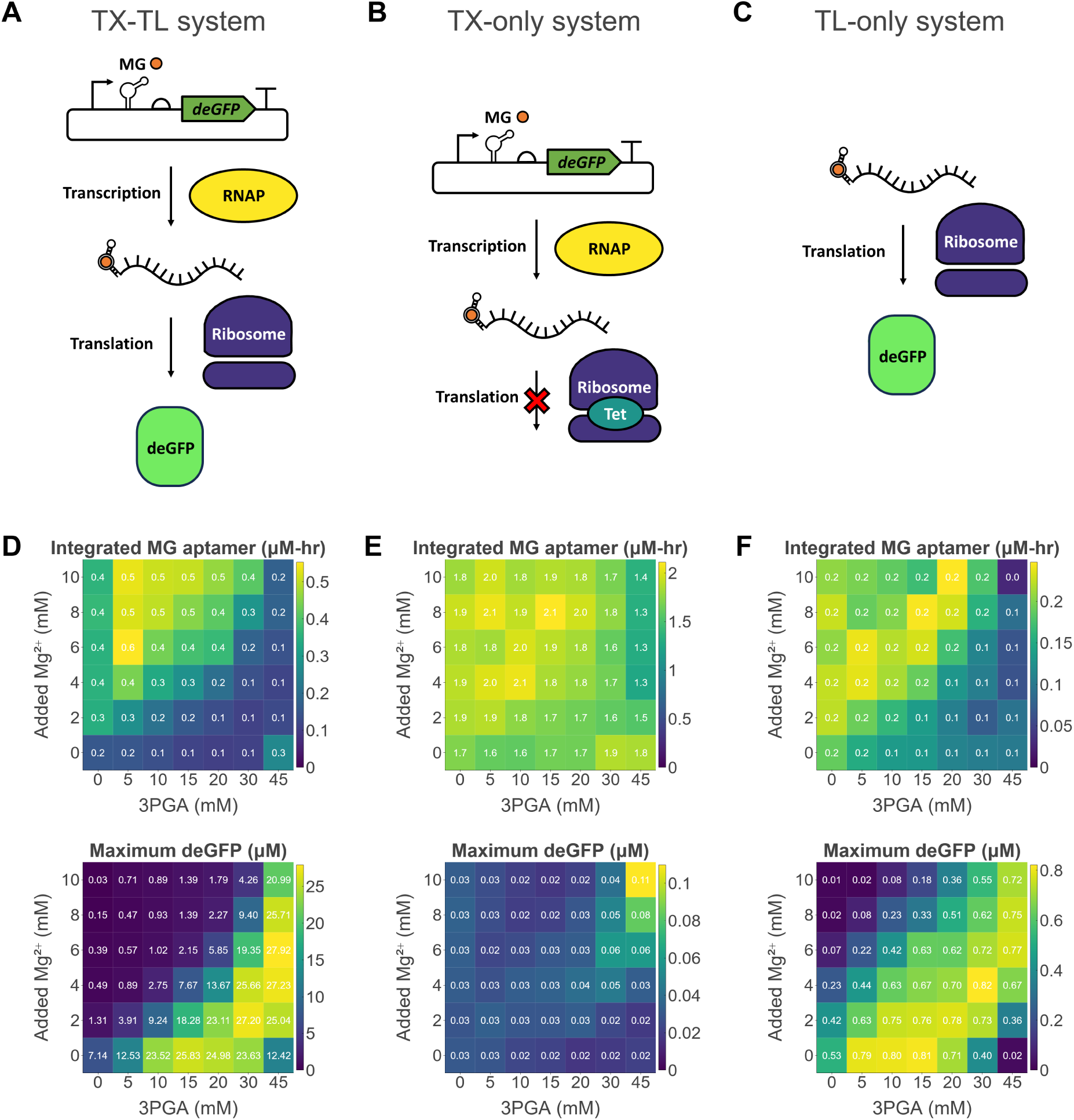
Decoupling of transcription (TX) and translation (TL). **(A)**, **(B)**, and **(C)** show experimental schemes corresponding to the data shown in **(D)**, **(E)**, and **(F)**, respectively. **(A)** Simplified view of TX-TL: P_OR1OR2_-MG aptamer-deGFP DNA is translated by *E. coli* RNA polymerase to mRNA, which fluoresces upon binding to Malachite Green (MG) dye. The mRNA is then translated by ribosomes into deGFP protein. **(B)** Simplified view of TX-only system: transcription proceeds as in (A), but translation is inhibited by tetracycline, which binds to ribosomes and inhibits translation initiation. **(C)** Simplified view of TL-only system: MG aptamer-deGFP mRNA, rather than DNA, is added to the system, and the mRNA is translated into deGFP protein. **(D)**, **(E)**, and **(F)** show integrated mRNA (top) and maximum deGFP measurements (bottom) for the (**D)** TX-TL system, **(E)** TX-only system, to which 200 μg/mL tetracycline was added, and **(F)** TL-only system, where 0.3 μM mRNA was added at the beginning of the reaction. All reactions shown used Batch 2 lysate, and the TX-TL and TX-only systems used 5 nM DNA. Each measurement is the average of three replicates for that experimental condition. Colorbars are scaled separately for each plot.

Using heatmaps to display integrated mRNA and maximum deGFP data, we found that in the TX-TL system (**Figure 6A**), the regime of Mg^2+^ and 3PGA concentrations that optimized transcription was distinct and different than the regime that optimized translation (**Figure 6D**). Specifically, optimal transcription preferred low 3PGA and high Mg^2+^ concentrations and optimal translation preferred high 3PGA and low Mg^2+^ concentrations; both findings are consistent with the trade-off curve. However, in the TX-only system (**Figure 6B**), MG aptamer expression was not optimized at low 3PGA and high Mg^2+^ as in the case of **Figure 6D**; rather, it was broadly optimal at 3PGA concentrations ranging from 0-30 mM and 0-10 mM added Mg^2+^ (**Figure 6E**). Overall transcription was also higher, likely due to the abundance of energy in the absence of translation. Meanwhile, in the TL-only system (**Figure 6C**), higher deGFP yields were achieved, as before, at low Mg^2+^ and high 3PGA concentrations, despite reduced mRNA degradation in the regime of high Mg^2+^ and low 3PGA concentrations (**Figure 6F**).

These data suggest that the TX-TL trade-off is caused by a combination of translation optimization at low Mg^2+^ and high 3PGA concentrations, increased MG aptamer stability at high Mg^2+^ and low 3PGA concentrations, and competition for resources in the TX-TL system. In other words, optimal translation occurs where conditions are ideal for that process, and optimal transcription occurs where translation is sub-optimal and leaves more fuel and energy resources for transcription. While these experiments did not shed light on the mechanism governing translation optimization at lower Mg^2+^ and higher 3PGA concentrations, they suggest that the TX-TL trade-off arises at the translational level in 3PGA fueled systems.

### The TX-TL trade-off is absent when TX-TL systems are supplied with no fuel or Mg^2+^ and additional energy

The TX-TL decoupling experiments had implied that the TX-TL trade-off arose, in part, due to competition for fuel and energy resources between transcription and translation. By this hypothesis, TX-TL systems with abundant energy should not fall along the trade-off curve, particularly in a regime of low Mg^2+^ concentration where both transcription and translation were favorable when not limited by a competing process (**Figures 6E, F**). To test this hypothesis, we next performed experiments where TX-TL systems were supplied with an excess of energy. To mitigate the potential effects of waste accumulation that came with adding high levels of central carbon-based fuel sources, we decided against using any central carbon fuel source (i.e. 3PGA, maltose, pyruvate, or succinate) for energy regeneration. As Mg^2+^ also had a translation-inhibition effect, for these experiments, we also decided against adding additional Mg^2+^. Instead, to explore the space of different energy sources and concentrations, we performed a three-dimensional panel of ATP versus GTP versus NTPs, where each solution was added at a final concentration of 0, 5, 10, or 15 mM. The results of these experiments are shown in **Figure 7**.

**Figure 7:**
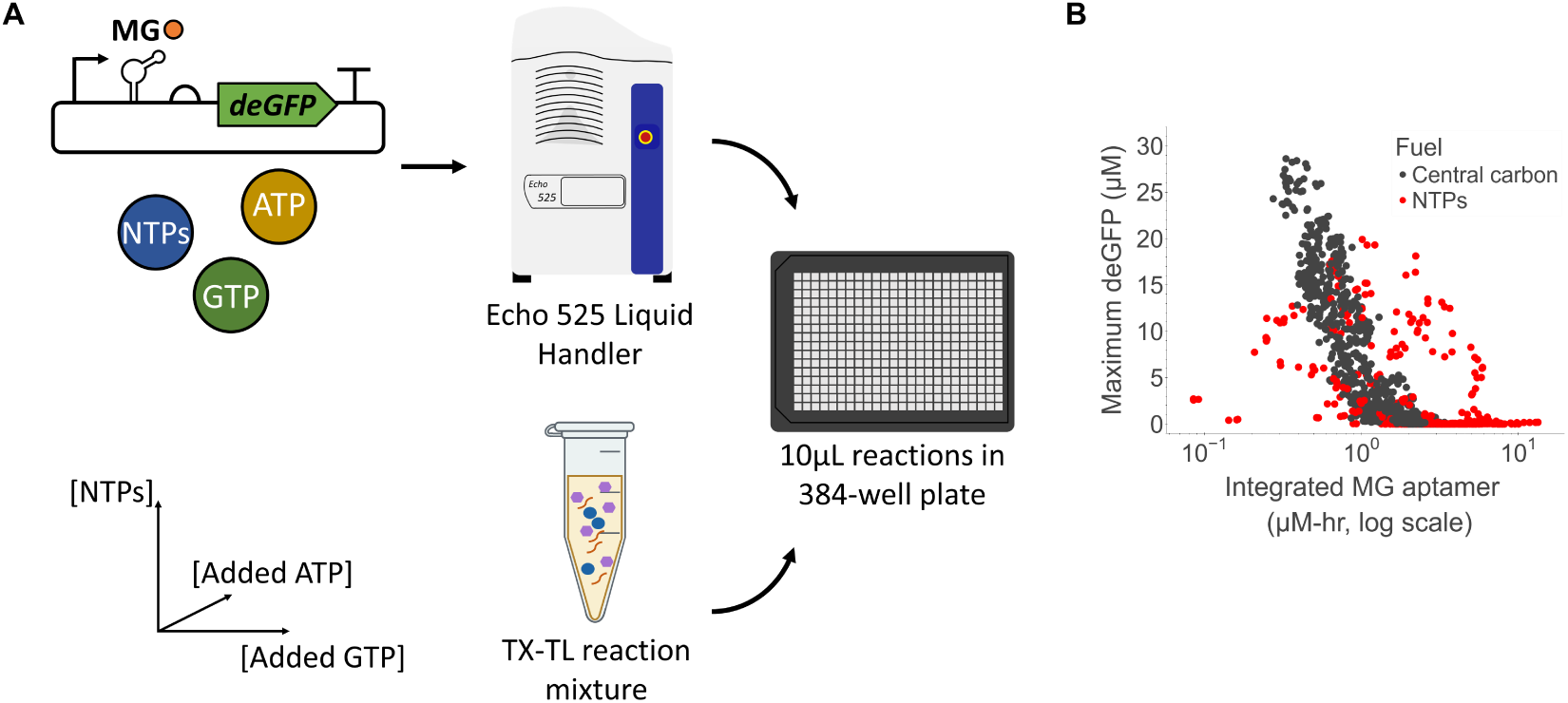
Absent TX-TL trade-off in the absence of fuel or Mg^2+^. **(A)** Overview of experimental workflow. **(B)** Maximum deGFP values versus integrated MG aptamer value. A log scale was chosen for the latter axis to prevent high MG aptamer values from skewing the data in a way that made it difficult to view the original trade-off data. Each point represents one of three replicates of a particular set of NTP, added ATP, and added GTP concentrations, colored by the fuel type. Here, the “central carbon” fuel data is the same as in Figure 1, and those reactions all contain 4.8 mM NTPs. The “NTPs” fuel data reflect experiments where no central fuel or Mg^2+^ has been added. The data shown reflect experiments performed in two batches of cell lysate, Batch 1 and Batch 2 (see Methods and Materials for details), and all experiments were performed using 5 nM DNA.

As expected, TX-TL systems supplied with additional energy molecules and no fuel or no Mg^2+^ did not fall along the previously found trade-off curve corresponding to that DNA concentration (5 nM), and overall, these NTP-fueled systems displayed a weaker trend of decreasing deGFP with increasing integrated MG aptamer (Spearman’s rho correlation of -0.47) compared to central carbon-fueled systems (Spearman’s rho correlation of -0.90). Additionally, NTP-fueled systems could achieve deGFP yields comparable to systems supplied with pyruvate or maltose and integrated MG aptamer yields 3-10X higher than in TX-TL systems supplied with those fuels and Mg^2+^. While there was no clear trend in ATP, GTP, or total NTP concentration for the data shown in **Figure 7B**, generally, as the concentration of NTPs increased, integrated MG aptamer values increased, and deGFP values decreased (**Figure S7**), which is consistent with a previous finding^28^. The decrease in deGFP yields despite the abundance of energy was likely due to NTP chelation of Mg^2+^, which is necessary for ribosomes and other translation machinery. Although Mg^2+^ was not directly added to the reactions, it was present at a concentration of around 4 mM due to the presence of S30B buffer (used in the preparation of cell lysate) in the reaction. Our results suggest that fuel-supplied TX-TL systems are indeed energy-limited, and that the TX-TL trade-off curve exists in part due to competition for a limited pool of energy for transcription and translation.

The data revealed three additional insights. First, systems with no fuel or no Mg^2+^ and high NTPs often exhibited unusual transcriptional and translational dynamics (**Figure S8**). Some systems exhibit long time delays in transcription (**Figure S8A**) or translation (**Figures S8E-F**). Other systems exhibited biphasic expression (**Figures S8B, D**) or steady state transcription rates as opposed to the typical transcription pulse (**Figures S8C**).

Second, although NTP-fueled TX-TL systems did not achieve protein yields as high as those achieved in systems with 3PGA, supplying systems with 10 mM NTPs helped achieve integrated MG aptamer and deGFP yields comparable to systems with about 5 mM NTPs and 10 mM 3PGA, the latter of which should have regenerated 10 mM ATP. This suggests that while energy regeneration is likely occurring in 3PGA-fueled systems, energy being regenerated is not being used efficiently. A large fraction of that regenerated energy, perhaps the majority, is likely fueling metabolic processes irrelevant to transcription or translation.

Third, if the TX-TL trade-off curve was indicative of a greater trade-off between efficient energy regeneration and waste minimization, any potential translation-inhibiting waste product was likely not produced as a result of total transcription, since relatively high maximum deGFP yields were achieved with high Integrated MG aptamer values in the case of NTP-fueled systems (**Figure 7B**). The waste product also did not appear to be solely a product of fast transcription; while there was negative correlation between the maximum rate of transcription and maximum deGFP in central carbon-fueled systems (Spearman’s rho correlation of -0.63), there was a positive correlation between the maximum rate of transcription and maximum deGFP in NTP-fueled systems (**Figure S9**). These data suggest that the waste was likely derived from the metabolism of central carbon fuels. Mg^2+^ versus fuel panels performed with all fuels in the regime of low DNA concentration (i.e. 1 nM) showed that in a transcription-limited regime, central carbon-fueled TX-TL systems indeed preferred low fuel concentrations for high deGFP yields (**Figures S10-S13**). These results suggest that these systems were not able to sufficiently utilize central carbon-fueled energy regeneration and thus could not rapidly make protein before waste accumulated, compared to systems with 5 nM DNA, thereby supporting the idea that waste products were, in part, central carbon metabolism-derived.

## Discussion

An improved understanding of cell-free TX-TL systems is essential for scaling up, expanding the scope of, and improving the performance of TX-TL applications in academic and industrial settings. By using metabolic perturbations as a tool to gain insight into the behavior of TX-TL systems in different metabolic regimes, this study has shed light on how Mg^2+^ and fuel contribute to TX-TL variability and how these two components and NTPs affect TX-TL performance. Our results indicate the presence of a trade-off between transcription and translation in *E. coli*-based TX-TL systems and show clear roles for how Mg^2+^, fuel, cell lysate batch, and other factors affect transcription and translation and give rise to TX-TL variability.

Armed with these insights, other users of TX-TL systems aiming to reduce variability and standardize their systems may consider calibrating the concentrations of the relevant small molecules (fuel, Mg^2+^, and/or DNA) for their application so that TX-TL curves for different systems have similar locations, shapes, and other characteristics. By characterizing different batches of cell lysate by the shape and location of their trade-off curves, for example, we can understand their respective transcriptional and translational limits. Beyond reducing variability and improving standardization, this study also has relevance for efforts aimed at increasing the predictability and performance of TX-TL systems for desired uses. A user can optimize transcription versus translation optimization in different batches of cell lysate by choosing the appropriate set of fuel types, fuel concentrations, and Mg^2+^ concentrations for a particular genetic program, whether that be a complex transcriptional circuit or a simple program for maximum protein yield. The data included with this paper provide hundreds of unique reaction conditions that can serve as a starting point for exploring conditions that are optimized for high protein yields, long reaction lifetime, specific temporal dynamics, or other desired behaviors.

In addition to providing insight into design considerations for TX-TL systems, the TX-TL trade-off curve, initially a surprising finding, has also provided a simple yet powerful way of gaining insight into fundamental trade-offs in fuel and energy metabolism. Taken together, our results suggest that the TX-TL trade-off is present in energy-limited systems and is indicative of a trade-off between energy regeneration and waste accumulation that results from fuel metabolism. In other words, while fuel molecules enable energy regeneration, their own metabolism also results in waste accumulation (through the generation of immediate or downstream products) that affects TX-TL processes. While the identities of the waste molecules remain unknown, our results suggest that in 3PGA-fueled systems, the waste is partially 3PGA-derived and affects translation. This insight, along with results from the TX-TL decoupling experiments, have helped us to create a new model of cell-free metabolism coupled to cell-free TX-TL (**Figure 8**). While this model has not been validated computationally, doing so may enable improved prediction of TX-TL performance, perhaps even in non-lysate-based TX-TL systems (e.g. PURE-based systems^16^), and this remains an active area of future work.

**Figure 8:**
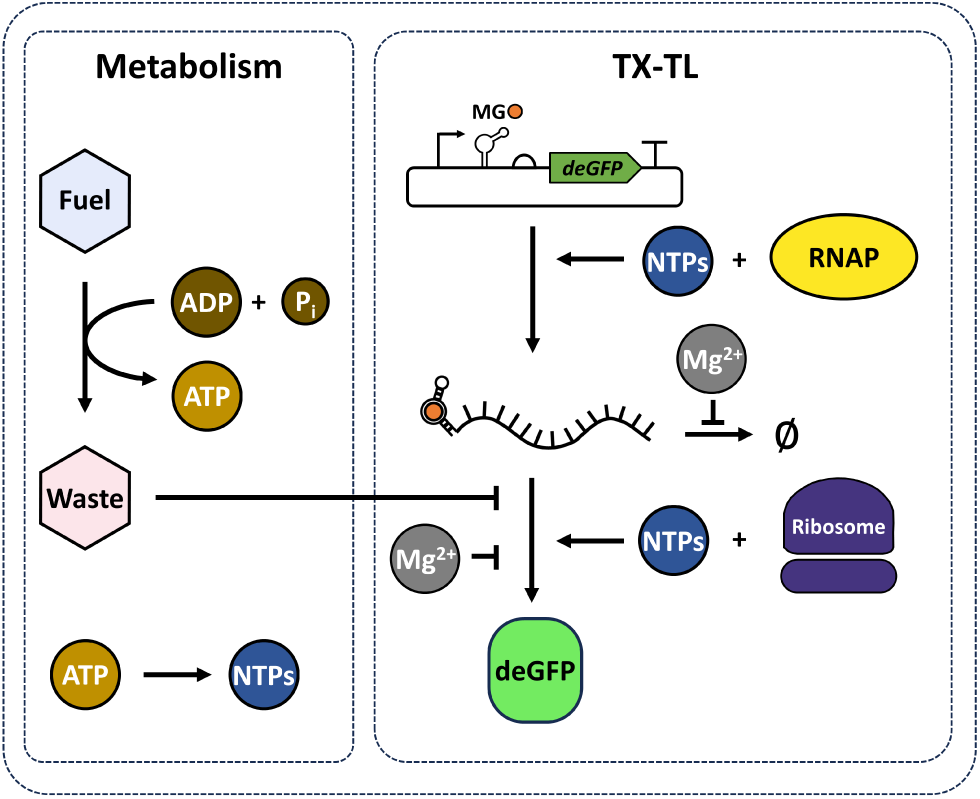
A new model for cell-free metabolism coupled to cell-free TX-TL. The conversion of fuel to waste regenerates energy, which can power transcription and translation, but the accumulation of waste inhibits translation. The fuel-energy-waste paradigm is derived from an existing model^33^. This model’s key contribution is providing roles for Mg^2+^ via its inhibition of mRNA degradation and translation.

In revealing the fundamental metabolic trade-offs in TX-TL systems, this work has also shed light on why previous studies aimed at increasing protein yield have been largely unsuccessful. Our data suggest that low Mg^2+^ concentrations, strong promoters, and high DNA concentrations – and high 3PGA concentrations where 3PGA is used – are necessary for high deGFP yields. However, it is not possible to remove more Mg^2+^ from the system without compromising the function of proteins in the cell lysate, nor is adding more fuel possible, as the inability of TX-TL systems to take advantage of faster energy regeneration will make the faster waste accumulation an issue at high fuel concentrations. Adding too much DNA may skew the systems towards transcription rather than translation. Previous studies aimed at improving deGFP yields have focused on other mechanisms of energy regeneration and waste mitigation, such as using combinations of fuels like 3PGA and maltose^34^, and such approaches and others, including at the level of plasmid design, will be necessary for vastly improving protein yields in lysate-based TX-TL systems.

Overall, by elucidating the effects of cell-free metabolism on TX-TL systems, this work lays a foundation for improving TX-TL performance, lifetime, standardization, and prediction. Ultimately, it represents one of many such steps that will be necessary for further improvement of TX-TL systems, whether that be for applications in biomanufacturing, gaining insight into natural biological systems, or the development of synthetic life.

## Materials and Methods

### Preparation of cell lysate

Cell lysate was prepared from *E. coli* BL21 Rosetta2 cells using the protocol established by Sun and coworkers^14^ with some modifications. Briefly, cells were grown sequentially in 6 mL, 100 mL, and 1 L cultures, where smaller cultures were used to inoculate larger ones. Once 6 L of culture had grown to roughly an OD600 of 2, cells were repeatedly centrifuged and washed with S30B buffer by re-suspension. Cells were then re-suspended in S30B buffer and subjected to cell lysis.

For the Batch 2 and Batch 3 lysates, cells were resuspended at 1.4 g cells per mL of S30B buffer and lysed by a French cell press at a pressure of 640 psi. For the Batch 1 lysate, cells were resuspended to 1 g cells per mL of S30B buffer and lysed by sonication at a probe frequency of 20 kHz and an amplitude of 50%. Cells were sonicated at 10-second ON/OFF intervals for a total of 60 seconds ON (6 ON/OFF cycles), resulting in approximately 300 Joules of energy delivered. After both methods of cell lysis, 3 μL of 1 M DTT was added per mL of cell lysate, and the lysates were then centrifuged to remove cell debris. All batches of lysate were then subjected to a 1-hour run-off reaction at 37°C, after which they were subjected to centrifugation to remove additional debris.

The Batch 2 and Batch 3 lysates were additionally subjected to dialysis at 4°C via Slide-a-Lyzer G3 cassettes (10kDa, 15mL) placed in a beaker containing S30B buffer and a magnetic stir bar to facilitate diffusion. Excess air was removed using a syringe. The Batch 2 lysate was dialyzed for 4 hours (where cassettes were transferred to a beaker of fresh S30B at the 2-hour mark) at 140 rpm. The Batch 3 lysate was dialyzed for 2 hours (where cassettes were transferred to a beaker of fresh S30B at the 35-minute mark) at 160 rpm.

Finally, both batches of cell lysate were aliquoted, flash-frozen in liquid nitrogen, and stored at - 80°C until further use. Bradford assays determined that the final protein concentrations for the cell lysates were 30-40 mg/mL.

### Cloning plasmids used to express deGFP

The P_OR1OR2_-MG aptamer-deGFP plasmid used to express deGFP was originally prepared by Dan Siegal and modified by Zoila Jurado via Site-Directed Mutagenesis to change the T7 promoter to a P_OR1OR2_ promoter and the T7 terminator to a T500 terminator. This plasmid is P_OR1OR2_-MG aptamer-UTR1-deGFP-T500. The P_Tet_-MG aptamer-UTR1-deGFP-T500 plasmid was subsequently constructed using Site-Directed Mutagenesis to change the P_OR1OR2_ promoter to a P_Tet_ promoter, to make expression results as comparable as possible. The P_T7_-MG aptamer-UTR1-deGFP-T7terminator plasmid was originally prepared by Dan Siegal and modified using Site-Directed Mutagenesis to add a few extra bases to the T7 promoter. The P_Tet_-F30Pepper-UTR1-mTurquoise2-ECK120029600 plasmid was constructed using Golden Gate Assembly of CIDAR MoClo part plasmids. All plasmid sequences are in the Supplementary Information section.

### Preparation of template DNA plasmids

In experiments where the P_OR1OR2_-MG aptamer-UTR1-deGFP-T500 plasmid was used, template DNA for the cell-free reactions was prepared from *E. coli* KL740 cells (purchased from Arbor Biosciences) that had been transformed with the plasmid described above. Cells were grown overnight at 30°C, in Lysogeny Broth (LB) medium supplemented with 100 µg/mL carbenicillin, and either Mini-prepped (using a Qiagen Mini-prep kit) or Midi-prepped (using a Macharey-Nagel NucleoBond Xtra Midi kit) the next day using the appropriate protocols. When Mini-prepping, DNA was eluted with nuclease-free water; when Midi-prepping, after the concentration step, DNA was eluted with IDTE Buffer (10 mM Tris-HCl, 0.1 mM EDTA).

The P_Tet_-MG aptamer-UTR1-deGFP-T500, P_T7_-MG aptamer-UTR1-deGFP-T7terminator, and P_Tet_-F30Pepper-UTR1-mTurquoise2-ECK120029600 plasmids were prepared from cells grown overnight at 37°C, in Lysogeny Broth (LB) medium supplemented with 100 µg/mL carbenicillin (for the deGFP plasmids) or 50 µg/mL kanamycin. The cells were Mini-prepped using a Qiagen Mini-prep kit, where DNA was eluted with nuclease-free water.

The P_T7_ -UTR1-deGFP-MG aptamer-T7terminator plasmid was constructed from plasmid pTXTL-T7p14-deGFP (Arbor Biosciences, myTXTL Toolbox 2.0 plasmid collection, now discontinued). The plasmid was modified by the insertion of a Malachite Green aptamer sequenced after the deGFP sequence with the use of the following primers: CGAGGGGATCCC GACTGGCGAGAGCCAGGTAACGAATGGATCCCTTAGGAGATCCGGCTG (ZJQ.A59F, forward primer) and GGGATCCATTCGTTACCTGGCTCTCGCCAGTCGGGATCCCCTCGAGTTAGATCCCGGC (ZJQ.A60R, reverse primer). The template plasmid was first added to a PCR reaction with these primers, using an annealing temperature of 60°C and an elongation time of 2.5 minutes. Next, the PCR reaction was subjected to a 1-hour DpnI digest to remove the template DNA via the addition of 0.25 µL of DpnI enzyme (purchased from New England Biolabs at 20000 units/mL) per 25 µL reaction and subsequent incubation at 37°C for 1 hour. Next, PCR clean-up was performed using the Qiagen Mini-prep kit to extract linear DNA. Finally, the linear DNA was re-circularized by performing a Gibson reaction using NEBuilder® HiFi DNA Assembly Master Mix, after which 0.5 µL of the reaction mix was then used to transform 50 µL of competent cells, which were grown overnight at 37°C on LB agar plates with 100 µg/mL carbenicillin. Transformed colonies were sequence verified for insertion of the Malachite Green sequence and subsequently cultured overnight at 37°C with 100 µg/mL carbenicillin and Mini-prepped for plasmid DNA extraction.

### Preparation of cell-free reactions

Cell-free reactions were prepared as per the protocol by Sun and coworkers^14^. Unless otherwise noted, each reaction consisted of the following: 33% (by volume) of cell lysate, 1.5 mM of each amino acid (except for Leucine, which was added at 1.25 mM), 4.8 nM of NTP mix (containing 1.5 mM each of ATP and GTP, 0.9 mM each of CTP and UTP, pH adjusted to 7.5 using KOH), 50 mM HEPES pH 8, 0.2 mg/mL tRNA, 0.26 mM Coenzyme A, 0.33 mM NAD^+^, 0.75 mM cyclic AMP (cAMP), 0.068 mM Folinic acid, 1 mM Spermidine, 10 μM Malachite Green Dye, and 30 mM 3PGA. Depending on the batch of cell lysate, a standard reaction used either 8 mM magnesium glutamate and 80 mM potassium glutamate (for lysate Batches 2 and 3) or 6 mM magnesium glutamate and 140 mM potassium glutamate (for lysate Batch 1); these salt concentrations were chosen because they optimized deGFP expression in TX-TL reactions supplied with 1 nM P_OR1OR2_-MG aptamer-deGFP plasmid for a particular lysate batch (**Figure S15**). For the Mg^2+^ versus Fuel panels, 3PGA and Mg^2+^ were omitted from the master mix, and the appropriate concentration of Mg^2+^ and fuel was added to reactions individually. Finally, depending on the experiments, a variable amount of DNA plasmid was added, and the remaining volume of the 10 μL reaction was filled with nuclease-free water.

As reactions were all performed in 384-well glass-bottom plates, water, DNA, and other added components were titrated into the TX-TL reaction using the Echo 525 liquid handler, while the remaining components were added to a bulk solution and electronically dispensed by a multi-dispense pipette. Unless otherwise specified, all reactions containing DNA were supplied with the P_OR1OR2_-MG aptamer-deGFP plasmid. To prevent the reaction from starting before all the components had been added in, all reagents were kept on ice until they were added to the reaction and added sequentially to reduce evaporation and/or degradation. First, a master mix minus the cell lysate was created and kept on ice in an Eppendorf tube; then, components to be added by Echo were added to the 384-well plate, where all stock concentrations were diluted to less than 500mM so that they could be effectively dispensed via Echo; next, the cell lysate was added to the master mix; finally, the master mix was pipetted to the wall of each well. Once the plate was covered with a plastic seal, it was centrifuged at 4000 x *g* for 1 minute at 4°C. Finally, the bottom of the plate was quickly cleaned by Kimwipe to remove any debris and immediately transferred to a plate BioTek H1MF plate reader that had reached its setpoint of 29°C.

### Preparation of no-fuel, no Mg^2+^ cell-free reactions

Reactions were prepared as described in the “Preparation of cell-free reactions” with minor changes. No fuel or additional Mg^2+^ were added to the reactions, apart from what was already there from the preparation of the cell lysate. For the addition of the NTP mix, we prepared a NTP stock solution at a molar ratio of 1.5:1.5:0.9:0.9 for ATP: GTP: CTP: UTP, pH adjusted to 7.5 using KOH. For the ATP and GTP solutions, we prepared stock solutions in nuclease-free water. Each solution was added at 0, 5, 10, or 15 mM.

### Dynamic measurements of fluorescent molecules

Fluorescence measurements were performed by a BioTek H1MF plate reader at 29°C, and measurements were read from the bottom of the plate every 3 minutes (preceded by 5 seconds of linear shaking) at excitation/emission wavelengths suitable for MG aptamer (610/650 nm, Gain 150) and deGFP (485/515 nm, Gain 61) for 18 hours. For the experiment corresponding to Figure 4, excitation/emission wavelengths suitable for F30-Pepper (580/620 nm, Gain 150) and mTurquoise2 (434/474 nm, Gain 61) were used.

For MG aptamer and deGFP, arbitrary fluorescence units were converted to micromolar concentrations using a calibration curve prepared using purified deGFP protein and MG aptamer mRNA. To create a deGFP fluorescence calibration curve, serial dilutions of eGFP (purchased from Cell Biolabs) were performed in 1x PBS, after which 3 sample replicates of 10 μL each were loaded onto a 384-well plate and read by a BioTek H1MF plate reader at 29°C and at excitation/emission wavelengths of 485/515 and a Gain of 61. 3 technical replicates were read over 3 minutes at 1-minute intervals to generate 4 points per replicate, and the average of these 12 points was used as a single point at a given concentration through which the calibration line was fit with a zero intercept.

To create a MG aptamer fluorescence calibration curve, mRNA of the MG aptamer sequence only – specifically rArCrUrGrGrArUrCrCrCrGrArCrUrGrGrCrGrArGrArGrCrCrArGrGrUrArArCrG rArArUrGrGrArUrCrCrArArU – was purchased from IDT DNA. The lyophilized mRNA was re-suspended in nuclease-free water at its concentration was determined by a Nanodrop 2000c. Next, serial dilutions of MG aptamer mRNA were performed in 1X PBS containing 10 μM Malachite Green dye, after which 3-4 sample replicates of 10 μL each were loaded onto a 384-well plate and read by a BioTek H1MF plate reader at 29°C and at excitation/emission wavelengths of 610/650 and a Gain of 150. Four technical replicates were read over 10 minutes at 2.5-minute intervals to generate 5 points per replicate, and the average of these 20 points was used as a single point at a given concentration through which the calibration line was fit with a zero intercept. The slope of the background-subtracted calibration curve was used to calibrate measurements.

### mRNA purification for a translation-only system

To obtain an mRNA sequence containing MG aptamer-UTR1-deGFP, a linear DNA template was first prepared, followed by DNA isolation, *in vitro* transcription, DNAse treatment, and mRNA isolation. First, for generation of the linear DNA template, primers of the following sequences were first used to amplify the relevant sequence off the P_OR1OR2_-MG aptamer-deGFP plasmid by PCR: CCAGAAAACCGAATTTTGCTGG and ATGATAAAGAAGACAGTCATAAGTGCG. A 1-mL PCR reaction was performed using NEB Q5 2x Master Mix and run for 30 cycles with an elongation time of 45 seconds and an annealing temperature of 65°C.

After the PCR reaction was performed, the PCR was aliquoted in three 1.7-mL Eppendorf tubes (i.e., 333 μL per tube) and DNA was precipitated by adding 33 μL of 3 M sodium acetate and 1 mL 100% ethanol per tube. After chilling the tubes at -80°C for 20 minutes, the tubes were centrifuged at 16000 x *g* for 30 minutes at 4°C. Finally, all supernatant was pipetted out, and residual ethanol was allowed to evaporate by leaving the tubes open and uncovered at room temperature. The DNA was resuspended in 50 μL nuclease-free water per tube.

*In vitro* transcription was performed using the HiScribe® T7 High Yield RNA Synthesis Kit, and instructions included in the kit’s manual were used for DNase I treatment of the reaction.

Finally, the synthesized mRNA was isolated using the PureLink RNA Mini kit. 100% ethanol was first added to the reaction to a final concentration of 35%, after which the kit’s “Protocol for RNA clean-up and purification from liquid samples” was used (omitting the Lysis step). mRNA concentration was determined by a Nanodrop 2000c.

### Translation inhibition using tetracycline

The translation inhibition experiments were performed in Batch 2 lysate by adding 200 μg/mL tetracycline to TX-TL reactions. Stock tetracycline solutions were created at 80 mg/mL in DMSO, where they were stored at -80°C until further use. For addition to TX-TL reactions, the stock solution was diluted to 5 mg/mL in nuclease-free water and subsequently dispensed using the Echo 525 liquid handler. To determine the optimal concentration of tetracycline to use for translation-inhibition, titrations were performed (see **Figure S14**).

### Data analysis

Fluorescence measurements of deGFP and MG aptamer were converted to micromolar concentrations via the calibration curves. Maximum deGFP concentration was determined to be the maximum deGFP concentration achieved for a given experimental condition at any point over the course of the 18-hour experiment. Integrated MG aptamer measurements were made by using the NumPy^40^ **trapz** function for numerical integration using the trapezoidal method.

Plots were generated using the Python package Bokeh^41^. The Jupyter notebooks that were used for creating tidy data and for subsequent data analysis and plot generation are included with this work, in addition to all the data that was collected for this study.

### Spearman’s rho correlation computation

For figures where a Spearman’s rho correlation value was computed, the SciPy^42^ **stats.spearmanr** function was used to compute correlation values. A Spearman rho correlation value has a minimum possible value of -1, corresponding to a perfect negative rank correlation between two variables, and a maximum possible value of 1, corresponding to a perfect positive rank correlation between two variables. A value of 0 suggests no rank correlation.

Where a bootstrapped Spearman’s rho correlation value was computed, the pseudocode below shows the algorithm used to perform the computation. Briefly, we first split our data into training and test sets in a 75/25 ratio, respectively, where the predictor variable was Mg^2+^ or 3PGA and the predicted data was the 2D trade-off data. We next trained a Partial Least Squares Regression (Canonical) model on the training data using the Python Scikit-learn^43^ package. This training simultaneously reduced the dimensionality of the trade-off data from 2D to 1D; re-dimensionalized the predictor variable (i.e., Mg^2+^ or 3PGA) so that it had a mean value of zero; and created a model capable of predicting position along the 1D trade-off data for the predictor variable. The model was then used to map the test data to the new 1D spaces. A Spearman rho correlation value was computed for the mapped test data between the 1D predictor variable and the 1D trade-off data using the SciPy^42^ **stats.spearmanr** function. This process was repeated 5000 times to randomize the splitting of data. The mean and standard deviation of this “bootstrapped” Spearman’s rho correlation value are reported for each plot where applicable.

**Algorithm 1.**
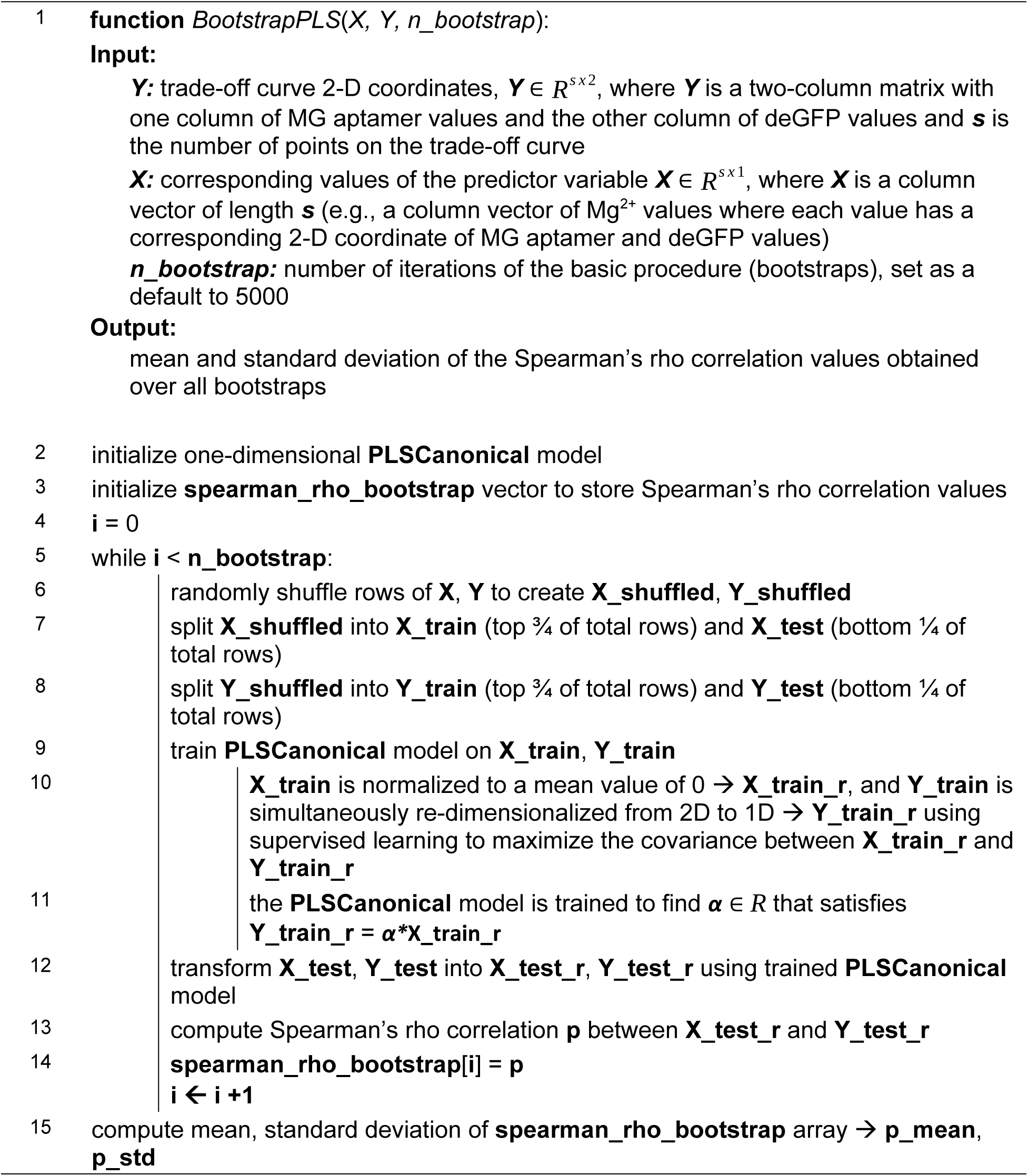
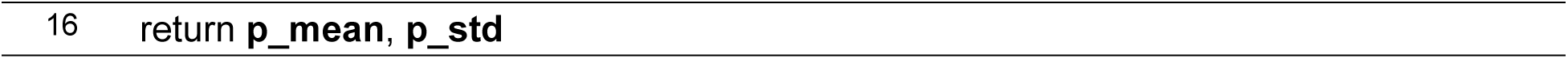
Bootstrapped Spearman’s Rho Correlation.

Although many cross decomposition methods exist, we chose Partial Least Squares (PLS) Regression, because it is better at preserving variance compared to methods like Principal Component Regression during the dimensionality reduction step (due to the addition of supervising learning). PLS is also particularly suitable for when there is collinearity among columns of the predictor or predicted data (as in the case of MG aptamer and deGFP, where we observed a negative correlation between the two variables). The Python Scikit-learn package has several functions for PLS: PLSCanonical, PLSSVD, and PLSRegression (which can implement the PLS1 and PLS2 algorithms); in the case of a one-dimensional PLS model as we have chosen, the underlying algorithm is the same for all of them, so we have arbitrarily chosen PLSCanonical.

Due to the re-dimensionalization of data during PLS regression, the bootstrapped Spearman’s rho correlations range from 0 to 1 (unlike the traditional Spearman’s rho correlation that ranges from -1 to 1). In the bootstrapped case, a value of 0 corresponds to no rank correlation and a value of 1 corresponds to a perfect rank correlation (with a direction of positive or negative stated in the Results section where applicable).

## Supporting information

Supplemental Figures and DNA sequences

Raw Data, Tidy Data, DNA GenBank files, Figures, and Jupyter Notebooks for Figure Generation and Analysis

## Acknowledgements

The work described here would not be possible without the initial ideas that came out of extensive discussions with William Poole. His suggestions to probe the roles of Mg^2+^ and different fuel sources helped generate new questions and analyses that ultimately led to the experiments discussed here, and we thank him for his insights. Apart from Poole, additional members of the lab – specifically Ankita Roychoudhury, David Alexander Johnson, Miryong (Miki) Yun, and Zoila Jurado – also contributed to the work shown here. Zoila was present in and contributed to many of these early discussions with William, and she also prepared the P_OR1OR2_-MG aptamer-deGFP and P_T7_ -deGFP-MG aptamer plasmids used throughout this study. All four individuals helped prepare the cell lysate batches used in the experiments here, and we thank them for their help. We thank Paul Freemont for his helpful comments, particularly those pertaining to cell-free metabolism. We also thank lab members John Marken, Yan Zhang, and Zoila Jurado for providing critical feedback on this manuscript. Finally, ChatGPT was used to write part of the introduction section, which has been an interesting and helpful endeavor.

Research was sponsored by the Army Research Office and was accomplished under Cooperative Agreement Number W911NF-22-2-0210. The views and conclusions contained in this document are those of the authors and should not be interpreted as representing the official policies, either expressed or implied, of the Army Research Office or the U.S. Government. The U.S. Government is authorized to reproduce and distribute reprints for Government purposes notwithstanding any copyright notation herein.

## Author Information

### Author Contributions

M.K. conceived and designed the study, performed experiments, acquired and analyzed the data, and wrote the manuscript. R.M.M. supervised the project and provided feedback on the manuscript.

## Supporting Information

The Supporting Information includes figures corresponding to experiments exploring:

○ the dependence of MG aptamer and deGFP fluorescence on pH, Mg^2+^, and 3PGA concentration (**Figure S1**)
○ the effects of Mg^2+^ concentration on the TX-TL trade-off by fuel type (**Figure S2**)
○ the effects of 3PGA concentration on the TX-TL trade-off across different lysate volume fractions (**Figure S3**)
○ the TX-TL trade-off in systems expressing MG aptamer and deGFP in different order under a P_T7_ promoter (**Figure S4**)
○ the TX-TL trade-off in systems expressing MG aptamer and deGFP under different promoters (**Figure S5**)
○ the scaling of transcription and TL with increasing DNA concentration (**Figure S6**)
○ the effects of total ATP versus total GTP concentration in NTP-fueled systems (**Figure S7**)
○ unusual and potentially desirable transcription and translation dynamics in TX-TL systems with no fuel or Mg^2+^ and with additional energy (**Figure S8**)
○ maximum deGFP versus maximum MG aptamer slope for central carbon- and NTP-fueled systems (**Figure S9**)
○ **h**eatmaps of integrated MG aptamer expression and deGFP expression at different Mg^2+^ and fuel concentrations at different DNA concentrations, for 3PGA, maltose, pyruvate, and succinate, respectively (**Figures S10-S13**)
○ tetracycline titrations (**Figure S14**)
○ salt calibrations for cell lysate batches (**Figure S15**)

The Supporting Information also includes plasmid sequences for most plasmids used in this study (except for the P_T7_-deGFP-MGapt plasmid), in addition to all data and code used for analysis and figure generation. The same data and code are also available on GitHub at the following link: https://github.com/mkapasiawala/txtl-tradeoff.

