## Supplemental Figures and DNA sequences for "Metabolic perturbations to an *E. coli*-based cell-free system reveal a trade-off between transcription and translation"

**Author list:** Manisha Kapasiawala[1]*, Richard M. Murray[1]

**Author address:**
[1]Division of Biology and Biological Engineering, California Institute of Technology, Pasadena, CA 91125, USA

### Table of contents

#### Supporting figures


- **Figure S1:** Exploring the dependence of MG aptamer and deGFP fluorescence on pH, $Mg^{2+}$, and 3PGA concentration.
- **Figure S2:** Effects of $Mg^{2+}$ concentration on the TX-TL trade-off by fuel type.
- **Figure S3:** Effects of 3PGA concentration on the TX-TL trade-off across different lysate volume fractions
- **Figure S4:** The TX-TL trade-off in systems expressing MG aptamer and deGFP in different order under a $P_{T7}$ promoter.
- **Figure S5:** The TX-TL trade-off in systems expressing MG aptamer and deGFP under different promoters.
- **Figure S6:** Scaling of transcription and translation with increasing DNA concentration.
- **Figure S7:** Exploring the effects of total ATP versus total GTP concentration in NTP-fueled systems.
- **Figure S8:** Examples of unusual and potentially desirable transcription and translation dynamics in TX-TL systems with no fuel or $Mg^{2+}$ and with additional energy.
- **Figure S9:** Maximum deGFP versus maximum MG aptamer slope for central carbon- and NTP-fueled systems
- **Figures S10-S13:** Heatmaps of integrated MG aptamer expression and deGFP expression at different $Mg^{2+}$ and fuel concentrations at different DNA concentrations, for 3PGA, maltose, pyruvate, and succinate, respectively
- **Figure S14:** Tetracycline titrations.
- **Figure S15:** Salt calibrations for cell lysate batches.


#### Plasmid sequences


- **$P_{OR1OR2}$-MG aptamer-deGFP**
- **$P_{T7}$-MG aptamer-deGFP**
- **$P_{Tet}$-MG aptamer-deGFP**
- **$P_{Tet}$-F30-Pepper-mTurquoise2**

### Supporting Figures

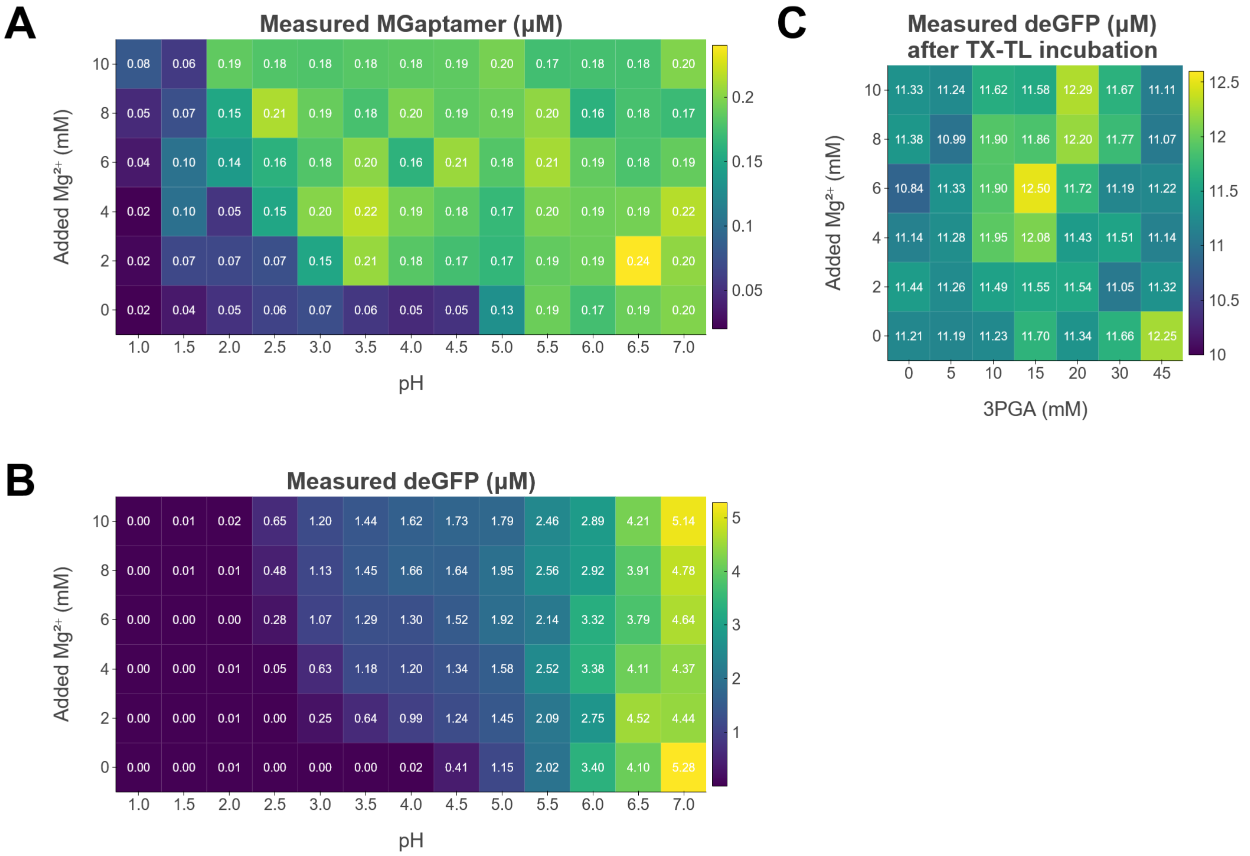


**Figure S1:** Exploring the dependence of MG aptamer and deGFP fluorescence on pH, $Mg^{2+}$, and 3PGA concentration. In **(A)** and **(B)**, either **(A)** 0.2 µM MG aptamer-UTR1-deGFP mRNA or **(B)** 6.02 µM purified deGFP was added to 9 uL of phosphate-buffered saline (PBS) of the appropriate pH, some amount of 100 mM Mg-glutamate to the appropriate concentration, and nuclease-free water to 10 µL. The 384-well plate was incubated for 18 hours in a plate-reader at 29°C, and the endpoint concentrations, as determined by the fluorescence calibration data, are indicated in the plots shown. **(C)** 10 µM purified deGFP was added to TX-TL reactions and incubated at 29°C for 18 hours. The final deGFP concentrations, as reported by the fluorescence values that were calibrated to micromolar units, are shown in the figure. Fluorescence calibrations were performed using concentrations of 2-40 µM purified deGFP in PBS, where the linear slope of the calibration was dominated by higher concentrations of deGFP compared to the concentrations used in (B) and (C). Thus, fluorescence values are lower than expected in (B), where a higher percentage of the deGFP is likely adsorbed onto the walls of the 384-well plate (compared to the calibration), and higher than expected in (C), where a lower percentage of deGFP is adsorbed (compared to the calibration) since the proteins from the cell lysate are competing for adsorption.

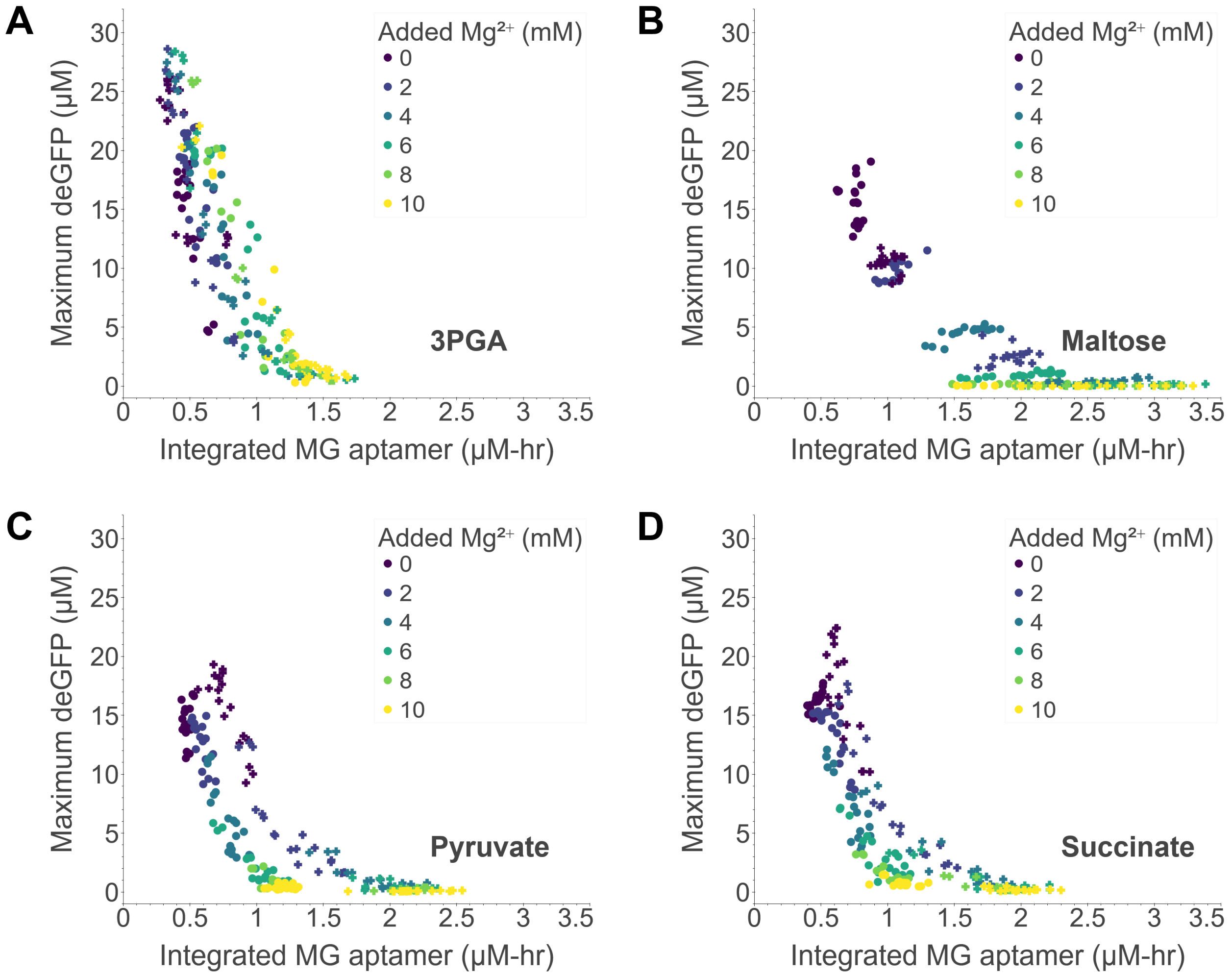


**Figure S2:** Effects of $Mg^{2+}$ concentration on the TX-TL trade-off by fuel type. Each point represents one of three replicates of a particular set of fuel and $Mg^{2+}$ concentrations, where fuel is either **(A)** 3PGA, **(B)** maltose, **(C)** pyruvate, or **(D)** succinate, with corresponding bootstrapped Spearman's rho correlations of 0.63 ± 0.05, 0.75 ± 0.04, 0.76 ± 0.04, and 0.78 ± 0.04, respectively. The data shown reflect experiments performed in two batches of cell lysate, Batch 1 (**o**) and Batch 2 (**+**) (see Methods and Materials for details), and all experiments were performed using 5 nM DNA.

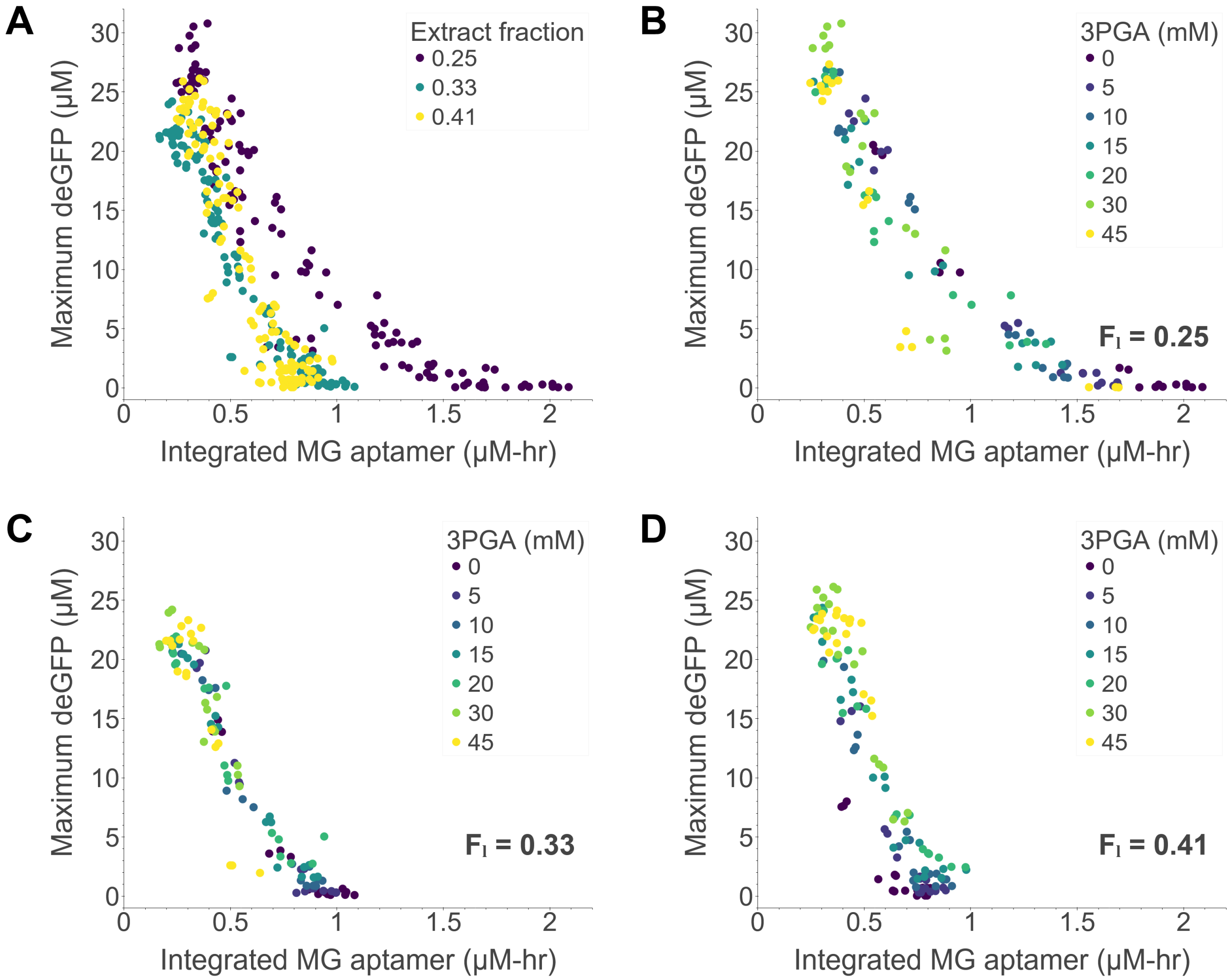


**Figure S3:** Effects of 3PGA concentration on the TX-TL trade-off across different lysate volume fractions. Each point represents one of three replicates of a particular set of 3PGA and $Mg^{2+}$ concentrations. **(A)** Trade-off curves from three different lysate volume fractions are overlaid, where the lysate volume fraction is the volume fraction of a TX-TL reaction that is made up of cell lysate. **(B) – (D)** Individual lysate fraction trade-off curves are shown and points are colored by 3PGA concentration, where the curves correspond to lysate fractions **(B)** $\mathbf{F_L}$ = 0.25, **(C)** $\mathbf{F_L}$ = 0.33, the nominal case; and **(D)** $\mathbf{F_L}$ = 0.41, with corresponding bootstrapped Spearman's rho correlations of 0.49 ± 0.09, 0.63 ± 0.06, and 0.63 ± 0.05, respectively. The data shown reflect experiments performed in cell lysate Batch 2 (see Methods and Materials for details), and all experiments were performed using 5 nM DNA.

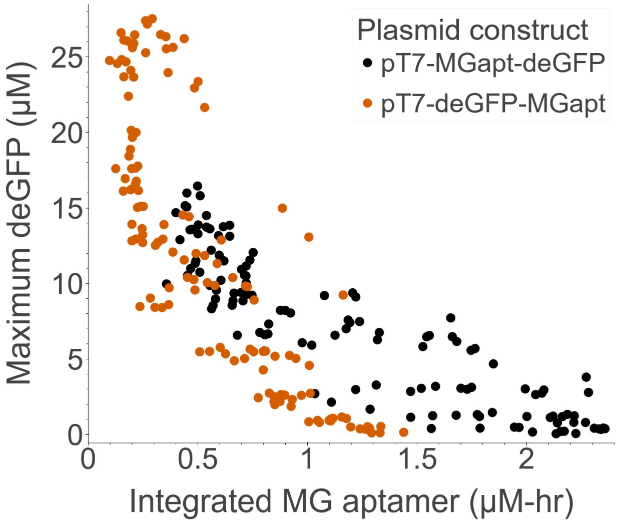


**Figure S4:** The TX-TL trade-off in systems expressing MG aptamer and deGFP in different order under a $P_{T7}$ promoter. Each point represents one of three replicates of a particular set of 3PGA and $Mg^{2+}$ concentrations. The data for $P_{T7}$-MG aptamer-deGFP are the same shown in **Figure 4B** and **S5**. The data shown reflect experiments performed in Batch 3 of cell lysate (see Methods and Materials for details), and all experiments were performed using 5 nM DNA.

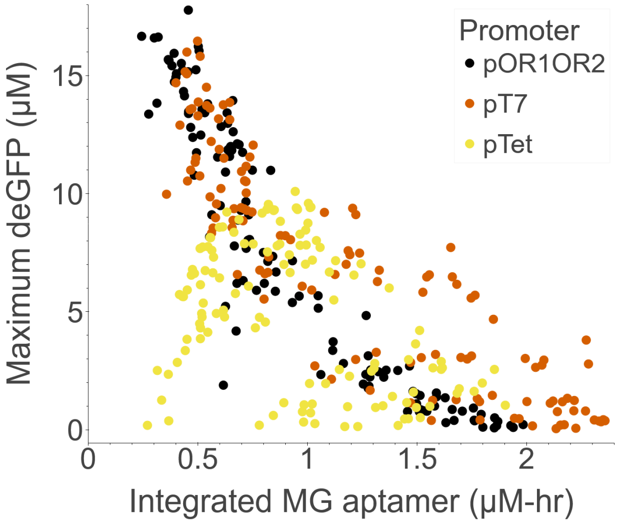


**Figure S5:** The TX-TL trade-off in systems expressing MG aptamer and deGFP under different promoters. Each point represents one of three replicates of a particular set of 3PGA and $Mg^{2+}$ concentrations. The data are the same shown in **Figure 4**, albeit with the data from panels **(A)**, **(B)**, and **(D)** overlaid and colored by promoter type. The data shown reflect experiments performed in Batch 3 of cell lysate (see Methods and Materials for details), and all experiments were performed using 5 nM DNA.

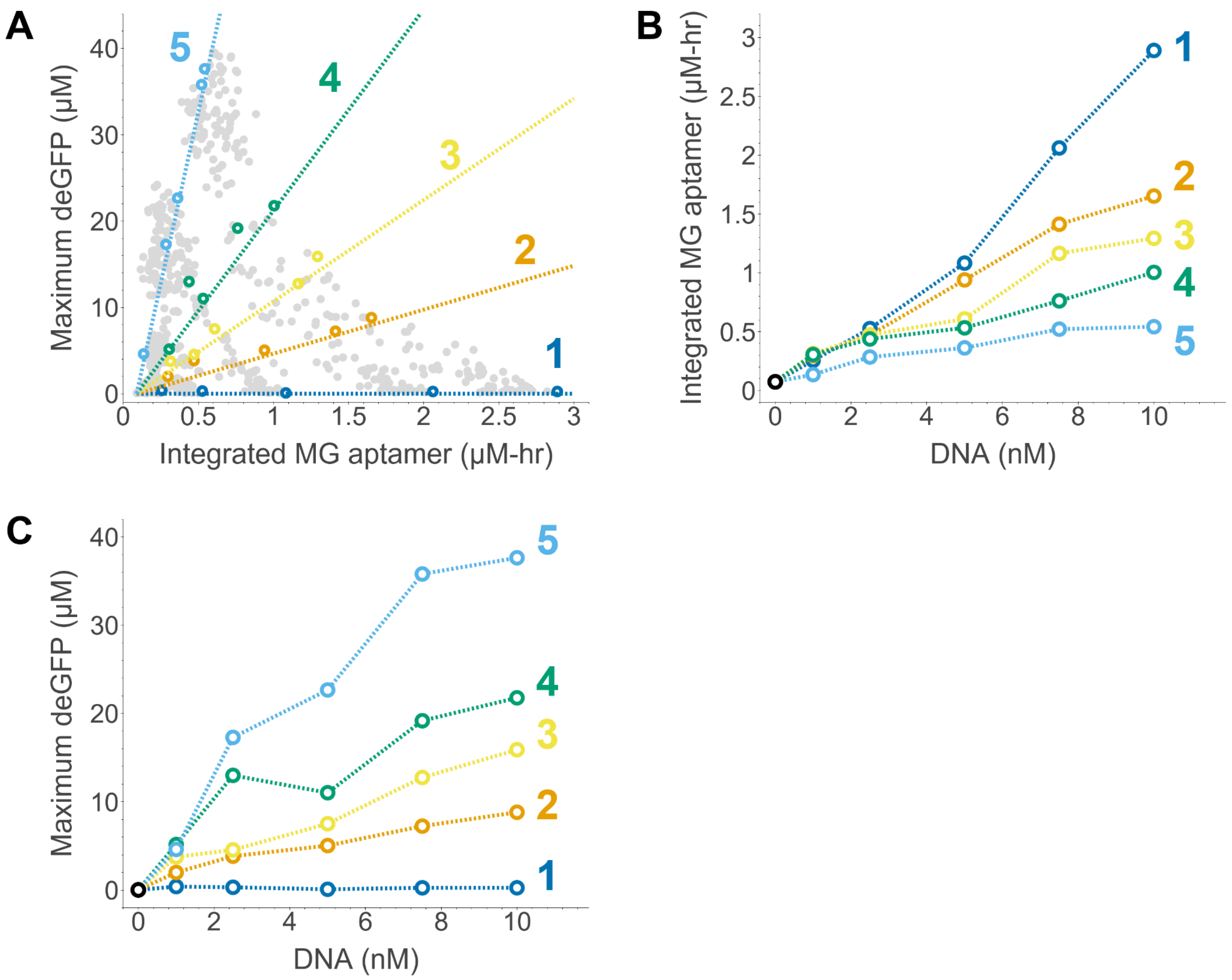


**Figure S6:** Scaling of transcription and translation with increasing DNA concentration. **(A)** Five lines were chosen to intersect the trade-off curves at roughly equi-angular intervals, and the point on each trade-off curve corresponding most closely to that line is highlighted. **(B)** Integrated MG aptamer versus DNA concentration for each of the intersecting lines for the points highlighted in (A). **(C)** Maximum deGFP versus DNA concentration for each of the intersecting lines for the points highlighted in (B). All experiments were performed in Batch 2 of cell lysate. The data in (A) are the same shown in **Figure 5.**

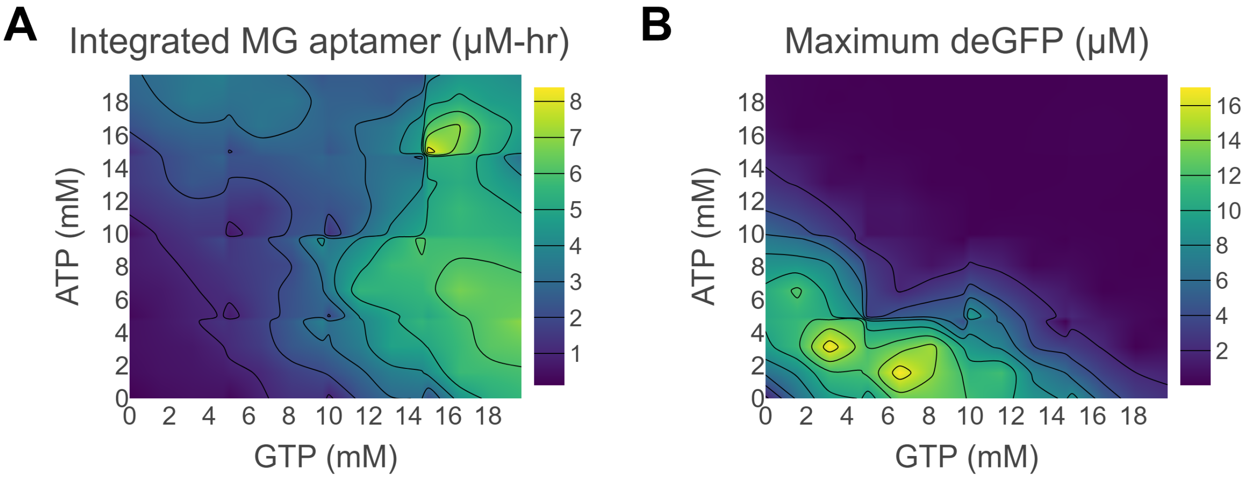


**Figure S7:** Exploring the effects of total ATP versus total GTP concentration in NTP-fueled systems. Here, the systems explored were TX-TL systems using 5 nM DNA to which no central carbon fuel nor $Mg^{2+}$ had been added. Systems fueled by NTPs were implemented using a three-way panel of ATP versus GTP versus NTP mix, where each solution was added at 0, 5, 10, or 15mM. As the NTP mix also contained ATP and GTP, the total concentrations of ATP and GTP were calculated for each system by summing the amount of ATP or GTP in the NTP mix with the added amount of ATP or GTP, respectively. Results were then averaged over three replicates and two batches of cell lysate (Batch 1 and Batch 2). **(A)** Integrated MG aptamer values for NTP-fueled systems. **(B)** Maximum deGFP values for the NTP-fueled systems.

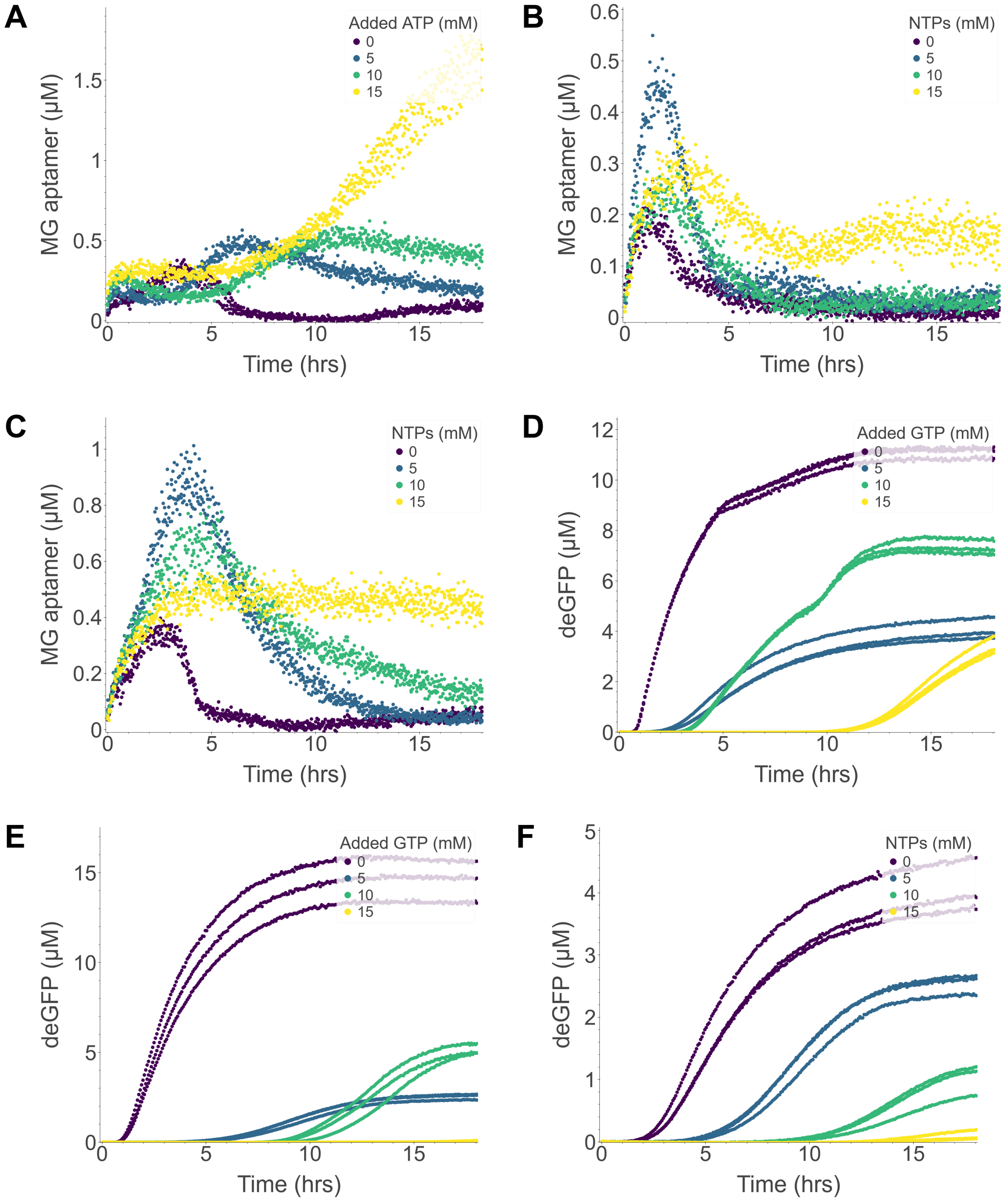


**Figure S8:** Examples of unusual and potentially desirable transcription and translation dynamics in TX-TL systems with no fuel or $Mg^{2+}$ and with additional energy. These systems have the following (NTP, Added ATP, Added GTP) concentrations in mM, where a dashed line (-) indicates that a concentration is being varied in that plot: **(A)** (0, -, 15), **(B)** (-, 5, 5), **(C)** (-, 5, 10), **(D)** (0, 5, -), **(E)** (5, 5, -), **(F)** (-, 5, 5). **(A)** shows an example of a system (at 15mM ATP)

exhibiting time-delayed transcription albeit with a high MG aptamer yield that continues increasing even at 18 hours. **(B)** and **(C)** show examples of systems with biphasic and sustained transcription (both at 15mM NTPs), respectively. **(D)** shows examples of systems with varying translation dynamics, ranging from a near-linear initial increase in deGFP (0mM GTP), biphasic translation (10mM GTP), slow translation (5mM GTP), and extremely time-delayed translation (15mM GTP). **(E)** demonstrates that TX-TL systems that have time-delayed translation do not necessarily have lower deGFP yields, as indicated in the comparison between the 5mM GTP and 10mM GTP cases. **(F)** shows that under certain regimes (here at 5mM ATP and 5mM GTP), TX-TL systems can exhibit similar translation dynamics, albeit with time delays that are proportional to the amount of NTPs added to the system. Each point represents one of three replicates of a particular set of NTP, Added ATP, and Added GTP concentrations. All experiments were performed at 5 nM DNA in systems using the Batch 2 lysate. Note: axes have different scales for each plot.

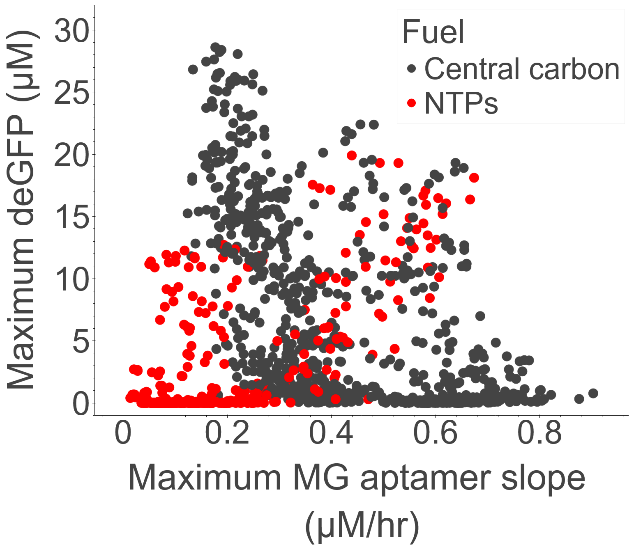


**Figure S9:** Maximum deGFP versus maximum MG aptamer slope for central carbon- and NTP-fueled systems. The data shown here are from the same experiments whose data is shown in **Figure 7B**. Each point represents one of three replicates of a particular set of NTP, added ATP, and added GTP concentrations, colored by the fuel type. Here, the “central carbon” fuel data is the same as in Figure 1, and those reactions all contain 4.8 mM NTPs. The “NTPs” fuel data reflect experiments where no central fuel or $Mg^{2+}$ has been added. Maximum MG aptamer slope was calculated for each experimental condition by computing a series of linear regressions between MG aptamer and time for points spanning a rolling window of 1 hour over a total period of 18 hours; the maximum slope of MG aptamer expression over the 18-hour period was plotted for each experimental condition. The data shown reflect experiments performed in two batches of cell lysate, Batch 1 and Batch 2 (see Methods and Materials for details), and all experiments were performed using 5 nM DNA.

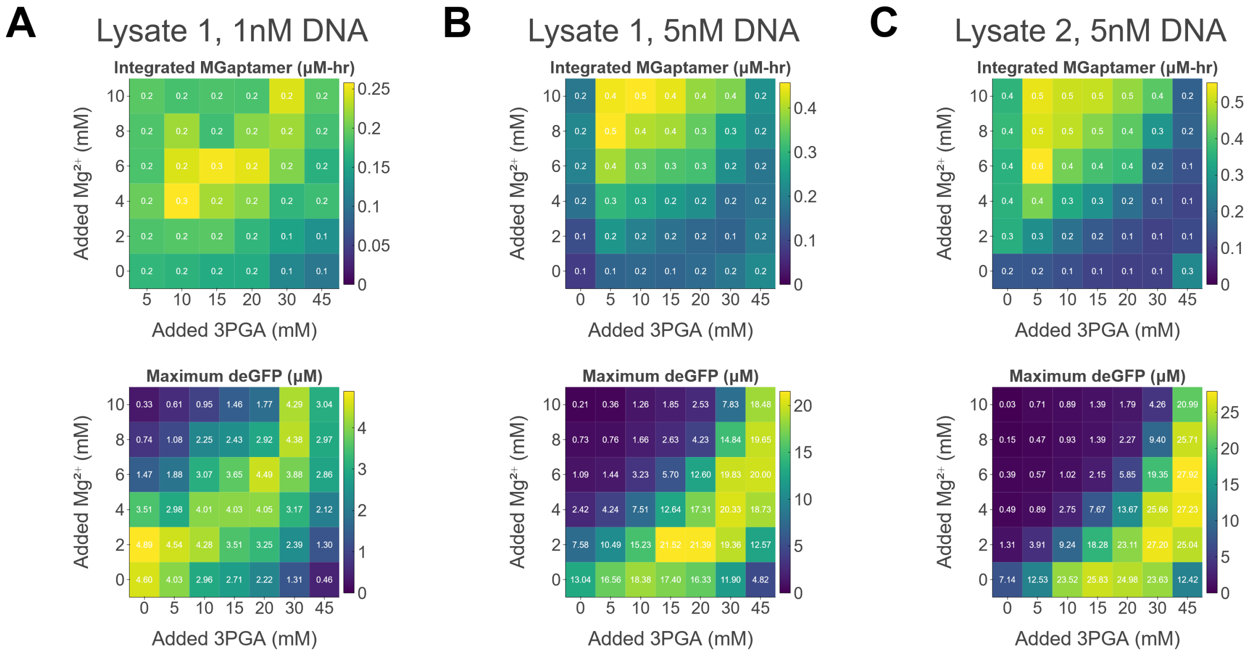


**Figure S10:** Heatmaps of integrated MG aptamer expression and deGFP expression at different $Mg^{2+}$ and 3PGA concentrations at different DNA concentrations. Experiments were performed using **(A)** Lysate Batch 1 and 1 nM DNA, **(B)** Lysate Batch 1 and 5 nM DNA, **(C)** Lysate Batch 2 and 5 nM DNA. Each value is an average of three replicates for that condition. Colorbars were scaled separately for each plot, and all colorbars were normalized to a minimum value of zero.

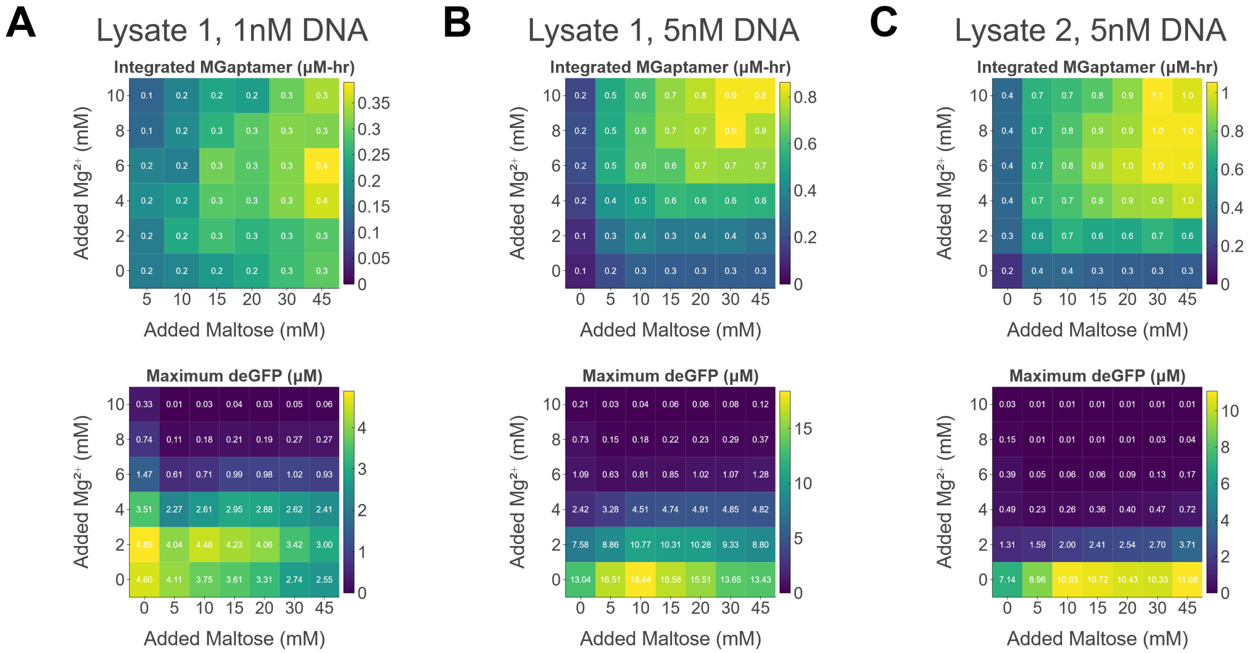


**Figure S11:** Heatmaps of integrated MG aptamer expression and deGFP expression at different $Mg^{2+}$ and maltose concentrations at different DNA concentrations. Experiments were performed using **(A)** Lysate Batch 1 and 1 nM DNA, **(B)** Lysate Batch 1 and 5 nM DNA, **(C)** Lysate Batch 2 and 5 nM DNA. Each value is an average of three replicates for that condition. Colorbars were scaled separately for each plot, and all colorbars were normalized to a minimum value of zero.

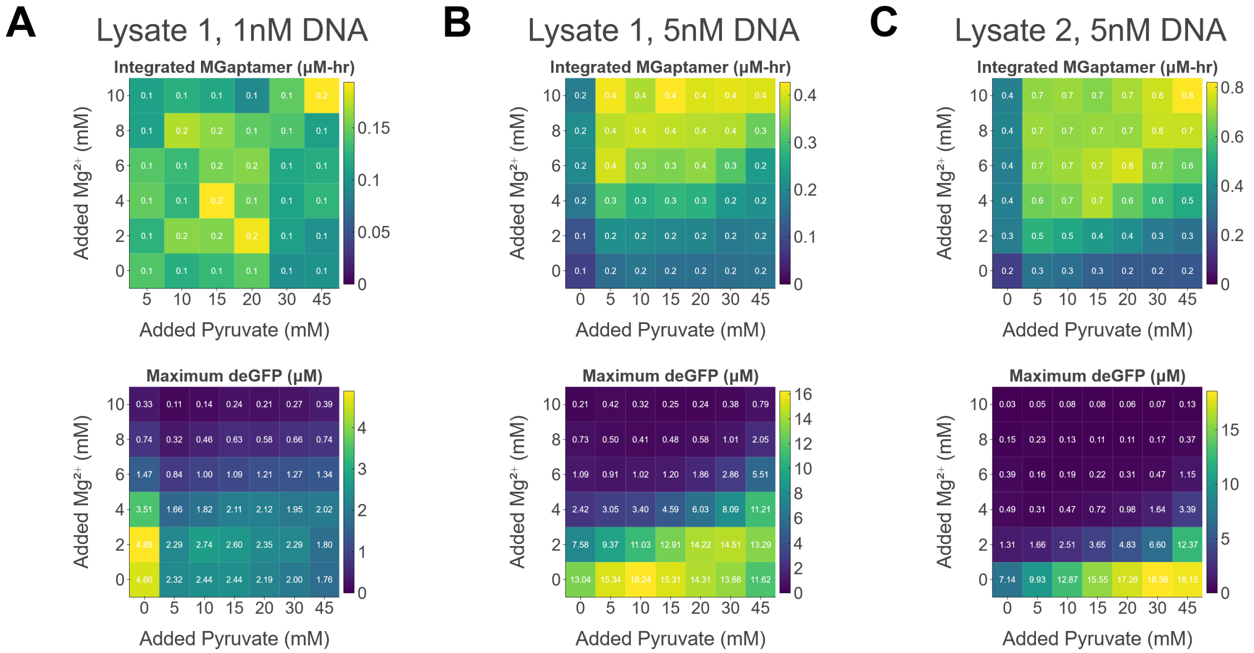


**Figure S12:** Heatmaps of integrated MG aptamer expression and deGFP expression at different $Mg^{2+}$ and pyruvate concentrations at different DNA concentrations. Experiments were performed using **(A)** Lysate Batch 1 and 1 nM DNA, **(B)** Lysate Batch 1 and 5 nM DNA, **(C)** Lysate Batch 2 and 5 nM DNA. Each value is an average of three replicates for that condition. Colorbars were scaled separately for each plot, and all colorbars were normalized to a minimum value of zero.

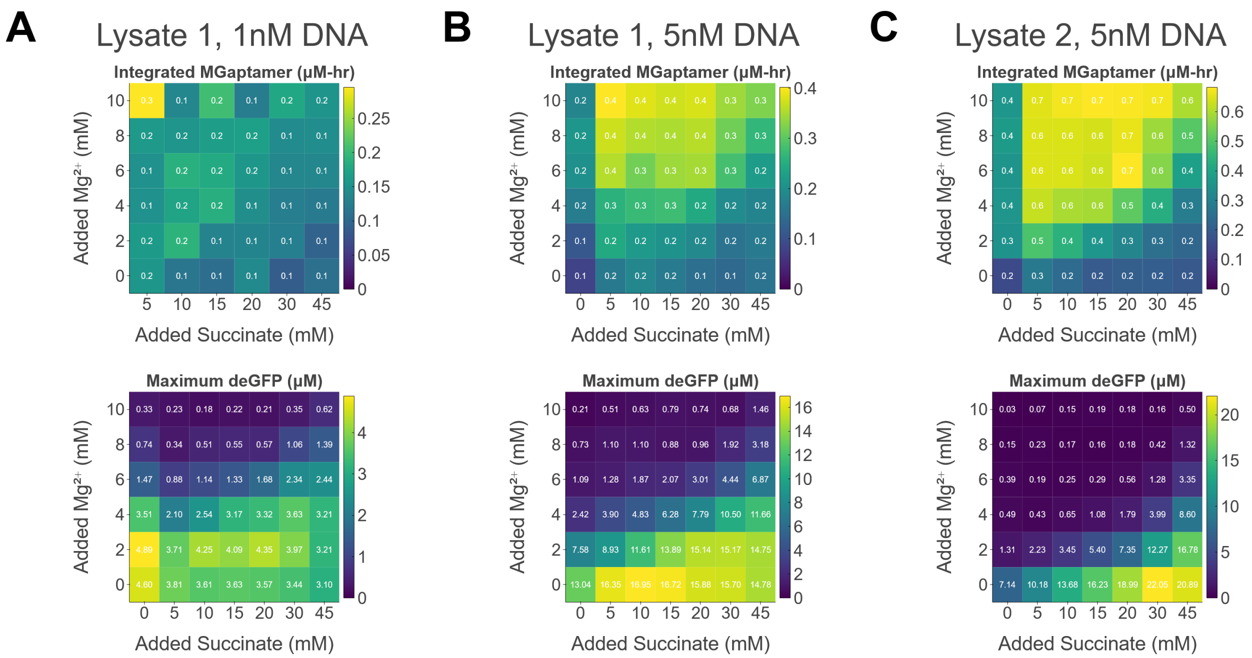


**Figure S13:** Heatmaps of integrated MG aptamer expression and deGFP expression at different $Mg^{2+}$ and succinate concentrations at different DNA concentrations. Experiments were performed using **(A)** Lysate Batch 1 and 1 nM DNA, **(B)** Lysate Batch 1 and 5 nM DNA, **(C)** Lysate Batch 2 and 5 nM DNA. Each value is an average of three replicates for that condition. Colorbars were scaled separately for each plot, and all colorbars were normalized to a minimum value of zero.

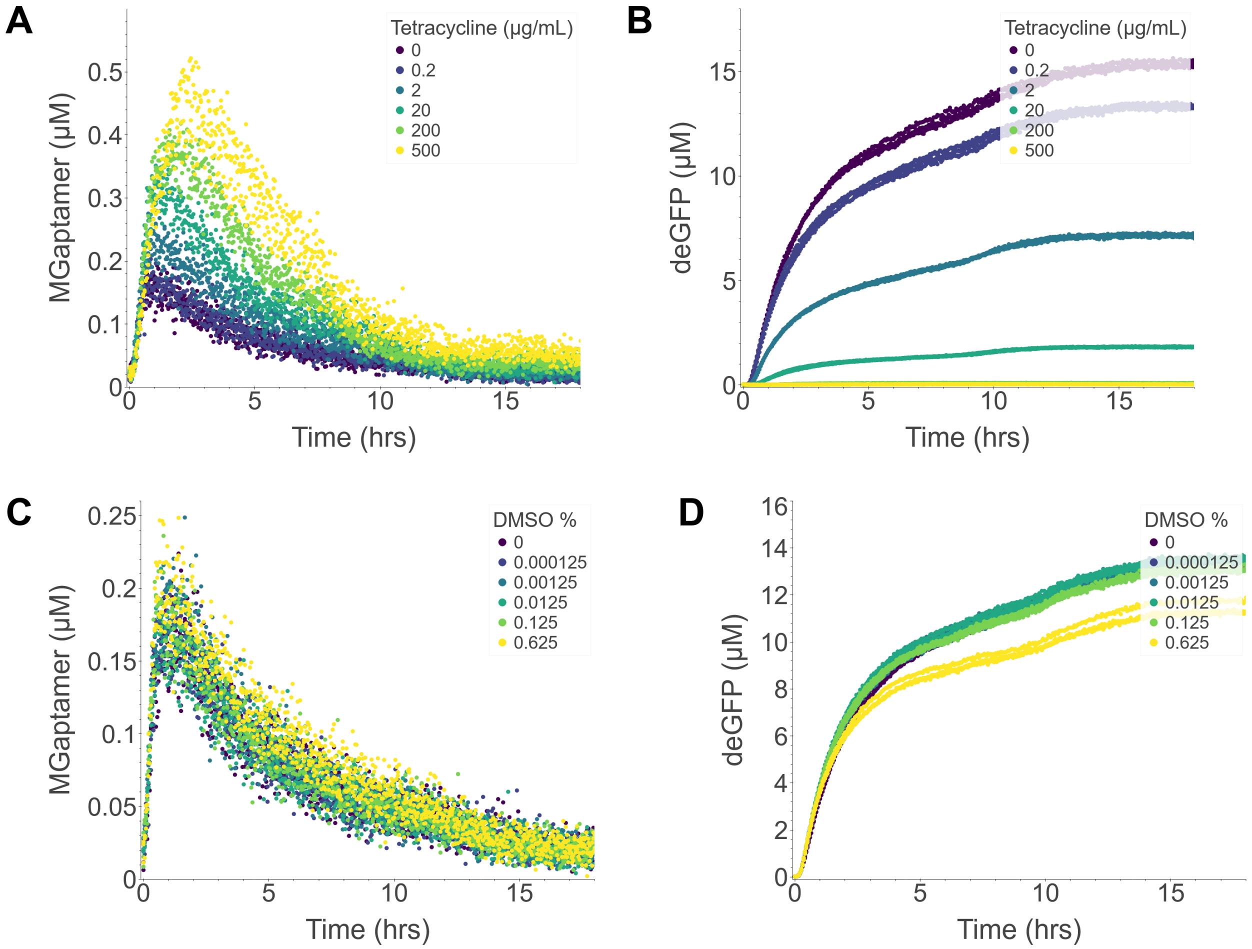


**Figure S14:** Tetracycline titrations. Titrations were performed to determine the concentration of tetracycline necessary for translation inhibition albeit without adding enough DMSO to the reaction to additionally reduce transcription or translation. Controls were performed by adding the equivalent amount of DMSO to make sure the tetracycline alone was impacting translation. **(A)** and **(B)** show transcription and translation data, respectively, for TX-TL reactions performed with 200 µg/mL tetracycline. **(C)** and **(D)** show transcription and translation data, respectively, for TX-TL reactions performed with the equivalent amount of DMSO for each of the conditions in **(A)** and **(B)**. TX-TL reactions were performed using the Batch 2 lysate with 5 nM DNA at 30 mM 3PGA and 8 mM $Mg^{2+}$.

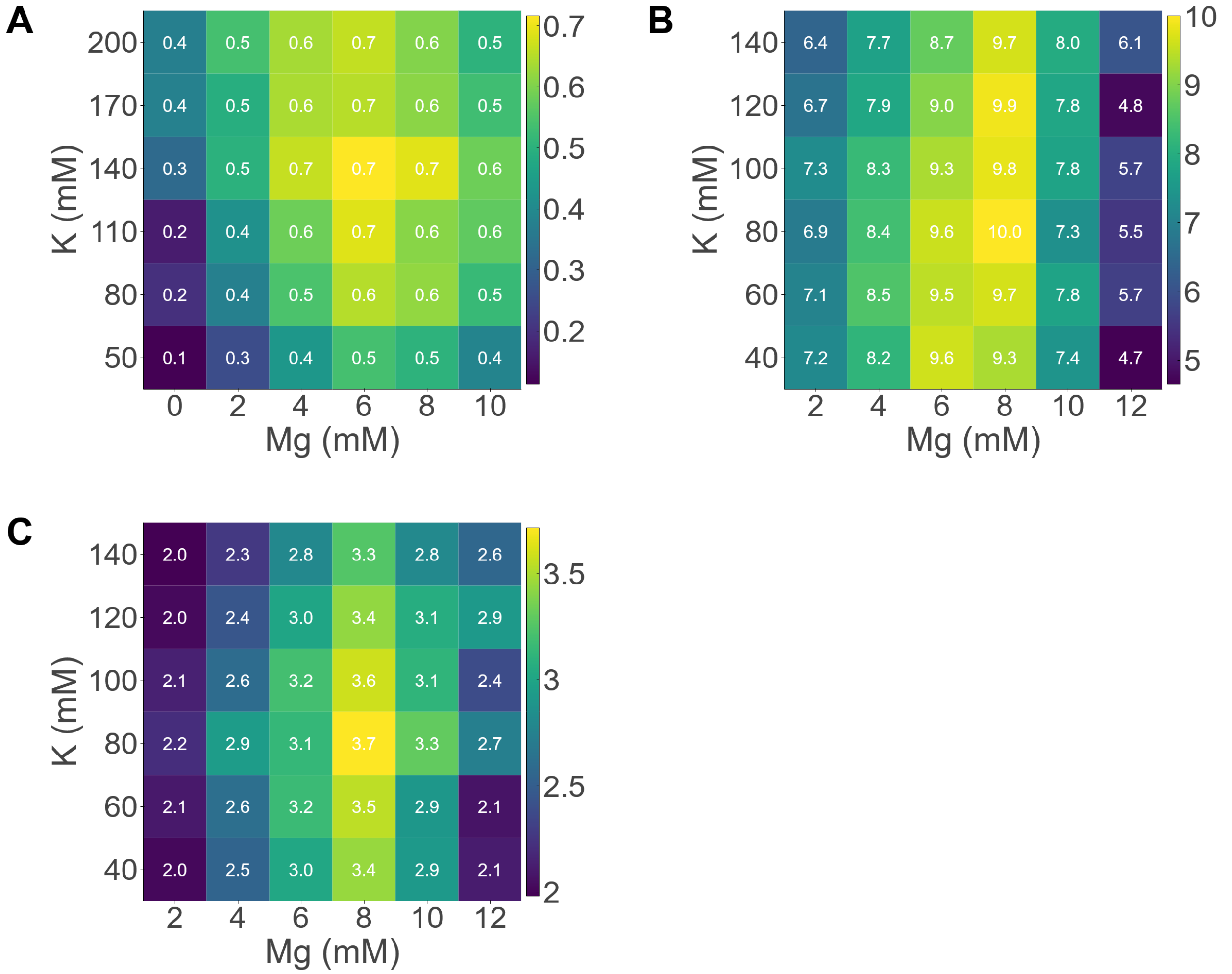


**Figure S15:** Salt calibrations for cell lysate batches. As each batch of cell lysate has different optimal $Mg^{2+}$ and $K^{+}$ concentrations, the optimal concentration for each batch was determined by a salt panel, where TX-TL reactions were performed with the standard formulation, albeit with different pairwise concentrations of Mg-glutamate and K-glutamate. The endpoint deGFP concentration (μM) was recorded after an 18-hour incubation at 29°C for lysate batches **(A)** 1, **(B)** 2, and **(C)** 3. Colorbars are scaled separately for each plot, and their range spans the minimum and maximum endpoint deGFP value for each experiment.

**Plasmid sequences**

The following are the sequences of four (out of five) plasmids used in this study. Promoters are highlighted in red, mRNA aptamer in blue, untranslated region (5' UTR) in yellow, protein sequence in green, and terminator sequence in magenta.

**$P_{OR1OR2}$-MG aptamer-UTR1-deGFP-T500 sequence (pZJQ.27):**

TGAGCTAACACCGTGCGTGTTGACAATTTTACCTCTGGCGGTGATAATGGTTGCAGCTAGC
GGGATCCCGACTGGCGAGAGCCAGGTAACGAATGGATCCAATAATTTTGTTTAACTTTAAG
AAGGAGATATACCATGGAGCTTTTCACTGGCGTTGTTCCCATCCTGGTCGAGCTGGACGG
CGACGTAAACGGCCACAAGTTCAGCGTGTCCGGCGAGGGCGAGGGCGATGCCACCTACG
GCAAGCTGACCCTGAAGTTCATCTGCACCACCGGCAAGCTGCCCGTGCCCTGGCCCACCC
TCGTGACCACCCTGACCTACGGCGTGCAGTGCTTCAGCCGCTACCCCGACCACATGAAGC
AGCACGACTTCTTCAAGTCCGCCATGCCCGAAGGCTACGTCCAGGAGCGCACCATCTTCT
TCAAGGACGACGGCAACTACAAGACCCGCGCCGAGGTGAAGTTCGAGGGCGACACCCTG
GTGAACCGCATCGAGCTGAAGGGCATCGACTTCAAGGAGGACGGCAACATCCTGGGGCA
CAAGCTGGAGTACAACTACAACAGCCACAACGTCTATATCATGGCCGACAAGCAGAAGAAC
GGCATCAAGGTGAACTTCAAGATCCGCCACAACATCGAGGACGGCAGCGTGCAGCTCGCC
GACCACTACCAGCAGAACACCCCCATCGGCGACGGCCCCGTGCTGCTGCCCGACAACCA
CTACCTGAGCACCCAGTCCGCCCTGAGCAAAGACCCCAACGAGAAGCGCGATCACATGGT
CCTGCTGGAGTTCGTGACCGCCGCCGGGATCTAACTCGAGCAAAGCCCGCCGAAAGGCG
GGCTTTTCTGTGTCGACCGATGCCCTTGAGAGCCTTCAACCCAGTCAGCTCCTTCCGGTG
GGCGCGGGGCATGACTATCGTCGCCGCACTTATGACTGTCTTCTTTATCATGCAACTCGTA
GGACAGGTGCCGGCAGCGCTCTTCCGCTTCCTCGCTCACTGACTCGCTGCGCTCGGTCGT
TCGGCTGCGGCGAGCGGTATCAGCTCACTCAAAGGCGGTAATACGGTTATCCACAGAATC
AGGGGATAACGCAGGAAAGAACATGTGAGCAAAAGGCCAGCAAAAGGCCAGGAACCGTAA
AAAGGCCGCGTTGCTGGCGTTTTTCCATAGGCTCCGCCCCCCTGACGAGCATCACAAAAA
TCGACGCTCAAGTCAGAGGTGGCGAAACCCGACAGGACTATAAAGATACCAGGCGTTTCC
CCCTGGAAGCTCCCTCGTGCGCTCTCCTGTTCCGACCCTGCCGCTTACCGGATACCTGTC
CGCCTTTCTCCCTTCGGGAAGCGTGGCGCTTTCTCAATGCTCACGCTGTAGGTATCTCAGT
TCGGTGTAGGTCGTTCGCTCCAAGCTGGGCTGTGTGCACGAACCCCCCGTTCAGCCCGAC
CGCTGCGCCTTATCCGGTAACTATCGTCTTGAGTCCAACCCGGTAAGACACGACTTATCGC
CACTGGCAGCAGCCACTGGTAACAGGATTAGCAGAGCGAGGTATGTAGGCGGTGCTACAG
AGTTCTTGAAGTGGTGGCCTAACTACGGCTACACTAGAAGGACAGTATTTGGTATCTGCGC
TCTGCTGAAGCCAGTTACCTTCGGAAAAAGAGTTGGTAGCTCTTGATCCGGCAAACAAACC
ACCGCTGGTAGCGGTGGTTTTTTTGTTTGCAAGCAGCAGATTACGCGCAGAAAAAAAGGAT
CTCAAGAAGATCCTTTGATCTTTTCTACGGGGTCTGACGCTCAGTGGAACGAAAACTCACG
TTAAGGGATTTTGGTCATGAGATTATCAAAAAGGATCTTCACCTAGATCCTTTTAAATTAAAA
ATGAAGTTTTAAATCAATCTAAAGTATATATGAGTAAACTTGGTCTGACAGTTACCAATGCTT
AATCAGTGAGGCACCTATCTCAGCGATCTGTCTATTTCGTTCATCCATAGTTGCCTGACTCC
CCGTCGTGTAGATAACTACGATACGGGAGGGCTTACCATCTGGCCCCAGTGCTGCAATGA
TACCGCGAGACCCACGCTCACCGGCTCCAGATTTATCAGCAATAAACCAGCCAGCCGGAA
GGGCCGAGCGCAGAAGTGGTCCTGCAACTTTATCCGCCTCCATCCAGTCTATTAATTGTTG

CCGGGAAGCTAGAGTAAGTAGTTCGCCAGTTAATAGTTTGCGCAACGTTGTTGCCATTGCT
ACAGGCATCGTGGTGTCACGCTCGTCGTTTGGTATGGCTTCATTCAGCTCCGGTTCCCAAC
GATCAAGGCGAGTTACATGATCCCCCATGTTGTGCAAAAAAGCGGTTAGCTCCTTCGGTCC
TCCGATCGTTGTCAGAAGTAAGTTGGCCGCAGTGTTATCACTCATGGTTATGGCAGCACTG
CATAATTCTCTTACTGTCATGCCATCCGTAAGATGCTTTTCTGTGACTGGTGAGTACTCAAC
CAAGTCATTCTGAGAATAGTGTATGCGGCGACCGAGTTGCTCTTGCCCGGCGTCAATACG
GGATAATACCGCGCCACATAGCAGAACTTTAAAAGTGCTCATCATTGGAAAACGTTCTTCG
GGGCGAAAACTCTCAAGGATCTTACCGCTGTTGAGATCCAGTTCGATGTAACCCACTCGTG
CACCCAACTGATCTTCAGCATCTTTTACTTTCACCAGCGTTTCTGGGTGAGCAAAAACAGG
AAGGCAAAATGCCGCAAAAAAGGGAATAAGGGCGACACGGAAATGTTGAATACTCATACTC
TTCCTTTTTCAATATTATTGAAGCATTTATCAGGGTTATTGTCTCATGAGCGGATACATATTT
GAATGTATTTAGAAAAATAAACAAATAGGGGTTCCGCGCACATTTCCCCGAAAAGTGCCAC
CTGACGTCTAAGAAACCATTATTATCATGACATTAACCTATAAAAATAGGCGTATCACGAGG
CCCTTTCGTCTTCAAGAATTCTGGCGAATCCTCTGACCAGCCAGAAAACGACCTTTCTGTG
GTGAAACCGGATGCTGCAATTCAGAGCGGCAGCAAGTGGGGGACAGCAGAAGACCTGAC
CGCCGCAGAGTGGATGTTTGACATGGTGAAGACTATCGCACCATCAGCCAGAAAACCGAA
TTTTGCTGGGTGGGCTAACGATATCCGCGGATCCCGACTGATGCGTGAACGTGACGGACG
TAACCACCGCGACATGTGTGTGCTGTTC

**$P_{T7}$-MG aptamer-UTR1-deGFP-T7terminator sequence (pMK18):**

TAATACGACTCACTATAGGGAGAGCTAGCGGGATCCCGACTGGCGAGAGCCAGGTAACGA
ATGGATCCAATAATTTTGTTTAACTTTAAGAAGGAGATATACCATGGAGCTTTTCACTGGCG
TTGTTCCCATCCTGGTCGAGCTGGACGGCGACGTAAACGGCCACAAGTTCAGCGTGTCCG
GCGAGGGCGAGGGCGATGCCACCTACGGCAAGCTGACCCTGAAGTTCATCTGCACCACC
GGCAAGCTGCCCGTGCCCTGGCCCACCCTCGTGACCACCCTGACCTACGGCGTGCAGTG
CTTCAGCCGCTACCCCGACCACATGAAGCAGCACGACTTCTTCAAGTCCGCCATGCCCGA
AGGCTACGTCCAGGAGCGCACCATCTTCTTCAAGGACGACGGCAACTACAAGACCCGCGC
CGAGGTGAAGTTCGAGGGCGACACCCTGGTGAACCGCATCGAGCTGAAGGGCATCGACT
TCAAGGAGGACGGCAACATCCTGGGGCACAAGCTGGAGTACAACTACAACAGCCACAACG
TCTATATCATGGCCGACAAGCAGAAGAACGGCATCAAGGTGAACTTCAAGATCCGCCACAA
CATCGAGGACGGCAGCGTGCAGCTCGCCGACCACTACCAGCAGAACACCCCCATCGGCG
ACGGCCCCGTGCTGCTGCCCGACAACCACTACCTGAGCACCCAGTCCGCCCTGAGCAAA
GACCCCAACGAGAAGCGCGATCACATGGTCCTGCTGGAGTTCGTGACCGCCGCCGGGAT
CTAACTCGAGCTAGCATAACCCCTTGGGGCCTCTAAACGGGTCTTGAGGGGTTTTTTGGTC
GACCGATGCCCTTGAGAGCCTTCAACCCAGTCAGCTCCTTCCGGTGGGCGCGGGGCATG
ACTATCGTCGCCGCACTTATGACTGTCTTCTTTATCATGCAACTCGTAGGACAGGTGCCGG
CAGCGCTCTTCCGCTTCCTCGCTCACTGACTCGCTGCGCTCGGTCGTTCGGCTGCGGCGA
GCGGTATCAGCTCACTCAAAGGCGGTAATACGGTTATCCACAGAATCAGGGGATAACGCA
GGAAAGAACATGTGAGCAAAAGGCCAGCAAAAGGCCAGGAACCGTAAAAAGGCCGCGTTG
CTGGCGTTTTTCCATAGGCTCCGCCCCCCTGACGAGCATCACAAAAATCGACGCTCAAGTC
AGAGGTGGCGAAACCCGACAGGACTATAAAGATACCAGGCGTTTCCCCCTGGAAGCTCCC
TCGTGCGCTCTCCTGTTCCGACCCTGCCGCTTACCGGATACCTGTCCGCCTTTCTCCCTTC
GGGAAGCGTGGCGCTTTCTCAATGCTCACGCTGTAGGTATCTCAGTTCGGTGTAGGTCGT
TCGCTCCAAGCTGGGCTGTGTGCACGAACCCCCCGTTCAGCCCGACCGCTGCGCCTTATC
CGGTAACTATCGTCTTGAGTCCAACCCGGTAAGACACGACTTATCGCCACTGGCAGCAGC
CACTGGTAACAGGATTAGCAGAGCGAGGTATGTAGGCGGTGCTACAGAGTTCTTGAAGTG
GTGGCCTAACTACGGCTACACTAGAAGGACAGTATTTGGTATCTGCGCTCTGCTGAAGCCA
GTTACCTTCGGAAAAAGAGTTGGTAGCTCTTGATCCGGCAAACAAACCACCGCTGGTAGC
GGTGGTTTTTTTGTTTGCAAGCAGCAGATTACGCGCAGAAAAAAAGGATCTCAAGAAGATC
CTTTGATCTTTTCTACGGGGTCTGACGCTCAGTGGAACGAAAACTCACGTTAAGGGATTTT
GGTCATGAGATTATCAAAAAGGATCTTCACCTAGATCCTTTTAAATTAAAAATGAAGTTTTAA
ATCAATCTAAAGTATATATGAGTAAACTTGGTCTGACAGTTACCAATGCTTAATCAGTGAGG
CACCTATCTCAGCGATCTGTCTATTTCGTTCATCCATAGTTGCCTGACTCCCCGTCGTGTAG
ATAACTACGATACGGGAGGGCTTACCATCTGGCCCCAGTGCTGCAATGATACCGCGAGAC
CCACGCTCACCGGCTCCAGATTTATCAGCAATAAACCAGCCAGCCGGAAGGGCCGAGCGC
AGAAGTGGTCCTGCAACTTTATCCGCCTCCATCCAGTCTATTAATTGTTGCCGGGAAGCTA
GAGTAAGTAGTTCGCCAGTTAATAGTTTGCGCAACGTTGTTGCCATTGCTACAGGCATCGT
GGTGTCACGCTCGTCGTTTGGTATGGCTTCATTCAGCTCCGGTTCCCAACGATCAAGGCG
AGTTACATGATCCCCCATGTTGTGCAAAAAAGCGGTTAGCTCCTTCGGTCCTCCGATCGTT
GTCAGAAGTAAGTTGGCCGCAGTGTTATCACTCATGGTTATGGCAGCACTGCATAATTCTC
TTACTGTCATGCCATCCGTAAGATGCTTTTCTGTGACTGGTGAGTACTCAACCAAGTCATTC
TGAGAATAGTGTATGCGGCGACCGAGTTGCTCTTGCCCGGCGTCAATACGGGATAATACC

GCGCCACATAGCAGAACTTTAAAAGTGCTCATCATTGGAAAACGTTCTTCGGGGCGAAAAC
TCTCAAGGATCTTACCGCTGTTGAGATCCAGTTCGATGTAACCCACTCGTGCACCCAACTG
ATCTTCAGCATCTTTTACTTTCACCAGCGTTTCTGGGTGAGCAAAAACAGGAAGGCAAAAT
GCCGCAAAAAAGGGAATAAGGGCGACACGGAAATGTTGAATACTCATACTCTTCCTTTTTC
AATATTATTGAAGCATTTATCAGGGTTATTGTCTCATGAGCGGATACATATTTGAATGTATTT
AGAAAAATAAACAAATAGGGGTTCCGCGCACATTTCCCCGAAAAGTGCCACCTGACGTCTA
AGAAACCATTATTATCATGACATTAACCTATAAAAATAGGCGTATCACGAGGCCCTTTCGTC
TTCAAGAATTCTGGCGAATCCTCTGACCAGCCAGAAAACGACCTTTCTGTGGTGAAACCGG
ATGCTGCAATTCAGAGCGGCAGCAAGTGGGGGACAGCAGAAGACCTGACCGCCGCAGAG
TGGATGTTTGACATGGTGAAGACTATCGCACCATCAGCCAGAAAACCGAATTTTGCTGGGT
GGGCTAACGATATCCGCGGATCCCGACTGATGCGTGAACGTGACGGACGTAACCACCGC
GACATGTGTGTGCTGTTC

**$P_{Tet}$-MG aptamer-UTR1-deGFP-T500 sequence (pMK19):**

TCCCTATCAGTGATAGAGATTGACATCCCTATCAGTGATAGAGATACTGAGCACGCTAGCG
GGATCCCGACTGGCGAGAGCCAGGTAACGAATGGATCCAATAATTTTGTTTAACTTTAAGA
AGGAGATATACCATGGAGCTTTTCACTGGCGTTGTTCCCATCCTGGTCGAGCTGGACGGC
GACGTAAACGGCCACAAGTTCAGCGTGTCCGGCGAGGGCGAGGGCGATGCCACCTACGG
CAAGCTGACCCTGAAGTTCATCTGCACCACCGGCAAGCTGCCCGTGCCCTGGCCCACCCT
CGTGACCACCCTGACCTACGGCGTGCAGTGCTTCAGCCGCTACCCCGACCACATGAAGCA
GCACGACTTCTTCAAGTCCGCCATGCCCGAAGGCTACGTCCAGGAGCGCACCATCTTCTT
CAAGGACGACGGCAACTACAAGACCCGCGCCGAGGTGAAGTTCGAGGGCGACACCCTGG
TGAACCGCATCGAGCTGAAGGGCATCGACTTCAAGGAGGACGGCAACATCCTGGGGCAC
AAGCTGGAGTACAACTACAACAGCCACAACGTCTATATCATGGCCGACAAGCAGAAGAAC
GGCATCAAGGTGAACTTCAAGATCCGCCACAACATCGAGGACGGCAGCGTGCAGCTCGCC
GACCACTACCAGCAGAACACCCCCATCGGCGACGGCCCCGTGCTGCTGCCCGACAACCA
CTACCTGAGCACCCAGTCCGCCCTGAGCAAAGACCCCAACGAGAAGCGCGATCACATGGT
CCTGCTGGAGTTCGTGACCGCCGCCGGGATCTAACTCGAGCAAAGCCCGCCGAAAGGCG
GGCTTTTCTGTGTCGACCGATGCCCTTGAGAGCCTTCAACCCAGTCAGCTCCTTCCGGTG
GGCGCGGGGCATGACTATCGTCGCCGCACTTATGACTGTCTTCTTTATCATGCAACTCGTA
GGACAGGTGCCGGCAGCGCTCTTCCGCTTCCTCGCTCACTGACTCGCTGCGCTCGGTCGT
TCGGCTGCGGCGAGCGGTATCAGCTCACTCAAAGGCGGTAATACGGTTATCCACAGAATC
AGGGGATAACGCAGGAAAGAACATGTGAGCAAAAGGCCAGCAAAAGGCCAGGAACCGTAA
AAAGGCCGCGTTGCTGGCGTTTTTCCATAGGCTCCGCCCCCCTGACGAGCATCACAAAAA
TCGACGCTCAAGTCAGAGGTGGCGAAACCCGACAGGACTATAAAGATACCAGGCGTTTCC
CCCTGGAAGCTCCCTCGTGCGCTCTCCTGTTCCGACCCTGCCGCTTACCGGATACCTGTC
CGCCTTTCTCCCTTCGGGAAGCGTGGCGCTTTCTCAATGCTCACGCTGTAGGTATCTCAGT
TCGGTGTAGGTCGTTCGCTCCAAGCTGGGCTGTGTGCACGAACCCCCCGTTCAGCCCGAC
CGCTGCGCCTTATCCGGTAACTATCGTCTTGAGTCCAACCCGGTAAGACACGACTTATCGC
CACTGGCAGCAGCCACTGGTAACAGGATTAGCAGAGCGAGGTATGTAGGCGGTGCTACAG
AGTTCTTGAAGTGGTGGCCTAACTACGGCTACACTAGAAGGACAGTATTTGGTATCTGCGC
TCTGCTGAAGCCAGTTACCTTCGGAAAAAGAGTTGGTAGCTCTTGATCCGGCAAACAAACC
ACCGCTGGTAGCGGTGGTTTTTTTGTTTGCAAGCAGCAGATTACGCGCAGAAAAAAAGGAT
CTCAAGAAGATCCTTTGATCTTTTCTACGGGGTCTGACGCTCAGTGGAACGAAAACTCACG
TTAAGGGATTTTGGTCATGAGATTATCAAAAAGGATCTTCACCTAGATCCTTTTAAATTAAAA
ATGAAGTTTTAAATCAATCTAAAGTATATATGAGTAAACTTGGTCTGACAGTTACCAATGCTT
AATCAGTGAGGCACCTATCTCAGCGATCTGTCTATTTCGTTCATCCATAGTTGCCTGACTCC
CCGTCGTGTAGATAACTACGATACGGGAGGGCTTACCATCTGGCCCCAGTGCTGCAATGA
TACCGCGAGACCCACGCTCACCGGCTCCAGATTTATCAGCAATAAACCAGCCAGCCGGAA
GGGCCGAGCGCAGAAGTGGTCCTGCAACTTTATCCGCCTCCATCCAGTCTATTAATTGTTG
CCGGGAAGCTAGAGTAAGTAGTTCGCCAGTTAATAGTTTGCGCAACGTTGTTGCCATTGCT
ACAGGCATCGTGGTGTCACGCTCGTCGTTTGGTATGGCTTCATTCAGCTCCGGTTCCCAAC
GATCAAGGCGAGTTACATGATCCCCCATGTTGTGCAAAAAAGCGGTTAGCTCCTTCGGTCC
TCCGATCGTTGTCAGAAGTAAGTTGGCCGCAGTGTTATCACTCATGGTTATGGCAGCACTG
CATAATTCTCTTACTGTCATGCCATCCGTAAGATGCTTTTCTGTGACTGGTGAGTACTCAAC
CAAGTCATTCTGAGAATAGTGTATGCGGCGACCGAGTTGCTCTTGCCCGGCGTCAATACG

GGATAATACCGCGCCACATAGCAGAACTTTAAAAGTGCTCATCATTGGAAAACGTTCTTCG
GGGCGAAAACTCTCAAGGATCTTACCGCTGTTGAGATCCAGTTCGATGTAACCCACTCGTG
CACCCAACTGATCTTCAGCATCTTTTACTTTCACCAGCGTTTCTGGGTGAGCAAAAACAGG
AAGGCAAAATGCCGCAAAAAAGGGAATAAGGGCGACACGGAAATGTTGAATACTCATACTC
TTCCTTTTTCAATATTATTGAAGCATTTATCAGGGTTATTGTCTCATGAGCGGATACATATTT
GAATGTATTTAGAAAAATAAACAAATAGGGGTTCCGCGCACATTTCCCCGAAAAGTGCCAC
CTGACGTCTAAGAAACCATTATTATCATGACATTAACCTATAAAAATAGGCGTATCACGAGG
CCCTTTCGTCTTCAAGAATTCTGGCGAATCCTCTGACCAGCCAGAAAACGACCTTTCTGTG
GTGAAACCGGATGCTGCAATTCAGAGCGGCAGCAAGTGGGGGACAGCAGAAGACCTGAC
CGCCGCAGAGTGGATGTTTGACATGGTGAAGACTATCGCACCATCAGCCAGAAAACCGAA
TTTTGCTGGGTGGGCTAACGATATCCGCGGATCCCGACTGATGCGTGAACGTGACGGACG
TAACCACCGCGACATGTGTGTGCTGTTC

**pTet-F30Pepper-UTR1-mTurquoise2-ECK120029600 sequence (pMK10):**

Here the F30 scaffold that flanks either side of the Pepper aptamer is highlighted in orange.

TCCCTATCAGTGATAGAGATTGACATCCCTATCAGTGATAGAGATACTGAGCACTACTTTGC
CATGTGTATGTGGGTTCGCCCACATACTCTGATGATCCCCAATCGTGGCGTGTCGGCCTG
CTTCGGCAGGCACTGGCGCCGGGATCATTCATGGCAAAATAATTTTGTTTAACTTTAAGAA
GGAGATATACCAATGGTAAGTAAGGGCGAAGAGTTGTTTACTGGGGTAGTCCCGATCCTG
GTCGAGCTGGATGGGGACGTTAACGGCCATAAATTTAGCGTGTCGGGTGAAGGCGAGGG
GGACGCGACCTACGGGAAACTGACCTTGAAGTTTATCTGTACGACCGGGAAACTTCCGGT
GCCATGGCCTACACTCGTCACGACCTTGAGCTGGGGTGTTCAATGCTTTGCACGTTACCCA
GACCATATGAAACAGCATGATTTCTTCAAAAGTGCAATGCCAGAAGGGTATGTTCAAGAGC
GCACCATCTTTTTTAAGGACGATGGCAACTATAAGACCCGTGCCGAAGTCAAGTTCGAAGG
TGACACCCTTGTCAACCGTATCGAGTTGAAAGGTATCGACTTTAAGGAAGATGGGAATATC
CTGGGTCATAAACTTGAATATAATTACTTTTCCGACAATGTTTATATCACAGCCGACAAACA
AAAAAACGGTATCAAGGCCAATTTCAAGATTCGGCATAACATCGAAGATGGGGGGTTCAA
CTCGCTGATCATTACCAGCAGAATACACCTATCGGCGACGGCCCAGTCTTATTACCTGATA
ATCATTACCTTAGCACCCAATCCAAACTCAGCAAGGATCCAAACGAAAAGCGCGATCACAT
GGTGCTTTTGGAGTTTGTCACTGCTGCTGGTATTACATTGGGTATGGATGAATTGTACAAA
GGGGGCGGCGGTAGCCATCATCATCATCACCATTAAAGGTTTCAGCCAAAAAACTTAAGAC
CGCCGGTCTTGTCCACTACCTTGCAGTAATGCGGTGGACAGGATCGGCGGTTTTCTTTTCT
CTTCTCAAGCTTATGTCTTCTACTAGTAGCGGCCGCTGCAGTCCGGCAAAAAAGGGCAAG
GTGTCACCACCCTGCCCTTTTTCTTTAAAACCGAAAAGATTACTTCGCGTTATGCAGGCTTC
CTCGCTCACTGACTCGCTGCGCTCGGTCGTTCGGCTGCGGCGAGCGGTATCAGCTCACTC
AAAGGCGGTAAGTCGCGTGTTATACGCCCGTTATCCATGGGTATGGACAGTTTTCCCTTTG
ATATGTAACGCACGTTGTGTCTCAAAATCTCTGATGTTACATTGCACAAGATAAAAATATATC
ATCATGAACAATAAAACTGTCTGCTTACATAAACAGTAATACAAGGGGTGTTATGAGCCATA
TTCAACGGGAAACGTCTTGCTCCCGTCCGCGCTTAAACTCCAACATGGACGCTGATTTATA
TGGGTATAAATGGGCTCGCGATAATGTCGGGCAATCAGGTGCGACAATCTATCGCTTGTAT
GGGAAGCCCGATGCGCCAGAGTTGTTTCTGAAACATGGCAAAGGTAGCGTTGCCAATGAT
GTTACAGATGAGATGGTCCGTCTCAACTGGCTGACGGAGTTTATGCCTCTCCCGACCATCA
AGCATTTTATCCGTACTCCTGATGATGCGTGGTTACTCACCACCGCGATTCCTGGGAAAAC
AGCCTTCCAGGTATTAGAAGAATATCCTGATTCAGGTGAAAATATTGTTGATGCGCTGGCC
GTGTTCCTGCGCCGGTTACATTCGATTCCTGTTTGTAATTGTCCTTTTAACAGCGATCGTGT
ATTTCGTCTTGCTCAGGCGCAATCACGCATGAATAACGGTTTGGTTGATGCGAGTGATTTT
GATGACGAGCGTAATGGCTGGCCTGTTGAACAAGTCTGGAAAGAAATGCACAAGCTCTTG
CCATTCTCACCGGATTCAGTCGTCACTCATGGTGATTTCTCACTTGATAACCTTATTTTTGA
CGAGGGGAAATTAATAGGTTGTATTGATGTTGGACGGGTCGGAATCGCAGACCGTTACCA
GGACCTTGCCATTCTTTGGAACTGCCTCGGTGAGTTTTCTCCTTCATTACAGAAACGGCTTT
TTCAAAAATATGGTATTGATAATCCTGATATGAATAAATTGCAGTTTCATTTGATGCTCGATG
AGTTTTTCTAATAATACTAGCAGAAATCATCCTTAGCGAAAGCTAAGGATTTTTTTTATCTGA
TTACCGCCTTTGAGTGAGCGTCGACCTAGTCCTAGCCAGGAACCGTAAAAAGGCCGCGTT
GCTGGCGTTTTTCCATAGGCTCCGCCCCCCTGACGAGCATCACAAAAATCGACGCTCAAG
TCAGAGGTGGCGAAACCCGACAGGACTATAAAGATACCAGGCGTTTCCCCCTGGAAGCTC

CCTCGTGCGCTCTCCTGTTCCGACCCTGCCGCTTACCGGATACCTGTCCGCCTTTCTCCCT
TCGGGAAGCGTGGCGCTTTCTCATAGCTCACGCTGTAGGTATCTCAGTTCGGTGTAGGTC
GTTCGCTCCAAGCTGGGCTGTGTGCACGAACCCCCCGTTCAGCCCGACCGCTGCGCCTTA
TCCGGTAACTATCGTCTTGAGTCCAACCCGGTAAGACACGACTTATCGCCACTGGCAGCA
GCCACTGGTAACAGGATTAGCAGAGCGAGGTATGTAGGCGGTGCTACAGAGTTCTTGAAG
TGGTGGCCTAACTACGGCTACACTAGAAGAACAGTATTTGGTATCTGCGCTCTGCTGAAGC
CAGTTACCTTCGGAAAAAGAGTTGGTAGCTCTTGATCCGGCAAACAAACCACCGCTGGTAG
CGGTGGTTTTTTTGTTTGCAAGCAGCAGATTACGCGCAGAAAAAAAGGATCTCAAGAAGAT
CCTTTGATCTTTTCTACGGGGTCTGACGCGCATAAATAGGCGTATCACGAGGCAGAATTTC
AGATAAAAAAAATCCTTAGCTTTCGCTAAGGATGATTTCTGGAATTCGCGGCCGCTTCTAGA
GACTAGTGGAAGACATGGAG
