## Supplementary material for "Metabolic perturbations to an *E. coli*-based cell-free system reveal a trade-off between transcription and translation": Raw Data, Tidy Data, DNA GenBank files, Figures, and Jupyter Notebooks for Figure Generation and Analysis: 20220224_SaltCalibration_AR_ZJ_Extract_w_PEG_and_Minimal_Energy_Solution.ipynb

```
# %% [code]
# analysis packages
import pandas as pd
import numpy as np

# plotting
import bokeh.io
import bokeh.plotting
from bokeh.resources import INLINE
from bokeh.themes import Theme
from bokeh.models import ColumnDataSource, Grid, LinearAxis, Plot, Text, Band, LogColorMapper, LogTicker, ColorBar, ColorMapper, Div, Label
from bokeh.layouts import gridplot, layout, row, column, Spacer
import bokeh.palettes
import seaborn as sns
import matplotlib as mpl
from cycler import cycler
import holoviews as hv
from holoviews.operation.timeseries import rolling
from bokeh.layouts import row
from base64 import b64encode 
from bokeh.models.tickers import FixedTicker
import plotly
import plotly.graph_objs as go
from plotly.offline import init_notebook_mode
import iqplot
init_notebook_mode()


bokeh.io.output_notebook(INLINE)
hv.extension("bokeh")

# %% [markdown]
## get data

# %% [code]
# file location
file_loc = "../Raw Data/20220224_SaltCalibration_AR_ZJ_Extract_w_PEG_and_Minimal_Energy_Solution/20220224_SaltCalibration_AR_ZJ_Extract_w_PEG_and_Minimal_Energy_Solution_Results.csv"
image_name = file_loc.split("/")[2].split('.')[0]
image_save = file_loc.split(file_loc.split("/")[3].split('.')[0])[0]
# read as dataframe
df = pd.read_csv(filepath_or_buffer=file_loc, sep=',', header=61)
# parameters of run
run_length_hrs = 16
interval = 5
n_rows = int(run_length_hrs*60/interval + 1)
# drop unnecessary rows and columns
df = df.drop(columns=[ x for x in df.columns if "Unnamed" in x ])
df = df.drop(columns=['Tｰ deGFP:485,515'])
df = df[0:n_rows]
# drop unnamed columns
df = df.drop(columns=[ x for x in df.columns if "Unnamed" in x ])
df.tail()

# %% [code]
# melt dataframe
df_melted = df.loc[df['Time'] == '16:00:06'].drop(columns=['Time']).melt()
df_melted['value'] = df_melted['value'].astype('float')


# add columns for Mg and K concentrations
Mg_vals = [0, 2, 4, 6, 8, 10]
K_vals = [50, 80, 110, 140, 170, 200]
K_vals = list(np.round(np.array(K_vals)))
Mg_list = []
K_list = []
for this_Mg in Mg_vals:
    for this_K in K_vals:
        Mg_list.append(this_Mg)
        K_list.append(this_K)
df_melted['Mg (mM)'] = Mg_list
df_melted['K (mM)'] = K_list
df_melted['value'] = df_melted['value']/1811.16
df_melted.head()

# %% [code]
# axis labels
new_group = df_melted.copy()
new_group['Mg (mM)'] = new_group['Mg (mM)'].astype('str')
new_group['K (mM)'] = new_group['K (mM)'].astype('str')
new_group['K (mM)'] = pd.Categorical(new_group['K (mM)'],categories=[str(i) for i in K_vals])
new_group['Mg (mM)'] = pd.Categorical(new_group['Mg (mM)'],categories=[str(i) for i in Mg_vals])
new_group = new_group.sort_values(['K (mM)', "Mg (mM)"])

# define rounding
value_dimension = hv.Dimension('value', value_format=lambda x: '%.1f' % x)  

# create holoviews heatmap
hv_fig = hv.HeatMap(data=new_group,
                    kdims=['Mg (mM)', 'K (mM)'],
                    vdims=['value']
).opts(
    frame_height=750,
    frame_width=750,
    colorbar=True,
    show_grid=False,
    cmap='Viridis',
    # title = f'Integrated MG aptamer (μM-hr)',
    # fontsize={'labels': 35, 'xticks': 30, 'yticks': 30, 'ticks':30}
    fontsize={'labels': 50, 'xticks': 45, 'yticks': 45, 'ticks':45, 'title':50}
)

# add numbers on top of grid
hv_fig = hv_fig * hv.Labels(hv_fig, vdims=value_dimension).opts(text_color='white', text_font_size='35px',fontsize={'ticks':20})

# render as a bokeh figure, make axes non-italic font
p = hv.render(hv_fig)
p.xaxis.axis_label_text_font_style = 'normal'
p.yaxis.axis_label_text_font_style = 'normal'
p.toolbar_location = None
bokeh.io.show(p)

# %% [code]
# save plot
file_save = '..\\Figures\\FigS15_a.png'
bokeh.io.reset_output()
# bokeh.io.curdoc().theme = theme
bokeh.io.export_png(p, filename = file_save)
```
