## Supplementary material for "Metabolic perturbations to an *E. coli*-based cell-free system reveal a trade-off between transcription and translation": Raw Data, Tidy Data, DNA GenBank files, Figures, and Jupyter Notebooks for Figure Generation and Analysis: 20230203_deGFP_stability_Mg_pH.ipynb

```
# %% [markdown]
# Tidy and Plot Data 

# %% [markdown]
In this experiment, we look at the results of Mg+2 and pH on deGFP fluorescence. Briefly, 0.935uM purified deGFP (from PEC) was added to pH-ed PBS solutions.

# %% [code]
# data analysis and computing packages
import numpy as np
import panel as pn
import scipy as sp
import pandas as pd
from datetime import datetime as dt
from datetime import timedelta
import csv
from itertools import permutations
from ast import literal_eval as make_tuple


# plotting
import bokeh.io
import bokeh.plotting
from bokeh.resources import INLINE
from bokeh.themes import Theme
from bokeh.models import ColumnDataSource, Grid, LinearAxis, Plot, Text, Band
import bokeh.palettes
import seaborn as sns
import matplotlib as mpl
from cycler import cycler
import holoviews as hv
from holoviews.operation.timeseries import rolling
from bokeh.layouts import row

# echo tools
import murraylab_tools.biotek as mt_biotek
import os

bokeh.io.output_notebook(INLINE)
hv.extension("bokeh")

# %% [code]
theme = Theme(json={'attrs': {
    
# apply defaults to Figure properties
'Figure': {
    'toolbar_location': 'right',
    'outline_line_color': None,
    'min_border_right': 10,
    'height':1000,
    'width':1500,
},    
    
# apply defaults to Grid properties
'Grid': {
    'grid_line_color': None,
},
    
# apply defaults to Title properties
'Title': {
    'text_font_size': '30pt',
    'align': 'center'
},
    
# # apply defaults to Plot properties
# 'Plot': {
#     'renderers': {'add_layout': {'legend[0]':'right'}},
# },
    
# apply defaults to Axis properties
'Axis': {
    'major_label_text_font_size': '25pt',
    'axis_label_text_font_size': '25pt',
    'axis_label_text_font_style': 'normal',
    'axis_label_standoff':30
},

# apply defaults to Legend properties
'Legend': {
    'background_fill_alpha': 0.8,
    'location': 'top_right',
    "label_text_font_size": '15pt',
    "click_policy": 'hide',
    "title_text_font_style": 'normal',
    "title_text_font_size": '18pt'
}}})

bokeh.io.curdoc().theme = theme
hv.renderer('bokeh').theme = theme

# %% [markdown]
# 

# %% [markdown]
## Tidy data

# %% [code]
# list of all relevant experiments
expt_lst = [
            '20230203_deGFP_stability_Mg_pH'
           ]
file_loc = "../Raw Data/"

# %% [code]
# load calibration data
calibration_dict = mt_biotek.calibration_data(filename = 'calibration.csv')
calibration_dict

# %% [code]
# iterate through experiments, make each one tidy :)
for this_expt in expt_lst:
    # specify input and output folders
    inputfolder = os.path.join(file_loc, this_expt)
    outputfolder = os.path.join(file_loc, this_expt)
    # specify file names
    data_name = this_expt
    data_filename = os.path.join(inputfolder, data_name+"_Results.csv")
    tidy_filename = os.path.join(outputfolder,data_name+"_Results_tidy.csv")
    # specify supplementary file containing metadata on what conditions pertain to each well
    supplementary_filename = os.path.join(inputfolder, data_name+'_supplementary_info.csv')
    # provide calibration data if available
    # calibration = mt_biotek.calibration_data("10/50/14")
    print(this_expt)
    mt_biotek.tidy_biotek_data(data_filename, 
                               supplementary_filename,
                               volume = 10, 
                               calibration_dict = calibration_dict,
                               convert_to_uM = True,
                               override_plate_reader_id='b1',
                               output_filename = tidy_filename)

# %% [code]
# read tidy dataframe
tidy_csv_loc = os.path.join(file_loc, this_expt,this_expt+"_Results_tidy.csv")
df = pd.read_csv(tidy_csv_loc)
df.head()

# %% [markdown]
# 

# %% [markdown]
## Plot results

# %% [code]
# endpoint dataframe
temp_df = df.loc[df['Time (sec)'] == 64806]

# drop unrelated columns, create new df
temp_df = temp_df.drop(columns=['Well'])
cols = list(temp_df.columns)
cols.remove('Replicate')
cols.remove('Measurement')
new_group = temp_df.groupby(cols).apply(lambda group2: np.mean(group2['Measurement'])).reset_index().fillna(method="ffill")
new_group = pd.DataFrame(new_group).rename({0:'Measurement'}, axis=1)

# make Mg and pH have string values so heatmap has categorical axes
new_group['Mg concentration (mM)'] = new_group['Mg concentration (mM)'].astype('str')
new_group['pH'] = new_group['pH'].astype('str')


# create holoviews heatmap
value_dimension = hv.Dimension('Measurement', value_format=lambda x: '%.2f' % x)    

# hv_fig = hv.HeatMap(data=new_group,
#                     kdims=['pH', ('Mg concentration (mM)', 'Added Mg⁺² (mM)') ],
#                     vdims=['Measurement']
# ).opts(
#     frame_height=500,
#     frame_width=1084,
#     colorbar=True,
#     show_grid=False,
#     cmap='Viridis',
#     title = 'Dependence of deGFP fluorescence on pH and Mg⁺²',
#     colorbar_opts={
#                    'title':'deGFP (uM)', 
#                    'title_text_font_style':'bold', 
#                    'title_text_font_size':'200px', 
#                    'width':200, 
#                    'title_text_align': 'left', 
#                    'major_tick_out':4,
#                    'title_standoff':10,
#                    'bar_line_width': 0,
#                    'orientation':'horizontal',
#                    'location': 'top_right',
#                    'margin': 200,
#                    'title_text_line_height':200,
#                    'padding':50
#                   }
# )
# hv_fig = hv_fig * hv.Labels(hv_fig, vdims=value_dimension).opts(text_color='white', text_font_size='24px',fontsize={'ticks':20})

# # render as bokeh figure, format
# hv_fig = hv.render(hv_fig)
# hv_fig.toolbar_location = None

# create holoviews heatmap
value_dimension = hv.Dimension('Measurement', value_format=lambda x: '%.2f' % x)    

hv_fig = hv.HeatMap(data=new_group,
                    kdims=['pH', ('Mg concentration (mM)', 'Added Mg²⁺ (mM)') ],
                    vdims=['Measurement']
).opts(
    frame_height=500,
    frame_width=1000,
    colorbar=True,
    show_grid=False,
    cmap='Viridis',
    title = f'Measured deGFP (μM)',
)
hv_fig = hv_fig * hv.Labels(hv_fig, vdims=value_dimension).opts(text_color='white', text_font_size='20px',fontsize={'ticks':20})

# render as bokeh figure, format
hv_fig = hv.render(hv_fig)
hv_fig.toolbar_location = None

# save plot
file_save = '../Figures/FigS1_b.png'
bokeh.io.reset_output()
bokeh.io.curdoc().theme = theme
bokeh.io.export_png(hv_fig, filename = file_save)
```
