## Supplementary material for "Metabolic perturbations to an *E. coli*-based cell-free system reveal a trade-off between transcription and translation": Raw Data, Tidy Data, DNA GenBank files, Figures, and Jupyter Notebooks for Figure Generation and Analysis: 20230228_TX_only_w_Tetracycline.ipynb

```
# %% [markdown]
# Tidy and Plot Data 

# %% [markdown]
In this experiment, we create a dose-response curve for TX-TL solutions with tetracycline to inhibit translation. This time, we use 80mg/mL tetracycline dissolved in 100% DMSO.

# %% [code]
# data analysis and computing packages
import numpy as np
import panel as pn
import scipy as sp
import pandas as pd
from datetime import datetime as dt
from datetime import timedelta
import csv
from itertools import permutations


# plotting
import bokeh.io
import bokeh.plotting
from bokeh.resources import INLINE
from bokeh.themes import Theme
from bokeh.models import ColumnDataSource, Grid, LinearAxis, Plot, Text, Band
import bokeh.palettes
import seaborn as sns
import matplotlib as mpl
from cycler import cycler
import holoviews as hv
from holoviews.operation.timeseries import rolling
from bokeh.layouts import row

# echo tools
import murraylab_tools.biotek as mt_biotek
import os

bokeh.io.output_notebook(INLINE)
hv.extension("bokeh")

# %% [code]
theme = Theme(json={'attrs': {
    
# apply defaults to Figure properties
'Figure': {
    'toolbar_location': 'right',
    'outline_line_color': None,
    'min_border_right': 10,
    'height':1000,
    'width':1500,
},    
    
# apply defaults to Grid properties
'Grid': {
    'grid_line_color': None,
},
    
# apply defaults to Title properties
'Title': {
    'text_font_size': '30pt',
    'align': 'center'
},
    
# # apply defaults to Plot properties
# 'Plot': {
#     'renderers': {'add_layout': {'legend[0]':'right'}},
# },
    
# apply defaults to Axis properties
'Axis': {
    'major_label_text_font_size': '50pt',
    'axis_label_text_font_size': '55pt',
    'axis_label_text_font_style': 'normal',
    'axis_label_standoff':30
},

# apply defaults to Legend properties
'Legend': {
    'background_fill_alpha': 0.8,
    'location': 'top_right',
    "label_text_font_size": '15pt',
    "click_policy": 'hide',
    "title_text_font_style": 'normal',
    "title_text_font_size": '36pt'
}}})

bokeh.io.curdoc().theme = theme
hv.renderer('bokeh').theme = theme

# %% [markdown]
# 

# %% [markdown]
## Tidy data

# %% [code]
# list of all relevant experiments
expt_lst = [
            '20230228_TX_only_w_Tetracycline'
           ]
file_loc = "../Raw Data/"

# %% [code]
# load calibration data
calibration_dict = mt_biotek.calibration_data(filename = 'calibration.csv')
calibration_dict

# %% [code]
# iterate through experiments, make each one tidy :)
for this_expt in expt_lst:
    # specify input and output folders
    inputfolder = os.path.join(file_loc, this_expt)
    outputfolder = os.path.join(file_loc, this_expt)
    # specify file names
    data_name = this_expt
    data_filename = os.path.join(inputfolder, data_name+"_Results.csv")
    tidy_filename = os.path.join(outputfolder,data_name+"_Results_tidy.csv")
    # specify supplementary file containing metadata on what conditions pertain to each well
    supplementary_filename = os.path.join(inputfolder, data_name+'_supplementary_info.csv')
    # provide calibration data if available
    # calibration = mt_biotek.calibration_data("10/50/14")
    print(this_expt)
    mt_biotek.tidy_biotek_data(data_filename, 
                               supplementary_filename,
                               volume = 10, 
                               calibration_dict = calibration_dict,
                               convert_to_uM = True,
                               override_plate_reader_id='b1',
                               output_filename = tidy_filename)

# %% [code]
# read tidy dataframe
tidy_csv_loc = os.path.join(file_loc, this_expt,this_expt+"_Results_tidy.csv")
df = pd.read_csv(tidy_csv_loc)
df.head()

# %% [markdown]
# 

# %% [markdown]
## Plot results

# %% [code]
def sample_averaged_timecourse_degfp(group):
    '''
    Takes in a dataframe, fixed variable, and changing variable. Returns a bokeh plot of time course data of deGFP dynamics with error band
    
    inputs
    ------
    df : Pandas dataframe
        Dataframe containing fluorescence time course data
    
    outputs
    -------
    p : Bokeh.plotting.figure instance
        A plot of the time course data
    '''
    
    # define variables
    changing_var = 'Tetratcycline concentration (ug/mL)'
    n_colors = 6
    
    # plot time rolling average of 3 replicates
    hv_fig1 = hv.Curve(
        data=group,
        kdims=[("Time (hr)", "Time (hrs)")],
        vdims=[('Measurement', 'deGFP (μM)'), 'Replicate', changing_var],
    ).groupby([changing_var, 'Replicate']
    ).collapse('Replicate',np.mean
    ).opts(
        color=hv.Cycle(list(bokeh.palettes.Viridis[n_colors])),
        frame_height=500,
        frame_width=750,
        line_width=4,
        line_color=hv.Cycle(list(bokeh.palettes.Viridis[n_colors]))
    ).overlay(
    ).opts(
        legend_position="right",
        legend_offset=(10, 300),
        legend_opts={'click_policy':'hide',
                    },
    ) 
    hv_fig1 = rolling(hv_fig1, rolling_window=10)
    
    
    # plot all data also
    hv_fig2 = hv.Points(
        data=group,
        kdims=[("Time (hr)", "Time (hrs)"), ('Measurement', 'deGFP (μM)')],
        vdims=[changing_var],
    ).groupby([changing_var]
    ).opts(
        color=hv.Cycle(list(bokeh.palettes.Viridis[n_colors])),
        size=10,
        frame_height=1000,
        frame_width=1200,
        line_color=hv.Cycle(list(bokeh.palettes.Viridis[n_colors])),
        alpha=1,
        xlim=(-0.1,18),
        ylim=(-0.1, 18)
    ).overlay(
    )
    
    # create bokeh figure from holoviews figure
    hv_fig = hv_fig1*hv_fig2
    p = hv.render(hv_fig2)

    p.legend.title = changing_var
    p.legend.label_text_font_size = '35pt'
    p.legend.title_text_font_size='40pt'
    p.legend.title_text_font_style='normal'
    p.legend.glyph_width=50
    p.legend.glyph_height=50

    return(p)

# %% [code]
def sample_averaged_timecourse_mgaptamer(group):
    '''
    Takes in a dataframe, fixed variable, and changing variable. Returns a bokeh plot of time course data of deGFP dynamics with error band
    
    inputs
    ------
    df : Pandas dataframe
        Dataframe containing fluorescence time course data
    
    outputs
    -------
    p : Bokeh.plotting.figure instance
        A plot of the time course data
    '''

    # define variables
    changing_var = 'Tetratcycline concentration (ug/mL)'
    n_colors = 6
    
    # plot time rolling average of 3 replicates
    hv_fig1 = hv.Curve(
        data=group,
        kdims=[("Time (hr)", "Time (hrs)")],
        vdims=[('Measurement', 'MGaptamer (μM)'), 'Replicate', changing_var],
    ).groupby([changing_var, 'Replicate']
    ).collapse('Replicate', np.mean
    ).opts(
        color=hv.Cycle(list(bokeh.palettes.Viridis[n_colors])),
        frame_height=500,
        frame_width=750,
        line_width=4,
        line_color=hv.Cycle(list(bokeh.palettes.Viridis[n_colors]))
    ).overlay(
    ).opts(
        legend_position="right",
        legend_offset=(10, 300),
        legend_opts={'click_policy':'hide',
                    },
    ) 
    hv_fig1 = rolling(hv_fig1, rolling_window=10)
    
    
    # plot all data also
    hv_fig2 = hv.Points(
        data=group,
        kdims=[("Time (hr)", "Time (hrs)"), ('Measurement', 'MGaptamer (μM)')],
        vdims=[changing_var],
    ).groupby([changing_var]
    ).opts(
        color=hv.Cycle(list(bokeh.palettes.Viridis[n_colors])),
        size=10,
        frame_height=1000,
        frame_width=1200,
        line_color=hv.Cycle(list(bokeh.palettes.Viridis[n_colors])),
        alpha=1,
        xlim=(-0.1,18),
        ylim=(0, 0.6)
    ).overlay(
    )
    
    # create bokeh figure from holoviews figure
    hv_fig = hv_fig1*hv_fig2
    p = hv.render(hv_fig2)

    p.legend.title = changing_var
    p.legend.label_text_font_size = '35pt'
    p.legend.title_text_font_size='40pt'
    p.legend.title_text_font_style='normal'
    p.legend.glyph_width=50
    p.legend.glyph_height=50

    return(p)

# %% [code]
# partition dataframe into separate ones for extracts and measurement type
df_groups = df.groupby(['Extract', 'Channel'])

# iterate through groups, make plots
suffixes = ['a', 'b']
j = 0
for name, group in df_groups:
    
    # extract info from name
    extract = name[0]
    channel = name[1]
    
    # make plots
    if channel == 'MGaptamer':
        
        # plot
        p = sample_averaged_timecourse_mgaptamer(group)
        # p.title.text = 'MGaptamer expression under ribosome inhibition'
        
    else:
        # plot
        p = sample_averaged_timecourse_degfp(group)
        # p.title.text = 'deGFP expression under ribosome inhibition'
          
        
    # add plot customizations
    p.legend.title = 'Tetracycline (μg/mL)'
    p.legend.click_policy="hide"
    p.toolbar_location = None
    p.title.text_font_size = '30pt'
    p.title.align = 'center'
    p.legend.title_text_font_style = 'normal'
    p.legend.title_text_font_size = '40pt'

    # save plot
    file_save = f'..//Figures/FigS14_{suffixes[j]}.png'
    bokeh.io.reset_output()
    bokeh.io.curdoc().theme = theme
    bokeh.io.export_png(p, filename = file_save)

    j += 1
```
