## Supplementary material for "Metabolic perturbations to an *E. coli*-based cell-free system reveal a trade-off between transcription and translation": Raw Data, Tidy Data, DNA GenBank files, Figures, and Jupyter Notebooks for Figure Generation and Analysis: Analysis_Main_Figures.ipynb

```
# %% [markdown]
# Main Figures

# %% [markdown]
### Import packages

# %% [code]
# data analysis and computing packages
import numpy as np
import panel as pn
import scipy as sp
from sklearn.utils import resample
from sklearn.linear_model import LinearRegression
import scipy.stats as stats
import pandas as pd
from datetime import datetime as dt
from datetime import timedelta
import csv
from itertools import permutations
from PIL import Image
import murraylab_tools.biotek as mt_biotek
from sklearn.cross_decomposition import PLSCanonical


# plotting
import bokeh.io
import bokeh.plotting
from bokeh.resources import INLINE
from bokeh.themes import Theme
from bokeh.models import ColumnDataSource, Grid, LinearAxis, Plot, Text, Band, LogColorMapper, LogTicker, ColorBar, ColorMapper, Div, Label
from bokeh.layouts import gridplot, layout, row, column, Spacer
import bokeh.palettes
import seaborn as sns
import matplotlib as mpl
from cycler import cycler
import holoviews as hv
from holoviews.operation.timeseries import rolling
from holoviews import dim
from bokeh.layouts import row
from base64 import b64encode 
from bokeh.models.tickers import FixedTicker
import plotly
import plotly.graph_objs as go
from plotly.offline import init_notebook_mode
import iqplot
init_notebook_mode()


bokeh.io.output_notebook(INLINE)
hv.extension("bokeh")

# %% [code]
import selenium
from selenium.webdriver.firefox.options import Options
from selenium.webdriver.firefox.service import Service
from selenium.webdriver.firefox.webdriver import WebDriver as Firefox

# service = Service()
service = Service(executable_path="C:\\Users\\manis\\anaconda3\\envs\\txtl_tradeoff_env\\txtl_tradeoff_env\\Scripts\\geckodriver.exe")

options = Options()
options.add_argument("--headless")
options.set_preference('layout.css.devPixelsPerPx', f'{1}')

webdriver = Firefox(service=service, options=options)

# %% [code]
%load_ext watermark
%watermark -v -p numpy,bokeh,jupyterlab

# %% [markdown]
### Set plot theme to standardize plot style

# %% [code]
theme = Theme(json={'attrs': {
    
# apply defaults to Figure properties
'figure': {
    'toolbar_location': None,
    'outline_line_color': None,
    'min_border_right': 10,
    'height':1000,
    'width':1200,
},    
    
# apply defaults to Grid properties
'Grid': {
    'grid_line_color': None,
},
    
# apply defaults to Title properties
'Title': {
    'text_font_size': '30pt',
    'align': 'center'
},
    
# # apply defaults to Plot properties
# 'Plot': {
#     'renderers': {'add_layout': {'legend[0]':'right'}},
# },
    
# apply defaults to Axis properties
'Axis': {
    'major_label_text_font_size': '50pt',
    'axis_label_text_font_size': '55pt',
    'axis_label_text_font_style': 'normal',
    'axis_label_standoff':30
},

# apply defaults to Legend properties
'Legend': {
    'background_fill_alpha': 0.8,
    'location': 'top_right',
    "label_text_font_size": '15pt',
    "click_policy": 'hide',
    "title_text_font_style": 'normal',
    "title_text_font_size": '18pt'
}}})

bokeh.io.curdoc().theme = theme
hv.renderer('bokeh').theme = theme

# %% [code]
# initialize labels as figures
p_label_a = bokeh.plotting.figure(width=100, height=110, toolbar_location=None, tools="")
p_label_b = bokeh.plotting.figure(width=100, height=110, toolbar_location=None, tools="")
p_label_c = bokeh.plotting.figure(width=100, height=110, toolbar_location=None, tools="")
p_label_d = bokeh.plotting.figure(width=100, height=110, toolbar_location=None, tools="")
p_label_e = bokeh.plotting.figure(width=100, height=110, toolbar_location=None, tools="")
p_label_f = bokeh.plotting.figure(width=100, height=110, toolbar_location=None, tools="")


# create labels for the figures
label_a = Label(x=0, y=5, x_units='screen', y_units='screen',
              text_font_size = '100px',
              text_font_style = 'bold',
              text_color = 'black',
              text='A', 
              background_fill_color='white', background_fill_alpha=1.0)
label_b = Label(x=0, y=5, x_units='screen', y_units='screen',
              text_font_size = '100px',
              text_font_style = 'bold',
              text_color = 'black',
              text='B', 
              background_fill_color='white', background_fill_alpha=1.0)
label_c = Label(x=0, y=5, x_units='screen', y_units='screen',
              text_font_size = '100px',
              text_font_style = 'bold',
              text_color = 'black',
              text='C', 
              background_fill_color='white', background_fill_alpha=1.0)
label_d = Label(x=0, y=5, x_units='screen', y_units='screen',
              text_font_size = '100px',
              text_font_style = 'bold',
              text_color = 'black',
              text='D', 
              background_fill_color='white', background_fill_alpha=1.0)
label_e = Label(x=0, y=5, x_units='screen', y_units='screen',
              text_font_size = '100px',
              text_font_style = 'bold',
              text_color = 'black',
              text='E', 
              background_fill_color='white', background_fill_alpha=1.0)
label_f = Label(x=0, y=5, x_units='screen', y_units='screen',
              text_font_size = '100px',
              text_font_style = 'bold',
              text_color = 'black',
              text='F', 
              background_fill_color='white', background_fill_alpha=1.0)

# add labels
p_label_a.add_layout(label_a)
p_label_b.add_layout(label_b)
p_label_c.add_layout(label_c)
p_label_d.add_layout(label_d)
p_label_e.add_layout(label_e)
p_label_f.add_layout(label_f)

# %% [markdown]
### Define functions for computing bootstrap Spearman's $\rho$

# %% [code]
def bootstrap_pls(X, Y, n_components, n_bootstrap):

    # create PLS object
    plsca = PLSCanonical(n_components=1)

    # store spearman rho values somewhere
    spearman_rho_bootstrap = np.zeros((n_bootstrap, X.shape[1]))

    # iterate through number of bootstraps
    for i in range(n_bootstrap):

        # randomly shuffle data and partition in half
        X_resampled, Y_resampled = resample(X, Y, random_state=i)
        n = len(X)
        X_train = X_resampled[:3 * n // 4]
        Y_train = Y_resampled[:3 * n // 4]
        X_test = X_resampled[n // 4 :]
        Y_test = Y_resampled[n // 4 :]

        # train data on half the data set, use model to fit the other half
        plsca.fit(X_train, Y_train)
        X_train_r, Y_train_r = plsca.transform(X_train, Y_train)
        X_test_r, Y_test_r = plsca.transform(X_test, Y_test)

        # define re-dimensionalized X and reduced-dimension Y
        x = X_test_r[:,0]
        y = Y_test_r[:,0]

        # compute spearman rho
        res = sp.stats.spearmanr(x, y)
        spearman_rho = res[0]
        spearman_rho_bootstrap[i] = spearman_rho
        
    return spearman_rho_bootstrap


def spearman_rho_bootstrap_values(X, Y):

    # Perform bootstrap PLS regression and calculate p-values
    n_components = 1
    n_bootstrap = 5000
    spearman_rho_bootstrap = bootstrap_pls(X, Y, n_components, n_bootstrap)
    
    # Calculate the mean and standard deviation of the bootstrap coefficients
    rho_mean = np.mean(spearman_rho_bootstrap, axis=0)
    rho_std = np.std(spearman_rho_bootstrap, axis=0)
    
    # Print the results
    print("Bootstrapped Spearman's rho Mean:", rho_mean[0])
    print("Bootstrapped Spearman's rho Std Dev:", rho_std[0])

# %% [markdown]
### Load data

# %% [code]
master_df = pd.read_csv('../Tidy Data/tidy_data_calibrated_full.csv')
master_df.head()

# %% [code]
master_df.tail()

# %% [markdown]
# 

# %% [markdown]
# Figures: 0, 1, 2, S2, 3, 7, S7, S8, S11, S12, S13

# %% [code]
# get relevant mgaptamer background subtraction data
# ar_zj lysate
arzj_bkgd_samples = master_df.loc[(master_df['DNA (nM)']==0) &
                      (master_df['DNA/mRNA batch'] < 5) &
                      (master_df['Extract']=='AR_ZJ') & 
                      (master_df['Fuel (mM)']==0) & 
                      (master_df['Mg (mM)']==0) & 
                      (master_df['Plasmid construct']=='pOR1OR2-MGapt-deGFP') & 
                      (master_df['mRNA (uM)'] == 0) & 
                      (master_df['Channel'] == 'MGaptamer')
                    ]
x = arzj_bkgd_samples['Time (hr)']
y = arzj_bkgd_samples['Measurement']
fn = np.polyfit(x, y, 5)
arzj_mgapt_fn = np.poly1d(fn)

# daj_mk_my lysate
dajmkmy_bkgd_samples = master_df.loc[(master_df['DNA (nM)']==0) &
                      (master_df['DNA/mRNA batch'] < 5) &
                      (master_df['Extract']=='DAJ_MK_MY') & 
                      (master_df['Fuel (mM)']==0) & 
                      (master_df['Mg (mM)']==0) & 
                      (master_df['Plasmid construct']=='pOR1OR2-MGapt-deGFP') & 
                      (master_df['mRNA (uM)'] == 0) & 
                      (master_df['Channel'] == 'MGaptamer')
                    ]
x = dajmkmy_bkgd_samples['Time (hr)']
y = dajmkmy_bkgd_samples['Measurement']
fn = np.polyfit(x, y, 5)
dajmkmy_mgapt_fn = np.poly1d(fn)
        
# create dictionary to store functions
mgaptamer_bkgd_subt_dict = {'AR_ZJ': arzj_mgapt_fn,
                            'DAJ_MK_MY': dajmkmy_mgapt_fn,
                            # '1 to 1 AR_ZJ + DAJ_MK_MY': 0.5*arzj_fn + 0.5*dajmkmy_fn,
                            # '1 to 2 AR_ZJ + DAJ_MK_MY': (1/3)*arzj_fn + (2/3)*dajmkmy_fn,
                            # '2 to 1 AR_ZJ + DAJ_MK_MY': (2/3)*arzj_fn + (1/3)*dajmkmy_fn
                           }

# %% [code]
# get relevant degfp background subtraction data
# ar_zj lysate
arzj_bkgd_samples = master_df.loc[(master_df['DNA (nM)']==0) &
                      (master_df['DNA/mRNA batch'] < 5) &
                      (master_df['Extract']=='AR_ZJ') & 
                      (master_df['Fuel (mM)']==0) & 
                      (master_df['Mg (mM)']==0) & 
                      (master_df['Plasmid construct']=='pOR1OR2-MGapt-deGFP') & 
                      (master_df['mRNA (uM)'] == 0) & 
                      (master_df['Channel'] == 'deGFP')
                    ]
x = arzj_bkgd_samples['Time (hr)']
y = arzj_bkgd_samples['Measurement']
fn = np.polyfit(x, y, 5)
arzj_degfp_fn = np.poly1d(fn)

# daj_mk_my lysate
dajmkmy_bkgd_samples = master_df.loc[(master_df['DNA (nM)']==0) &
                      (master_df['DNA/mRNA batch'] < 5) &
                      (master_df['Extract']=='DAJ_MK_MY') & 
                      (master_df['Fuel (mM)']==0) & 
                      (master_df['Mg (mM)']==0) & 
                      (master_df['Plasmid construct']=='pOR1OR2-MGapt-deGFP') & 
                      (master_df['mRNA (uM)'] == 0) & 
                      (master_df['Channel'] == 'deGFP')
                    ]
x = dajmkmy_bkgd_samples['Time (hr)']
y = dajmkmy_bkgd_samples['Measurement']
fn = np.polyfit(x, y, 5)
dajmkmy_degfp_fn = np.poly1d(fn)
        
# create dictionary to store functions
degfp_bkgd_subt_dict = {'AR_ZJ': arzj_degfp_fn,
                            'DAJ_MK_MY': dajmkmy_degfp_fn,
                            # '1 to 1 AR_ZJ + DAJ_MK_MY': 0.5*arzj_fn + 0.5*dajmkmy_fn,
                            # '1 to 2 AR_ZJ + DAJ_MK_MY': (1/3)*arzj_fn + (2/3)*dajmkmy_fn,
                            # '2 to 1 AR_ZJ + DAJ_MK_MY': (2/3)*arzj_fn + (1/3)*dajmkmy_fn
                           }

# %% [code]
# get dataframe that contains relevant data
temp_df = master_df.drop(index=master_df[((master_df['Fuel'] !='Succinate') & (master_df['Fuel'] !='Pyruvate')  & (master_df['Fuel'] !='Maltose')  & (master_df['Fuel'] !='3PGA') & (master_df['Fuel'] !='NTPs') & (master_df['Fuel'] !='No fuel')) |
                                                 ((master_df['Extract'] !='AR_ZJ') & (master_df['Extract'] !='DAJ_MK_MY')) |
                                                 (master_df['DNA (nM)'] != 5) |
                                                 (master_df['Plasmid construct'] != 'pOR1OR2-MGapt-deGFP') | 
                                                 (master_df['mRNA (uM)'] != 0) | 
                                                 (master_df['DNA/mRNA batch'] >= 5)
                                                ].index).groupby(['Extract','Fuel', 'DNA (nM)', 'Channel', 'NTPs (mM)', 'Added GTP (mM)', 'Added ATP (mM)', 'Replicate'])


# create list to store dataframes
integrated_mgapt_dfs = []
max_degfp_dfs = []

# iterate through groups
for name, group in temp_df:
                
    # get relevant information
    extract = name[0]
    fuel = name[1]
    dna_conc = name[2]
    fluor_mol = name[3]
    
    # background subtraction
    if fluor_mol == 'MGaptamer':
        bkgd_sub_fn = mgaptamer_bkgd_subt_dict[list(group['Extract'])[0]]
    elif fluor_mol == 'deGFP':
        bkgd_sub_fn = degfp_bkgd_subt_dict[list(group['Extract'])[0]]
    group['Measurement'] = group['Measurement'] - bkgd_sub_fn(group['Time (hr)'])    
    
    # drop unrelated columns, create new df
    # group = group.drop(columns=['Well'])
    # cols = list(group.columns)
    # cols.remove('Replicate')
    # cols.remove('Measurement')
    # new_group = group.groupby(cols).apply(lambda group2: np.mean(group2['Measurement'])).reset_index().ffill()#fillna(method="ffill")
    # new_group = pd.DataFrame(new_group).rename({0:'Measurement'}, axis=1)
    new_group = group.copy()

    if fuel=='None':
        print(new_group.head())
        
    # perform an integration
    if fluor_mol == 'MGaptamer':
        cols2 = list(new_group.columns)
        cols2.remove('Measurement')
        cols2.remove('Time (hr)')
        cols2.remove('Time (sec)')
        # new_group = new_group.groupby(cols2).apply(lambda group2: np.trapz(y=group2['Measurement'])).reset_index().ffill()#fillna(method="ffill")
        timepoints = sorted(list(set(new_group['Time (hr)'])))
        dx = timepoints[2] - timepoints[1]
        new_group = new_group.groupby(cols2).apply(lambda group2: np.trapz(y=group2['Measurement'], dx=dx)).reset_index().ffill()
        new_group = pd.DataFrame(new_group).rename({0:'Measurement'}, axis=1)
    elif fluor_mol == 'deGFP':
        new_group = new_group.drop(columns=['Time (sec)'])
        cols = list(new_group.columns)
        cols.remove('Measurement')
        cols.remove('Time (hr)')
        new_group = new_group.groupby(cols).apply(lambda group2: np.max(group2['Measurement'])).reset_index().ffill()#fillna(method="ffill")
        new_group = pd.DataFrame(new_group).rename({0:'Measurement'}, axis=1)
        
    # axis labels
    new_group['Fuel (mM)'] = new_group['Fuel (mM)'].astype('str')
    new_group['Mg (mM)'] = new_group['Mg (mM)'].astype('str')
    
    # drop all but relevant columns
    irrelevant_cols = list(new_group.columns)
    for i in ['Extract', 'DNA (nM)', 'Fuel (mM)', 'Mg (mM)', 'Fuel', 'NTPs (mM)', 'Added ATP (mM)', 'Added GTP (mM)', 'Replicate','Measurement']:
        irrelevant_cols.remove(i)
    new_group = new_group.drop(columns=irrelevant_cols)
    
    # rename measurement column, add relevant data to relevant list of dataframes
    if fluor_mol == 'MGaptamer':
        new_group = pd.DataFrame(new_group).rename({'Measurement':'Integrated MGaptamer (μM-hr)'}, axis=1)
        integrated_mgapt_dfs.append(new_group)
    elif fluor_mol == 'deGFP':
        new_group = pd.DataFrame(new_group).rename({'Measurement':'Maximum deGFP (μM)'}, axis=1)
        max_degfp_dfs.append(new_group)

# %% [code]
# combine list of dataframes into single dataframe for each channel
combined_integrated_mgapt_df = pd.concat(integrated_mgapt_dfs)
combined_max_degfp_df = pd.concat(max_degfp_dfs)

# combine all data into one dataframe via merge
max_degfp_integrated_mgapt_df = combined_integrated_mgapt_df.merge(combined_max_degfp_df, on=['Fuel', 'DNA (nM)', 'Mg (mM)', 'Fuel (mM)', 'Extract', 'Added ATP (mM)', 'Added GTP (mM)', 'NTPs (mM)', 'Replicate'])
max_degfp_integrated_mgapt_df.head()

# %% [code]
# plotting parameters
xlim = (0, 3.5)
ylim = (-1, 32)

# %% [markdown]
# Figure 0

# %% [code]
# create holoviews scatterplot
hv_fig = hv.Points(data=max_degfp_integrated_mgapt_df.loc[((max_degfp_integrated_mgapt_df['Fuel']!='NTPs') & (max_degfp_integrated_mgapt_df['Fuel']!='No fuel'))],
                    kdims=[('Integrated MGaptamer (μM-hr)', 'Int. MGapt'), ('Maximum deGFP (μM)', 'Max. deGFP')],
                    vdims=['Fuel', 'DNA (nM)', 'Mg (mM)', 'Fuel (mM)', 'Extract']
).opts(
    frame_height=1000,
    frame_width=1000,
    show_grid=False,
    color='#444444',
    size=20,
    xlim=(0.1, 3.5),
    ylim=(-1.2, 30)
    # xformatter='%.0f',
)

# render as bokeh figure, format
hv_fig = hv.render(hv_fig)
hv_fig.toolbar_location = None
hv_fig.xaxis.axis_label_text_font_size= '115pt'
hv_fig.yaxis.axis_label_text_font_size= '115pt'

# remove lines and numbers
hv_fig.xaxis.major_tick_line_color = None  # turn off x-axis major ticks
hv_fig.xaxis.minor_tick_line_color = None  # turn off x-axis minor ticks
hv_fig.yaxis.major_tick_line_color = None  # turn off y-axis major ticks
hv_fig.yaxis.minor_tick_line_color = None  # turn off y-axis minor ticks
hv_fig.xaxis.major_label_text_font_size = '0pt'  # preferred method for removing tick labels
hv_fig.yaxis.major_label_text_font_size = '0pt'  # preferred method for removing tick labels

# adjust line width
hv_fig.xaxis.axis_line_width = 5
hv_fig.yaxis.axis_line_width = 5

# save plot
file_save = '..\\Figures\\Fig0_b.png'
bokeh.io.reset_output()
bokeh.io.curdoc().theme = theme
bokeh.io.export_png(hv_fig, filename = file_save)

# %% [code]
def fig0():
    
    # Read in PNG images
    png_paths = ["..\\Figures\\Fig0_a.png", 
                   "..\\Figures\\Fig0_b.png"]
    
    # Create Divs for each subfigure
    divs = []
    for i, path in enumerate(png_paths):
        with open(path, "rb") as f:
            img_data = f.read()
        img_src = f"data:image/png;base64,{b64encode(img_data).decode()}"
        div_content = f'<img src="{img_src}">' # style="max-width:100%; max-height:100%">'
        div = Div(text=div_content)
        divs.append(div)
    
    # create panel of images
    panel = row([divs[0], row(Spacer(width=10)), column([Spacer(height=600), divs[1]])])
    
    # save plot
    file_save = '..\\Figures\\Fig0.png'
    bokeh.io.reset_output()
    bokeh.io.curdoc().theme = theme
    bokeh.io.export_png(panel, filename = file_save)

# %% [code]
fig0()

# %% [markdown]
# 

# %% [markdown]
# Figure 1

# %% [markdown]
### Figure 1c

# %% [code]
# create holoviews plot
color_legend_title = 'Fuel'
plotting_df = max_degfp_integrated_mgapt_df.copy()
plotting_df = plotting_df.loc[plotting_df['Fuel']!='NTPs']


hv_fig = hv.Points(data=plotting_df,
                    kdims=[('Integrated MGaptamer (μM-hr)', 'Integrated MG aptamer (μM-hr)'), 'Maximum deGFP (μM)'],
                    vdims=['Fuel', 'DNA (nM)', 'Mg (mM)', 'Fuel (mM)', 'Extract'],
                   
).opts(
    frame_height=1000,
    frame_width=1200,
    show_grid=False,
    cmap=['#009E73','#56B4E9','#000000','#D55E00','#F0E442',], # list(bokeh.palettes.Viridis[5]),
    color=color_legend_title,
    size=20,
    xlim=xlim,
    ylim=ylim,
    # tools=['hover'],
    # xformatter='%.0f',
    legend_opts={'title': 'Fuel',
                 'label_text_font_size':'45pt',
                 'title_text_font_size':'50pt',
                 'title_text_font_style': 'normal',
                 'glyph_width':50,
                 'glyph_height':50
                }
)

# render as bokeh figure, format
hv_fig = hv.render(hv_fig)

# save plot
file_save = '..\\Figures\\Fig1_c.png'
bokeh.io.reset_output()
bokeh.io.curdoc().theme = theme
bokeh.io.export_png(hv_fig, filename = file_save, webdriver=webdriver)

# %% [markdown]
### Fig 1d

# %% [code]
# create holoviews heatmap
color_legend_title = 'Fuel (mM)'
plotting_df = max_degfp_integrated_mgapt_df.copy()
plotting_df[color_legend_title] = pd.Categorical(plotting_df[color_legend_title],categories=['0', '5', '10', '15', '20', '30', '45'])
plotting_df = plotting_df.loc[plotting_df['Fuel']!='NTPs']
plotting_df = plotting_df.sort_values(color_legend_title)

hv_fig = hv.Points(data=plotting_df,
                    kdims=[('Integrated MGaptamer (μM-hr)', 'Integrated MG aptamer (μM-hr)'), 'Maximum deGFP (μM)'],
                    vdims=['Fuel', 'DNA (nM)', 'Mg (mM)', 'Fuel (mM)', 'Extract'],
                   
).opts(
    frame_height=1000,
    frame_width=1200,
    show_grid=False,
    cmap=list(bokeh.palettes.Viridis[7]),
    color=color_legend_title,
    size=20,
    # tools=['hover'],
    xlim=xlim,
    ylim=ylim,
    # xformatter='%.0f',
    legend_opts={'title': 'Fuel (mM)',
                 'label_text_font_size':'45pt',
                 'title_text_font_size':'50pt',
                 'title_text_font_style': 'normal',
                 'glyph_width':50,
                 'glyph_height':50
                }
)

# render as bokeh figure, format
hv_fig = hv.render(hv_fig)
hv_fig.legend.spacing = 25

# save plot
file_save = '..\\Figures\\Fig1_d.png'
bokeh.io.reset_output()
bokeh.io.curdoc().theme = theme
bokeh.io.export_png(hv_fig, filename = file_save)

# %% [code]
# compute statistics to show strength of relation
X = np.array(plotting_df['Fuel (mM)'].astype(int)).reshape(-1, 1)
Y = np.array([plotting_df['Integrated MGaptamer (μM-hr)'], plotting_df['Maximum deGFP (μM)']]).T
spearman_rho_bootstrap_values(X, Y)

# %% [markdown]
### Fig 1

# %% [code]
def fig1():
    
    # Read in PNG images
    png_paths = ["..\\Figures\\Fig1_a.png", 
                   "..\\Figures\\Fig1_b.png", 
                   "..\\Figures\\Fig1_c.png",
                 "..\\Figures\\Fig1_d.png"
                ]
    
    # Create Divs for each subfigure
    divs = []
    for i, path in enumerate(png_paths):
        with open(path, "rb") as f:
            img_data = f.read()
        img_src = f"data:image/png;base64,{b64encode(img_data).decode()}"
        if i == 0:
            div_content = f'<img src="{img_src}" margin:0; style="max-width:90%; max-height:90%">'
        else:
            div_content = f'<img src="{img_src}">' # style="max-width:100%; max-height:100%">'
        div = Div(text=div_content)
        divs.append(div)
    
    # create panel of images
    panel = column(row([p_label_a, column(Spacer(height=100),divs[0])]), 
                   Spacer(height=100), 
                   row([row([p_label_b, column(Spacer(height=100),divs[1])]), column([row([p_label_c, divs[2]]), 
                                                                                      Spacer(height=100), 
                                                                                      row([p_label_d, divs[3]]),
                                                                                      Spacer(height=50)
                                                                                     ])]))
    
    # save plot
    file_save = '..\\Figures\\Fig1.png'
    bokeh.io.reset_output()
    bokeh.io.curdoc().theme = theme
    bokeh.io.export_png(panel, filename = file_save)

# %% [code]
fig1()

# %% [markdown]
# Figure 2

# %% [markdown]
### Fig 2a

# %% [code]
# create holoviews heatmap
color_legend_title = 'Mg (mM)'
plotting_df = max_degfp_integrated_mgapt_df.copy()
plotting_df[color_legend_title] = pd.Categorical(plotting_df[color_legend_title],categories=['0', '2', '4', '6', '8', '10'])
plotting_df = plotting_df.loc[plotting_df['Fuel']!='NTPs']
plotting_df = plotting_df.sort_values(color_legend_title)

hv_fig = hv.Points(data=plotting_df,
                    kdims=[('Integrated MGaptamer (μM-hr)', 'Integrated MG aptamer (μM-hr)'), 'Maximum deGFP (μM)'],
                    vdims=['Fuel', 'DNA (nM)', 'Mg (mM)', 'Fuel (mM)', 'Extract'],
                   
).opts(
    frame_height=1000,
    frame_width=1200,
    show_grid=False,
    cmap=list(bokeh.palettes.Viridis[6]),
    # marker=dim('Extract').categorize({'DAJ_MK_MY': 'circle', "AR_ZJ": 'square'}),
    color=color_legend_title,
    size=20,
    xlim=xlim,
    ylim=ylim,
    # tools=['hover'],
    # xformatter='%.0f',
    legend_opts={'title': 'Added Mg²⁺ (mM)',
                 'label_text_font_size':'45pt',
                 'title_text_font_size':'50pt',
                 'title_text_font_style': 'normal',
                 'glyph_width':50,
                 'glyph_height':50
                }
)

# render as bokeh figure, format
hv_fig = hv.render(hv_fig)
hv_fig.legend.spacing = 25

# save plot
file_save = '..\\Figures\\Fig2_a.png'
bokeh.io.reset_output()
bokeh.io.curdoc().theme = theme
bokeh.io.export_png(hv_fig, filename = file_save, webdriver=webdriver)

# %% [code]
# compute statistics to show strength of relation
X = np.array(plotting_df['Mg (mM)'].astype(int)).reshape(-1, 1)
Y = np.array([plotting_df['Integrated MGaptamer (μM-hr)'], plotting_df['Maximum deGFP (μM)']]).T
spearman_rho_bootstrap_values(X, Y)

# %% [markdown]
### Fig 2b

# %% [code]
# create holoviews plot
color_legend_title = 'Extract'
plotting_df = max_degfp_integrated_mgapt_df.copy()
plotting_df = plotting_df.loc[plotting_df['Fuel']!='NTPs']
plotting_df = plotting_df.replace({'AR_ZJ': 'Batch 1', 'DAJ_MK_MY': 'Batch 2'})


hv_fig = hv.Points(data=plotting_df,
                    kdims=[('Integrated MGaptamer (μM-hr)', 'Integrated MG aptamer (μM-hr)'), 'Maximum deGFP (μM)'],
                    vdims=['Fuel', 'DNA (nM)', 'Mg (mM)', 'Fuel (mM)', 'Extract'],
                   
).opts(
    frame_height=1000,
    frame_width=1200,
    show_grid=False,
    cmap=[list(bokeh.palettes.Viridis[6])[0], list(bokeh.palettes.Viridis[6])[5]],
    color=color_legend_title,
    size=20,
    xlim=xlim,
    ylim=ylim,
    # tools=['hover'],
    # xformatter='%.0f',
    legend_opts={'title': 'Lysate batch',
                 'label_text_font_size':'45pt',
                 'title_text_font_size':'50pt',
                 'title_text_font_style': 'normal',
                 'glyph_width':50,
                 'glyph_height':50
                }
)

# render as bokeh figure, format
hv_fig = hv.render(hv_fig)
# bokeh.io.show(hv_fig)

# save plot
file_save = '..\\Figures\\Fig2_b.png'
bokeh.io.reset_output()
bokeh.io.curdoc().theme = theme
bokeh.io.export_png(hv_fig, filename = file_save, webdriver=webdriver)

# %% [code]
def fig2():
    
    # Read in PNG images
    png_paths = ["..\\Figures\\Fig2_a.png", 
                   "..\\Figures\\Fig2_b.png"]
    
    # Create Divs for each subfigure
    divs = []
    for i, path in enumerate(png_paths):
        with open(path, "rb") as f:
            img_data = f.read()
        img_src = f"data:image/png;base64,{b64encode(img_data).decode()}"
        div_content = f'<img src="{img_src}">' # style="max-width:100%; max-height:100%">'
        div = Div(text=div_content)
        divs.append(div)
    
    # create panel of images
    panel = row([row([p_label_a, divs[0]]), row(Spacer(width=100)), row([p_label_b, divs[1]])])
    
    # save plot
    file_save = '..\\Figures\\Fig2.png'
    bokeh.io.reset_output()
    bokeh.io.curdoc().theme = theme
    bokeh.io.export_png(panel, filename = file_save)

# %% [code]
fig2()

# %% [markdown]
# Fig S2

# %% [code]
# create holoviews heatmap
labels = ['a', 'b', 'c', 'd']

# iterate through fuels
for i, value in enumerate(['3PGA', 'Maltose', 'Pyruvate', 'Succinate']):

    # plot points colored by Mg
    color_legend_title = 'Mg (mM)'
    this_fuel = value
    plotting_df = max_degfp_integrated_mgapt_df.copy()
    plotting_df[color_legend_title] = pd.Categorical(plotting_df[color_legend_title],categories=['0', '2', '4', '6', '8', '10'])
    plotting_df = plotting_df.loc[plotting_df['Fuel'] == this_fuel]
    plotting_df = plotting_df.sort_values(['Extract', color_legend_title])
    
    hv_fig = hv.Points(data=plotting_df,
                        kdims=[('Integrated MGaptamer (μM-hr)', 'Integrated MG aptamer (μM-hr)'), 'Maximum deGFP (μM)'],
                        vdims=['Fuel', 'DNA (nM)', 'Mg (mM)', 'Fuel (mM)', 'Extract'],
                       
    ).opts(
        frame_height=1000,
        frame_width=1200,
        show_grid=False,
        cmap=list(bokeh.palettes.Viridis[6]),
        color=color_legend_title,
        marker=dim('Extract').categorize({'DAJ_MK_MY': 'plus', "AR_ZJ": 'circle'}),
        size=20,
        # xlim=(0.2, 3.3),
        # ylim=(-2, 30),
        xlim=xlim,
        ylim=ylim,
        # xformatter='%.0f',
        
        legend_opts={'title': 'Added Mg²⁺ (mM)',
                     'label_text_font_size':'45pt',
                     'title_text_font_size':'50pt',
                     'title_text_font_style': 'normal',
                     'glyph_width':50,
                     'glyph_height':50,
                    }
    )
    
    # render as bokeh figure, format
    hv_fig = hv.render(hv_fig)
    hv_fig.legend.spacing = 25
    # mytext = Label(x=0.1, y=-0.5, text=this_fuel, x_units='data', y_units='data', text_font_size='55pt', text_font_style='bold')
    mytext = Label(x=2.3, y=2.5, text=this_fuel, x_units='data', y_units='data', text_font_size='55pt', text_font_style='bold')

    
    hv_fig.add_layout(mytext)
    # bokeh.io.show(hv_fig)
    
    # save plot
    file_save = f'..\\Figures\\FigS2_{labels[i]}.png'
    bokeh.io.reset_output()
    bokeh.io.curdoc().theme = theme
    bokeh.io.export_png(hv_fig, filename = file_save)

    # compute statistics to show strength of relation
    print(this_fuel)
    X = np.array(plotting_df['Mg (mM)'].astype(int)).reshape(-1, 1)
    Y = np.array([plotting_df['Integrated MGaptamer (μM-hr)'], plotting_df['Maximum deGFP (μM)']]).T
    spearman_rho_bootstrap_values(X, Y)
    print('\n')

# %% [code]
def figS2():
    
    # Read in PNG images
    png_paths = ["..\\Figures\\FigS2_a.png", 
                "..\\Figures\\FigS2_b.png",
                "..\\Figures\\FigS2_c.png",
                "..\\Figures\\FigS2_d.png"]
    
    # Create Divs for each subfigure
    divs = []
    for i, path in enumerate(png_paths):
        with open(path, "rb") as f:
            img_data = f.read()
        img_src = f"data:image/png;base64,{b64encode(img_data).decode()}"
        div_content = f'<img src="{img_src}">' # style="max-width:100%; max-height:100%">'
        div = Div(text=div_content)
        divs.append(div)
    
    # create panel of images
    panel = column(row([row([p_label_a, divs[0]]), row(Spacer(width=100)), row([p_label_b, divs[1]])]),
                   Spacer(height=100),
                   row([row([p_label_c, divs[2]]), row(Spacer(width=100)), row([p_label_d, divs[3]])]),
                  )
    
    # save plot
    file_save = '..\\Figures\\FigS2.png'
    bokeh.io.reset_output()
    bokeh.io.curdoc().theme = theme
    bokeh.io.export_png(panel, filename = file_save)

# %% [code]
figS2()

# %% [markdown]
# Figure 3

# %% [code]
# create holoviews heatmap
labels = ['a', 'b', 'c', 'd']

# iterate through fuels
for i, value in enumerate(['3PGA', 'Maltose', 'Pyruvate', 'Succinate']):

    # plot points colored by Mg
    color_legend_title = 'Fuel (mM)'
    this_fuel = value
    plotting_df = max_degfp_integrated_mgapt_df.copy()
    plotting_df[color_legend_title] = pd.Categorical(plotting_df[color_legend_title],categories=['5', '10', '15', '20', '30', '45'])
    plotting_df = plotting_df.loc[plotting_df['Fuel'] == this_fuel]
    plotting_df = plotting_df.sort_values(['Extract', color_legend_title])
    
    hv_fig = hv.Points(data=plotting_df,
                        kdims=[('Integrated MGaptamer (μM-hr)', 'Integrated MG aptamer (μM-hr)'), 'Maximum deGFP (μM)'],
                        vdims=['Fuel', 'DNA (nM)', 'Mg (mM)', 'Fuel (mM)', 'Extract'],
                       
    ).opts(
        frame_height=1000,
        frame_width=1200,
        show_grid=False,
        cmap=list(bokeh.palettes.Viridis[6]),
        color=color_legend_title,
        size=20,
        xlim=xlim,
        ylim=ylim,
        # xformatter='%.0f',
        marker=dim('Extract').categorize({'DAJ_MK_MY': 'plus', "AR_ZJ": 'circle'}),
        legend_opts={'title': f'{value} (mM)',
                     'label_text_font_size':'45pt',
                     'title_text_font_size':'50pt',
                     'title_text_font_style': 'normal',
                     'glyph_width':50,
                     'glyph_height':50
                    }
    )
    
    # render as bokeh figure, format
    hv_fig = hv.render(hv_fig)
    hv_fig.legend.spacing = 25
    mytext = Label(x=0.3, y=-1.5, text=this_fuel, x_units='data', y_units='data', text_font_size='55pt', text_font_style='bold')
    mytext = Label(x=2.3, y=2.5, text=this_fuel, x_units='data', y_units='data', text_font_size='55pt', text_font_style='bold')
    # hv_fig.add_layout(mytext)
    # bokeh.io.show(hv_fig)
    
    # save plot
    file_save = f'..\\Figures\\Fig3_{labels[i]}.png'
    bokeh.io.reset_output()
    bokeh.io.curdoc().theme = theme
    bokeh.io.export_png(hv_fig, filename = file_save)

    # compute statistics to show strength of relation
    print(value)
    X = np.array(plotting_df['Fuel (mM)'].astype(int)).reshape(-1, 1)
    Y = np.array([plotting_df['Integrated MGaptamer (μM-hr)'], plotting_df['Maximum deGFP (μM)']]).T
    spearman_rho_bootstrap_values(X, Y)
    print('\n')

# %% [code]
def fig3():
    
    # Read in PNG images
    png_paths = ["..\\Figures\\Fig3_a.png", 
                "..\\Figures\\Fig3_b.png",
                "..\\Figures\\Fig3_c.png",
                "..\\Figures\\Fig3_d.png"]
    
    # Create Divs for each subfigure
    divs = []
    for i, path in enumerate(png_paths):
        with open(path, "rb") as f:
            img_data = f.read()
        img_src = f"data:image/png;base64,{b64encode(img_data).decode()}"
        div_content = f'<img src="{img_src}">' # style="max-width:100%; max-height:100%">'
        div = Div(text=div_content)
        divs.append(div)
    
    # create panel of images
    panel = column(row([row([p_label_a, divs[0]]), row(Spacer(width=100)), row([p_label_b, divs[1]])]),
                   Spacer(height=100),
                   row([row([p_label_c, divs[2]]), row(Spacer(width=100)), row([p_label_d, divs[3]])]),
                  )
    
    # save plot
    file_save = '..\\Figures\\Fig3.png'
    bokeh.io.reset_output()
    bokeh.io.curdoc().theme = theme
    bokeh.io.export_png(panel, filename = file_save)

# %% [code]
fig3()

# %% [markdown]
# Figure 7

# %% [code]
# pick correct data
plotting_df = max_degfp_integrated_mgapt_df.copy()
plotting_df = plotting_df.drop(index=plotting_df[(plotting_df['Fuel'] =='No fuel')].index)

fuel_type = []
for i in plotting_df['Fuel']:
    if i=='NTPs':
        fuel_type.append('NTPs')
    else:
        fuel_type.append('Central carbon')
plotting_df['Fuel type'] = fuel_type
explicit_mapping = {'NTPs': 'red', 'Central carbon': '#444444'}

# create plot
hv_fig = hv.Points(data=plotting_df,
                    kdims=[('Integrated MGaptamer (μM-hr)', 'Integrated MG aptamer \n (μM-hr, log scale)'), 'Maximum deGFP (μM)'],
                    vdims=['Fuel', 'DNA (nM)', 'Mg (mM)', 'Fuel (mM)', 'Extract', 'Fuel type'],
                   
).opts(
    frame_height=1000,
    frame_width=1300,
    show_grid=False,
    cmap=explicit_mapping,
    color='Fuel type',
    size=23,
    # tools=['hover'],
    # xformatter='%.0f',
    ylim=ylim,
    logx=True,
    legend_opts={'title': 'Fuel',
                 'label_text_font_size':'55pt',
                 'title_text_font_size':'60pt',
                 'title_text_font_style': 'normal',
                 'glyph_width':50,
                 'glyph_height':50
                }
)

# render as bokeh figure, format
hv_fig = hv.render(hv_fig)
hv_fig.xaxis.axis_label_text_font_size = '65pt'
hv_fig.xaxis.major_label_text_font_size = '60pt'
hv_fig.yaxis.axis_label_text_font_size = '65pt'
hv_fig.yaxis.major_label_text_font_size = '60pt'
# hv_fig.axis[0].ticker = FixedTicker(ticks=[0, 5, 10, 25, 50, 100, 200])

# save plot
file_save = '..\\Figures\\Fig7_b.png'
bokeh.io.reset_output()
bokeh.io.curdoc().theme = theme
bokeh.io.export_png(hv_fig, filename = file_save, webdriver=webdriver)

# %% [code]
# traditional spearman's rho between deGFP and MG aptamer
for i in ['Central carbon', 'NTPs']:
    print(i)
    plotting_df_this_fuel = plotting_df.loc[plotting_df['Fuel type']==i]
    res = sp.stats.spearmanr(plotting_df_this_fuel['Integrated MGaptamer (μM-hr)'], plotting_df_this_fuel['Maximum deGFP (μM)'])
    print(f"Spearman's rho correlation: {res[0]}")
    print('\n')

# %% [code]
def fig7():
    
    # Read in PNG images
    png_paths = ["..\\Figures\\Fig7_a.png", 
                   "..\\Figures\\Fig7_b.png"]
    
    # Create Divs for each subfigure
    divs = []
    for i, path in enumerate(png_paths):
        with open(path, "rb") as f:
            img_data = f.read()
        img_src = f"data:image/png;base64,{b64encode(img_data).decode()}"
        div_content = f'<img src="{img_src}">' # style="max-width:100%; max-height:100%">'
        div = Div(text=div_content)
        divs.append(div)
    
    # create panel of images
    panel = row([row([p_label_a, divs[0]]), row(Spacer(width=100)), row([p_label_b, column([Spacer(height=400), divs[1]])])])
    
    # save plot
    file_save = '..\\Figures\\Fig7.png'
    bokeh.io.reset_output()
    bokeh.io.curdoc().theme = theme
    bokeh.io.export_png(panel, filename = file_save)

# %% [code]
fig7()

# %% [markdown]
# Figure S7

# %% [code]
# get ntp data
ntps_df = max_degfp_integrated_mgapt_df.copy()
ntps_df = ntps_df.drop(index=ntps_df[
                                     (ntps_df['Fuel'] !='NTPs') |
                                     (ntps_df['NTPs (mM)'] == 4.8) 
                                    ].index)

# add total atp, gtp columns
ntps_df['Total ATP (mM)'] = ntps_df['NTPs (mM)']*0.3125 + ntps_df[ 'Added ATP (mM)']
ntps_df['Total GTP (mM)'] = ntps_df['NTPs (mM)']*0.3125 + ntps_df[ 'Added GTP (mM)']

ntps_df.head()

# %% [code]
# split
ntps_df_mgapt = ntps_df.drop(columns=['Maximum deGFP (μM)'])
ntps_df_degfp = ntps_df.drop(columns=['Integrated MGaptamer (μM-hr)'])

# average over extracts for mg aptamer
cols = list(ntps_df_mgapt.columns)
cols.remove('Extract')
cols.remove('Replicate')
cols.remove('Integrated MGaptamer (μM-hr)')
ntps_df_mgapt = ntps_df_mgapt.groupby(cols).apply(lambda group: np.mean(group['Integrated MGaptamer (μM-hr)'])).reset_index().ffill()
ntps_df_mgapt = pd.DataFrame(ntps_df_mgapt).rename({0:'Integrated MGaptamer (μM-hr)'}, axis=1)

# average over extracts for degfp
cols = list(ntps_df_degfp.columns)
cols.remove('Extract')
cols.remove('Replicate')
cols.remove('Maximum deGFP (μM)')
ntps_df_degfp = ntps_df_degfp.groupby(cols).apply(lambda group: np.mean(group['Maximum deGFP (μM)'])).reset_index().ffill()
ntps_df_degfp = pd.DataFrame(ntps_df_degfp).rename({0:'Maximum deGFP (μM)'}, axis=1)

# combine
ntps_df = ntps_df_mgapt.merge(ntps_df_degfp, on=['Fuel', 'Mg (mM)', 'Fuel (mM)', 
                                                 'NTPs (mM)','Added ATP (mM)', 
                                                 'Added GTP (mM)', 'Total ATP (mM)', 'Total GTP (mM)']
                             )

# %% [code]
# create contour plot
x=sorted(set(ntps_df_degfp['Total GTP (mM)']))
y=sorted(set(ntps_df_degfp['Total ATP (mM)']))
z = []

# iterate through x,y values and create 2-D array for z
for this_x in x:
    temp_lst = []
    for this_y in y:
        try:
            val = float(ntps_df_degfp.loc[(ntps_df_degfp['Total GTP (mM)']==this_x) & (ntps_df_degfp['Total ATP (mM)']==this_y)]['Maximum deGFP (μM)'])
            temp_lst.append(val)
        except:
            temp_lst.append(None)
    z.append(temp_lst)
z = np.array(z)

# %% [code]
# create figure
fig = go.Figure(data =
    go.Contour(
        z=z.T,
        x=x, # horizontal axis
        y=y, # vertical axis
        connectgaps=True,
        colorbar=dict(
            # title='deGFP (μM)', # title here
            titleside='right',
            titlefont=dict(
                size=75,
                family='Arial, sans-serif'),
            tickfont_size=50,
            thickness=65,
            thicknessmode='pixels',
            len=0.95,
            ticks='',
        ),
        contours_coloring='heatmap',
        line_width=2,
        colorscale = 'Viridis'
    )
)

# update plot settings
fig.update_layout(
    width=1200,
    height=1000,
    title={
        'text':"\t Maximum deGFP (μM)",
        'yanchor': 'top',
        'automargin': True, 
        'x':0.5,
    },
    title_font_family="Arial, sans-serif",
    title_font_color="#444444",
    legend_title_font_color="#444444",
    font=dict(
        family="Arial, sans-serif",
        size=55,
        color="#444444"
    ),
    xaxis_title="GTP (mM)",
    yaxis_title="ATP (mM)",
    xaxis_title_standoff=35,
    yaxis_title_standoff=50,
    title_pad={'b':25, 't':25,}
)

# %% [code]
#Plot and save html
file_save = '..\\Figures\\FigS7_b.png'
fig.write_image(file_save)

# %% [code]
# create contour plot
x=sorted(set(ntps_df_mgapt['Total GTP (mM)']))
y=sorted(set(ntps_df_mgapt['Total ATP (mM)']))
z = []

# iterate through x,y values and create 2-D array for z
for this_x in x:
    temp_lst = []
    for this_y in y:
        try:
            val = float(ntps_df_mgapt.loc[(ntps_df_mgapt['Total GTP (mM)']==this_x) & (ntps_df_degfp['Total ATP (mM)']==this_y)]['Integrated MGaptamer (μM-hr)'])
            temp_lst.append(val)
        except:
            temp_lst.append(None)
    z.append(temp_lst)
z = np.array(z)

# create figure
fig = go.Figure(data =
    go.Contour(
        z=z.T,
        x=x, # horizontal axis
        y=y, # vertical axis
        connectgaps=True,
        colorbar=dict(
            titleside='right',
            titlefont=dict(
                size=75,
                family='Arial, sans-serif'),
            tickfont_size=50,
            thickness=65,
            thicknessmode='pixels',
            len=0.95,
            ticks='',
        ),
        contours_coloring='heatmap',
        line_width=2,
        colorscale = 'Viridis'
    )
)

# update plot settings
fig.update_layout(
    width=1150,
    height=1000,
    title={
        'text':"\t Integrated MG aptamer (μM-hr)",
        'yanchor': 'top',
        'automargin': True, 
        'x':0.5,
    },
    title_font_family="Arial, sans-serif",
    title_font_color="#444444",
    legend_title_font_color="#444444",
    font=dict(
        family="Arial, sans-serif",
        size=55,
        color="#444444"
    ),
    xaxis_title="GTP (mM)",
    yaxis_title="ATP (mM)",
    xaxis_title_standoff=35,
    yaxis_title_standoff=50,
    title_pad={'b':25, 't':25,}
)

#Plot and save html
file_save = '..\\Figures\\FigS7_a.png'
fig.write_image(file_save)

# %% [code]
def figS7():
    
    # Read in PNG images
    png_paths = ["..\\Figures\\FigS7_a.png", 
                   "..\\Figures\\FigS7_b.png"]
    
    # Create Divs for each subfigure
    divs = []
    for i, path in enumerate(png_paths):
        with open(path, "rb") as f:
            img_data = f.read()
        img_src = f"data:image/png;base64,{b64encode(img_data).decode()}"
        div_content = f'<img src="{img_src}">' # style="max-width:100%; max-height:100%">'
        div = Div(text=div_content)
        divs.append(div)
    
    # create panel of images
    panel = row([row([p_label_a, divs[0]]), row(Spacer(width=100)), row([p_label_b, divs[1]])])
    
    # save plot
    file_save = '..\\Figures\\FigS7.png'
    bokeh.io.reset_output()
    bokeh.io.curdoc().theme = theme
    bokeh.io.export_png(panel, filename = file_save)

# %% [code]
figS7()

# %% [markdown]
# 

# %% [markdown]
# Figure S8

# %% [code]
# define function to extract data
def get_timecourse_data(df):

    # create dataframes
    integrated_mgapt_dfs = []
    max_degfp_dfs = []

    # get dataframe that contains relevant data
    temp_df = df.groupby(['Channel', 'Replicate'])
    
    # iterate through groups
    for name, group in temp_df:
                    
        # get relevant information
        fluor_mol = name[0]
        replicate = name[1]
        
        # background subtraction
        if fluor_mol == 'MGaptamer':
            bkgd_sub_fn = mgaptamer_bkgd_subt_dict['DAJ_MK_MY']
        elif fluor_mol == 'deGFP':
            bkgd_sub_fn = degfp_bkgd_subt_dict['DAJ_MK_MY']
        group['Measurement'] = group['Measurement'] - bkgd_sub_fn(group['Time (hr)'])    
        new_group = group.copy()
            
        # axis labels
        new_group['Mg (mM)'] = new_group['Mg (mM)'].astype('str')
                
        # drop all but relevant columns
        irrelevant_cols = list(new_group.columns)
        for i in ['Channel', 'Time (hr)','Mg (mM)', 'NTPs (mM)', 'Added ATP (mM)', 'Added GTP (mM)', 'Replicate', 'Measurement']:
            irrelevant_cols.remove(i)
        new_group = new_group.drop(columns=irrelevant_cols)
        
        # rename measurement column, add relevant data to relevant list of dataframes
        if fluor_mol == 'MGaptamer':
            integrated_mgapt_dfs.append(new_group)
        elif fluor_mol == 'deGFP':
            max_degfp_dfs.append(new_group)

    # return lists of dataframes
    return pd.concat(integrated_mgapt_dfs), pd.concat(max_degfp_dfs)

# %% [code]
def sample_averaged_timecourse(group, changing_var):
    '''
    Takes in a dataframe, fixed variable, and changing variable. Returns a bokeh plot of time course data of deGFP dynamics with error band
    
    inputs
    ------
    df : Pandas dataframe
        Dataframe containing fluorescence time course data
    changing_var : str
        Name of changing variable to be used in legend
    
    outputs
    -------
    p : Bokeh.plotting.figure instance
        A plot of the time course data
    '''
    
    # define variables
    n_colors = 4

    # define fluorescence channel
    channel = list(group['Channel'])[0]

    # final channel name
    if channel == 'MGaptamer':
        final_channel_name = 'MG aptamer (μM)'
        ylim = (-0.01, np.max(group['Measurement'])*1.1)
    elif channel == 'deGFP':
        final_channel_name = 'deGFP (μM)'
        ylim = (-0.01, np.max(group['Measurement'])*1.1)

    # if ntps as changing variable, sort the dataframe by ntp value ascending
    if changing_var == 'NTPs (mM)':
        group[changing_var] = group[changing_var].astype('int')
        group = group.sort_values(changing_var)    
    
    # plot time rolling average of 3 replicates
    hv_fig1 = hv.Curve(
        data=group,
        kdims=[("Time (hr)", "Time (hrs)")],
        vdims=[('Measurement', final_channel_name), 'Replicate', changing_var],
    ).groupby([changing_var, 'Replicate']
    ).collapse('Replicate',np.mean
    ).opts(
        color=hv.Cycle(list(bokeh.palettes.Viridis[n_colors])),
        frame_height=1000,
        frame_width=1200,
        line_width=4,
        line_color=hv.Cycle(list(bokeh.palettes.Viridis[n_colors]))
    ).overlay(
    ).opts(
        # legend_position="right",
        # legend_offset=(10, 300),
        legend_opts={'click_policy':'hide',
                    },
    ) 
    hv_fig1 = rolling(hv_fig1, rolling_window=10)
    
    
    # plot all data also
    hv_fig2 = hv.Points(
        data=group,
        kdims=[("Time (hr)", "Time (hrs)"), ('Measurement', final_channel_name)],
        vdims=[changing_var],
    ).groupby([changing_var]
    ).opts(
        color=hv.Cycle(list(bokeh.palettes.Viridis[n_colors])),
        size=10,
        frame_height=1000,
        frame_width=1200,
        xlim=(-0.1, 18.1),
        ylim=ylim,
        line_color=hv.Cycle(list(bokeh.palettes.Viridis[n_colors])),
        alpha=1
    ).overlay(
    )
    
    # create bokeh figure from holoviews figure
    hv_fig = hv_fig1*hv_fig2
    p = hv.render(hv_fig2)

    # p.legend.location = 'top_right'
    p.legend.title = changing_var
    p.legend.label_text_font_size = '35pt'
    p.legend.title_text_font_size='40pt'
    p.legend.title_text_font_style='normal'
    p.legend.glyph_width=50
    p.legend.glyph_height=50

    return(p)

# %% [code]
# get dataframe that contains relevant data
temp_df = master_df.drop(index=master_df[((master_df['Fuel'] !='NTPs')) |
                                                 ((master_df['Extract'] !='DAJ_MK_MY')) |
                                                 (master_df['DNA (nM)'] != 5) |
                                                 # (master_df['Channel'] != 'MGaptamer') | 
                                                 (master_df['Plasmid construct'] != 'pOR1OR2-MGapt-deGFP') | 
                                                 (master_df['mRNA (uM)'] != 0) | 
                                                 (master_df['DNA/mRNA batch'] >= 5) 
                                                ].index)

# name-save suffix
subplot_suffix = ['a', 'b', 'c', 'd', 'e', 'f']


# list of dataframes
list_of_dfs = [temp_df.loc[(temp_df['NTPs (mM)']==0) & (temp_df['Added GTP (mM)']==15)], #  0mM NTPs, 15mM GTP --> delayed TX
               temp_df.loc[(temp_df['Added ATP (mM)']==5) & (temp_df['Added GTP (mM)']==5)], # 5mM ATP, 5mM GTP --> biphasic TX
               temp_df.loc[(temp_df['Added ATP (mM)']==5) & (temp_df['Added GTP (mM)']==10)], # 5mM ATP, 10mM GTP --> sustained TX
               temp_df.loc[(temp_df['NTPs (mM)']==0) & (temp_df['Added ATP (mM)']==5)], #  0mM NTPs, 5mM ATP --> TL curve changes
               temp_df.loc[(temp_df['NTPs (mM)']==5) & (temp_df['Added ATP (mM)']==5)], # 5mM NTPs, 5mM ATP --> very interesting because a late start gets higher TL
               temp_df.loc[(temp_df['Added ATP (mM)']==5) & (temp_df['Added GTP (mM)']==5)] # 5mM ATP, 5mM GTP --> delayed TL
]
changing_var = ['Added ATP (mM)',
                'NTPs (mM)',
                'NTPs (mM)',
                'Added GTP (mM)',
                'Added GTP (mM)',
                'NTPs (mM)'
]

# iterate through datarframes
for index, this_df in enumerate(list_of_dfs):

    # get tx, tl data
    integrated_mgapt_dfs, max_degfp_dfs = get_timecourse_data(this_df)

    # tx plots
    if index < 3:
        p = sample_averaged_timecourse(integrated_mgapt_dfs, changing_var[index])

    # tl plots
    else:
        p = sample_averaged_timecourse(max_degfp_dfs, changing_var[index])

    # save figure
    file_save = f'..\\Figures\\FigS8_{subplot_suffix[index]}.png'
    bokeh.io.reset_output()
    bokeh.io.curdoc().theme = theme
    bokeh.io.export_png(p, filename = file_save)

# %% [code]
def figS8():
    
    # Read in PNG images
    png_paths = ["..\\Figures\\FigS8_a.png", 
                "..\\Figures\\FigS8_b.png",
                "..\\Figures\\FigS8_c.png",
                "..\\Figures\\FigS8_d.png",
                 "..\\Figures\\FigS8_e.png",
                 "..\\Figures\\FigS8_f.png"
                ]
    
    # Create Divs for each subfigure
    divs = []
    for i, path in enumerate(png_paths):
        with open(path, "rb") as f:
            img_data = f.read()
        img_src = f"data:image/png;base64,{b64encode(img_data).decode()}"
        # if i > 2:
        #     div_content = f'<img src="{img_src}" margin:0; style="max-width:50%; max-height:50%">'
        # else:
        div_content = f'<img src="{img_src}">' # style="max-width:100%; max-height:100%">'
        div = Div(text=div_content)
        divs.append(div)

    panel = column(row([row([p_label_a, divs[0]]), row(Spacer(width=100)), row([p_label_b, divs[1]]), row(Spacer(width=100)), row([p_label_c, divs[2]])]),
                   Spacer(height=100),
                   row([row([p_label_d, divs[3]]), row(Spacer(width=100)), row([p_label_e, divs[4]]), row(Spacer(width=100)), row([p_label_f, divs[5]])])
                  )
    panel = column(row([row([p_label_a, divs[0]]), row(Spacer(width=100)), row([p_label_b, divs[1]])]), 
                   Spacer(height=100),
                   row([row([p_label_c, divs[2]]), row(Spacer(width=100)), row([p_label_d, divs[3]])]), 
                   Spacer(height=100),
                   row([row([p_label_e, divs[4]]), row(Spacer(width=100)), row([p_label_f, divs[5]])]), 
                   Spacer(height=100),
                  )
 
    # save plot
    file_save = '..\\Figures\\FigS8.png'
    bokeh.io.reset_output()
    bokeh.io.curdoc().theme = theme
    bokeh.io.export_png(panel, filename = file_save)

# %% [code]
figS8()

# %% [markdown]
# 

# %% [markdown]
# Figures S10, S11, S12, S13

# %% [code]
# get relevant mgaptamer background subtraction data
# ar_zj lysate
arzj_bkgd_samples = master_df.loc[(master_df['DNA (nM)']==0) &
                      (master_df['DNA/mRNA batch'] < 5) &
                      (master_df['Extract']=='AR_ZJ') & 
                      (master_df['Fuel (mM)']==0) & 
                      (master_df['Mg (mM)']==0) & 
                      (master_df['Plasmid construct']=='pOR1OR2-MGapt-deGFP') & 
                      (master_df['mRNA (uM)'] == 0) & 
                      (master_df['Channel'] == 'MGaptamer')
                    ]
x = arzj_bkgd_samples['Time (hr)']
y = arzj_bkgd_samples['Measurement']
fn = np.polyfit(x, y, 5)
arzj_mgapt_fn = np.poly1d(fn)

# daj_mk_my lysate
dajmkmy_bkgd_samples = master_df.loc[(master_df['DNA (nM)']==0) &
                      (master_df['DNA/mRNA batch'] < 5) &
                      (master_df['Extract']=='DAJ_MK_MY') & 
                      (master_df['Fuel (mM)']==0) & 
                      (master_df['Mg (mM)']==0) & 
                      (master_df['Plasmid construct']=='pOR1OR2-MGapt-deGFP') & 
                      (master_df['mRNA (uM)'] == 0) & 
                      (master_df['Channel'] == 'MGaptamer')
                    ]
x = dajmkmy_bkgd_samples['Time (hr)']
y = dajmkmy_bkgd_samples['Measurement']
fn = np.polyfit(x, y, 5)
dajmkmy_mgapt_fn = np.poly1d(fn)
        
# create dictionary to store functions
mgaptamer_bkgd_subt_dict = {'AR_ZJ': arzj_mgapt_fn,
                            'DAJ_MK_MY': dajmkmy_mgapt_fn,
                            # '1 to 1 AR_ZJ + DAJ_MK_MY': 0.5*arzj_fn + 0.5*dajmkmy_fn,
                            # '1 to 2 AR_ZJ + DAJ_MK_MY': (1/3)*arzj_fn + (2/3)*dajmkmy_fn,
                            # '2 to 1 AR_ZJ + DAJ_MK_MY': (2/3)*arzj_fn + (1/3)*dajmkmy_fn
                           }

# %% [code]
# get relevant degfp background subtraction data
# ar_zj lysate
arzj_bkgd_samples = master_df.loc[(master_df['DNA (nM)']==0) &
                      (master_df['DNA/mRNA batch'] < 5) &
                      (master_df['Extract']=='AR_ZJ') & 
                      (master_df['Fuel (mM)']==0) & 
                      (master_df['Mg (mM)']==0) & 
                      (master_df['Plasmid construct']=='pOR1OR2-MGapt-deGFP') & 
                      (master_df['mRNA (uM)'] == 0) & 
                      (master_df['Channel'] == 'deGFP')
                    ]
x = arzj_bkgd_samples['Time (hr)']
y = arzj_bkgd_samples['Measurement']
fn = np.polyfit(x, y, 5)
arzj_degfp_fn = np.poly1d(fn)

# daj_mk_my lysate
dajmkmy_bkgd_samples = master_df.loc[(master_df['DNA (nM)']==0) &
                      (master_df['DNA/mRNA batch'] < 5) &
                      (master_df['Extract']=='DAJ_MK_MY') & 
                      (master_df['Fuel (mM)']==0) & 
                      (master_df['Mg (mM)']==0) & 
                      (master_df['Plasmid construct']=='pOR1OR2-MGapt-deGFP') & 
                      (master_df['mRNA (uM)'] == 0) & 
                      (master_df['Channel'] == 'deGFP')
                    ]
x = dajmkmy_bkgd_samples['Time (hr)']
y = dajmkmy_bkgd_samples['Measurement']
fn = np.polyfit(x, y, 5)
dajmkmy_degfp_fn = np.poly1d(fn)
        
# create dictionary to store functions
degfp_bkgd_subt_dict = {'AR_ZJ': arzj_degfp_fn,
                            'DAJ_MK_MY': dajmkmy_degfp_fn,
                            # '1 to 1 AR_ZJ + DAJ_MK_MY': 0.5*arzj_fn + 0.5*dajmkmy_fn,
                            # '1 to 2 AR_ZJ + DAJ_MK_MY': (1/3)*arzj_fn + (2/3)*dajmkmy_fn,
                            # '2 to 1 AR_ZJ + DAJ_MK_MY': (2/3)*arzj_fn + (1/3)*dajmkmy_fn
                           }

# %% [code]
# plot heatmaps
def create_heatmaps(combined_integrated_mgapt_df, combined_max_degfp_df, fuel, dna_conc, extract):

    # MG APTAMER PLOT
    # axis labels
    new_group = combined_integrated_mgapt_df.copy()
    new_group['Fuel (mM)'] = new_group['Fuel (mM)'].astype('str')
    new_group['Mg (mM)'] = new_group['Mg (mM)'].astype('str')
    if dna_conc == '1':
        new_group['Fuel (mM)'] = pd.Categorical(new_group['Fuel (mM)'],categories=['5', '10', '15', '20', '30', '45'])
    else: 
        new_group['Fuel (mM)'] = pd.Categorical(new_group['Fuel (mM)'],categories=['0', '5', '10', '15', '20', '30', '45'])
    new_group['Mg (mM)'] = pd.Categorical(new_group['Mg (mM)'],categories=['0', '2', '4', '6', '8', '10'])
    new_group = new_group.sort_values(['Fuel (mM)', "Mg (mM)"])
    new_group = new_group.reset_index().drop(labels=['index'], axis=1)
    
    # create holoviews heatmap
    value_dimension = hv.Dimension('Integrated MGaptamer (μM-hr)', value_format=lambda x: '%.1f' % x)   
    # plot range max
    this_max = np.max(new_group['Integrated MGaptamer (μM-hr)'])
    hv_fig = hv.HeatMap(data=new_group,
                        kdims=[('Fuel (mM)', f'Added {fuel} (mM)'), ('Mg (mM)', 'Added Mg²⁺ (mM)') ],
                        vdims=hv.Dimension('Integrated MGaptamer (μM-hr)', range=(0, this_max))
    ).opts(
        frame_height=500,
        frame_width=500,
        colorbar=True,
        show_grid=False,
        cmap='Viridis',
        title = f'Integrated MGaptamer (μM-hr)',
        fontsize={'labels': 35, 'xticks': 30, 'yticks': 30, 'ticks':30}
        
    )
    hv_fig = hv_fig * hv.Labels(hv_fig, vdims=value_dimension).opts(text_color='white', text_font_size='20px',fontsize={'ticks':20})
    
    # render as bokeh figure, format
    hv_fig1 = hv.render(hv_fig)
    hv_fig1.toolbar_location = None
    hv_fig1.title.align = 'center'


    # DEGFP PLOT
    # define rounding
    value_dimension = hv.Dimension('Maximum deGFP (μM)', value_format=lambda x: '%.2f' % x)    
    
    # plot range max
    new_group = combined_max_degfp_df.copy()
    new_group['Fuel (mM)'] = pd.Categorical(new_group['Fuel (mM)'],categories=['0', '5', '10', '15', '20', '30', '45'])
    new_group['Mg (mM)'] = pd.Categorical(new_group['Mg (mM)'],categories=['0', '2', '4', '6', '8', '10'])
    new_group = new_group.sort_values(['Fuel (mM)', "Mg (mM)"])
    new_group = new_group.reset_index().drop(labels=['index'], axis=1)
    this_max = np.max(new_group['Maximum deGFP (μM)'])
    
    # create holoviews heatmap
    hv_fig = hv.HeatMap(data=new_group,
                        kdims=[('Fuel (mM)', f'Added {fuel} (mM)'), ('Mg (mM)', 'Added Mg²⁺ (mM)') ],
                        vdims=hv.Dimension('Maximum deGFP (μM)', range=(0, this_max))
    # ).aggregate(function=np.max
    ).opts(
        frame_height=500,
        frame_width=500,
        colorbar=True,
        show_grid=False,
        cmap='Viridis',
        title = f'Maximum deGFP (μM)',
        fontsize={'labels': 35, 'xticks': 30, 'yticks': 30, 'ticks':30}
    )
    hv_fig = hv_fig * hv.Labels(hv_fig, vdims=value_dimension).opts(text_color='white', text_font_size='20px',fontsize={'ticks':20})
    
    # render as bokeh figure, format
    hv_fig2 = hv.render(hv_fig)
    hv_fig2.toolbar_location = None
    hv_fig2.title.align = 'center'
    
    # add plot title
    if extract == 'AR_ZJ':
        title = Div(text = f'<p style="color: #444444; font-size: 75px; margin:0;"> Lysate 1, {dna_conc}nM DNA </p>', align='center')
    else:
        title = Div(text = f'<p style="color: #444444; font-size: 75px; margin:0;"> Lysate 2, {dna_conc}nM DNA </p>', align='center')
    p = column(title, hv_fig1, Spacer(height=50), hv_fig2)

    return(p)

# %% [code]
# function to extract data
def create_max_degfp_integrated_mgapt_df(master_df, fuel, extract, dna_conc):

    # get dataframe for particular fuel, extract, and dna conc
    # subset only relevant fuel
    subset_df1 = master_df.drop(index=master_df[(master_df['Fuel'] != 'No fuel') |
                                             (master_df['Extract'] !=extract) |
                                             (master_df['DNA (nM)'] != dna_conc) |
                                             (master_df['Plasmid construct'] != 'pOR1OR2-MGapt-deGFP') | 
                                             (master_df['mRNA (uM)'] != 0) | 
                                             (master_df['DNA/mRNA batch'] >= 5)
                                            ].index)
    subset_df2 = master_df.drop(index=master_df[(master_df['Fuel'] !=fuel) |
                                             (master_df['Extract'] !=extract) |
                                             (master_df['DNA (nM)'] != dna_conc) |
                                             (master_df['Plasmid construct'] != 'pOR1OR2-MGapt-deGFP') | 
                                             (master_df['mRNA (uM)'] != 0) | 
                                             (master_df['DNA/mRNA batch'] >= 5)
                                            ].index)
    combined_df = pd.concat([subset_df1, subset_df2], axis=0)
    combined_df = combined_df.replace('No fuel', fuel)
    temp_df = combined_df.groupby(['Extract','Fuel', 'DNA (nM)', 'Channel', 'NTPs (mM)', 'Added GTP (mM)', 'Added ATP (mM)'], observed=False)

    # create list to store dataframes
    integrated_mgapt_dfs = []
    max_degfp_dfs = []
    
    # iterate through groups
    for name, group in temp_df:
                    
        # get relevant information
        extract = name[0]
        fuel = name[1]
        dna_conc = name[2]
        fluor_mol = name[3]
        
        # background subtraction
        if fluor_mol == 'MGaptamer':
            bkgd_sub_fn = mgaptamer_bkgd_subt_dict[list(group['Extract'])[0]]
        elif fluor_mol == 'deGFP':
            bkgd_sub_fn = degfp_bkgd_subt_dict[list(group['Extract'])[0]]
        group['Measurement'] = group['Measurement'] - bkgd_sub_fn(group['Time (hr)'])    
        
        # drop unrelated columns, create new df
        group = group.drop(columns=['Well'])
        cols = list(group.columns)
        cols.remove('Replicate')
        cols.remove('Measurement')
        new_group = group.groupby(cols, observed=False).apply(lambda group2: np.mean(group2['Measurement'])).reset_index().ffill()#fillna(method="ffill")
        new_group = pd.DataFrame(new_group).rename({0:'Measurement'}, axis=1)
    
        if fuel=='None':
            print(new_group.head())
            
        # perform an integration
        if fluor_mol == 'MGaptamer':
            cols2 = list(new_group.columns)
            cols2.remove('Measurement')
            cols2.remove('Time (hr)')
            cols2.remove('Time (sec)')
            # new_group = new_group.groupby(cols2).apply(lambda group2: np.trapz(y=group2['Measurement'])).reset_index().ffill()#fillna(method="ffill")
            timepoints = sorted(list(set(new_group['Time (hr)'])))
            dx = timepoints[2] - timepoints[1]
            new_group = new_group.groupby(cols2, observed=False).apply(lambda group2: np.trapz(y=group2['Measurement'], dx=dx)).reset_index().ffill()
            new_group = pd.DataFrame(new_group).rename({0:'Measurement'}, axis=1)
        elif fluor_mol == 'deGFP':
            new_group = new_group.drop(columns=['Time (sec)'])
            cols = list(new_group.columns)
            cols.remove('Measurement')
            cols.remove('Time (hr)')
            new_group = new_group.groupby(cols, observed=False).apply(lambda group2: np.max(group2['Measurement'])).reset_index().ffill()#fillna(method="ffill")
            new_group = pd.DataFrame(new_group).rename({0:'Measurement'}, axis=1)
            
        # axis labels
        new_group['Fuel (mM)'] = new_group['Fuel (mM)'].astype('str')
        new_group['Mg (mM)'] = new_group['Mg (mM)'].astype('str')
        
        # drop all but relevant columns
        irrelevant_cols = list(new_group.columns)
        for i in ['Extract', 'DNA (nM)', 'Fuel (mM)', 'Mg (mM)', 'Fuel', 'NTPs (mM)', 'Added ATP (mM)', 'Added GTP (mM)', 'Measurement']:
            irrelevant_cols.remove(i)
        new_group = new_group.drop(columns=irrelevant_cols)
        
        # rename measurement column, add relevant data to relevant list of dataframes
        if fluor_mol == 'MGaptamer':
            new_group = pd.DataFrame(new_group).rename({'Measurement':'Integrated MGaptamer (μM-hr)'}, axis=1)
            integrated_mgapt_dfs.append(new_group)
        elif fluor_mol == 'deGFP':
            new_group = pd.DataFrame(new_group).rename({'Measurement':'Maximum deGFP (μM)'}, axis=1)
            max_degfp_dfs.append(new_group)

    # combine list of dataframes into single dataframe for each channel
    combined_integrated_mgapt_df = pd.concat(integrated_mgapt_dfs)
    combined_max_degfp_df = pd.concat(max_degfp_dfs)
    
    # combine all data into one dataframe via merge
    # max_degfp_integrated_mgapt_df = combined_integrated_mgapt_df.merge(combined_max_degfp_df, on=['Fuel', 'DNA (nM)', 'Mg (mM)', 'Fuel (mM)', 'Extract', 'Added ATP (mM)', 'Added GTP (mM)', 'NTPs (mM)'])
    max_degfp_integrated_mgapt_df = combined_integrated_mgapt_df.join(combined_max_degfp_df.set_index(['Fuel', 'DNA (nM)', 'Mg (mM)', 'Fuel (mM)', 'Extract', 'Added ATP (mM)', 'Added GTP (mM)', 'NTPs (mM)']), 
                                                                      on=['Fuel', 'DNA (nM)', 'Mg (mM)', 'Fuel (mM)', 'Extract', 'Added ATP (mM)', 'Added GTP (mM)', 'NTPs (mM)'])

    return(combined_integrated_mgapt_df, combined_max_degfp_df)

# %% [code]
# figure labels
fig_labels = ['10', '11', '12', '13']

# iterate through fuels
for index, fuel in enumerate(['3PGA', 'Maltose', 'Pyruvate', 'Succinate']):
    
    # iterate through extracts
    for extract in ['AR_ZJ', 'DAJ_MK_MY']:
        
        # iterate through dna concentrations
        if extract=='AR_ZJ':
            
            # iterate through two dna concs
            for dna_conc in [1, 5]:
                combined_integrated_mgapt_df, combined_max_degfp_df = create_max_degfp_integrated_mgapt_df(master_df, fuel, extract, dna_conc)
                # for plotting
                if dna_conc == 1:
                    subfig_label = 'a'
                else:
                    subfig_label = 'b'
                # give information to plotting function
                p = create_heatmaps(combined_integrated_mgapt_df, combined_max_degfp_df, fuel, dna_conc, extract)
                # save image
                file_save = f'..\\Figures\\FigS{fig_labels[index]}_{subfig_label}.png'
                bokeh.io.reset_output()
                bokeh.io.curdoc().theme = theme
                bokeh.io.export_png(p, filename = file_save)   
        else:
            dna_conc = 5
            combined_integrated_mgapt_df, combined_max_degfp_df = create_max_degfp_integrated_mgapt_df(master_df, fuel, extract, dna_conc)
            # give information to plotting function
            p = create_heatmaps(combined_integrated_mgapt_df, combined_max_degfp_df, fuel, dna_conc, extract)
            subfig_label = 'c'

            # save image
            file_save = f'..\\Figures\\FigS{fig_labels[index]}_{subfig_label}.png'
            bokeh.io.reset_output()
            bokeh.io.curdoc().theme = theme
            bokeh.io.export_png(p, filename = file_save)        

# %% [code]
def figS10_S11_S12_S13():

    # iterate through image numbers
    for img in ['10', '11', '12', '13']:
        
    
    
        # Read in PNG images
        png_paths = [f"..\\Figures\\FigS{img}_a.png", 
                    f"..\\Figures\\FigS{img}_b.png",
                    f"..\\Figures\\FigS{img}_c.png",
                    ]
        
        # Create Divs for each subfigure
        divs = []
        for i, path in enumerate(png_paths):
            with open(path, "rb") as f:
                img_data = f.read()
            img_src = f"data:image/png;base64,{b64encode(img_data).decode()}"
            # if i > 2:
            #     div_content = f'<img src="{img_src}" margin:0; style="max-width:50%; max-height:50%">'
            # else:
            div_content = f'<img src="{img_src}">' # style="max-width:100%; max-height:100%">'
            div = Div(text=div_content)
            divs.append(div)    
    
        panel = row([row([p_label_a, Spacer(width=50), divs[0]]), 
                            row(Spacer(width=100)), 
                            row([p_label_b, Spacer(width=50), divs[1]]), 
                            row(Spacer(width=100)), 
                            row([p_label_c, Spacer(width=50), divs[2]])])                   
    
        # save plot
        file_save = f'..\\Figures\\FigS{img}.png'
        bokeh.io.reset_output()
        bokeh.io.curdoc().theme = theme
        bokeh.io.export_png(panel, filename = file_save)

# %% [code]
figS10_S11_S12_S13()

# %% [markdown]
# 

# %% [markdown]
# Figure S9

# %% [code]
# define rolling window slope function
def compute_slope(window):
    # Prepare data for linear regression
    x = np.linspace(0,1,len(window)).reshape(-1, 1)
    y = window.values
    
    # Fit linear regression model
    model = LinearRegression()
    model.fit(x, y)
    
    # Return the slope
    return model.coef_[0]

# %% [code]
# get dataframe that contains relevant data
temp_df = master_df.drop(index=master_df[((master_df['Fuel'] !='Succinate') & (master_df['Fuel'] !='Pyruvate')  & (master_df['Fuel'] !='Maltose')  & (master_df['Fuel'] !='3PGA') & (master_df['Fuel'] !='NTPs') & (master_df['Fuel'] !='No fuel')) |
                                                 ((master_df['Extract'] !='AR_ZJ') & (master_df['Extract'] !='DAJ_MK_MY')) |
                                                 (master_df['DNA (nM)'] != 5) |
                                                 # (master_df['Channel'] != 'MGaptamer') | 
                                                 (master_df['Plasmid construct'] != 'pOR1OR2-MGapt-deGFP') | 
                                                 (master_df['mRNA (uM)'] != 0) | 
                                                 (master_df['DNA/mRNA batch'] >= 5)
                                                ].index).groupby(['Extract','Fuel', 'DNA (nM)', 'Channel', 'NTPs (mM)', 'Added GTP (mM)', 'Added ATP (mM)', 'Replicate'])


# create list to store dataframes
integrated_mgapt_dfs = []
max_degfp_dfs = []

# iterate through groups
for name, group in temp_df:
                
    # get relevant information
    extract = name[0]
    fuel = name[1]
    dna_conc = name[2]
    fluor_mol = name[3]
    
    # background subtraction
    if fluor_mol == 'MGaptamer':
        bkgd_sub_fn = mgaptamer_bkgd_subt_dict[list(group['Extract'])[0]]
    elif fluor_mol == 'deGFP':
        bkgd_sub_fn = degfp_bkgd_subt_dict[list(group['Extract'])[0]]
    group['Measurement'] = group['Measurement'] - bkgd_sub_fn(group['Time (hr)'])    
    new_group = group.copy()

    # pick window size so that it is roughly an hour
    if fuel=='NTPs':
        if extract=='AR_ZJ':
            window_size=19
        else:
            window_size=16
    else:
        window_size=21
        
    # perform an integration
    if fluor_mol == 'MGaptamer':
        cols2 = list(new_group.columns)
        cols2.remove('Measurement')
        cols2.remove('Time (hr)')
        cols2.remove('Time (sec)')
        # # new_group = new_group.groupby(cols2).apply(lambda group2: np.trapz(y=group2['Measurement'])).reset_index().ffill()#fillna(method="ffill")
        # timepoints = sorted(list(set(new_group['Time (hr)'])))
        # dx = timepoints[2] - timepoints[1]
        new_group = new_group.groupby(cols2).apply(lambda group2: group2['Measurement'].rolling(window=window_size).apply(compute_slope, raw=False).max()).reset_index().ffill()
        new_group = pd.DataFrame(new_group).rename({0:'Measurement'}, axis=1)
    elif fluor_mol == 'deGFP':
        new_group = new_group.drop(columns=['Time (sec)'])
        cols = list(new_group.columns)
        cols.remove('Measurement')
        cols.remove('Time (hr)')
        new_group = new_group.groupby(cols).apply(lambda group2: np.max(group2['Measurement'])).reset_index().ffill()#fillna(method="ffill")
        new_group = pd.DataFrame(new_group).rename({0:'Measurement'}, axis=1)
        
    # axis labels
    new_group['Fuel (mM)'] = new_group['Fuel (mM)'].astype('str')
    new_group['Mg (mM)'] = new_group['Mg (mM)'].astype('str')
    
    # drop all but relevant columns
    irrelevant_cols = list(new_group.columns)
    for i in ['Extract', 'DNA (nM)', 'Fuel (mM)', 'Mg (mM)', 'Fuel', 'NTPs (mM)', 'Added ATP (mM)', 'Added GTP (mM)', 'Replicate','Measurement']:
        irrelevant_cols.remove(i)
    new_group = new_group.drop(columns=irrelevant_cols)
    
    # rename measurement column, add relevant data to relevant list of dataframes
    if fluor_mol == 'MGaptamer':
        new_group = pd.DataFrame(new_group).rename({'Measurement':'Integrated MGaptamer (μM-hr)'}, axis=1)
        integrated_mgapt_dfs.append(new_group)
    elif fluor_mol == 'deGFP':
        new_group = pd.DataFrame(new_group).rename({'Measurement':'Maximum deGFP (μM)'}, axis=1)
        max_degfp_dfs.append(new_group)

# combine list of dataframes into single dataframe for each channel
combined_integrated_mgapt_df = pd.concat(integrated_mgapt_dfs)
combined_max_degfp_df = pd.concat(max_degfp_dfs)

# combine all data into one dataframe via merge
max_degfp_integrated_mgapt_df = combined_integrated_mgapt_df.merge(combined_max_degfp_df, on=['Fuel', 'DNA (nM)', 'Mg (mM)', 'Fuel (mM)', 'Extract', 'Added ATP (mM)', 'Added GTP (mM)', 'NTPs (mM)', 'Replicate'])
max_degfp_integrated_mgapt_df.head()

# %% [code]
# pick correct data
plotting_df = max_degfp_integrated_mgapt_df.copy()
plotting_df = plotting_df.drop(index=plotting_df[(plotting_df['Fuel'] =='No fuel')].index)

fuel_type = []
for i in plotting_df['Fuel']:
    if i=='NTPs':
        fuel_type.append('NTPs')
    else:
        fuel_type.append('Central carbon')
plotting_df['Fuel type'] = fuel_type
explicit_mapping = {'NTPs': 'red', 'Central carbon': '#444444'}

# create plot
hv_fig = hv.Points(data=plotting_df,
                    kdims=[('Integrated MGaptamer (μM-hr)', 'Maximum MG aptamer slope \n(μM/hr)'), 'Maximum deGFP (μM)'], 
                    vdims=['Fuel', 'DNA (nM)', 'Mg (mM)', 'Fuel (mM)', 'Extract', 'Fuel type'],
                   
).opts(
    frame_height=1000,
    frame_width=1300,
    show_grid=False,
    cmap=explicit_mapping,
    color='Fuel type',
    size=23,
    # tools=['hover'],
    # xformatter='%.0f',
    ylim=ylim,
    # logx=True,
    legend_opts={'title': 'Fuel',
                 'label_text_font_size':'55pt',
                 'title_text_font_size':'60pt',
                 'title_text_font_style': 'normal',
                 'glyph_width':50,
                 'glyph_height':50
                }
)

# render as bokeh figure, format
hv_fig = hv.render(hv_fig)
hv_fig.xaxis.axis_label_text_font_size = '65pt'
hv_fig.xaxis.major_label_text_font_size = '60pt'
hv_fig.yaxis.axis_label_text_font_size = '65pt'
hv_fig.yaxis.major_label_text_font_size = '60pt'
# hv_fig.axis[0].ticker = FixedTicker(ticks=[0, 5, 10, 25, 50, 100, 200])

# save plot
file_save = '..\\Figures\\FigS9.png'
bokeh.io.reset_output()
bokeh.io.curdoc().theme = theme
bokeh.io.export_png(hv_fig, filename = file_save, webdriver=webdriver)

# %% [code]
# traditional spearman's rho between deGFP and MG aptamer
for i in ['Central carbon', 'NTPs']:
    print(i)
    plotting_df_this_fuel = plotting_df.loc[plotting_df['Fuel type']==i]
    res = sp.stats.spearmanr(plotting_df_this_fuel['Integrated MGaptamer (μM-hr)'], plotting_df_this_fuel['Maximum deGFP (μM)'])
    print(f"Spearman's rho correlation: {res[0]}")
    print('\n')

# %% [markdown]
# 

# %% [markdown]
# Figure S3, 5, S6

# %% [code]
# get relevant mgaptamer background subtraction data
# daj_mk_my lysate
dajmkmy_bkgd_samples = master_df.loc[(master_df['DNA (nM)']==0) &
                      (master_df['DNA/mRNA batch'] == 6) &
                      (master_df['Extract']=='DAJ_MK_MY') & 
                      (master_df['Fuel (mM)']==0) & 
                      (master_df['Mg (mM)']==0) & 
                      (master_df['Plasmid construct']=='pOR1OR2-MGapt-deGFP') & 
                      (master_df['mRNA (uM)'] == 0) & 
                      (master_df['Channel'] == 'MGaptamer') &
                      (master_df['Tetratcycline (ug/mL)'] == 0)
                    ]
x = dajmkmy_bkgd_samples['Time (hr)']
y = dajmkmy_bkgd_samples['Measurement']
fn = np.polyfit(x, y, 5)
dajmkmy_mgapt_fn = np.poly1d(fn)
        
# create dictionary to store functions
mgaptamer_bkgd_subt_dict2 = {'DAJ_MK_MY': dajmkmy_mgapt_fn,
                            }

dajmkmy_bkgd_samples = master_df.loc[(master_df['DNA (nM)']==0) &
                      (master_df['DNA/mRNA batch'] == 6) &
                      (master_df['Extract']=='DAJ_MK_MY') & 
                      (master_df['Fuel (mM)']==0) & 
                      (master_df['Mg (mM)']==0) & 
                      (master_df['Plasmid construct']=='pOR1OR2-MGapt-deGFP') & 
                      (master_df['mRNA (uM)'] == 0) & 
                      (master_df['Channel'] == 'deGFP') &
                      (master_df['Tetratcycline (ug/mL)'] == 0)
                    ]
x = dajmkmy_bkgd_samples['Time (hr)']
y = dajmkmy_bkgd_samples['Measurement']
fn = np.polyfit(x, y, 5)
dajmkmy_degfp_fn = np.poly1d(fn)
        
# create dictionary to store functions
degfp_bkgd_subt_dict2 = {'DAJ_MK_MY': dajmkmy_degfp_fn,
                            }

# %% [code]
# get relevant samples, do background subtraction
temp_df = master_df.drop(index=master_df[(master_df['DNA/mRNA batch'] != 6) |
                                         (master_df['Extract'] != 'DAJ_MK_MY') |
                                         (master_df['Tetratcycline (ug/mL)'] != 0)
                         ].index)
temp_df.head()

# %% [code]
# get dataframe that contains relevant data
temp_df = temp_df.groupby(['Extract fraction','Fuel', 'DNA (nM)', 'Channel', 'Replicate'])

# create list to store dataframes
integrated_mgapt_dfs = []
max_degfp_dfs = []

# iterate through groups
for name, group in temp_df:
                
    # get relevant information
    extract_fraction = name[0]
    fuel = name[1]
    dna_conc = name[2]
    fluor_mol = name[3]

    # background subtraction
    if fluor_mol == 'MGaptamer':
        bkgd_sub_fn = mgaptamer_bkgd_subt_dict2[list(group['Extract'])[0]]
    elif fluor_mol == 'deGFP':
        bkgd_sub_fn = degfp_bkgd_subt_dict2[list(group['Extract'])[0]]
    group['Measurement'] = group['Measurement'] - bkgd_sub_fn(group['Time (hr)'])   
    
    # drop unrelated columns, create new df
    group = group.drop(columns=['Well'])
    cols = list(group.columns)
    cols.remove('Replicate')
    cols.remove('Measurement')
    new_group = group.groupby(cols).apply(lambda group2: np.mean(group2['Measurement'])).reset_index().ffill()
    new_group = pd.DataFrame(new_group).rename({0:'Measurement'}, axis=1)
    new_group = group.copy()
        
    # perform an integration
    if fluor_mol == 'MGaptamer':
        cols2 = list(new_group.columns)
        cols2.remove('Measurement')
        cols2.remove('Time (hr)')
        cols2.remove('Time (sec)')
        timepoints = sorted(list(set(new_group['Time (hr)'])))
        dx = timepoints[2] - timepoints[1]
        new_group = new_group.groupby(cols2).apply(lambda group2: np.trapz(y=group2['Measurement'], dx=dx)).reset_index().ffill()
        new_group = pd.DataFrame(new_group).rename({0:'Measurement'}, axis=1)
    elif fluor_mol == 'deGFP':
        new_group = new_group.drop(columns=['Time (sec)'])
        cols = list(new_group.columns)
        cols.remove('Measurement')
        cols.remove('Time (hr)')
        new_group = new_group.groupby(cols).apply(lambda group2: np.max(group2['Measurement'])).reset_index().ffill()
        new_group = pd.DataFrame(new_group).rename({0:'Measurement'}, axis=1)
        
    # axis labels
    new_group['Fuel (mM)'] = new_group['Fuel (mM)'].astype('str')
    new_group['Mg (mM)'] = new_group['Mg (mM)'].astype('str')
    
    # drop all but relevant columns
    irrelevant_cols = list(new_group.columns)
    for i in ['Extract fraction', 'DNA (nM)', 'Fuel (mM)', 'Mg (mM)', 'Fuel', 'Measurement', 'Replicate']:
        irrelevant_cols.remove(i)
    new_group = new_group.drop(columns=irrelevant_cols)
    
    # rename measurement column, add relevant data to relevant list of dataframes
    if fluor_mol == 'MGaptamer':
        new_group = pd.DataFrame(new_group).rename({'Measurement':'Integrated MGaptamer (μM-hr)'}, axis=1)
        integrated_mgapt_dfs.append(new_group)
    elif fluor_mol == 'deGFP':
        new_group = pd.DataFrame(new_group).rename({'Measurement':'Maximum deGFP (μM)'}, axis=1)
        max_degfp_dfs.append(new_group)

# %% [code]
# combine list of dataframes into single dataframe for each channel
combined_integrated_mgapt_df = pd.concat(integrated_mgapt_dfs)
combined_max_degfp_df = pd.concat(max_degfp_dfs)

# combine all data into one dataframe via merge
max_degfp_integrated_mgapt_df = combined_integrated_mgapt_df.merge(combined_max_degfp_df, on=['Fuel', 'DNA (nM)', 'Mg (mM)', 'Fuel (mM)', 'Extract fraction', 'Replicate'])
max_degfp_integrated_mgapt_df.head()

# %% [markdown]
### Fig S3a

# %% [code]
# create holoviews plot
color_legend_title = 'Extract fraction'

# get lysate ratio samples at 5nM DNA
plotting_df = max_degfp_integrated_mgapt_df.loc[max_degfp_integrated_mgapt_df['DNA (nM)']==5]
plotting_df['Extract fraction'] = plotting_df['Extract fraction'].astype('str')
plotting_df[color_legend_title] = pd.Categorical(plotting_df[color_legend_title],categories=['0.25', '0.33', '0.41'])
plotting_df = plotting_df.sort_values(color_legend_title)

# plot data
hv_fig = hv.Points(data=plotting_df,
                    kdims=[('Integrated MGaptamer (μM-hr)', 'Integrated MG aptamer (μM-hr)'), 'Maximum deGFP (μM)'],
                    vdims=['Fuel', 'DNA (nM)', 'Mg (mM)', 'Fuel (mM)', 'Extract fraction'],
                   
).opts(
    frame_height=1000,
    frame_width=1200,
    show_grid=False,
    cmap=list(bokeh.palettes.Viridis[3]),
    color=color_legend_title,
    size=20,
    # xlim=xlim,
    xlim=(0, 2.2),
    ylim=ylim,
    # tools=['hover'],
    # xformatter='%.0f',
    legend_opts={'title': 'Extract fraction',
                 'label_text_font_size':'45pt',
                 'title_text_font_size':'50pt',
                 'title_text_font_style': 'normal',
                 'glyph_width':50,
                 'glyph_height':50
                }
)

# render as bokeh figure, format
hv_fig = hv.render(hv_fig)
hv_fig.legend.spacing = 25

# bokeh.io.show(hv_fig)

# save plot
file_save = '..\\Figures\\FigS3_a.png'
bokeh.io.reset_output()
bokeh.io.curdoc().theme = theme
bokeh.io.export_png(hv_fig, filename = file_save, webdriver=webdriver)

# %% [code]
# create holoviews heatmap
labels = ['b', 'c', 'd']

# get approrpriate data
plotting_df = max_degfp_integrated_mgapt_df.loc[max_degfp_integrated_mgapt_df['DNA (nM)']==5]
plotting_df['Extract fraction'] = plotting_df['Extract fraction'].astype('str')

# iterate through extract fractions
for i, value in enumerate(['0.25', '0.33', '0.41']):

    # plot points colored by Mg
    color_legend_title = 'Fuel (mM)'
    plotting_df2 = plotting_df.copy()
    plotting_df2 = plotting_df2.loc[plotting_df2['Extract fraction']==value]
    plotting_df2[color_legend_title] = pd.Categorical(plotting_df2[color_legend_title],categories=['0', '5', '10', '15', '20', '30', '45'])
    plotting_df2 = plotting_df2.sort_values(color_legend_title)
    
    hv_fig = hv.Points(data=plotting_df2,
                        kdims=[('Integrated MGaptamer (μM-hr)', 'Integrated MG aptamer (μM-hr)'), 'Maximum deGFP (μM)'],
                        vdims=['Fuel', 'DNA (nM)', 'Mg (mM)', 'Fuel (mM)', 'Extract fraction'],
                       
    ).opts(
        frame_height=1000,
        frame_width=1200,
        show_grid=False,
        cmap=list(bokeh.palettes.Viridis[7]),
        color=color_legend_title,
        size=20,
        xlim=(0, 2.2),
        # ylim=(-2, 33),
        # xlim=xlim,
        ylim=ylim,
        # xformatter='%.0f',
        legend_opts={'title': '3PGA (mM)',
                     'label_text_font_size':'45pt',
                     'title_text_font_size':'50pt',
                     'title_text_font_style': 'normal',
                     'glyph_width':50,
                     'glyph_height':50
                    }
    )
    
    # render as bokeh figure, format
    hv_fig = hv.render(hv_fig)
    hv_fig.legend.spacing = 25
    mytext = Label(x=1.6, y=3, text=f'Fₗ = {value}', x_units='data', y_units='data', text_font_size='55pt', text_font_style='bold')
    
    hv_fig.add_layout(mytext)
    # bokeh.io.show(hv_fig)
    
    # save plot
    file_save = f'..\\Figures\\FigS3_{labels[i]}.png'
    bokeh.io.reset_output()
    bokeh.io.curdoc().theme = theme
    bokeh.io.export_png(hv_fig, filename = file_save)

    # compute statistics to show strength of relation
    print(f'Cell lysate fraction: {i}')
    X = np.array(plotting_df2['Fuel (mM)'].astype(int)).reshape(-1, 1)
    Y = np.array([plotting_df2['Integrated MGaptamer (μM-hr)'], plotting_df2['Maximum deGFP (μM)']]).T
    spearman_rho_bootstrap_values(X, Y)
    print('\n')

# %% [code]
def figS3():
    
    # Read in PNG images
    png_paths = ["..\\Figures\\FigS3_a.png", 
                "..\\Figures\\FigS3_b.png",
                "..\\Figures\\FigS3_c.png",
                "..\\Figures\\FigS3_d.png"]
    
    # Create Divs for each subfigure
    divs = []
    for i, path in enumerate(png_paths):
        with open(path, "rb") as f:
            img_data = f.read()
        img_src = f"data:image/png;base64,{b64encode(img_data).decode()}"
        div_content = f'<img src="{img_src}">' # style="max-width:100%; max-height:100%">'
        div = Div(text=div_content)
        divs.append(div)
    
    # create panel of images
    panel = column(row([row([p_label_a, divs[0]]), row(Spacer(width=100)), row([p_label_b, divs[1]])]),
                   Spacer(height=100),
                   row([row([p_label_c, divs[2]]), row(Spacer(width=100)), row([p_label_d, divs[3]])]),
                  )
    
    # save plot
    file_save = '..\\Figures\\FigS3.png'
    bokeh.io.reset_output()
    bokeh.io.curdoc().theme = theme
    bokeh.io.export_png(panel, filename = file_save)

# %% [code]
figS3()

# %% [markdown]
#

# %% [markdown]
# Figure 5

# %% [markdown]
### Fig 5a

# %% [code]
# create holoviews plot
color_legend_title = 'DNA (nM)'

# get lysate ratio samples at 5nM DNA
plotting_df = max_degfp_integrated_mgapt_df.loc[(max_degfp_integrated_mgapt_df['Extract fraction']==0.33) & (max_degfp_integrated_mgapt_df['DNA (nM)']!=0)]
plotting_df[color_legend_title] = plotting_df[color_legend_title].astype('str')

plotting_df[color_legend_title] = pd.Categorical(plotting_df[color_legend_title],categories=['1.0', '2.5', '5.0', '7.5', '10.0'])
plotting_df = plotting_df.sort_values(color_legend_title)

# plot data
hv_fig = hv.Points(data=plotting_df,
                    kdims=[('Integrated MGaptamer (μM-hr)', 'Integrated MG aptamer (μM-hr)'), 'Maximum deGFP (μM)'],
                    vdims=['Fuel', 'DNA (nM)', 'Mg (mM)', 'Fuel (mM)', 'Extract fraction'],
                   
).opts(
    frame_height=1000,
    frame_width=1200,
    show_grid=False,
    cmap=list(bokeh.palettes.Viridis[5]),
    color=color_legend_title,
    size=20,
    xlim = (0, 3.1),
    ylim = (-1, 44),
    # tools=['hover'],
    # xformatter='%.0f',
    legend_opts={'title': color_legend_title,
                 'label_text_font_size':'45pt',
                 'title_text_font_size':'50pt',
                 'title_text_font_style': 'normal',
                 'glyph_width':50,
                 'glyph_height':50
                }
)

# render as bokeh figure, format
hv_fig = hv.render(hv_fig)
hv_fig.legend.spacing = 25

# save plot
file_save = '..\\Figures\\Fig5_a.png'
bokeh.io.reset_output()
bokeh.io.curdoc().theme = theme
bokeh.io.export_png(hv_fig, filename = file_save, webdriver=webdriver)

# %% [markdown]
### Fig 5b

# %% [code]
# create holoviews plot
color_legend_title = 'Fuel (mM)'

# get lysate ratio samples at 5nM DNA
plotting_df = max_degfp_integrated_mgapt_df.loc[(max_degfp_integrated_mgapt_df['Extract fraction']==0.33) & (max_degfp_integrated_mgapt_df['DNA (nM)']!=0)]
plotting_df[color_legend_title] = plotting_df[color_legend_title].astype('str')

plotting_df[color_legend_title] = pd.Categorical(plotting_df[color_legend_title],categories=['0', '5', '10', '15', '20', '30', '45'])
plotting_df = plotting_df.sort_values(color_legend_title)

# plot data
hv_fig = hv.Points(data=plotting_df,
                    kdims=['Integrated MGaptamer (μM-hr)', 'Maximum deGFP (μM)'],
                    vdims=['Fuel', 'DNA (nM)', 'Mg (mM)', 'Fuel (mM)', 'Extract fraction'],
                   
).opts(
    frame_height=1000,
    frame_width=1200,
    show_grid=False,
    cmap=list(bokeh.palettes.Viridis[7]),
    color=color_legend_title,
    size=20,
    xlim = (0, 3.1),
    ylim = (-1, 44),
    # tools=['hover'],
    # xformatter='%.0f',
    legend_opts={'title': '3PGA (mM)',
                 'label_text_font_size':'45pt',
                 'title_text_font_size':'50pt',
                 'title_text_font_style': 'normal',
                 'glyph_width':50,
                 'glyph_height':50
                }
)

# render as bokeh figure, format
hv_fig = hv.render(hv_fig)
hv_fig.legend.spacing = 25

# bokeh.io.show(hv_fig)

# save plot
file_save = '..\\Figures\\Fig5_b.png'
bokeh.io.reset_output()
bokeh.io.curdoc().theme = theme
bokeh.io.export_png(hv_fig, filename = file_save, webdriver=webdriver)

# %% [code]
# iterate through dna concs
for i in [1.0, 2.5, 5, 7.5, 10]:
    plotting_df2 = plotting_df.loc[plotting_df['DNA (nM)']==i]

    # print dna
    print(f'DNA concentration (nM): {i}')
    
    # compute statistics to show strength of relation
    X = np.array(plotting_df2['Fuel (mM)'].astype(int)).reshape(-1, 1)
    Y = np.array([plotting_df2['Integrated MGaptamer (μM-hr)'], plotting_df2['Maximum deGFP (μM)']]).T
    spearman_rho_bootstrap_values(X, Y)

    print('\n')

# %% [code]
def fig5():
    
    # Read in PNG images
    png_paths = ["..\\Figures\\Fig5_a.png", 
                   "..\\Figures\\Fig5_b.png"]
    
    # Create Divs for each subfigure
    divs = []
    for i, path in enumerate(png_paths):
        with open(path, "rb") as f:
            img_data = f.read()
        img_src = f"data:image/png;base64,{b64encode(img_data).decode()}"
        div_content = f'<img src="{img_src}">' # style="max-width:100%; max-height:100%">'
        div = Div(text=div_content)
        divs.append(div)
    
    # create panel of images
    panel = row([row([p_label_a, divs[0]]), row(Spacer(width=100)), row([p_label_b, divs[1]])])
    
    # save plot
    file_save = '..\\Figures\\Fig5.png'
    bokeh.io.reset_output()
    bokeh.io.curdoc().theme = theme
    bokeh.io.export_png(panel, filename = file_save)

# %% [code]
fig5()

# %% [markdown]
#

# %% [markdown]
# Figure S6

# %% [markdown]
### Fig S6a

# %% [code]
# create holoviews plot
color_legend_title = 'DNA (nM)'

# get lysate ratio samples at 5nM DNA
plotting_df = max_degfp_integrated_mgapt_df.loc[(max_degfp_integrated_mgapt_df['Extract fraction']==0.33) & (max_degfp_integrated_mgapt_df['DNA (nM)']!=0)]
plotting_df[color_legend_title] = plotting_df[color_legend_title].astype('str')

plotting_df[color_legend_title] = pd.Categorical(plotting_df[color_legend_title],categories=['1.0', '2.5', '5.0', '7.5', '10.0'])
plotting_df = plotting_df.sort_values(color_legend_title)

# plot data
hv_fig = hv.Points(data=plotting_df,
                    kdims=[('Integrated MGaptamer (μM-hr)', 'Integrated MG aptamer (μM-hr)'),'Maximum deGFP (μM)'],
                    vdims=['Fuel', 'DNA (nM)', 'Mg (mM)', 'Fuel (mM)', 'Extract fraction'],
                   
).opts(
    frame_height=1000,
    frame_width=1200,
    show_grid=False,
    color=list(bokeh.palettes.Greys[6])[4],
    size=20,
    xlim = (0, 3.1),
    ylim = (-1, 44)
)

# Plot the regression line
k = 0
h = np.min(plotting_df['Integrated MGaptamer (μM-hr)'].loc[plotting_df['DNA (nM)']!='0.0'])
x_line = np.array([h, 3])
slopes = []
for i in [0, 20, 40, 59, 80]:
    slopes.append(np.sin(np.deg2rad(i))/np.cos(np.deg2rad(i))*14)

# define linear function
def linear_function_y(x, slope, h, k):
    return(k + slope*(x-h))
    
# iterate through slopes
for index,slope in enumerate(slopes):

    # get data for max tx for each condition
    if index != 4:
        plotting_df2 = plotting_df.loc[((plotting_df['Maximum deGFP (μM)'] >= linear_function_y(plotting_df['Integrated MGaptamer (μM-hr)'], slope, h, k)) &
                                       (plotting_df['Maximum deGFP (μM)'] < linear_function_y(plotting_df['Integrated MGaptamer (μM-hr)'], slopes[index+1], h, k))
                                      )]
    else:
        plotting_df2 = plotting_df.loc[(plotting_df['Maximum deGFP (μM)'] >= linear_function_y(plotting_df['Integrated MGaptamer (μM-hr)'], slope, h, k))]

    # iterate through DNA concs
    for dna_conc in ['1.0', '2.5', '5.0', '7.5', '10.0']:

        # plot maximum mgapt point for each cutoff
        temp_df = plotting_df2.loc[plotting_df2['DNA (nM)']==dna_conc]
        temp_df = temp_df.loc[temp_df['Integrated MGaptamer (μM-hr)'] == np.max(temp_df['Integrated MGaptamer (μM-hr)'])]

        # plot point
        hv_fig2 = hv.Points(data=temp_df,
                            kdims=['Integrated MGaptamer (μM-hr)', 'Maximum deGFP (μM)'],
                            vdims=['Fuel', 'DNA (nM)', 'Mg (mM)', 'Fuel (mM)', 'Extract fraction'],
                           
        ).opts(
            frame_height=1000,
            frame_width=1200,
            show_grid=False,
            color=list(bokeh.palettes.Colorblind[5])[index],
            size=20,
            fill_color = 'white',
            line_width=10,
        )

        # add point to plot
        hv_fig = hv_fig*hv_fig2


# render as bokeh figure, format
hv_fig = hv.render(hv_fig)
hv_fig.legend.spacing = 25

# Plot the regression line
k = 0
h = np.min(plotting_df['Integrated MGaptamer (μM-hr)'])
x_line = np.array([h, 3])
slopes = []
for i in [0, 20, 40, 59, 80]:
    slopes.append(np.sin(np.deg2rad(i))/np.cos(np.deg2rad(i))*14)
for index,slope in enumerate(slopes):    
    y_line = k + slope * (x_line - h)
    hv_fig.line(x_line, y_line, color=list(bokeh.palettes.Colorblind[5])[index], line_width=10, line_dash = 'dashed')

# add labels for dashed lines
xy_vals = [(2.9, 3), (2.2, 13), (1.70, 20), (1.15, 28), (0.4, 38.5)]
xy_vals = [(2.9, 3), (2.7, 15), (2.20, 26), (1.4, 35), (0.4, 38.5)]
for index, text in enumerate(np.linspace(1,5,5)):
    x, y = xy_vals[index]
    x = x-0.1
    # y = y - 4
    mytext = Label(x=x, y=y, text=f'{int(text)}', x_units='data', y_units='data', text_font_size='80pt', text_font_style='bold', text_color=list(bokeh.palettes.Colorblind[5])[index])
    hv_fig.add_layout(mytext)

# remove labels
hv_fig.legend.visible = False

# save plot
file_save = '..\\Figures\\FigS6_a.png'
bokeh.io.reset_output()
bokeh.io.curdoc().theme = theme
bokeh.io.export_png(hv_fig, filename = file_save, webdriver=webdriver)

# %% [markdown]
### Fig S5b

# %% [code]
# plotting things
xlim = (-0.4, 11.9)
ylim=(-0.1, 3.2)

# get lysate ratio samples at 5nM DNA
color_legend_title = 'DNA categorical'
plotting_df = max_degfp_integrated_mgapt_df.loc[(max_degfp_integrated_mgapt_df['Extract fraction']==0.33) & (max_degfp_integrated_mgapt_df['DNA (nM)']!=0)]
plotting_df = max_degfp_integrated_mgapt_df.loc[(max_degfp_integrated_mgapt_df['Extract fraction']==0.33)]

plotting_df['Mg (mM)'] = plotting_df['Mg (mM)'].astype('int')
plotting_df['Fuel (mM)'] = plotting_df['Fuel (mM)'].astype('int')
plotting_df['DNA categorical'] = plotting_df['DNA (nM)']
plotting_df[color_legend_title] = plotting_df[color_legend_title].astype('str')
plotting_df[color_legend_title] = pd.Categorical(plotting_df[color_legend_title],categories=['0.0','1.0', '2.5', '5.0', '7.5', '10.0'])
plotting_df = plotting_df.sort_values(color_legend_title)

# create plot
hv_fig = hv.Points(data=plotting_df,
                    kdims=['DNA (nM)',('Integrated MGaptamer (μM-hr)', 'Integrated MG aptamer (μM-hr)')],
                    vdims=['Fuel', 'DNA (nM)', 'Mg (mM)', 'Fuel (mM)', 'DNA categorical'],
                   
).opts(
    frame_height=1000,
    frame_width=1200,
    show_grid=False,
    # cmap=list(bokeh.palettes.Viridis[6]),
    color= '#444444',
    size=30,
    # tools=['hover'],
    xformatter='%.0f',
    # fill_alpha = 0,
    line_width = 5,
    xlim=xlim,
    ylim=ylim
)


# Plot the regression line
k = 0
h = np.min(plotting_df['Integrated MGaptamer (μM-hr)'].loc[plotting_df['DNA (nM)']!=0.0])
x_line = np.array([h, 3])
slopes = []
for i in [0, 20, 40, 59, 80]:
    slopes.append(np.sin(np.deg2rad(i))/np.cos(np.deg2rad(i))*14)

# define linear function
def linear_function_y(x, slope, h, k):
    return(k + slope*(x-h))

# define linear function
def linear_function_x(y, slope, h, k):
    return(h + (y-k)/slope)
    
# iterate through slopes
for index,slope in enumerate(slopes):

    # get data for max tx for each condition
    plotting_df = max_degfp_integrated_mgapt_df.loc[(max_degfp_integrated_mgapt_df['Extract fraction']==0.33) & (max_degfp_integrated_mgapt_df['DNA (nM)']!=0.0)]
    if index != 4:
        plotting_df2 = plotting_df.loc[((plotting_df['Maximum deGFP (μM)'] >= linear_function_y(plotting_df['Integrated MGaptamer (μM-hr)'], slope, h, k)) &
                                       (plotting_df['Maximum deGFP (μM)'] < linear_function_y(plotting_df['Integrated MGaptamer (μM-hr)'], slopes[index+1], h, k))
                                      )]
    else:
        plotting_df2 = plotting_df.loc[(plotting_df['Maximum deGFP (μM)'] >= linear_function_y(plotting_df['Integrated MGaptamer (μM-hr)'], slope, h, k))]


    # format data, pick max mgapt
    plotting_df2 = plotting_df2.groupby('DNA (nM)').apply(lambda group2: np.max(group2['Integrated MGaptamer (μM-hr)'])).reset_index().ffill()
    plotting_df2 = pd.DataFrame(plotting_df2).rename({0:'Integrated MGaptamer (μM-hr)'}, axis=1)


    # create plot
    hv_fig2 = hv.Points(data=plotting_df2,
                        kdims=['DNA (nM)',('Integrated MGaptamer (μM-hr)', 'Integrated MG aptamer (μM-hr)')],
                       
    ).opts(
        frame_height=1000,
        frame_width=1200,
        show_grid=False,
        # cmap=list(bokeh.palettes.Viridis[6]),
        # color= color_legend_title,
        size=30,
        # tools=['hover'],
        # xformatter='%.0f',
        xlim=xlim,
        ylim=ylim,
        # marker='square',
        line_color = list(bokeh.palettes.Colorblind[5])[index],
        fill_color = 'white',
        # fill_alpha=0,
        line_width = 10,
    )
    
    # create plot
    k = 0
    h = np.min(plotting_df['Integrated MGaptamer (μM-hr)'].loc[plotting_df['DNA (nM)']!='0.0'])
    plotting_df = max_degfp_integrated_mgapt_df.loc[(max_degfp_integrated_mgapt_df['Extract fraction']==0.33)]
    if index != 4:
        plotting_df2 = plotting_df.loc[((plotting_df['Maximum deGFP (μM)'] >= linear_function_y(plotting_df['Integrated MGaptamer (μM-hr)'], slope, h, k)) &
                                       (plotting_df['Maximum deGFP (μM)'] < linear_function_y(plotting_df['Integrated MGaptamer (μM-hr)'], slopes[index+1], h, k))
                                      )]
    else:
        plotting_df2 = plotting_df.loc[(plotting_df['Maximum deGFP (μM)'] >= linear_function_y(plotting_df['Integrated MGaptamer (μM-hr)'], slope, h, k))]
    plotting_df2 = plotting_df2.groupby('DNA (nM)').apply(lambda group2: np.max(group2['Integrated MGaptamer (μM-hr)'])).reset_index().ffill()
    plotting_df2 = pd.DataFrame(plotting_df2).rename({0:'Integrated MGaptamer (μM-hr)'}, axis=1)

    # add zero if not there
    if 0.0 not in set(list(plotting_df2['DNA (nM)'])):
        plotting_df2.loc[-1] = [0.0, 0.074226]  # adding a row
        plotting_df2.index = plotting_df2.index + 1  # shifting index
        plotting_df2 = plotting_df2.sort_index()  # sorting by index

    hv_fig3 = hv.Curve(data=plotting_df2,
                        kdims=['DNA (nM)'],
                        vdims=[('Integrated MGaptamer (μM-hr)', 'Integrated MG aptamer (μM-hr)')]
    ).opts(
        frame_height=1000,
        frame_width=1200,
        show_grid=False,
        line_color = list(bokeh.palettes.Colorblind[5])[index],
        line_width = 10,
        line_dash = 'dashed'
    )

    hv_fig4 =  hv.Points(data=plotting_df2.loc[plotting_df2['DNA (nM)']==0.0],
                        kdims=['DNA (nM)',('Integrated MGaptamer (μM-hr)', 'Integrated MG aptamer (μM-hr)')],
                       
    ).opts(
        frame_height=1000,
        frame_width=1200,
        show_grid=False,
        size=30,
        line_color = 'black',
        fill_color = 'white',
        line_width = 10,
    )


    if index==0:
        hv_fig = hv_fig3*hv_fig2*hv_fig4
    else:
        hv_fig = hv_fig*hv_fig3*hv_fig2*hv_fig4

# render as bokeh figure, format
hv_fig = hv.render(hv_fig)

# add labels for dashed lines
x = 10.5
y_vals = [2.8, 1.7, 1.2, 0.8, 0.3]
for index, text in enumerate(np.linspace(1,5,5)):
    y = y_vals[index]
    mytext = Label(x=x, y=y, text=f'{int(text)}', x_units='data', y_units='data', text_font_size='80pt', text_font_style='bold', text_color=list(bokeh.palettes.Colorblind[5])[index])
    hv_fig.add_layout(mytext)

# save plot
file_save = '..\\Figures\\FigS6_b.png'
bokeh.io.reset_output()
bokeh.io.curdoc().theme = theme
bokeh.io.export_png(hv_fig, filename = file_save, webdriver=webdriver)

# %% [markdown]
### Fig S5c

# %% [code]
# get lysate ratio samples at 5nM DNA
color_legend_title = 'DNA categorical'
plotting_df = max_degfp_integrated_mgapt_df.loc[(max_degfp_integrated_mgapt_df['Extract fraction']==0.33) & (max_degfp_integrated_mgapt_df['DNA (nM)']!=0)]
plotting_df = max_degfp_integrated_mgapt_df.loc[(max_degfp_integrated_mgapt_df['Extract fraction']==0.33)]
plotting_df['Mg (mM)'] = plotting_df['Mg (mM)'].astype('int')
plotting_df['Fuel (mM)'] = plotting_df['Fuel (mM)'].astype('int')
plotting_df['DNA categorical'] = plotting_df['DNA (nM)']
plotting_df[color_legend_title] = plotting_df[color_legend_title].astype('str')
plotting_df[color_legend_title] = pd.Categorical(plotting_df[color_legend_title],categories=['0.0','1.0', '2.5', '5.0', '7.5', '10.0'])
plotting_df = plotting_df.sort_values(color_legend_title)

# create plot
hv_fig = hv.Points(data=plotting_df,
                    kdims=['DNA (nM)','Maximum deGFP (μM)'],
                    vdims=['Fuel', 'DNA (nM)', 'Mg (mM)', 'Fuel (mM)', 'DNA categorical'],
                   
).opts(
    frame_height=1000,
    frame_width=1200,
    show_grid=False,
    # cmap=list(bokeh.palettes.Viridis[6]),
    color= '#444444',
    size=30,
    # tools=['hover'],
    xformatter='%.0f',
    # fill_alpha = 0,
    line_width = 5,
    xlim=(-0.4, 11.9),
)


# Plot the regression line
k = 0
h = np.min(plotting_df['Integrated MGaptamer (μM-hr)'].loc[plotting_df['DNA (nM)']!=0.0])
x_line = np.array([h, 3])
slopes = []
for i in [0, 20, 40, 59, 80]:
    slopes.append(np.sin(np.deg2rad(i))/np.cos(np.deg2rad(i))*14)

# define linear function
def linear_function_y(x, slope, h, k):
    return(k + slope*(x-h))

# define linear function
def linear_function_x(y, slope, h, k):
    return(h + (y-k)/slope)
    
# iterate through slopes
for index,slope in enumerate(slopes):

    # get data for max tx for each condition
    plotting_df = max_degfp_integrated_mgapt_df.loc[(max_degfp_integrated_mgapt_df['Extract fraction']==0.33) & (max_degfp_integrated_mgapt_df['DNA (nM)']!=0.0)]

    # PLOT POINTS
    if index != 4:
        plotting_df2 = plotting_df.loc[((plotting_df['Maximum deGFP (μM)'] >= linear_function_y(plotting_df['Integrated MGaptamer (μM-hr)'], slope, h, k)) &
                                       (plotting_df['Maximum deGFP (μM)'] < linear_function_y(plotting_df['Integrated MGaptamer (μM-hr)'], slopes[index+1], h, k))
                                      )]
    else:
        plotting_df2 = plotting_df.loc[(plotting_df['Maximum deGFP (μM)'] >= linear_function_y(plotting_df['Integrated MGaptamer (μM-hr)'], slope, h, k))]

    # format data, pick max mgapt
    plotting_df2 = plotting_df2.groupby('DNA (nM)').apply(lambda group2: np.max(group2.loc[group2['Integrated MGaptamer (μM-hr)'].idxmax()]['Maximum deGFP (μM)'])).reset_index().ffill()
    plotting_df2 = pd.DataFrame(plotting_df2).rename({0:'Maximum deGFP (μM)'}, axis=1)


    # create plot
    hv_fig2 = hv.Points(data=plotting_df2,
                        kdims=['DNA (nM)','Maximum deGFP (μM)'],
                       
    ).opts(
        frame_height=1000,
        frame_width=1200,
        show_grid=False,
        size=30,
        xlim=(-0.4, 11.9),
        line_color = list(bokeh.palettes.Colorblind[5])[index],
        fill_color = 'white',
        line_width = 10,
    )
    
    # get data to make curve
    k = 0
    h = np.min(plotting_df['Integrated MGaptamer (μM-hr)'].loc[plotting_df['DNA (nM)']!='0.0'])
    plotting_df = max_degfp_integrated_mgapt_df.loc[(max_degfp_integrated_mgapt_df['Extract fraction']==0.33)]
    if index != 4:
        plotting_df2 = plotting_df.loc[((plotting_df['Maximum deGFP (μM)'] >= linear_function_y(plotting_df['Integrated MGaptamer (μM-hr)'], slope, h, k)) &
                                       (plotting_df['Maximum deGFP (μM)'] < linear_function_y(plotting_df['Integrated MGaptamer (μM-hr)'], slopes[index+1], h, k))
                                      )]
    else:
        plotting_df2 = plotting_df.loc[(plotting_df['Maximum deGFP (μM)'] >= linear_function_y(plotting_df['Integrated MGaptamer (μM-hr)'], slope, h, k))]
    plotting_df2 = plotting_df2.groupby('DNA (nM)').apply(lambda group2: np.max(group2.loc[group2['Integrated MGaptamer (μM-hr)'].idxmax()]['Maximum deGFP (μM)'])).reset_index().ffill()
    plotting_df2 = pd.DataFrame(plotting_df2).rename({0:'Maximum deGFP (μM)'}, axis=1)

    # add zero if not there
    if 0.0 not in set(list(plotting_df2['DNA (nM)'])):
        plotting_df2.loc[-1] = [0.0, 0.022721]  # adding a row
        plotting_df2.index = plotting_df2.index + 1  # shifting index
        plotting_df2 = plotting_df2.sort_index()  # sorting by index

    # ADD LINE
    hv_fig3 = hv.Curve(data=plotting_df2,
                        kdims=['DNA (nM)'],
                        vdims=['Maximum deGFP (μM)'],
    ).opts(
        frame_height=1000,
        frame_width=1200,
        show_grid=False,
        line_color = list(bokeh.palettes.Colorblind[5])[index],
        line_width = 10,
        line_dash = 'dashed'
    )

    # ADD ZERO POINT
    hv_fig4 =  hv.Points(data=plotting_df2.loc[plotting_df2['DNA (nM)']==0.0],
                        kdims=['DNA (nM)','Maximum deGFP (μM)'],
                       
    ).opts(
        frame_height=1000,
        frame_width=1200,
        show_grid=False,
        size=30,
        xlim=(-1,11),
        ylim=(-2, 42),
        # marker='square',
        line_color = 'black',
        fill_color = 'white',
        line_width = 10,
    )


    if index==0:
        hv_fig = hv_fig3*hv_fig2*hv_fig4
    else:
        hv_fig = hv_fig*hv_fig3*hv_fig2*hv_fig4
        
# render as bokeh figure, format
hv_fig = hv.render(hv_fig)

# add labels for dashed lines
x = 10.5
y_vals = [0, 8, 16, 22, 38]
for index, text in enumerate(np.linspace(1,5,5)):
    y = y_vals[index]
    mytext = Label(x=x, y=y, text=f'{int(text)}', x_units='data', y_units='data', text_font_size='80pt', text_font_style='bold', text_color=list(bokeh.palettes.Colorblind[5])[index])
    hv_fig.add_layout(mytext)

# save plot
file_save = '..\\Figures\\FigS6_c.png'
bokeh.io.reset_output()
bokeh.io.curdoc().theme = theme
bokeh.io.export_png(hv_fig, filename = file_save, webdriver=webdriver)

# %% [code]
def figS6():
    
    # Read in PNG images
    png_paths = ["..\\Figures\\FigS6_a.png", 
                "..\\Figures\\FigS6_b.png",
                "..\\Figures\\FigS6_c.png"]
    
    # Create Divs for each subfigure
    divs = []
    for i, path in enumerate(png_paths):
        with open(path, "rb") as f:
            img_data = f.read()
        img_src = f"data:image/png;base64,{b64encode(img_data).decode()}"
        div_content = f'<img src="{img_src}">' # style="max-width:100%; max-height:100%">'
        div = Div(text=div_content)
        divs.append(div)
    
    # create panel of images
    panel = column(row([row([p_label_a, divs[0]]), row(Spacer(width=100)), row([p_label_b, divs[1]])]),
                   Spacer(height=100),
                   row([row([p_label_c, divs[2]]), row(Spacer(width=100))]),
                  )
    
    # save plot
    file_save = '..\\Figures\\FigS6.png'
    bokeh.io.reset_output()
    bokeh.io.curdoc().theme = theme
    bokeh.io.export_png(panel, filename = file_save)

# %% [code]
figS6()

# %% [markdown]
# 

# %% [markdown]
# Figures 4, S4, S5

# %% [code]
# get relevant mgaptamer background subtraction data
# por1or2 construct
por1or2_mgapt_bkgd_samples = master_df.loc[(master_df['DNA (nM)']==0) &
                      (master_df['DNA/mRNA batch'] ==6) &
                      (master_df['Extract']=='DAJ_MK_MY2') & 
                      (master_df['Fuel (mM)']==0) & 
                      (master_df['Mg (mM)']==0) & 
                      (master_df['Plasmid construct']=='pOR1OR2-MGapt-deGFP') & 
                      (master_df['mRNA (uM)'] == 0) & 
                      (master_df['Channel'] == 'MGaptamer')
                    ]
x = por1or2_mgapt_bkgd_samples['Time (hr)']
y = por1or2_mgapt_bkgd_samples['Measurement']
fn = np.polyfit(x, y, 5)
por1or2_mgapt_fn = np.poly1d(fn)
por1or2_degfp_bkgd_samples = master_df.loc[(master_df['DNA (nM)']==0) &
                      (master_df['DNA/mRNA batch'] ==6) &
                      (master_df['Extract']=='DAJ_MK_MY2') & 
                      (master_df['Fuel (mM)']==0) & 
                      (master_df['Mg (mM)']==0) & 
                      (master_df['Plasmid construct']=='pOR1OR2-MGapt-deGFP') & 
                      (master_df['mRNA (uM)'] == 0) & 
                      (master_df['Channel'] == 'deGFP')
                    ]
x = por1or2_degfp_bkgd_samples['Time (hr)']
y = por1or2_degfp_bkgd_samples['Measurement']
fn = np.polyfit(x, y, 5)
por1or2_degfp_fn = np.poly1d(fn)
        
# create dictionary to store functions
bkgd_subt_dict = {'MGaptamer': por1or2_mgapt_fn,
                'deGFP': por1or2_degfp_fn,
                }

# %% [code]
# get relevant samples, do background subtraction
temp_df = master_df.drop(index=master_df[(((master_df['DNA/mRNA batch'] != 6) & 
                                           (master_df['Plasmid construct'] != 'pOR1OR2')) & 
                                          (master_df['Plasmid construct'] != 'pTet-MGapt-deGFP') & 
                                          (master_df['Plasmid construct'] != 'pT7-MGapt-deGFP w/ 10uM T7RNAP') &
                                          (master_df['Plasmid construct'] != 'pT7-deGFP-MGapt w/ 10uM T7RNAP')
                                         ) |
                                         (master_df['Extract'] != 'DAJ_MK_MY2') 
                         ].index).groupby(['Extract','Fuel', 'DNA (nM)', 'Channel','Plasmid construct'])

# create list to store dataframes
integrated_mgapt_dfs = []
max_degfp_dfs = []

# iterate through groups
for name, group in temp_df:
                
    # get relevant information
    extract = name[0]
    fuel = name[1]
    dna_conc = name[2]
    fluor_mol = name[3]
    
    # background subtraction
    bkgd_sub_fn = bkgd_subt_dict[fluor_mol]
    group['Measurement'] = group['Measurement'] - bkgd_sub_fn(group['Time (hr)'])    
    
    # drop unrelated columns, create new df
    group = group.drop(columns=['Well'])
    cols = list(group.columns)
    cols.remove('Replicate')
    cols.remove('Measurement')
    new_group = group.groupby(cols).apply(lambda group2: np.mean(group2['Measurement'])).reset_index().ffill()#fillna(method="ffill")
    new_group = pd.DataFrame(new_group).rename({0:'Measurement'}, axis=1)
    new_group = group.copy()

    if fuel=='None':
        print(new_group.head())
        
    # perform an integration
    if fluor_mol == 'MGaptamer':
        cols2 = list(new_group.columns)
        cols2.remove('Measurement')
        cols2.remove('Time (hr)')
        cols2.remove('Time (sec)')
        timepoints = sorted(list(set(new_group['Time (hr)'])))
        dx = timepoints[2] - timepoints[1]
        new_group = new_group.groupby(cols2).apply(lambda group2: np.trapz(y=group2['Measurement'], dx=dx)).reset_index().ffill()
        new_group = pd.DataFrame(new_group).rename({0:'Measurement'}, axis=1)
    elif fluor_mol == 'deGFP':
        new_group = new_group.drop(columns=['Time (sec)'])
        cols = list(new_group.columns)
        cols.remove('Measurement')
        cols.remove('Time (hr)')
        new_group = new_group.groupby(cols).apply(lambda group2: np.max(group2['Measurement'])).reset_index().ffill()#fillna(method="ffill")
        new_group = pd.DataFrame(new_group).rename({0:'Measurement'}, axis=1)
        
    # axis labels
    new_group['Fuel (mM)'] = new_group['Fuel (mM)'].astype('str')
    new_group['Mg (mM)'] = new_group['Mg (mM)'].astype('str')
    
    # drop all but relevant columns
    irrelevant_cols = list(new_group.columns)
    for i in ['Extract', 'DNA (nM)', 'Fuel (mM)', 'Mg (mM)', 'Fuel', 'Measurement', 'Plasmid construct', 'Replicate']:
        irrelevant_cols.remove(i)
    new_group = new_group.drop(columns=irrelevant_cols)
    
    # rename measurement column, add relevant data to relevant list of dataframes
    if fluor_mol == 'MGaptamer':
        new_group = pd.DataFrame(new_group).rename({'Measurement':'Integrated MGaptamer (μM-hr)'}, axis=1)
        integrated_mgapt_dfs.append(new_group)
    elif fluor_mol == 'deGFP':
        new_group = pd.DataFrame(new_group).rename({'Measurement':'Maximum deGFP (μM)'}, axis=1)
        max_degfp_dfs.append(new_group)

# %% [code]
# combine list of dataframes into single dataframe for each channel
combined_integrated_mgapt_df = pd.concat(integrated_mgapt_dfs)
combined_max_degfp_df = pd.concat(max_degfp_dfs)

# combine all data into one dataframe via merge
max_degfp_integrated_mgapt_df = combined_integrated_mgapt_df.merge(combined_max_degfp_df, on=['Fuel', 'DNA (nM)', 'Mg (mM)', 'Fuel (mM)', 'Extract', 'Plasmid construct', 'Replicate'])
max_degfp_integrated_mgapt_df.head()

# %% [code]
# get promoters
max_degfp_integrated_mgapt_df['Promoter'] = [i.split('-')[0] for i in max_degfp_integrated_mgapt_df['Plasmid construct']]
max_degfp_integrated_mgapt_df.head()

# %% [code]
# ranges for plots
xlim2 = (0, 2.4)
ylim2 = (-0.56, 18)

# %% [code]
# create holoviews plot
color_legend_title = 'Promoter'

# get lysate ratio samples at 5nM DNA
plotting_df = max_degfp_integrated_mgapt_df.loc[(max_degfp_integrated_mgapt_df['DNA (nM)']==5) &
                                                (max_degfp_integrated_mgapt_df['Plasmid construct']!='pT7-deGFP-MGapt w/ 10uM T7RNAP')
                                                ]

# plot data
hv_fig = hv.Points(data=plotting_df,
                    kdims=[('Integrated MGaptamer (μM-hr)', 'Integrated MG aptamer (μM-hr)'), 'Maximum deGFP (μM)'],
                    vdims=['Fuel', 'DNA (nM)', 'Mg (mM)', 'Fuel (mM)', 'Promoter', 'Plasmid construct'],
                   
).opts(
    frame_height=1000,
    frame_width=1200,
    show_grid=False,
    cmap=['#000000','#D55E00','#F0E442'],
    color=color_legend_title,
    size=20,
    # xlim=(0.15, 2.6),
    # ylim=(-2, 19),
    xlim=xlim2,
    ylim=ylim2,
    # tools=['hover'],
    # xformatter='%.0f',
    legend_opts={'title': color_legend_title,
                 'label_text_font_size':'45pt',
                 'title_text_font_size':'50pt',
                 'title_text_font_style': 'normal',
                 'glyph_width':50,
                 'glyph_height':50
                }
)

# render as bokeh figure, format
hv_fig = hv.render(hv_fig)
hv_fig.legend.spacing = 25

# bokeh.io.show(hv_fig)

# save plot
file_save = '..\\Figures\\FigS5.png'
bokeh.io.reset_output()
bokeh.io.curdoc().theme = theme
bokeh.io.export_png(hv_fig, filename = file_save, webdriver=webdriver)

# %% [code]
# create holoviews plot
color_legend_title = 'Plasmid construct'

# get lysate ratio samples at 5nM DNA
plotting_df = max_degfp_integrated_mgapt_df.loc[(max_degfp_integrated_mgapt_df['DNA (nM)']==5) &
                                                (max_degfp_integrated_mgapt_df['Plasmid construct']!='pOR1OR2-MGapt-deGFP') &
                                                (max_degfp_integrated_mgapt_df['Plasmid construct']!='pTet-MGapt-deGFP')
                                                ]
plotting_df['Plasmid construct'] = [i.split(' w/')[0] for i in plotting_df['Plasmid construct']]


# plot data
hv_fig = hv.Points(data=plotting_df,
                    kdims=[('Integrated MGaptamer (μM-hr)', 'Integrated MG aptamer (μM-hr)'), 'Maximum deGFP (μM)'],
                    vdims=['Fuel', 'DNA (nM)', 'Mg (mM)', 'Fuel (mM)', 'Promoter', 'Plasmid construct'],
                   
).opts(
    frame_height=1000,
    frame_width=1200,
    show_grid=False,
    cmap=['#000000','#D55E00','#F0E442'],
    color=color_legend_title,
    size=20,
    # xlim=(0.15, 2.6),
    # ylim=(-2, 19),
    xlim=xlim2,
    ylim=(-0.56, 28.5),
    # tools=['hover'],
    # xformatter='%.0f',
    legend_opts={'title': color_legend_title,
                 'label_text_font_size':'45pt',
                 'title_text_font_size':'50pt',
                 'title_text_font_style': 'normal',
                 'glyph_width':50,
                 'glyph_height':50
                }
)

# render as bokeh figure, format
hv_fig = hv.render(hv_fig)
hv_fig.legend.spacing = 25

# bokeh.io.show(hv_fig)

# save plot
file_save = '..\\Figures\\FigS4.png'
bokeh.io.reset_output()
bokeh.io.curdoc().theme = theme
bokeh.io.export_png(hv_fig, filename = file_save, webdriver=webdriver)

# %% [code]
# create holoviews heatmap
labels = ['a', 'c', 'b']

# get appropriate data
plotting_df = max_degfp_integrated_mgapt_df.loc[(max_degfp_integrated_mgapt_df['DNA (nM)']==5) &
                                                (max_degfp_integrated_mgapt_df['Plasmid construct']!='pT7-deGFP-MGapt w/ 10uM T7RNAP')
                                                ]

# iterate through extract fractions
for i, value in enumerate(['pOR1OR2', 'pTet', 'pT7']):

    # plot points colored by Mg
    color_legend_title = 'Fuel (mM)'
    plotting_df2 = plotting_df.copy()
    plotting_df2 = plotting_df2.loc[plotting_df2['Promoter']==value]
    
    hv_fig = hv.Points(data=plotting_df2,
                        kdims=[('Integrated MGaptamer (μM-hr)', 'Integrated MG aptamer (μM-hr)'), 'Maximum deGFP (μM)'],
                        vdims=['Fuel', 'DNA (nM)', 'Mg (mM)', 'Fuel (mM)'],
                       
    ).opts(
        frame_height=1000,
        frame_width=1200,
        show_grid=False,
        cmap=list(bokeh.palettes.Viridis[7]),
        color=color_legend_title,
        size=20,
        # xlim=(0.15, 2.6),
        # ylim=(-2, 19),
        xlim=xlim2,
        ylim=ylim2,
        # xformatter='%.0f',
        
        legend_opts={'title': '3PGA (mM)',
                     'label_text_font_size':'45pt',
                     'title_text_font_size':'50pt',
                     'title_text_font_style': 'normal',
                     'glyph_width':50,
                     'glyph_height':50
                    }
    )
    
    # render as bokeh figure, format
    hv_fig = hv.render(hv_fig)
    hv_fig.legend.spacing = 25

    # add label
    if i<2:
        mytext = Label(x=1.7, y=3, text=f'{value}', x_units='data', y_units='data', text_font_size='55pt', text_font_style='bold')
    else:
        mytext = Label(x=0.1, y=0.5, text=f'{value}', x_units='data', y_units='data', text_font_size='55pt', text_font_style='bold')
    
    # hv_fig.add_layout(mytext)
    # bokeh.io.show(hv_fig)
    
    # save plot
    file_save = f'..\\Figures\\Fig4_{labels[i]}.png'
    bokeh.io.reset_output()
    bokeh.io.curdoc().theme = theme
    bokeh.io.export_png(hv_fig, filename = file_save)

    # compute statistics to show strength of relation
    print(value)
    X = np.array(plotting_df2['Fuel (mM)'].astype(int)).reshape(-1, 1)
    Y = np.array([plotting_df2['Integrated MGaptamer (μM-hr)'], plotting_df2['Maximum deGFP (μM)']]).T
    spearman_rho_bootstrap_values(X, Y)
    print('\n')

# %% [code]
# get relevant f30-pepper background subtraction data
f30pepper_bkgd_samples = master_df.loc[(master_df['DNA (nM)']==0) &
                      (master_df['DNA/mRNA batch'] ==1) &
                      (master_df['Extract']=='DAJ_MK_MY') & 
                      (master_df['Fuel (mM)']==0) & 
                      (master_df['Mg (mM)']==0) & 
                      (master_df['Plasmid construct']=='pTet-F30Pepper-mTurquoise2') & 
                      (master_df['mRNA (uM)'] == 0) & 
                      (master_df['Channel'] == 'Pepper')
                    ]
x = f30pepper_bkgd_samples['Time (hr)']
y = f30pepper_bkgd_samples['Measurement']
fn = np.polyfit(x, y, 5)
f30pepper_bkgd_fn = np.poly1d(fn)

# get relevant mturquoise2 background subtraction data
mturquoise2_bkgd_samples = master_df.loc[(master_df['DNA (nM)']==0) &
                      (master_df['DNA/mRNA batch'] ==1) &
                      (master_df['Extract']=='DAJ_MK_MY') & 
                      (master_df['Fuel (mM)']==0) & 
                      (master_df['Mg (mM)']==0) & 
                      (master_df['Plasmid construct']=='pTet-F30Pepper-mTurquoise2') & 
                      (master_df['mRNA (uM)'] == 0) & 
                      (master_df['Channel'] == 'mTurquoise2')
                    ]
x = mturquoise2_bkgd_samples['Time (hr)']
y = mturquoise2_bkgd_samples['Measurement']
fn = np.polyfit(x, y, 5)
mturquoise2_bkgd_fn = np.poly1d(fn)
        
# create dictionary to store functions
bkgd_subt_dict = {'Pepper': f30pepper_bkgd_fn,
                'mTurquoise2': mturquoise2_bkgd_fn,
                }

# %% [code]
# get relevant samples, do background subtraction
temp_df = master_df.drop(index=master_df[master_df['Plasmid construct'] != 'pTet-F30Pepper-mTurquoise2'].index).groupby(['Extract','Fuel', 'DNA (nM)', 'Channel','Plasmid construct'])

# create list to store dataframes
integrated_mgapt_dfs = []
max_degfp_dfs = []

# iterate through groups
for name, group in temp_df:
                
    # get relevant information
    extract = name[0]
    fuel = name[1]
    dna_conc = name[2]
    fluor_mol = name[3]  

    # background subtraction
    bkgd_sub_fn = bkgd_subt_dict[fluor_mol]
    group['Measurement'] = group['Measurement'] - bkgd_sub_fn(group['Time (hr)'])    
    
    # drop unrelated columns, create new df
    group = group.drop(columns=['Well'])
    cols = list(group.columns)
    cols.remove('Replicate')
    cols.remove('Measurement')
    new_group = group.groupby(cols).apply(lambda group2: np.mean(group2['Measurement'])).reset_index().ffill()
    new_group = pd.DataFrame(new_group).rename({0:'Measurement'}, axis=1)
    new_group = group.copy()
        
    # perform an integration
    if fluor_mol == 'Pepper':
        cols2 = list(new_group.columns)
        cols2.remove('Measurement')
        cols2.remove('Time (hr)')
        cols2.remove('Time (sec)')
        timepoints = sorted(list(set(new_group['Time (hr)'])))
        dx = timepoints[2] - timepoints[1]
        new_group = new_group.groupby(cols2).apply(lambda group2: np.trapz(y=group2['Measurement'], dx=dx)).reset_index().ffill()
        new_group = pd.DataFrame(new_group).rename({0:'Measurement'}, axis=1)
    elif fluor_mol == 'mTurquoise2':
        new_group = new_group.drop(columns=['Time (sec)'])
        cols = list(new_group.columns)
        cols.remove('Measurement')
        cols.remove('Time (hr)')
        new_group = new_group.groupby(cols).apply(lambda group2: np.max(group2['Measurement'])).reset_index().ffill()
        new_group = pd.DataFrame(new_group).rename({0:'Measurement'}, axis=1)
        
    # axis labels
    new_group['Fuel (mM)'] = new_group['Fuel (mM)'].astype('str')
    new_group['Mg (mM)'] = new_group['Mg (mM)'].astype('str')
    
    # drop all but relevant columns
    irrelevant_cols = list(new_group.columns)
    for i in ['Extract', 'DNA (nM)', 'Fuel (mM)', 'Mg (mM)', 'Fuel', 'Measurement', 'Replicate']:
        irrelevant_cols.remove(i)
    new_group = new_group.drop(columns=irrelevant_cols)
    
    # rename measurement column, add relevant data to relevant list of dataframes
    if fluor_mol == 'Pepper':
        new_group = pd.DataFrame(new_group).rename({'Measurement':'Integrated F30-Pepper (AFU-hr)'}, axis=1)
        integrated_mgapt_dfs.append(new_group)
    elif fluor_mol == 'mTurquoise2':
        new_group = pd.DataFrame(new_group).rename({'Measurement':'Maximum mTurquoise2 (AFU)'}, axis=1)
        max_degfp_dfs.append(new_group)

# %% [code]
# combine list of dataframes into single dataframe for each channel
combined_integrated_mgapt_df = pd.concat(integrated_mgapt_dfs)
combined_max_degfp_df = pd.concat(max_degfp_dfs)

# combine all data into one dataframe via merge
max_degfp_integrated_mgapt_df = combined_integrated_mgapt_df.merge(combined_max_degfp_df, on=['Fuel', 'DNA (nM)', 'Mg (mM)', 'Fuel (mM)', 'Extract', 'Replicate'])
max_degfp_integrated_mgapt_df.head()

# %% [code]
# plot points colored by Mg
color_legend_title = 'Fuel (mM)'
plotting_df = max_degfp_integrated_mgapt_df.copy()
plotting_df = plotting_df.loc[plotting_df['DNA (nM)']==5]

hv_fig = hv.Points(data=plotting_df,
                    kdims=['Integrated F30-Pepper (AFU-hr)', 'Maximum mTurquoise2 (AFU)'],
                    vdims=['Fuel', 'DNA (nM)', 'Mg (mM)', 'Fuel (mM)'],
                   
).opts(
    frame_height=1000,
    frame_width=1200,
    show_grid=False,
    cmap=list(bokeh.palettes.Viridis[7]),
    color=color_legend_title,
    size=20,
    xformatter='%.0f',
    xlim = (0, 220000),
    ylim = (-200, 11600),
    legend_opts={'title': '3PGA (mM)',
                 'label_text_font_size':'45pt',
                 'title_text_font_size':'50pt',
                 'title_text_font_style': 'normal',
                 'glyph_width':50,
                 'glyph_height':50
                }
)

# render as bokeh figure, format
hv_fig = hv.render(hv_fig)
hv_fig.legend.spacing = 25

# save plot
file_save = '..\\Figures\\Fig4_d.png'
bokeh.io.reset_output()
bokeh.io.curdoc().theme = theme
bokeh.io.export_png(hv_fig, filename = file_save)

# %% [code]
# compute statistics to show strength of relation
X = np.array(plotting_df['Fuel (mM)'].astype(int)).reshape(-1, 1)
Y = np.array([plotting_df['Integrated F30-Pepper (AFU-hr)'], plotting_df['Maximum mTurquoise2 (AFU)']]).T
spearman_rho_bootstrap_values(X, Y)

# %% [code]
def fig4():
    
    # Read in PNG images
    png_paths = ["..\\Figures\\Fig4_a.png", 
                "..\\Figures\\Fig4_b.png",
                "..\\Figures\\Fig4_c.png",
                "..\\Figures\\Fig4_d.png"]
    
    # Create Divs for each subfigure
    divs = []
    for i, path in enumerate(png_paths):
        with open(path, "rb") as f:
            img_data = f.read()
        img_src = f"data:image/png;base64,{b64encode(img_data).decode()}"
        div_content = f'<img src="{img_src}">' # style="max-width:100%; max-height:100%">'
        div = Div(text=div_content)
        divs.append(div)
    
    # create panel of images
    panel = column(row([row([p_label_a, divs[0]]), row(Spacer(width=100)), row([p_label_b, divs[1]])]),
                   Spacer(height=100),
                   row([row([p_label_c, divs[2]]), row(Spacer(width=100)), row([p_label_d, divs[3]])]),
                  )
    
    # save plot
    file_save = '..\\Figures\\Fig4.png'
    bokeh.io.reset_output()
    bokeh.io.curdoc().theme = theme
    bokeh.io.export_png(panel, filename = file_save)

# %% [code]
fig4()

# %% [markdown]
# 

# %% [markdown]
# Figure 6

# %% [code]
# get relevant degfp background subtraction data
# mgapt samples
mgapt_bkgd_samples = master_df.loc[(master_df['DNA (nM)']==0) &
                      (master_df['DNA/mRNA batch'] < 5) &
                      (master_df['Extract']=='DAJ_MK_MY') & 
                      (master_df['Fuel (mM)']==0) & 
                      (master_df['Mg (mM)']==0) & 
                      (master_df['Plasmid construct']=='pOR1OR2-MGapt-deGFP') & 
                      (master_df['mRNA (uM)'] == 0) & 
                      (master_df['Channel'] == 'MGaptamer')
                    ]
x = mgapt_bkgd_samples['Time (hr)']
y = mgapt_bkgd_samples['Measurement']
fn = np.polyfit(x, y, 5)
mgapt_fn = np.poly1d(fn)

# degfp samples
degfp_bkgd_samples = master_df.loc[(master_df['DNA (nM)']==0) &
                      (master_df['DNA/mRNA batch'] < 5) &
                      (master_df['Extract']=='DAJ_MK_MY') & 
                      (master_df['Fuel (mM)']==0) & 
                      (master_df['Mg (mM)']==0) & 
                      (master_df['Plasmid construct']=='pOR1OR2-MGapt-deGFP') & 
                      (master_df['mRNA (uM)'] == 0) & 
                      (master_df['Channel'] == 'deGFP')
                    ]
x = degfp_bkgd_samples['Time (hr)']
y = degfp_bkgd_samples['Measurement']
fn = np.polyfit(x, y, 5)
degfp_fn = np.poly1d(fn)
        
# create dictionary to store functions
bkgd_subt_dict = {
                    'MGaptamer': mgapt_fn,
                    'deGFP': degfp_fn,
                  }

# %% [code]
# get appropriate data
tx_df = master_df.drop(index=master_df[(master_df['Fuel'] !='3PGA') |
                                        (master_df['Extract'] !='DAJ_MK_MY') |
                                        (master_df['DNA (nM)'] != 5) |
                                        (master_df['Plasmid construct'] != 'pOR1OR2-MGapt-deGFP') | 
                                        (master_df['mRNA (uM)'] != 0) | 
                                        (master_df['DNA/mRNA batch'] != 6) |
                                        (master_df['Tetratcycline (ug/mL)'] != 200)
                                        ].index)
tl_df = master_df.drop(index=master_df[(master_df['Fuel'] !='3PGA') |
                                        (master_df['Extract'] !='DAJ_MK_MY') |
                                        (master_df['DNA (nM)'] != 0) |
                                        (master_df['Plasmid construct'] != 'pOR1OR2-MGapt-deGFP') | 
                                        (master_df['mRNA (uM)'] != 0.3) | 
                                        (master_df['DNA/mRNA batch'] >= 5)
                                        ].index)

# %% [code]
# subset only relevant fuel
subset_df1 = master_df.drop(index=master_df[(master_df['Fuel'] !='No fuel') |
                                        (master_df['Extract'] !='DAJ_MK_MY') |
                                        (master_df['DNA (nM)'] != 5) |
                                        (master_df['Plasmid construct'] != 'pOR1OR2-MGapt-deGFP') | 
                                        (master_df['mRNA (uM)'] != 0) | 
                                        (master_df['DNA/mRNA batch'] >= 5)
                                        ].index)
subset_df2 = master_df.drop(index=master_df[(master_df['Fuel'] !='3PGA') |
                                        (master_df['Extract'] !='DAJ_MK_MY') |
                                        (master_df['DNA (nM)'] != 5) |
                                        (master_df['Plasmid construct'] != 'pOR1OR2-MGapt-deGFP') | 
                                        (master_df['mRNA (uM)'] != 0) | 
                                        (master_df['DNA/mRNA batch'] >= 5)
                                        ].index)
txtl_df = pd.concat([subset_df1, subset_df2], axis=0)

# %% [code]
# figure suffixes
suffixes = ['a', 'b', 'c']
titles = ['TX-TL system', 'TX-only system', 'TL-only system']

# iterate through data
for index, df in enumerate([txtl_df, tx_df, tl_df]):

    # get dataframe that contains only non-NTP fuels
    temp_df = df.groupby(['Channel'])

    # iterate through channels
    for name, group in temp_df:

        # if mgapt
        if name[0] == 'MGaptamer':
        
            # define rounding
            value_dimension = hv.Dimension('Measurement', value_format=lambda x: '%.1f' % x)    
            
            # background subtract
            group['Measurement'] = group['Measurement'] - mgapt_fn(group['Time (hr)'])
            
            # drop unrelated columns, create new df
            group = group.drop(columns=['Well'])
            cols = list(group.columns)
            cols.remove('Replicate')
            cols.remove('Measurement')
            new_group = group.groupby(cols).apply(lambda group2: np.mean(group2['Measurement'])).reset_index().ffill()
            new_group = pd.DataFrame(new_group).rename({0:'Measurement'}, axis=1)
            
            # perform an integration
            cols2 = list(new_group.columns)
            cols2.remove('Measurement')
            cols2.remove('Time (hr)')
            cols2.remove('Time (sec)')
            timepoints = sorted(list(set(new_group['Time (hr)'])))
            dx = timepoints[2] - timepoints[1]
            new_group = new_group.groupby(cols2).apply(lambda group2: np.trapz(y=group2['Measurement'], dx=dx)).reset_index().ffill()
            new_group = pd.DataFrame(new_group).rename({0:'Measurement'}, axis=1)
            
            # axis labels
            new_group['Fuel (mM)'] = new_group['Fuel (mM)'].astype('str')
            new_group['Mg (mM)'] = new_group['Mg (mM)'].astype('str')
            new_group['Fuel (mM)'] = pd.Categorical(new_group['Fuel (mM)'],categories=['0', '5', '10', '15', '20', '30', '45'])
            new_group['Mg (mM)'] = pd.Categorical(new_group['Mg (mM)'],categories=['0', '2', '4', '6', '8', '10'])
            new_group = new_group.sort_values(['Fuel (mM)', "Mg (mM)"])

            # plot range max
            this_max = np.max(new_group['Measurement'])
            
            # create holoviews heatmap
            hv_fig = hv.HeatMap(data=new_group,
                                kdims=[('Fuel (mM)', '3PGA (mM)'), ('Mg (mM)', 'Added Mg²⁺ (mM)') ],
                                vdims=hv.Dimension('Measurement', range=(0, this_max))
            ).opts(
                frame_height=750,
                frame_width=750,
                colorbar=True,
                show_grid=False,
                cmap='Viridis',
                title = f'Integrated MG aptamer (μM-hr)',
                # fontsize={'labels': 35, 'xticks': 30, 'yticks': 30, 'ticks':30}
                fontsize={'labels': 50, 'xticks': 45, 'yticks': 45, 'ticks':45, 'title':50}
            )
            hv_fig = hv_fig * hv.Labels(hv_fig, vdims=value_dimension).opts(text_color='white', text_font_size='35px',fontsize={'ticks':20})
            
            # render as bokeh figure, format
            hv_fig1 = hv.render(hv_fig)
            hv_fig1.toolbar_location = None
            hv_fig1.title.align = 'center'

        # elif degfp
        elif name[0] == 'deGFP':    
    
            # background subtract
            group['Measurement'] = group['Measurement'] - degfp_fn(group['Time (hr)'])
            
            # define rounding
            value_dimension = hv.Dimension('Measurement', value_format=lambda x: '%.2f' % x)    
            
            # drop unrelated columns, create new df
            group = group.drop(columns=['Well'])
            cols = list(group.columns)
            cols.remove('Replicate')
            cols.remove('Measurement')
            new_group = group.groupby(cols).apply(lambda group2: np.mean(group2['Measurement'])).reset_index().ffill()
            new_group = pd.DataFrame(new_group).rename({0:'Measurement'}, axis=1)
            
            # axis labels
            new_group['Fuel (mM)'] = new_group['Fuel (mM)'].astype('str')
            new_group['Mg (mM)'] = new_group['Mg (mM)'].astype('str')
            new_group['Fuel (mM)'] = pd.Categorical(new_group['Fuel (mM)'],categories=['0', '5', '10', '15', '20', '30', '45'])
            new_group['Mg (mM)'] = pd.Categorical(new_group['Mg (mM)'],categories=['0', '2', '4', '6', '8', '10'])
            new_group = new_group.sort_values(['Fuel (mM)', 'Mg (mM)'])

            # plot range max
            this_max = np.max(new_group['Measurement'])
            
            # create holoviews heatmap
            hv_fig = hv.HeatMap(data=new_group,
                                kdims=[('Fuel (mM)', '3PGA (mM)'), ('Mg (mM)', 'Added Mg²⁺ (mM)') ],
                                vdims=hv.Dimension('Measurement', range=(0, this_max))
            ).aggregate(function=np.max
            ).opts(
                frame_height=750,
                frame_width=750,
                colorbar=True,
                show_grid=False,
                cmap='Viridis',
                title = f'Maximum deGFP (μM)',
                # fontsize={'labels': 35, 'xticks': 30, 'yticks': 30, 'ticks':30}
                fontsize={'labels': 50, 'xticks': 45, 'yticks': 45, 'ticks':45, 'title':50}
            )
            hv_fig = hv_fig * hv.Labels(hv_fig, vdims=value_dimension).opts(text_color='white', text_font_size='35px',fontsize={'ticks':20})
            # render as bokeh figure, format
            hv_fig2 = hv.render(hv_fig)
            hv_fig2.toolbar_location = None
            hv_fig2.title.align = 'center'
            
    # put together heatmaps, label them
    title = Div(text = f'<p style="color: #444444; font-size: 150px; margin:0;"> {titles[index]} </p>', align='center')
    p = column(title, hv_fig1, Spacer(height=75), hv_fig2)
    p = column(hv_fig1, Spacer(height=75), hv_fig2)

    # save heatmaps
    file_save = f'..\\Figures\\Fig6_{suffixes[index]}.png'
    bokeh.io.reset_output()
    bokeh.io.curdoc().theme = theme
    bokeh.io.export_png(p, filename = file_save)

# %% [code]
def fig6():
    
    # Read in PNG images
    png_paths = ["..\\Figures\\Fig6_a.png", 
                "..\\Figures\\Fig6_b.png",
                "..\\Figures\\Fig6_c.png",
                "..\\Figures\\Fig6_d.png",
                 "..\\Figures\\Fig6_e.png",
                 "..\\Figures\\Fig6_f.png"
                ]
    
    # Create Divs for each subfigure
    divs = []
    for i, path in enumerate(png_paths):
        with open(path, "rb") as f:
            img_data = f.read()
        img_src = f"data:image/png;base64,{b64encode(img_data).decode()}"
        # if i > 2:
        #     div_content = f'<img src="{img_src}" margin:0; style="max-width:50%; max-height:50%">'
        # else:
        div_content = f'<img src="{img_src}">' # style="max-width:100%; max-height:100%">'
        div = Div(text=div_content)
        divs.append(div)
    
    # create panel of images
    title_txtl = Div(text = '<p style="color: #444444; font-size: 100px; margin:0;"> TX-TL system </p>', align='center')
    title_tx = Div(text = '<p style="color: #444444; font-size: 100px; margin:0;"> TX-only system </p>', align='center')
    title_tl = Div(text = '<p style="color: #444444; font-size: 100px; margin:0;"> TL-only system </p>', align='center')

    # put together figure
    panel = column(row([row([p_label_a, Spacer(width=50), column(title_txtl, divs[3])]), 
                        row(Spacer(width=180)), 
                        row([p_label_b, Spacer(width=50), column(title_tx, divs[4])]), 
                        row(Spacer(width=180)), 
                        row([p_label_c, Spacer(width=50), column(title_tl, divs[5])])]),
                   Spacer(height=100),
                   row([row([p_label_d, divs[0]]), row(Spacer(width=100)), row([p_label_e, divs[1]]), row(Spacer(width=100)), row([p_label_f, divs[2]])])
                  )
    
    # save plot
    file_save = '..\\Figures\\Fig6.png'
    bokeh.io.reset_output()
    bokeh.io.curdoc().theme = theme
    bokeh.io.export_png(panel, filename = file_save)

# %% [code]
fig6()

# %% [markdown]
# 

# %% [markdown]
# Figure S1

# %% [code]
def figS1():
    
    # Read in PNG images
    png_paths = ["..\\Figures\\FigS1_a.png", 
                   "..\\Figures\\FigS1_b.png", 
                   "..\\Figures\\FigS1_c.png"]

    # initialize labels as figures
    p_label_a_small = bokeh.plotting.figure(width=75, height=85, toolbar_location=None, tools="")
    p_label_b_small = bokeh.plotting.figure(width=75, height=85, toolbar_location=None, tools="")
    p_label_c_small = bokeh.plotting.figure(width=75, height=85, toolbar_location=None, tools="")    
    
    # create labels for the figures
    label_a = Label(x=0, y=5, x_units='screen', y_units='screen',
                  text_font_size = '75px',
                  text_font_style = 'bold',
                  text_color = 'black',
                  text='A', 
                  background_fill_color='white', background_fill_alpha=1.0)
    label_b = Label(x=0, y=5, x_units='screen', y_units='screen',
                  text_font_size = '75px',
                  text_font_style = 'bold',
                  text_color = 'black',
                  text='B', 
                  background_fill_color='white', background_fill_alpha=1.0)
    label_c = Label(x=0, y=5, x_units='screen', y_units='screen',
                  text_font_size = '75px',
                  text_font_style = 'bold',
                  text_color = 'black',
                  text='C', 
                  background_fill_color='white', background_fill_alpha=1.0)
    # add labels
    p_label_a_small.add_layout(label_a)
    p_label_b_small.add_layout(label_b)
    p_label_c_small.add_layout(label_c)
    
    # Create Divs for each subfigure
    divs = []
    for i, path in enumerate(png_paths):
        with open(path, "rb") as f:
            img_data = f.read()
        img_src = f"data:image/png;base64,{b64encode(img_data).decode()}"
        div_content = f'<img src="{img_src}">' # style="max-width:100%; max-height:100%">'
        div = Div(text=div_content)
        divs.append(div)
    
    # create panel of images
    panel = row([column([row([p_label_a_small, divs[0]]), 
                         Spacer(height=100), 
                         row([p_label_b_small, divs[1]])]),
                 row([p_label_c_small, divs[2]]),
                ])
    
    # save plot
    file_save = '..\\Figures\\FigS1.png'
    bokeh.io.reset_output()
    bokeh.io.curdoc().theme = theme
    bokeh.io.export_png(panel, filename = file_save)

# %% [code]
figS1()

# %% [markdown]
# 

# %% [markdown]
# Figure S14

# %% [code]
def figS14():
    
    # Read in PNG images
    png_paths = ["..\\Figures\\FigS14_a.png", 
                "..\\Figures\\FigS14_b.png",
                "..\\Figures\\FigS14_c.png",
                "..\\Figures\\FigS14_d.png"]
    
    # Create Divs for each subfigure
    divs = []
    for i, path in enumerate(png_paths):
        with open(path, "rb") as f:
            img_data = f.read()
        img_src = f"data:image/png;base64,{b64encode(img_data).decode()}"
        div_content = f'<img src="{img_src}">' # style="max-width:100%; max-height:100%">'
        div = Div(text=div_content)
        divs.append(div)
    
    # create panel of images
    panel = column(row([row([p_label_a, divs[0]]), row(Spacer(width=100)), row([p_label_b, divs[1]])]),
                   Spacer(height=100),
                   row([row([p_label_c, divs[2]]), row(Spacer(width=200)), row([p_label_d, divs[3]])]),
                  )
    
    # save plot
    file_save = '..\\Figures\\FigS14.png'
    bokeh.io.reset_output()
    bokeh.io.curdoc().theme = theme
    bokeh.io.export_png(panel, filename = file_save)

# %% [code]
figS14()

# %% [markdown]
# 

# %% [markdown]
# Figure S15

# %% [code]
def figS15():
    
    # Read in PNG images
    png_paths = ["..\\Figures\\FigS15_a.png", 
                   "..\\Figures\\FigS15_b.png", 
                   "..\\Figures\\FigS15_c.png"]

    # initialize labels as figures
    p_label_a_small = bokeh.plotting.figure(width=75, height=85, toolbar_location=None, tools="")
    p_label_b_small = bokeh.plotting.figure(width=75, height=85, toolbar_location=None, tools="")
    p_label_c_small = bokeh.plotting.figure(width=75, height=85, toolbar_location=None, tools="")    
    
    # create labels for the figures
    label_a = Label(x=0, y=5, x_units='screen', y_units='screen',
                  text_font_size = '75px',
                  text_font_style = 'bold',
                  text_color = 'black',
                  text='A', 
                  background_fill_color='white', background_fill_alpha=1.0)
    label_b = Label(x=0, y=5, x_units='screen', y_units='screen',
                  text_font_size = '75px',
                  text_font_style = 'bold',
                  text_color = 'black',
                  text='B', 
                  background_fill_color='white', background_fill_alpha=1.0)
    label_c = Label(x=0, y=5, x_units='screen', y_units='screen',
                  text_font_size = '75px',
                  text_font_style = 'bold',
                  text_color = 'black',
                  text='C', 
                  background_fill_color='white', background_fill_alpha=1.0)
    # add labels
    p_label_a_small.add_layout(label_a)
    p_label_b_small.add_layout(label_b)
    p_label_c_small.add_layout(label_c)
    
    # Create Divs for each subfigure
    divs = []
    for i, path in enumerate(png_paths):
        with open(path, "rb") as f:
            img_data = f.read()
        img_src = f"data:image/png;base64,{b64encode(img_data).decode()}"
        div_content = f'<img src="{img_src}">' # style="max-width:100%; max-height:100%">'
        div = Div(text=div_content)
        divs.append(div)
    
    # create panel of images
    panel = row([column([row([p_label_a_small, divs[0]]), 
                         Spacer(height=100), 
                         row([p_label_c_small, divs[2]])]),
                 row(Spacer(width=100)),
                 row([p_label_b_small, divs[1]]),
                ])
    
    # save plot
    file_save = '..\\Figures\\FigS15.png'
    bokeh.io.reset_output()
    bokeh.io.curdoc().theme = theme
    bokeh.io.export_png(panel, filename = file_save)

# %% [code]
figS15()
```
