## Supplementary material for "Metabolic perturbations to an *E. coli*-based cell-free system reveal a trade-off between transcription and translation": Raw Data, Tidy Data, DNA GenBank files, Figures, and Jupyter Notebooks for Figure Generation and Analysis: environment.yml

```
name: txtl_tradeoff_env
channels:
  - anaconda
  - conda-forge
  - defaults
dependencies:
  - abseil-cpp=20211102.0=hd77b12b_0
  - anyio=4.2.0=py312haa95532_0
  - aom=3.6.0=hd77b12b_0
  - argon2-cffi=21.3.0=pyhd3eb1b0_0
  - argon2-cffi-bindings=21.2.0=py312h2bbff1b_0
  - arrow-cpp=14.0.2=ha81ea56_1
  - asciitree=0.3.3=py_2
  - asttokens=2.0.5=pyhd3eb1b0_0
  - async-lru=2.0.4=py312haa95532_0
  - attrs=23.1.0=py312haa95532_0
  - aws-c-auth=0.6.19=h2bbff1b_0
  - aws-c-cal=0.5.20=h2bbff1b_0
  - aws-c-common=0.8.5=h2bbff1b_0
  - aws-c-compression=0.2.16=h2bbff1b_0
  - aws-c-event-stream=0.2.15=hd77b12b_0
  - aws-c-http=0.6.25=h2bbff1b_0
  - aws-c-io=0.13.10=h2bbff1b_0
  - aws-c-mqtt=0.7.13=h2bbff1b_0
  - aws-c-s3=0.1.51=h2bbff1b_0
  - aws-c-sdkutils=0.1.6=h2bbff1b_0
  - aws-checksums=0.1.13=h2bbff1b_0
  - aws-crt-cpp=0.18.16=hd77b12b_0
  - aws-sdk-cpp=1.10.55=hd77b12b_0
  - babel=2.11.0=py312haa95532_0
  - beautifulsoup4=4.12.2=py312haa95532_0
  - blas=1.0=mkl
  - bleach=4.1.0=pyhd3eb1b0_0
  - blosc=1.21.3=h6c2663c_0
  - bokeh=3.3.4=py312hfc267ef_0
  - boost-cpp=1.82.0=h59b6b97_2
  - bottleneck=1.3.7=py312he558020_0
  - brotli=1.0.9=h2bbff1b_7
  - brotli-bin=1.0.9=h2bbff1b_7
  - brotli-python=1.0.9=py312hd77b12b_7
  - bzip2=1.0.8=h2bbff1b_5
  - c-ares=1.19.1=h2bbff1b_0
  - ca-certificates=2024.7.2=haa95532_0
  - certifi=2024.7.4=py312haa95532_0
  - cffi=1.16.0=py312h2bbff1b_0
  - cfitsio=3.470=h2bbff1b_7
  - cftime=1.6.2=py312he558020_1
  - charls=2.2.0=h6c2663c_0
  - charset-normalizer=2.0.4=pyhd3eb1b0_0
  - click=8.1.7=py312haa95532_0
  - cloudpickle=2.2.1=py312haa95532_0
  - colorcet=3.1.0=py312haa95532_0
  - comm=0.2.1=py312haa95532_0
  - contourpy=1.2.0=py312h59b6b97_0
  - cramjam=2.7.0=py312h15f02fc_0
  - cycler=0.11.0=pyhd3eb1b0_0
  - cytoolz=0.12.2=py312h2bbff1b_0
  - dask=2023.11.0=py312haa95532_0
  - dask-core=2023.11.0=py312haa95532_0
  - datashader=0.16.0=py312haa95532_0
  - dav1d=1.2.1=h2bbff1b_0
  - debugpy=1.6.7=py312hd77b12b_0
  - decorator=5.1.1=pyhd3eb1b0_0
  - defusedxml=0.7.1=pyhd3eb1b0_0
  - distributed=2023.11.0=py312haa95532_0
  - et_xmlfile=1.1.0=py312haa95532_1
  - exceptiongroup=1.2.0=pyhd8ed1ab_2
  - executing=0.8.3=pyhd3eb1b0_0
  - expat=2.5.0=hd77b12b_0
  - fasteners=0.16.3=pyhd3eb1b0_0
  - fastparquet=2023.8.0=py312he558020_0
  - firefox=123.0=h63175ca_0
  - fonttools=4.25.0=pyhd3eb1b0_0
  - freetype=2.12.1=ha860e81_0
  - fsspec=2023.10.0=py312haa95532_0
  - geckodriver=0.34.0=h611cf2b_0
  - gflags=2.2.2=hd77b12b_1
  - giflib=5.2.1=h8cc25b3_3
  - git=2.44.0=h57928b3_0
  - glog=0.5.0=hd77b12b_1
  - grpc-cpp=1.48.2=hfe90ff0_1
  - h11=0.14.0=pyhd8ed1ab_0
  - h5netcdf=1.2.0=py312haa95532_0
  - h5py=3.9.0=py312h59a1360_0
  - hdf4=4.2.13=h712560f_2
  - hdf5=1.12.1=h51c971a_3
  - heapdict=1.0.1=pyhd3eb1b0_0
  - holoviews=1.18.3=py312haa95532_0
  - hvplot=0.9.2=py312haa95532_0
  - hypothesis=6.82.0=py312haa95532_0
  - icc_rt=2022.1.0=h6049295_2
  - icu=73.1=h6c2663c_0
  - idna=3.4=py312haa95532_0
  - imagecodecs=2023.1.23=py312hd5bf116_1
  - imageio=2.33.1=py312haa95532_0
  - importlib-metadata=7.0.1=py312haa95532_0
  - iniconfig=1.1.1=pyhd3eb1b0_0
  - intel-openmp=2023.1.0=h59b6b97_46320
  - ipykernel=6.28.0=py312haa95532_0
  - ipython=8.20.0=py312haa95532_0
  - ipywidgets=8.1.2=py312haa95532_0
  - jedi=0.18.1=py312haa95532_1
  - jinja2=3.1.3=py312haa95532_0
  - joblib=1.2.0=py312haa95532_0
  - jpeg=9e=h2bbff1b_1
  - json5=0.9.6=pyhd3eb1b0_0
  - jsonschema=4.19.2=py312haa95532_0
  - jsonschema-specifications=2023.7.1=py312haa95532_0
  - jupyter=1.0.0=py312haa95532_9
  - jupyter-lsp=2.2.0=py312haa95532_0
  - jupyter_client=8.6.0=py312haa95532_0
  - jupyter_console=6.6.3=py312haa95532_1
  - jupyter_core=5.5.0=py312haa95532_0
  - jupyter_events=0.8.0=py312haa95532_0
  - jupyter_server=2.10.0=py312haa95532_0
  - jupyter_server_terminals=0.4.4=py312haa95532_1
  - jupyterlab_pygments=0.1.2=py_0
  - jupyterlab_server=2.25.1=py312haa95532_0
  - jupyterlab_widgets=3.0.10=py312haa95532_0
  - kaleido-core=0.2.1=h8ffe710_0
  - kiwisolver=1.4.4=py312hd77b12b_0
  - krb5=1.20.1=h5b6d351_0
  - lazy_loader=0.3=py312haa95532_0
  - lcms2=2.12=h83e58a3_0
  - lerc=3.0=hd77b12b_0
  - libaec=1.0.4=h33f27b4_1
  - libavif=0.11.1=h2bbff1b_0
  - libboost=1.82.0=h3399ecb_2
  - libbrotlicommon=1.0.9=h2bbff1b_7
  - libbrotlidec=1.0.9=h2bbff1b_7
  - libbrotlienc=1.0.9=h2bbff1b_7
  - libclang=14.0.6=default_hb5a9fac_1
  - libclang13=14.0.6=default_h8e68704_1
  - libcurl=8.5.0=h86230a5_0
  - libdeflate=1.17=h2bbff1b_1
  - libevent=2.1.12=h56d1f94_1
  - libffi=3.4.4=hd77b12b_0
  - libiconv=1.16=h2bbff1b_3
  - libnetcdf=4.8.1=h6685c40_4
  - libpng=1.6.39=h8cc25b3_0
  - libpq=12.17=h906ac69_0
  - libprotobuf=3.20.3=h23ce68f_0
  - libsodium=1.0.18=h62dcd97_0
  - libssh2=1.10.0=he2ea4bf_2
  - libthrift=0.15.0=h4364b78_2
  - libtiff=4.5.1=hd77b12b_0
  - libwebp-base=1.3.2=h2bbff1b_0
  - libxml2=2.13.1=h24da03e_2
  - libxslt=1.1.41=h0739af5_0
  - libzip=1.8.0=h289538f_1
  - libzopfli=1.0.3=ha925a31_0
  - linkify-it-py=2.0.0=py312haa95532_0
  - llvmlite=0.42.0=py312hf2fb9eb_0
  - locket=1.0.0=py312haa95532_0
  - lxml=5.2.1=py312h395c83e_1
  - lz4=4.3.2=py312h2bbff1b_0
  - lz4-c=1.9.4=h2bbff1b_0
  - m2w64-gcc-libgfortran=5.3.0=6
  - m2w64-gcc-libs=5.3.0=7
  - m2w64-gcc-libs-core=5.3.0=7
  - m2w64-gmp=6.1.0=2
  - m2w64-libwinpthread-git=5.0.0.4634.697f757=2
  - markdown=3.4.1=py312haa95532_0
  - markdown-it-py=2.2.0=py312haa95532_1
  - markupsafe=2.1.3=py312h2bbff1b_0
  - mathjax=2.7.7=h57928b3_3
  - matplotlib=3.8.0=py312haa95532_0
  - matplotlib-base=3.8.0=py312hc7c4135_0
  - matplotlib-inline=0.1.6=py312haa95532_0
  - mdit-py-plugins=0.3.0=py312haa95532_0
  - mdurl=0.1.0=py312haa95532_0
  - mistune=2.0.4=py312haa95532_0
  - mkl=2023.1.0=h6b88ed4_46358
  - mkl-service=2.4.0=py312h2bbff1b_1
  - mkl_fft=1.3.8=py312h2bbff1b_0
  - mkl_random=1.2.4=py312h59b6b97_0
  - mpmath=1.3.0=py312haa95532_0
  - msgpack-python=1.0.3=py312h59b6b97_0
  - msys2-conda-epoch=20160418=1
  - multipledispatch=0.6.0=py312haa95532_0
  - munkres=1.1.4=py_0
  - mypy=1.8.0=py312h2bbff1b_0
  - mypy_extensions=1.0.0=py312haa95532_0
  - nbclient=0.8.0=py312haa95532_0
  - nbconvert=7.10.0=py312haa95532_0
  - nbformat=5.9.2=py312haa95532_0
  - nest-asyncio=1.6.0=py312haa95532_0
  - netcdf4=1.6.2=py312had6fc2f_0
  - networkx=3.1=py312haa95532_0
  - notebook=7.0.8=py312haa95532_0
  - notebook-shim=0.2.3=py312haa95532_0
  - numba=0.59.0=py312hc7c4135_0
  - numcodecs=0.12.1=py312h3469f8a_0
  - numexpr=2.8.7=py312h96b7d27_0
  - numpy=1.26.4=py312hfd52020_0
  - numpy-base=1.26.4=py312h4dde369_0
  - openjpeg=2.4.0=h4fc8c34_0
  - openpyxl=3.0.10=py312h2bbff1b_0
  - openssl=3.0.14=h827c3e9_0
  - orc=1.7.4=h623e30f_1
  - outcome=1.3.0.post0=pyhd8ed1ab_0
  - overrides=7.4.0=py312haa95532_0
  - packaging=23.1=py312haa95532_0
  - pandas=2.1.4=py312hc7c4135_0
  - pandoc=2.12=haa95532_3
  - pandocfilters=1.5.0=pyhd3eb1b0_0
  - panel=1.3.8=py312haa95532_0
  - param=2.0.2=py312haa95532_0
  - parso=0.8.3=pyhd3eb1b0_0
  - partd=1.4.1=py312haa95532_0
  - pathspec=0.10.3=py312haa95532_0
  - patsy=0.5.3=py312haa95532_0
  - pillow=10.2.0=py312h2bbff1b_0
  - pip=23.3.1=py312haa95532_0
  - platformdirs=3.10.0=py312haa95532_0
  - plotly=5.21.0=pyhd8ed1ab_0
  - pluggy=1.0.0=py312haa95532_1
  - ply=3.11=py312haa95532_1
  - prometheus_client=0.14.1=py312haa95532_0
  - prompt-toolkit=3.0.43=py312haa95532_0
  - prompt_toolkit=3.0.43=hd3eb1b0_0
  - psutil=5.9.0=py312h2bbff1b_0
  - pure_eval=0.2.2=pyhd3eb1b0_0
  - pyarrow=14.0.2=py312had8a6e9_0
  - pycparser=2.21=pyhd3eb1b0_0
  - pyct=0.5.0=py312haa95532_0
  - pygments=2.15.1=py312haa95532_1
  - pyparsing=3.0.9=py312haa95532_0
  - pyqt=5.15.10=py312hd77b12b_0
  - pyqt5-sip=12.13.0=py312h2bbff1b_0
  - pysocks=1.7.1=py312haa95532_0
  - pytest=7.4.0=py312haa95532_0
  - python=3.12.2=h1d929f7_0
  - python-dateutil=2.8.2=pyhd3eb1b0_0
  - python-fastjsonschema=2.16.2=py312haa95532_0
  - python-json-logger=2.0.7=py312haa95532_0
  - python-kaleido=0.2.1=pyhd8ed1ab_0
  - python-lmdb=1.4.1=py312hd77b12b_0
  - python-tzdata=2023.3=pyhd3eb1b0_0
  - pytz=2023.3.post1=py312haa95532_0
  - pyviz_comms=3.0.0=py312haa95532_0
  - pywin32=305=py312h2bbff1b_0
  - pywinpty=2.0.10=py312h5da7b33_0
  - pyyaml=6.0.1=py312h2bbff1b_0
  - pyzmq=25.1.2=py312hd77b12b_0
  - qt-main=5.15.2=h19c9488_10
  - qtconsole=5.5.1=py312haa95532_0
  - qtpy=2.4.1=py312haa95532_0
  - re2=2022.04.01=hd77b12b_0
  - referencing=0.30.2=py312haa95532_0
  - requests=2.31.0=py312haa95532_1
  - rfc3339-validator=0.1.4=py312haa95532_0
  - rfc3986-validator=0.1.1=py312haa95532_0
  - rich=13.3.5=py312haa95532_1
  - rpds-py=0.10.6=py312h062c2fa_0
  - scikit-image=0.22.0=py312h20b63e8_0
  - scikit-learn=1.3.0=py312hc7c4135_2
  - scipy=1.11.3=py312h3d2928d_0
  - seaborn=0.12.2=py312haa95532_0
  - selenium=4.18.1=pyhd8ed1ab_0
  - selenium-manager=4.18.1=h975169c_0
  - send2trash=1.8.2=py312haa95532_0
  - setuptools=68.2.2=py312haa95532_0
  - sip=6.7.12=py312hd77b12b_0
  - six=1.16.0=pyhd3eb1b0_1
  - snappy=1.1.10=h6c2663c_1
  - sniffio=1.3.0=py312haa95532_0
  - sortedcontainers=2.4.0=pyhd3eb1b0_0
  - soupsieve=2.5=py312haa95532_0
  - sqlite=3.41.2=h2bbff1b_0
  - stack_data=0.2.0=pyhd3eb1b0_0
  - statsmodels=0.14.0=py312he558020_0
  - svgwrite=1.4.3=pyhd8ed1ab_0
  - sympy=1.12=py312haa95532_0
  - tbb=2021.8.0=h59b6b97_0
  - tblib=1.7.0=pyhd3eb1b0_0
  - tenacity=8.2.2=py312haa95532_1
  - terminado=0.17.1=py312haa95532_0
  - threadpoolctl=2.2.0=pyh0d69192_0
  - tifffile=2023.4.12=py312haa95532_0
  - tinycss2=1.2.1=py312haa95532_0
  - tk=8.6.12=h2bbff1b_0
  - toolz=0.12.0=py312haa95532_0
  - tornado=6.3.3=py312h2bbff1b_0
  - tqdm=4.65.0=py312hfc267ef_0
  - traitlets=5.7.1=py312haa95532_0
  - trio=0.24.0=py312haa95532_0
  - trio-websocket=0.11.1=pyhd8ed1ab_0
  - typing_extensions=4.9.0=py312haa95532_1
  - tzdata=2024a=h04d1e81_0
  - uc-micro-py=1.0.1=py312haa95532_0
  - ucrt=10.0.22621.0=h57928b3_0
  - ujson=5.4.0=py312hd77b12b_0
  - urllib3=2.1.0=py312haa95532_1
  - utf8proc=2.6.1=h2bbff1b_1
  - vc=14.2=h21ff451_1
  - vc14_runtime=14.38.33130=h82b7239_18
  - vs2015_runtime=14.38.33130=hcb4865c_18
  - wcwidth=0.2.5=pyhd3eb1b0_0
  - webencodings=0.5.1=py312haa95532_2
  - websocket-client=0.58.0=py312haa95532_4
  - wheel=0.41.2=py312haa95532_0
  - widgetsnbextension=4.0.10=py312haa95532_0
  - win_inet_pton=1.1.0=py312haa95532_0
  - winpty=0.4.3=4
  - wsproto=1.2.0=pyhd8ed1ab_0
  - xarray=2024.2.0=pyhd8ed1ab_0
  - xlrd=2.0.1=pyhd3eb1b0_1
  - xyzservices=2022.9.0=py312haa95532_1
  - xz=5.4.6=h8cc25b3_0
  - yaml=0.2.5=he774522_0
  - zarr=2.13.3=py312haa95532_0
  - zeromq=4.3.5=hd77b12b_0
  - zfp=1.0.0=hd77b12b_0
  - zict=3.0.0=py312haa95532_0
  - zipp=3.17.0=py312haa95532_0
  - zlib=1.2.13=h8cc25b3_0
  - zstd=1.5.5=hd43e919_0
  - pip:
    - altair==5.2.0
    - appdirs==1.4.4
    - argparse==1.4.0
    - arrow==1.3.0
    - arviz==0.17.0
    - awscli==1.32.58
    - bebi103==0.1.20
    - biopython==1.83
    - black==21.12b0
    - blackcellmagic==0.0.3
    - botocore==1.34.58
    - cairocffi==1.7.1
    - cairosvg==2.7.1
    - chardet==5.2.0
    - cmdstanpy==1.2.1
    - colorama==0.4.4
    - commonmark==0.9.1
    - cssselect2==0.7.0
    - dill==0.3.8
    - dnaplotlib==1.0
    - docutils==0.16
    - fqdn==1.5.1
    - greenlet==3.0.3
    - iqplot==0.3.5
    - isoduration==20.11.0
    - jmespath==1.0.1
    - jsonpointer==2.4
    - jupyter-bokeh==4.0.0
    - jupyter-cache==1.0.0
    - jupyterlab==4.0.13
    - jupyterlab-spellchecker==0.8.4
    - jupytext==1.16.1
    - multiprocess==0.70.16
    - polars==0.20.14
    - prettytable==3.10.0
    - pyasn1==0.5.1
    - pydna==5.2.0
    - pyfiglet==1.0.2
    - pyperclip==1.8.2
    - reportlab==4.2.2
    - rsa==4.7.2
    - s3transfer==0.10.0
    - sqlalchemy==2.0.28
    - stanio==0.3.0
    - svg42pdf==0.2.2
    - tabulate==0.9.0
    - toml==0.10.2
    - tomli==1.2.3
    - types-python-dateutil==2.8.19.20240106
    - typing-extensions==4.10.0
    - uri-template==1.3.0
    - vega-datasets==0.9.0
    - watermark==2.4.3
    - webcolors==1.13
    - xarray-datatree==0.0.14
    - xarray-einstats==0.7.0
prefix: C:\Users\manis\anaconda3\envs\txtl_tradeoff_env\txtl_tradeoff_env
```
