## Supplementary material for "Metabolic perturbations to an *E. coli*-based cell-free system reveal a trade-off between transcription and translation": Raw Data, Tidy Data, DNA GenBank files, Figures, and Jupyter Notebooks for Figure Generation and Analysis: Tidy_and_Combine_Data.ipynb

```
# %% [markdown]
# Tidying data from titration experiments

# %% [code]
# data analysis and computing packages
import numpy as np
import pandas as pd

# echo tools
import murraylab_tools.biotek as mt_biotek
import os

# %% [markdown]
# Create tidy data CSVs for each of the experiments

# %% [code]
# list of all relevant experiments
expt_lst = [
            '20220513_TXTL_WP_ZJ_Titration_of_3PGA_MGapt_deGFP',
            '20220514_TXTL_WP_ZJ_Titration_of_Maltose_MGapt_deGFP',
            '20220515_TXTL_WP_ZJ_Titration_of_Succinate_MGapt_deGFP',
            '20220517_TXTL_WP_ZJ_Titration_of_Pyruvate_MGapt_deGFP',
            '20220519_TXTL_WP_ZJ_Titration_of_NTPs_ATP_GTP_and_Fuel_Neg_Ctrls_MGapt_deGFP',
            '20220714_TXTL_WP_ZJ_Titration_of_3PGA_MGapt_deGFP_5nM_DNA',
            '20220716_TXTL_WP_ZJ_Titration_of_Maltose_MGapt_deGFP_5nM_DNA',
            '20220717_TXTL_WP_ZJ_Titration_of_Succinate_MGapt_deGFP_5nM_DNA',
            '20220718_TXTL_WP_ZJ_Titration_of_Pyruvate_MGapt_deGFP_5nM_DNA',
            '20220720_TXTL_WP_ZJ_Titration_of_NTPs_ATP_GTP_and_Fuel_Neg_Ctrls_MGapt_deGFP_5nM_DNA',
            '20220801_TXTL_WP_ZJ_Titration_of_3PGA_MGapt_deGFP_5nM_DNA_Dialyzed_Extract',
            '20220803_TXTL_WP_ZJ_Titration_of_Maltose_MGapt_deGFP_5nM_DNA_Dialyzed_Extract',
            '20220804_TXTL_WP_ZJ_Titration_of_Succinate_MGapt_deGFP_5nM_DNA_Dialyzed_Extract',
            '20220805_TXTL_WP_ZJ_Titration_of_Pyruvate_MGapt_deGFP_5nM_DNA_Dialyzed_Extract',
            '20220809_TXTL_WP_ZJ_Titration_of_NTPs_ATP_GTP_and_Fuel_Neg_Ctrls_MGapt_deGFP_5nM_DNA_Dialyzed_Extract',
            '20220816_TXTL_WP_ZJ_Validation_Data',
            '20230224_TL_only_3PGA_5nM_DNA_Dialyzed_Extract',
           ]
file_loc = "..\\Raw Data\\"

# %% [code]
# load calibration data
calibration_dict = mt_biotek.calibration_data(filename = 'calibration.csv')
calibration_dict

# %% [code]
# iterate through experiments, make each one tidy :)
for this_expt in expt_lst:
    # specify input and output folders
    inputfolder = os.path.join(file_loc, this_expt+'\\')
    outputfolder = os.path.join(file_loc, this_expt+'\\')
    # specify file names
    data_name = this_expt
    data_filename = os.path.join(inputfolder, data_name+"_Results.csv")
    tidy_filename = os.path.join(outputfolder,data_name+"_Results_tidy.csv")
    # specify supplementary file containing metadata on what conditions pertain to each well
    supplementary_filename = os.path.join(inputfolder, data_name+'_supplementary_info.csv')
    # provide calibration data if available
    print(this_expt)
    mt_biotek.tidy_biotek_data(data_filename, 
                               supplementary_filename,
                               volume = 10, 
                               calibration_dict = calibration_dict,
                               convert_to_uM = True,
                               override_plate_reader_id='b1',
                               output_filename = tidy_filename)

# %% [code]
# list of all relevant experiments
expt_lst2 = [
            '20231011_TXTL_Titration_of_3PGA_MGapt_deGFP_2.5nM_DNA_Dialyzed_Extract',
            '20231016_TXTL_Titration_of_3PGA_MGapt_deGFP_7.5nM_DNA_Dialyzed_Extract',
            '20231017_TXTL_Titration_of_3PGA_MGapt_deGFP_10nM_DNA_Dialyzed_Extract',
            '20231018_TXTL_Titration_of_3PGA_MGapt_deGFP_5nM_DNA_Dialyzed_Extract_0.41',
            '20231019_TXTL_Titration_of_3PGA_MGapt_deGFP_5nM_DNA_Dialyzed_Extract_0.25',
            '20231020_TXTL_Titration_of_3PGA_MGapt_deGFP_5nM_DNA_Dialyzed_Extract',
            '20240223_TXTL_Titration_of_3PGA_MGapt_deGFP_1nM_DNA_Dialyzed_Extract',
            '20231122_TXTL_Titration_of_3PGA_Pepperapt_mTurq2_5nM_DNA_Dialyzed_Extract',
            '20231212_TXTL_Titration_of_3PGA_MGapt_deGFP_5nM_DNA_Dialyzed_Extract',
            '20240214_TXTL_Titration_of_3PGA_pMK19_5nM_DNA_Dialyzed_Extract',
            '20240216_TXTL_Titration_of_3PGA_pMK18_5nM_DNA_Dialyzed_Extract',
            '20240426_TX_only_w_Tetracycline_3PGA_5nM_DNA_Dialyzed_Extract',
            '20240726_TXTL_Titration_of_3PGA_pZJQ35_5nM_DNA_Dialyzed_Extract'
           ]
file_loc = "../Raw Data/"

# %% [code]
# load calibration data
calibration_dict = mt_biotek.calibration_data(filename = 'calibration2.csv')
calibration_dict

# %% [code]
# iterate through experiments, make each one tidy :)
for this_expt in expt_lst2:
    # specify input and output folders
    inputfolder = os.path.join(file_loc, this_expt)
    outputfolder = os.path.join(file_loc, this_expt)
    # specify file names
    data_name = this_expt
    data_filename = os.path.join(inputfolder, data_name+"_Results.csv")
    tidy_filename = os.path.join(outputfolder,data_name+"_Results_tidy.csv")
    # specify supplementary file containing metadata on what conditions pertain to each well
    supplementary_filename = os.path.join(inputfolder, data_name+'_supplementary_info.csv')
    # provide calibration data if available
    print(this_expt)
    mt_biotek.tidy_biotek_data(data_filename, 
                               supplementary_filename,
                               volume = 10, 
                               calibration_dict = calibration_dict,
                               convert_to_uM = True,
                               override_plate_reader_id='b1',
                               output_filename = tidy_filename)

# %% [code]
# list of all relevant experiments
expt_lst3 = [
            '20230501_TX_only_w_Tetracycline_3PGA_5nM_DNA_Dialyzed_Extract'
           ]
file_loc = "../Raw Data/"

# %% [code]
# load calibration data
calibration_dict = mt_biotek.calibration_data(filename = 'calibration3.csv')
calibration_dict

# %% [code]
# iterate through experiments, make each one tidy :)
for this_expt in expt_lst3:
    # specify input and output folders
    inputfolder = os.path.join(file_loc, this_expt)
    outputfolder = os.path.join(file_loc, this_expt)
    # specify file names
    data_name = this_expt
    data_filename = os.path.join(inputfolder, data_name+"_Results.csv")
    tidy_filename = os.path.join(outputfolder,data_name+"_Results_tidy.csv")
    # specify supplementary file containing metadata on what conditions pertain to each well
    supplementary_filename = os.path.join(inputfolder, data_name+'_supplementary_info.csv')
    # provide calibration data if available
    print(this_expt)
    mt_biotek.tidy_biotek_data(data_filename, 
                               supplementary_filename,
                               volume = 10, 
                               calibration_dict = calibration_dict,
                               convert_to_uM = True,
                               override_plate_reader_id='b1',
                               output_filename = tidy_filename)

# %% [markdown]
#  

# %% [markdown]
# Create master CSV with all tidy data appended to it

# %% [code]
# iterate through experiments
for index, this_expt in enumerate(expt_lst+expt_lst2+expt_lst3):
    # extract tidy data csv for each
    tidy_csv_loc = os.path.join(file_loc, this_expt,this_expt+"_Results_tidy.csv")
    temp_df = pd.read_csv(tidy_csv_loc)
    # if first data set, create new dataframe from it
    if index==0:
        master_df = temp_df
    # else, append data to existing dataframe
    else:
        master_df = pd.concat([master_df, temp_df])

# reset dataframe index
master_df.reset_index(inplace=True, drop=True)

# %% [code]
# check to make sure operation worked okay
master_df.tail()

# %% [code]
master_df.to_csv('../Tidy Data/tidy_data_calibrated_full.csv', index=False)
```
