## Supplementary material for "Metabolic perturbations to an *E. coli*-based cell-free system reveal a trade-off between transcription and translation": Raw Data, Tidy Data, DNA GenBank files, Figures, and Jupyter Notebooks for Figure Generation and Analysis: Fig0.pptx

### Slide 1
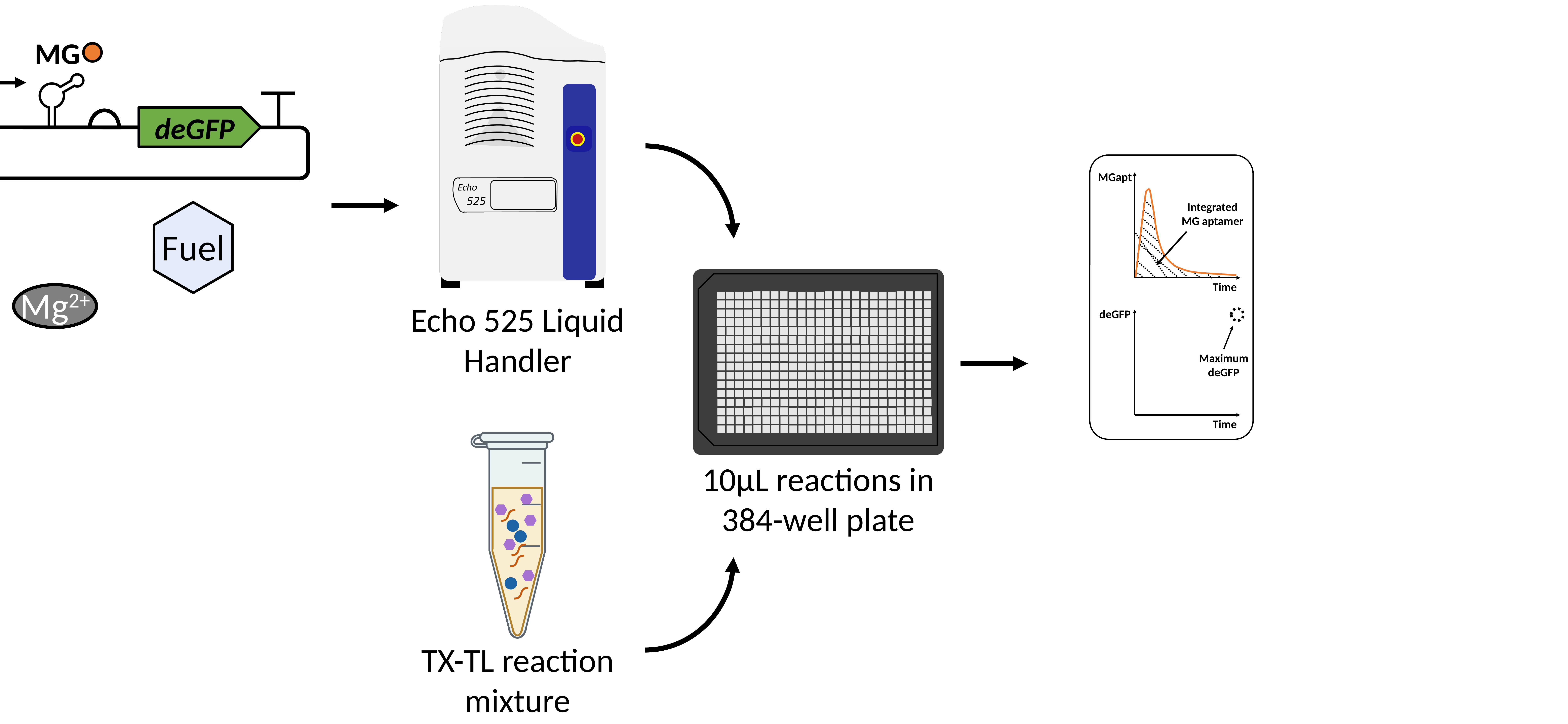

MG
deGFP
MGapt
Integrated MG aptamer
*
Fuel
Time
Mg^2+^
Echo 525 Liquid Handler
deGFP
Maximum deGFP
Time
TX-TL reaction mixture
10µL reactions in 384-well plate

### Slide 2
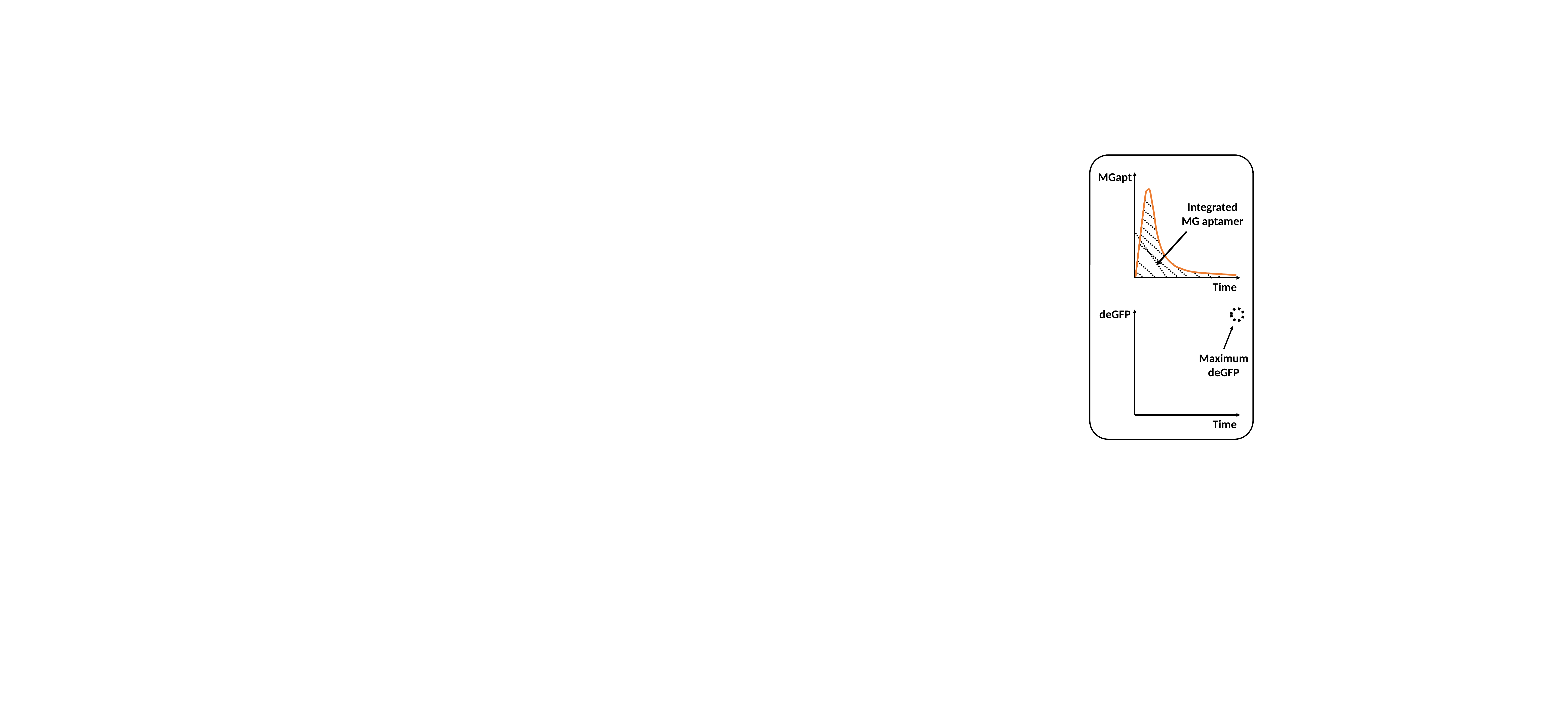

MGapt
Integrated MG aptamer
Time
deGFP
Maximum deGFP
Time

### Slide 3
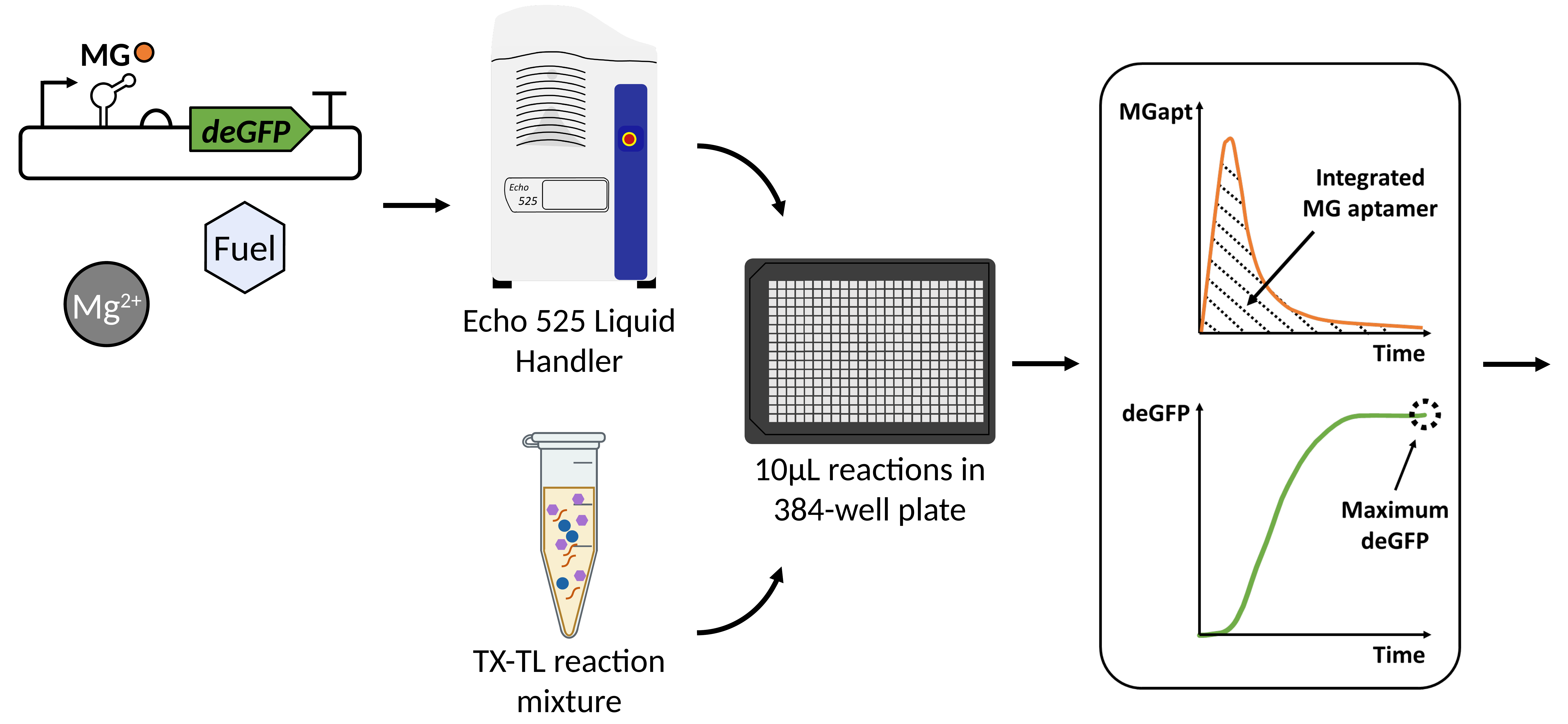

MG
deGFP
*
Fuel
Mg^2+^
Echo 525 Liquid Handler
TX-TL reaction mixture
10µL reactions in 384-well plate

### Slide 4
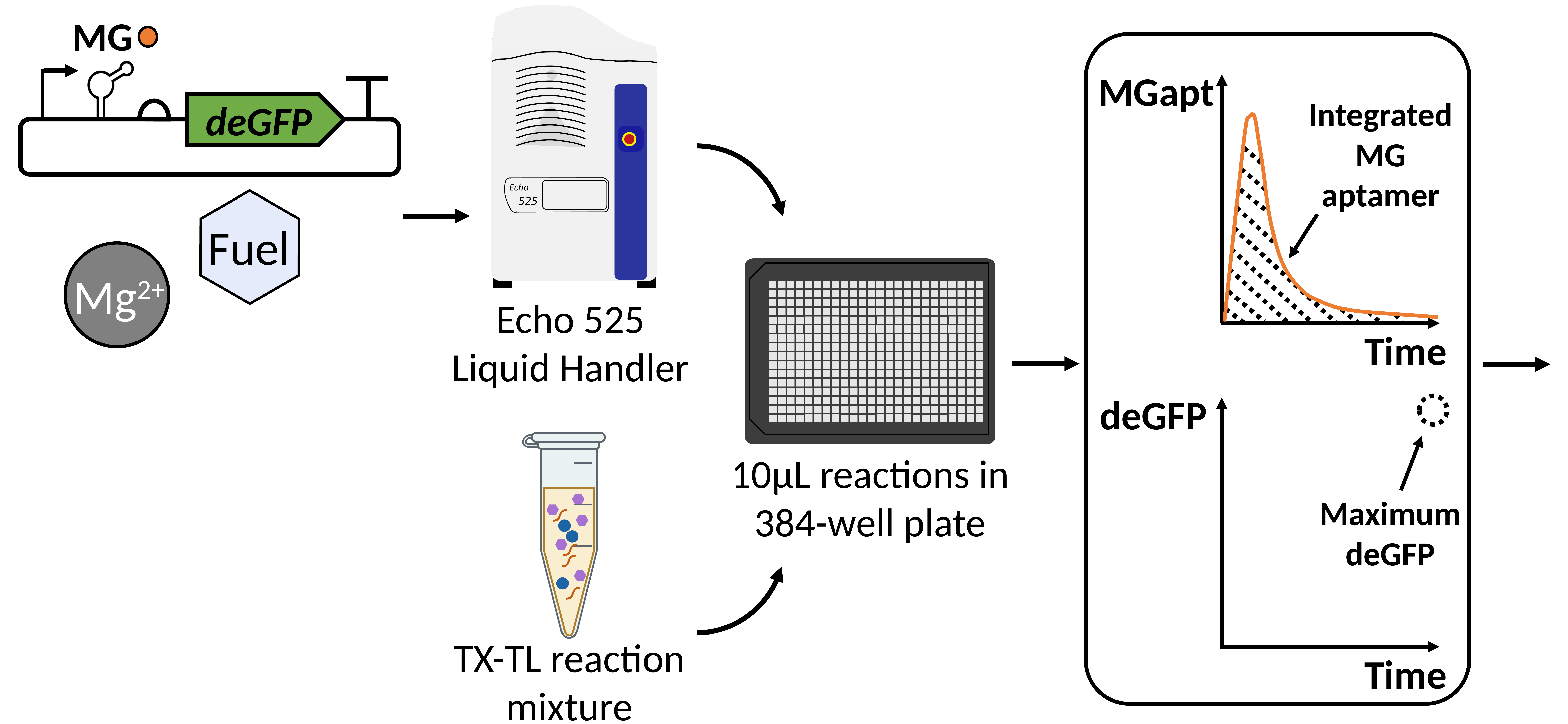

MG
deGFP
MGapt
Integrated MG aptamer
*
Fuel
Mg^2+^
Echo 525 Liquid Handler
Time
deGFP
TX-TL reaction mixture
10µL reactions in 384-well plate
Maximum deGFP
Time

### Slide 5
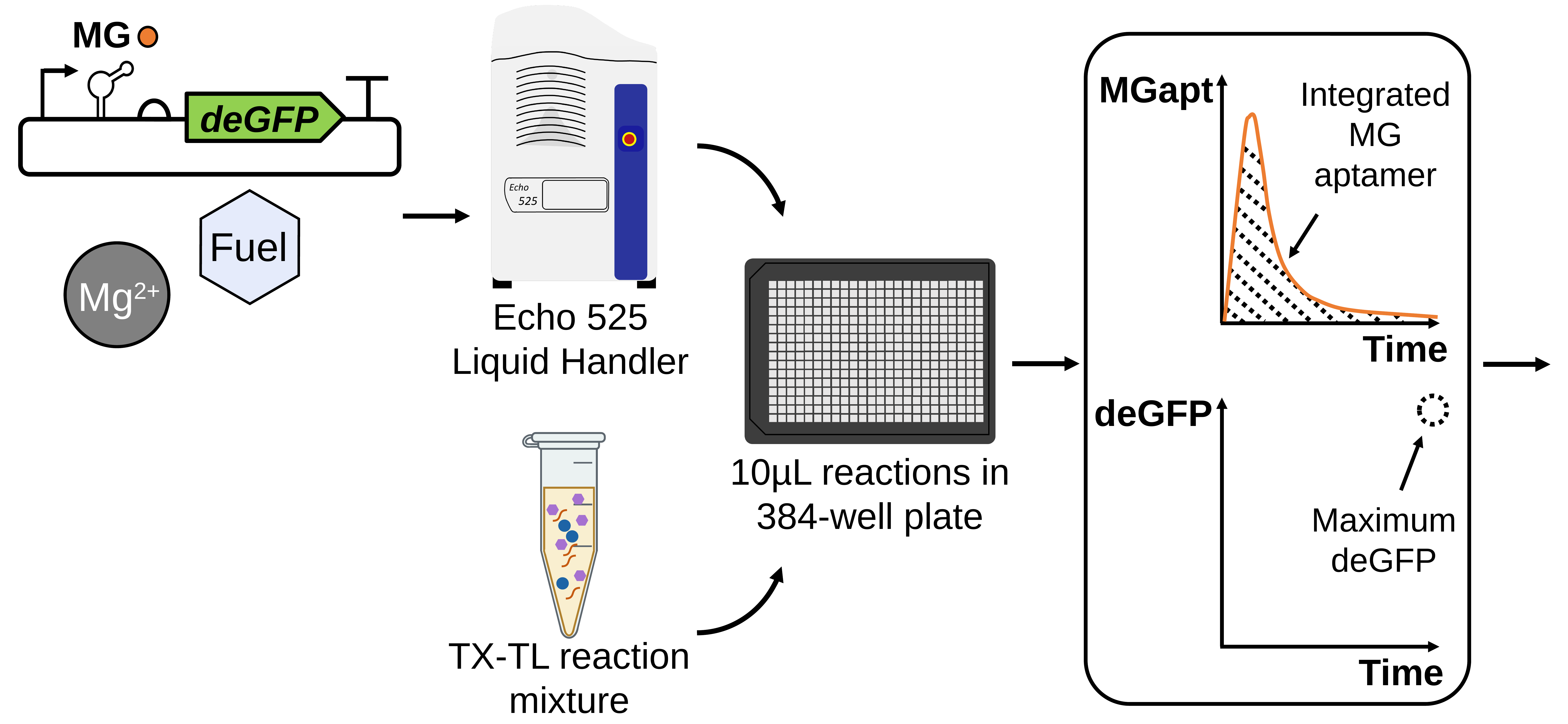

MG
deGFP
MGapt
Integrated MG aptamer
*
Fuel
Mg^2+^
Echo 525 Liquid Handler
Time
deGFP
TX-TL reaction mixture
10µL reactions in 384-well plate
Maximum deGFP
Time
