## Supplementary material for "Metabolic perturbations to an *E. coli*-based cell-free system reveal a trade-off between transcription and translation": Raw Data, Tidy Data, DNA GenBank files, Figures, and Jupyter Notebooks for Figure Generation and Analysis: Fig1_b.pptx

### Slide 1
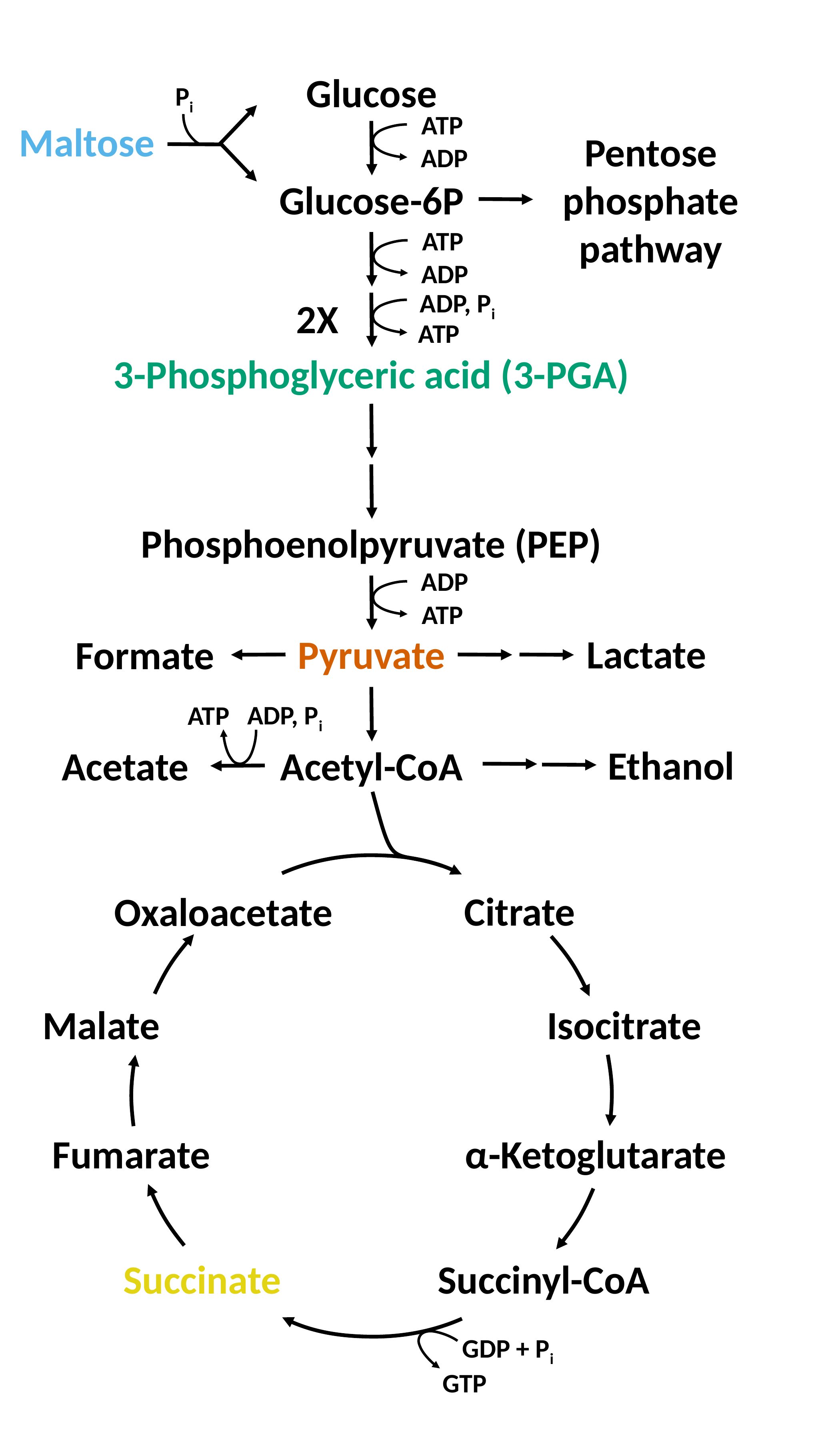

Glucose
P_i_
ATP
Maltose
Pentose phosphate pathway
ADP
Glucose-6P
ATP
ADP
ADP, P_i_
2X
ATP
3-Phosphoglyceric acid (3-PGA)
Phosphoenolpyruvate (PEP)
ADP
ATP
Lactate
Pyruvate
Formate
ADP, P_i_
ATP
Ethanol
Acetyl-CoA
Acetate
Citrate
Oxaloacetate
Malate
Isocitrate
Fumarate
α-Ketoglutarate
Succinate
Succinyl-CoA
GDP + P_i_
GTP
