## Supplementary material for "Metabolic perturbations to an *E. coli*-based cell-free system reveal a trade-off between transcription and translation": Raw Data, Tidy Data, DNA GenBank files, Figures, and Jupyter Notebooks for Figure Generation and Analysis: Fig6_def.pptx

### Slide 1
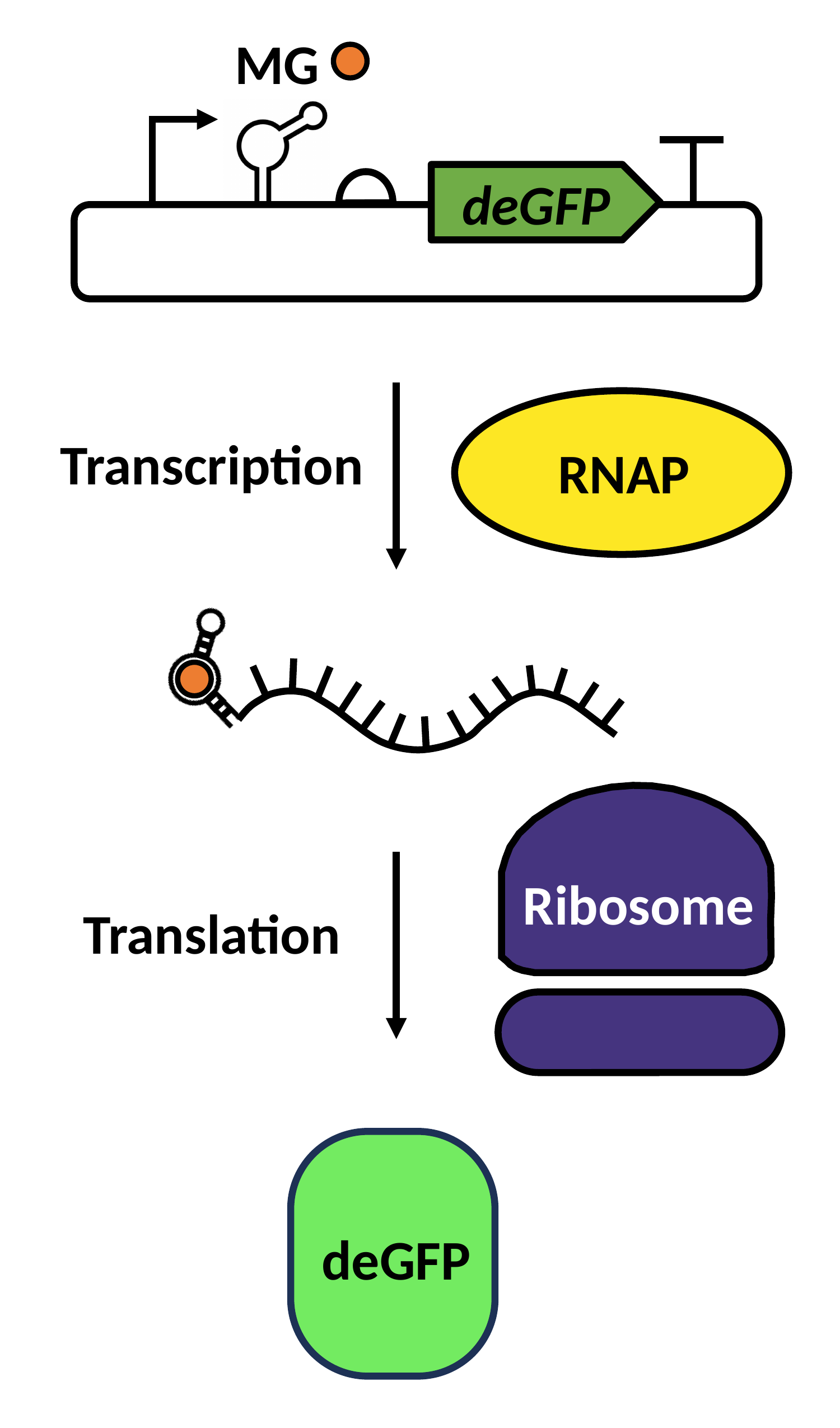

MG
deGFP
Transcription
RNAP
Ribosome
Ribosome
Translation
deGFP

### Slide 2
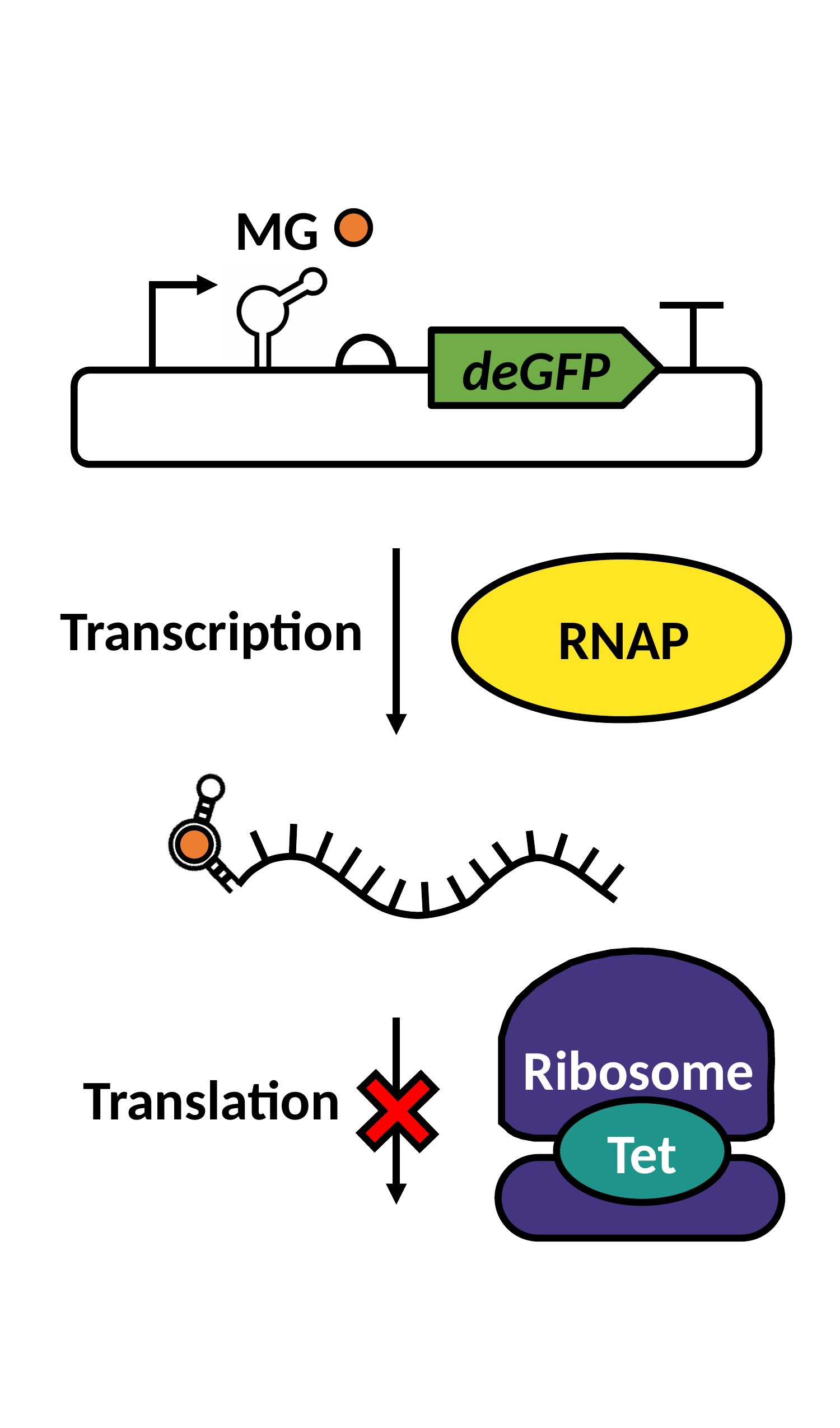

MG
deGFP
Transcription
RNAP
Ribosome
Ribosome
Translation
Tet

### Slide 3
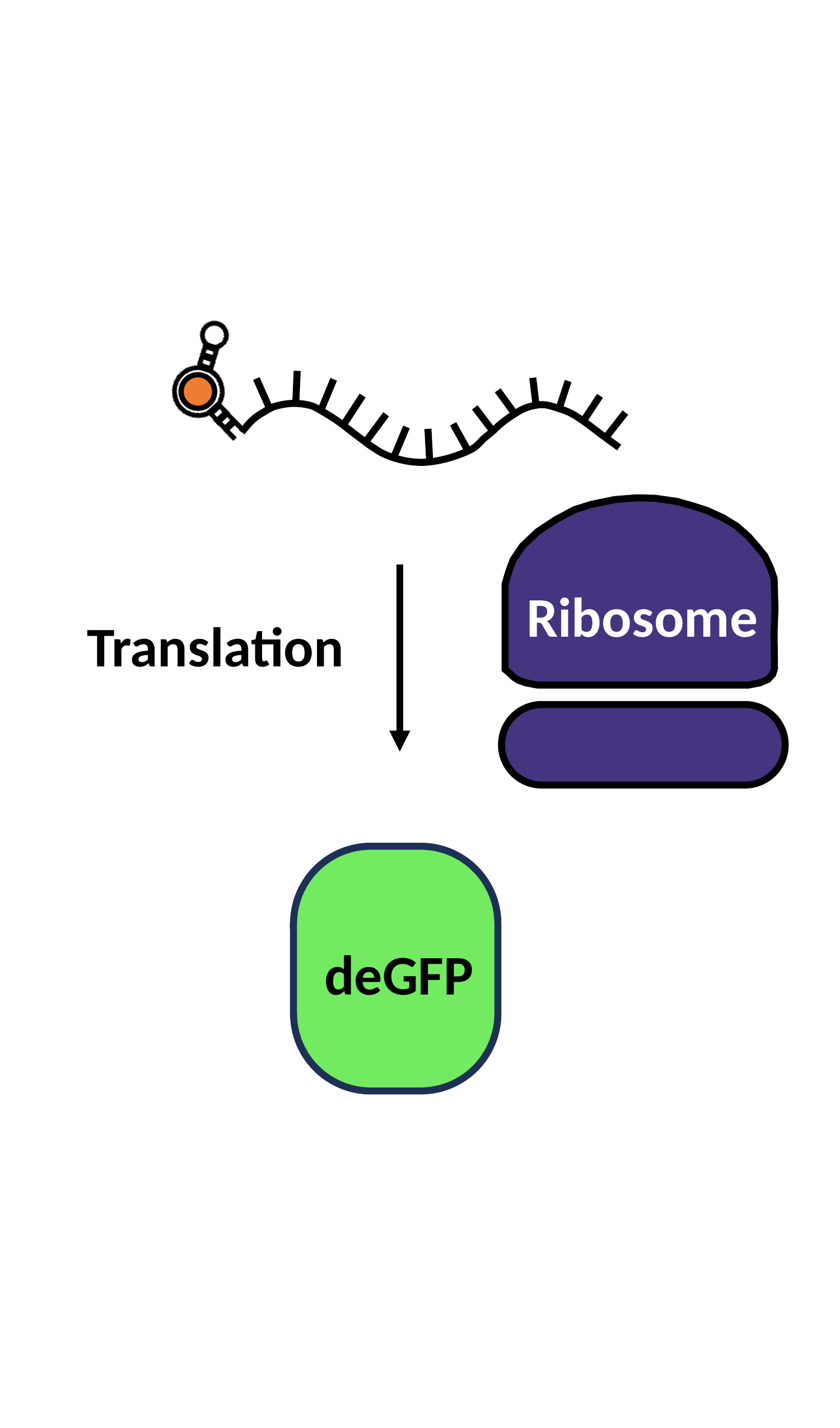

Ribosome
Ribosome
Translation
deGFP
deGFP
