## Supplementary material for "Metabolic perturbations to an *E. coli*-based cell-free system reveal a trade-off between transcription and translation": Raw Data, Tidy Data, DNA GenBank files, Figures, and Jupyter Notebooks for Figure Generation and Analysis: Fig7_a.pptx

### Slide 1
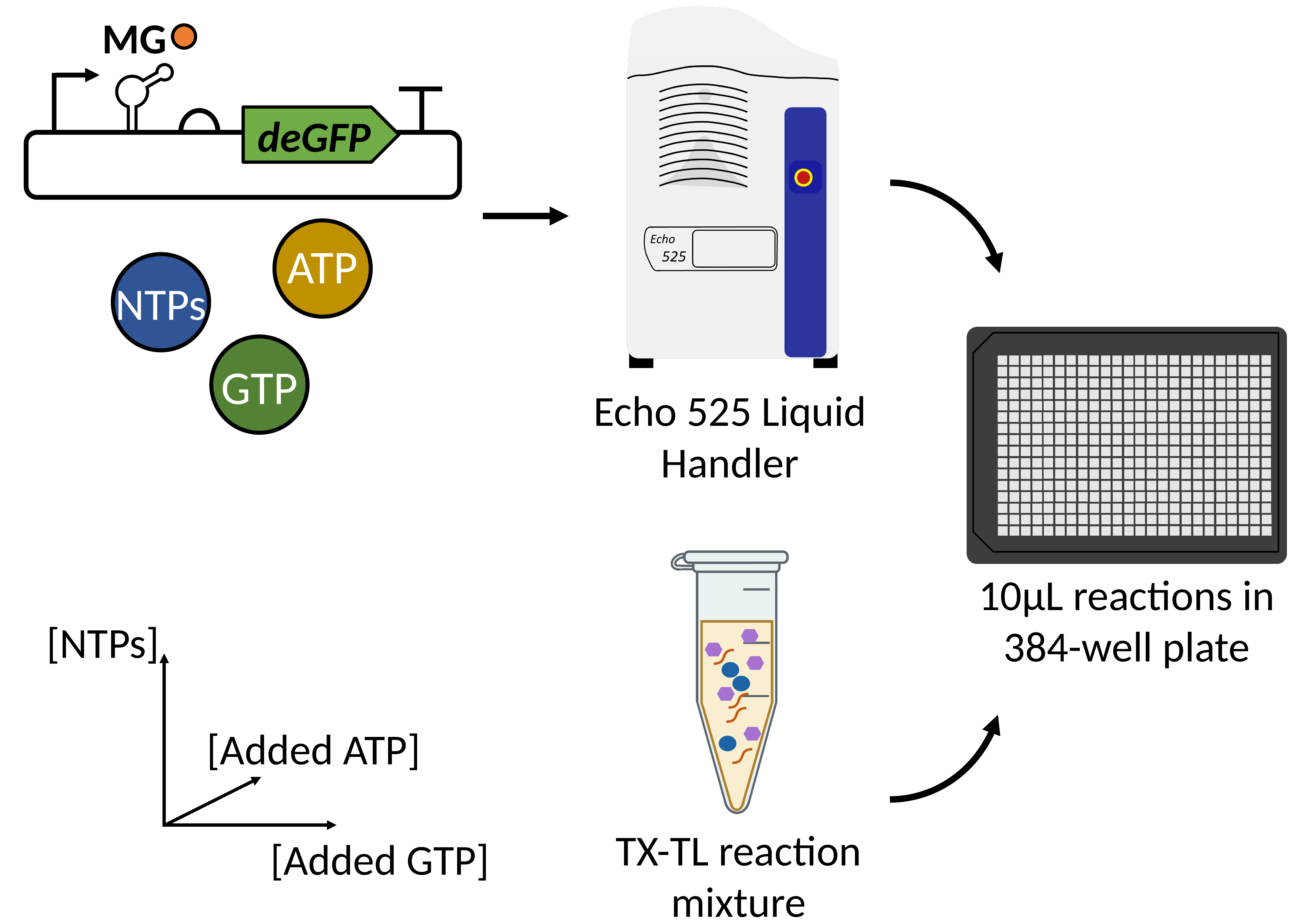

MG
deGFP
ATP
NTPs
GTP
Echo 525 Liquid Handler
TX-TL reaction mixture
10µL reactions in 384-well plate
[NTPs]
[Added ATP]
[Added GTP]
