## Supplementary material for "Metabolic perturbations to an *E. coli*-based cell-free system reveal a trade-off between transcription and translation": Raw Data, Tidy Data, DNA GenBank files, Figures, and Jupyter Notebooks for Figure Generation and Analysis: readme.txt

Description: In collaboration with William Poole, I am doing experiments to evaluate the performance of cell etxract using
varyng amounts of key components. Specifically, I am testing how extract behaves at 25%, 50%, 150%, and 200% of the levels
of NTPs, ATP, Mg, and 3-PGA, and 1-5mM Pi (inorganic phosphate). The data will further be used for parameter inference for 
William's minimal metabolomics model. We use a Jupyter notebook to compute the final concentrations of these components in 
the TXTL reaction mixture. Next, we create a minimal energy solution without these components. We also create an NTP 
solution without ATP, an ATP solution, and a buffer for the extract (to test the quality of MiniPrepped DNA). 
	In this experiment, we re-perform the salt calibrations as before, albeit using a minimal energy solution and 
adding other components back in, rather than using a pre-mixed energy solution.
	We repeat the 2022.02.23 calibration albeit using the correct concentrations of K and only the new energy solution.


Minimal energy solution (5.498mL/27.49):

	Component		Protocol Amount		Amount for 200uL (/27.49)
	---------		---------------		-------------------------
	HEPES			3.6mL				130.96uL
	tRNA			576uL				20.96uL
	CoA			576uL				20.96uL
	cAMP			170uL				6.18uL
	Folinic acid		288uL				10.48uL
	Spermidine		144uL				5.24uL
	H2O			144uL				5.24uL


NTPs w/o ATP:
To prepare the individual NTP solutions,
	- 760mM GTP: Mix 500mg GTP and 910uL H2O
	- 813mM UTP: Mix 300mg UTP and 480uL H2O
	- 621mM CTP: Mix 300mg CTP and 708uL H2O

	Component		Protocol Amount (uL)		Final Concentration (mM)
	---------		--------------------		-------------------------
	GTP				52.8				221
	UTP				29.6				133
	CTP				38.8				133
	15% KOH				60				N/A


NTP solution:

	Component		Protocol Amount 		Final Concentration (mM)
	---------		--------------- 		-------------------------
	ATP			     145mg				156
	GTP			     141mg				156
	UTP			     87.7mg				94
	CTP			     84.3mg				94
	15% KOH			    374.8uL				N/A
	H2O			to 1592.7uL				N/A


ATP solution:
	- 540mM ATP: Mix 471mg ATP and 235uL H2O


Buffer for AR/ZJ 2021 Extract:

	Component		Protocol Amount (uL)
	---------		--------------------
	Mg-glut				9.44
	K-glut				25.12
	AA				94.18
	Energy sol.			26.92
	H2O				1.36


Experiment Protocol:
	1. Create a minimal master mix (for each minimal energy solution) that will be added to all samples. 
	The composition is computed from the same Jupyter notebook used to create the Echo picklist.
	   (42 samples * 6.70uL/sample * 1.1 excess = 281.4uL)
		Component			Protocol Amount (uL)
		---------			--------------------
		Extract				138.6 (6 tubes)
		PEG					21.00
		Minimal energy sol.			16.08
		Amino acids				104.97
		NAD					0.79
	2. We use the Echo to load whatever is not in the minimal master mix into our sample wells, as defined by
	   our Echo picklist. We make sure to leave one well space between wells with different samples.
		Source Well		Component			Volume (uL)
		-----------	-------------------------		-----------
		    B01		Mg-glutamate, 400mM			28
		    B03		K-glutamate, 1200mM			62
		    B05		3-PGA, 467mM				47
		    B07		NTPs, 125mM				38
		    B09		DNA (71 ng/uL)				34
		    B11			H2O				65
	3. Load 6.70uL of minimal master mix into each sample.
	4. Perform the following BioTek protocol.
		  Start Kinetic	Runtime 16:00:00 (HH:MM:SS), Interval 0:05:00, 193 Reads							
			- Linear shake: 0:05 (MM:SS), frequency: 360 cpm (6 mm)							
    			- Read deGFP														
			  Filter Set 1							
	    			Excitation: 485,  Emission: 515							
	    			Optics: Bottom,  Gain: 61							
			  Filter Set 2							
	    			Excitation: 485,  Emission: 515							
	    			Optics: Bottom,  Gain: 100							
			  Light Source: Xenon Flash,  Lamp Energy: High							
			  Read Speed: Normal,  Delay: 100 msec,  Measurements/Data Point: 10							
			  Read Height: 7 mm							


Results: The extract performs best at around 6mM Mg and 140mM K. We can use this an a new optimal for our titration experiments.
Interestingly, performance is poor compared to the previous day's experiment, but this could be due to shitty DNA.
