## Supplementary material for "Metabolic perturbations to an *E. coli*-based cell-free system reveal a trade-off between transcription and translation": Raw Data, Tidy Data, DNA GenBank files, Figures, and Jupyter Notebooks for Figure Generation and Analysis: readme.txt

Description: In collaboration with William Poole and Zoila Jurado, I am doing experiments to probe cell extract metabolism.
In this experiment and subsequent experiments, we vary the amount of fuel we are using at the following levels: 5mM, 10mM,
15mM, 20mM, 30mM, and 45mM. We are effectively performing a panel of fuel vs Mg to understand the complex interplay between
fuel and magnesium. We use a Jupyter notebook to compute the final concentrations of these components in the TXTL 
reaction mixture. 
	We create the following solutions for our experiments: a minimal energy solution without 3-PGA and NTPs; ATP 
solution, GTP solution, NTP solution, 3-PGA solution (from Miki), pyruvate solution, succinate solution, and maltose
solution. We also create a malachite green solution to measure mRNA dynamics.
	In this experiment, we perform a 3-PGA vs Magnesium panel.


Minimal energy solution (5.774mL/7.33):

	Component		Protocol Amount		Amount for 788uL (/7.33)
	---------		---------------		-------------------------
	HEPES			3.6mL				491.13uL
	tRNA			576uL				78.58uL
	CoA			576uL				78.58uL
	cAMP			170uL				23.19uL
	Folinic acid		288uL				39.29uL
	Spermidine		144uL				19.65uL
	NAD+			276uL				37.65uL
	H2O			144uL				19.65uL


GTP (760mM):

	Component		Protocol Amount		Actual amount (/4.237)
	---------		---------------		-------------------------
	GTP			   500mg			118mg
	H2O			   1245uL			294uL


3-PGA (1400mM):

	Component		Protocol Amount
	---------		---------------	
	3-PGA			    3g
	2M Tris			    5.04mL
	H2O			    3.156mL


Malachite Green Dye (62.5mM):

	Component		Protocol Amount
	---------		---------------	
	Malachite green		0.0929g
	H2O			1050uL


Malachite Green Dye (500uM):

	Component		Protocol Amount
	---------		---------------	
	62.5mM MG (above)	8uL
	H2O			992  uL


Pyruvate (1250mM, pH ~ 7.5):

	Component		Protocol Amount
	---------		---------------	
	Pyruvate		    0.77g
	2M Tris-HCl pH 7.4	    4.257mL
	Tris base (2M)		    350uL
	KOH (5M)		    100uL


Maltose (1400mM, pH 7.44):

	Component		Protocol Amount
	---------		---------------	
	Maltose			    2.52g
	2M Tris-HCl pH 7.4	    4.305mL
	Tris base (2M)		    90uL


Succinate (1400mM, pH 7.54):

	Component		Protocol Amount
	---------		---------------	
	Succinate		    1.89g
	2M Tris-HCl pH 7.4	    5mL
	HCl			    55uL


Experiment Protocol:
	1. Create a minimal master mix that will be added to all samples. 
	The composition is computed from the same Jupyter notebook used to create the Echo picklist.
	   (108 samples * 7.466uL/sample * 1.1 excess = 887uL)
		Component				Protocol Amount (uL)
		---------				--------------------
		Extract					392 (15 tubes)
		PEG (40%)				59.40
		Minimal energy sol. (24.862 x)		47.78
		Amino acids				297
		K(3000mM)				55.44
		Malachite Green Dye			23.76
		NTPs (500mM)				11.51
	2. We use the Echo to load whatever is not in the minimal master mix into our sample wells, as defined by
	   our Echo picklist. We make sure to leave one well space between wells with different samples.
		Source Well		Component			Volume (uL)
		-----------	-------------------------		-----------
		    M01		3-PGA, diluted 3X to 467mM		65	
		    M02		3-PGA, diluted 3X to 467mM		30	
		    M04		Mg-glutamate, 100mM			65    
		    M05		Mg-glutamate, 400mM			35
		    M07		DNA,  144 ng/uL				40
		    M09		H2O					65
		    M10		H2O					65
		    M11		H2O					65
		    M12		H2O					65
	3. Load 7.47uL of minimal master mix into each sample.
	4. Perform the following BioTek protocol.
		  Start Kinetic	Runtime 18:00:00 (HH:MM:SS), Interval 0:03:00, 193 Reads							
			- Linear shake: 0:05 (MM:SS), frequency: 360 cpm (6 mm)	
			- Read MGaptmer
			  Filter Set 1							
	    			Excitation: 610,  Emission: 650							
	    			Optics: Bottom,  Gain: 150													
			  Light Source: Xenon Flash,  Lamp Energy: High							
			  Read Speed: Normal,  Delay: 100 msec,  Measurements/Data Point: 10							
			  Read Height: 9.5 mm							
    			- Read deGFP														
			  Filter Set 1							
	    			Excitation: 485,  Emission: 515							
	    			Optics: Bottom,  Gain: 61													
			  Light Source: Xenon Flash,  Lamp Energy: High							
			  Read Speed: Normal,  Delay: 100 msec,  Measurements/Data Point: 10							
			  Read Height: 7 mm		


Results:
- We see a clear positive correlation between Mg and 3-PGA concentration. That is, as we increase [3-PGA], a higher 
[Mg] is required to achieve the same concentration of deGFP expression.
- As we increase Mg concentration, we see an interesting biphasic protein expression, especially at lower [3-PGA].
- MG aptamer read poor due to low DNA concentration. Data collected but not shown here.						
