## Supplementary material for "Metabolic perturbations to an *E. coli*-based cell-free system reveal a trade-off between transcription and translation": Raw Data, Tidy Data, DNA GenBank files, Figures, and Jupyter Notebooks for Figure Generation and Analysis: readme.txt

Description: In collaboration with William Poole and Zoila Jurado, I am doing experiments to probe cell extract metabolism.
In this experiment and subsequent experiments, we vary the amount of fuel we are using at the following levels: 5mM, 10mM,
15mM, 20mM, 30mM, and 45mM. We are effectively performing a panel of fuel vs Mg to understand the complex interplay between
fuel and magnesium. We use a Jupyter notebook to compute the final concentrations of these components in the TXTL 
reaction mixture. 
	We create the following solutions for our experiments: a minimal energy solution without 3-PGA and NTPs; ATP 
solution, GTP solution, NTP solution, 3-PGA solution (from Miki), pyruvate solution, succinate solution, and maltose
solution. We also create a malachite green solution to measure mRNA dynamics.
	In this experiment, we perform an NTP by ATP by GTP panel at fixed 2mM Mg+2 (previous optimal for other fuel sources).
Here we do not measure mRNA dynamics since it doesn't seem to work well for this extract anyways.


Minimal energy solution (5.774mL/7.33):

	Component		Protocol Amount		Amount for 788uL (/7.33)
	---------		---------------		-------------------------
	HEPES			3.6mL				491.13uL
	tRNA			576uL				78.58uL
	CoA			576uL				78.58uL
	cAMP			170uL				23.19uL
	Folinic acid		288uL				39.29uL
	Spermidine		144uL				19.65uL
	NAD+			276uL				37.65uL
	H2O			144uL				19.65uL


GTP (760mM):

	Component		Protocol Amount		Actual amount (/4.237)
	---------		---------------		-------------------------
	GTP			   500mg			118mg
	H2O			   1245uL			294uL


3-PGA (1400mM):

	Component		Protocol Amount
	---------		---------------	
	3-PGA			    3g
	2M Tris			    5.04mL
	H2O			    3.156mL


Malachite Green Dye (62.5mM):

	Component		Protocol Amount
	---------		---------------	
	Malachite green		0.0929g
	H2O			1050uL


Malachite Green Dye (500uM):

	Component		Protocol Amount
	---------		---------------	
	62.5mM MG (above)	8uL
	H2O			992  uL


Pyruvate (1250mM, pH ~ 7.5):

	Component		Protocol Amount
	---------		---------------	
	Pyruvate		    0.77g
	2M Tris-HCl pH 7.4	    4.257mL
	Tris base (2M)		    350uL
	KOH (5M)		    100uL


Maltose (1400mM, pH 7.44):

	Component		Protocol Amount
	---------		---------------	
	Maltose			    2.52g
	2M Tris-HCl pH 7.4	    4.305mL
	Tris base (2M)		    90uL


Succinate (1400mM, pH 7.54):

	Component		Protocol Amount
	---------		---------------	
	Succinate		    1.89g
	2M Tris-HCl pH 7.4	    5mL
	HCl			    55uL


Experiment Protocol:
	1. Create a minimal master mix that will be added to all samples. 
	The composition is computed from the same Jupyter notebook used to create the Echo picklist.
	   (228 samples * 7.169uL/sample * 1.1 excess = 1635uL)
		Component				Protocol Amount (uL)
		---------				--------------------
		Extract					828 (30 tubes)
		PEG (40%)				125.4
		Minimal energy sol. (24.862 x)		100.9 (3 tubes)
		Amino acids				627
		K(3000mM)				117				
	2. We use the Echo to load whatever is not in the minimal master mix into our sample wells, as defined by
	   our Echo picklist. We make sure to leave one well space between wells with different samples.
		Source Well		Component			Volume (uL)
		-----------	-------------------------		-----------
		    J01		NTPs, diluted 3X to 167mM		65	
		    J02		NTPs, diluted 3X to 167mM		65	
		    J03		NTPs, diluted 3X to 167mM		35    
		    J05		ATP, diluted 3X to 180mM		65
		    J06		ATP, diluted 3X to 180mM		65
		    J08		GTP, diluted 4X to 190mM		65
		    J09		GTP, diluted 4X to 190mM		65
		    J11		Mg-glutamate, diluted 2X to 200mM	60
		    J13		DNA,  283 ng/uL				40
		    J15		Malachite Green Dye (500uM)		32
		    J17		H2O					65
		    J18		H2O					65
		    J19		H2O					65
		    J20		H2O					65
		    J21		H2O					65
		    J22		H2O					65
		    J23		H2O					65
		    J24		H2O					65
	3. Load 7.17uL of minimal master mix into each sample.
	4. Perform the following BioTek protocol.
		  Start Kinetic	Runtime 18:00:00 (HH:MM:SS), Interval 0:03:10							
			- Linear shake: 0:05 (MM:SS), frequency: 360 cpm (6 mm)							
    			- Read deGFP														
			  Filter Set 1							
	    			Excitation: 485,  Emission: 515							
	    			Optics: Bottom,  Gain: 61													
			  Light Source: Xenon Flash,  Lamp Energy: High							
			  Read Speed: Normal,  Delay: 100 msec,  Measurements/Data Point: 10							
			  Read Height: 7 mm

Results:
- Increased NTP concentrations increase the time it takes for protein expression to begin	
- Increased GTP, but not ATP, concentrations improve protein yields
- Biphasic protein production occurs in absence of fuel				
