## Supplementary material for "Metabolic perturbations to an *E. coli*-based cell-free system reveal a trade-off between transcription and translation": Raw Data, Tidy Data, DNA GenBank files, Figures, and Jupyter Notebooks for Figure Generation and Analysis: readme.txt

Description: In collaboration with William Poole and Zoila Jurado, I am doing experiments to probe cell extract metabolism.
In this experiment and subsequent experiments, we vary the amount of fuel we are using at the following levels: 5mM, 10mM,
15mM, 20mM, 30mM, and 45mM. We are effectively performing a panel of fuel vs Mg to understand the complex interplay between
fuel and magnesium. We use a Jupyter notebook to compute the final concentrations of these components in the TXTL 
reaction mixture. 
	We create the following solutions for our experiments: a minimal energy solution without 3-PGA and NTPs; ATP 
solution, GTP solution, NTP solution, 3-PGA solution (from Miki), pyruvate solution, succinate solution, and maltose
solution. We also create a malachite green solution to measure mRNA dynamics.
	In this experiment, we perform an NTP by ATP by GTP panel at fixed 0mM Mg+2 (previous optimal for other fuel 
sources at 5nM DNA).


Minimal energy solution (5.774mL/7.33):

	Component		Protocol Amount		Amount for 788uL (/7.33)
	---------		---------------		-------------------------
	HEPES			3.6mL				491.13uL
	tRNA			576uL				78.58uL
	CoA			576uL				78.58uL
	cAMP			170uL				23.19uL
	Folinic acid		288uL				39.29uL
	Spermidine		144uL				19.65uL
	NAD+			276uL				37.65uL
	H2O			144uL				19.65uL


GTP (760mM):

	Component		Protocol Amount		Actual amount (/4.237)
	---------		---------------		-------------------------
	GTP			   500mg			118mg
	H2O			   1245uL			294uL


3-PGA (1400mM):

	Component		Protocol Amount
	---------		---------------	
	3-PGA			    3g
	2M Tris			    5.04mL
	H2O			    3.156mL


Malachite Green Dye (62.5mM):

	Component		Protocol Amount
	---------		---------------	
	Malachite green		0.0929g
	H2O			1050uL


Malachite Green Dye (500uM):

	Component		Protocol Amount
	---------		---------------	
	62.5mM MG (above)	8uL
	H2O			992  uL


Pyruvate (1250mM, pH ~ 7.5):

	Component		Protocol Amount
	---------		---------------	
	Pyruvate		    0.77g
	2M Tris-HCl pH 7.4	    4.257mL
	Tris base (2M)		    350uL
	KOH (5M)		    100uL


Maltose (1400mM, pH 7.44):

	Component		Protocol Amount
	---------		---------------	
	Maltose			    2.52g
	2M Tris-HCl pH 7.4	    4.305mL
	Tris base (2M)		    90uL


Succinate (1400mM, pH 7.54):

	Component		Protocol Amount
	---------		---------------	
	Succinate		    1.89g
	2M Tris-HCl pH 7.4	    5mL
	HCl			    55uL


Experiment Protocol:
	1. Create a minimal master mix that will be added to all samples. 
	The composition is computed from the same Jupyter notebook used to create the Echo picklist.
	   (228 samples * 7.369uL/sample * 1.1 excess = 1848uL)
		Component				Protocol Amount (uL)
		---------				--------------------
		Extract					828 (30 tubes)
		PEG (40%)				125.4
		Minimal energy sol. (24.862 x)		100.9 (3 tubes)
		Amino acids (6mM)			627
		K(3000mM)				117	
		Malachite Green Dye (500uM)		50.16		
	2. Prepare diluted NTP solutions.
		- 60uL NTPs + 110uL H2O
		- 50uL ATP + 87.5uL H2O
		- 35uL GTP + 105uL H2O
	2. We use the Echo to load whatever is not in the minimal master mix into our sample wells, as defined by
	   our Echo picklist. We make sure to leave one well space between wells with different samples.
		Source Well		Component			Volume (uL)
		-----------	-------------------------		-----------
		    O01		NTPs, diluted 2.5X to 200mM		65	
		    O02		NTPs, diluted 2.5X to 200mM		60	    
		    O04		ATP, diluted 2.75X to 196mM		65
		    O05		ATP, diluted 2.75X to 196mM		55
		    O07		GTP, diluted 4X to 190mM		65
		    O08		GTP, diluted 4X to 190mM		55
		    O10		Mg-glutamate, 100mM			42
		    O12		DNA,  425 ng/uL				65
		    O13		DNA,  425 ng/uL				32
		    O15		H2O					65
		    O16		H2O					65
		    O17		H2O					65
		    O18		H2O					65
		    O22		H2O					65
		    O23		H2O					65
		    O24		H2O					65
	3. Load 7.37uL of minimal master mix into each sample.
	4. Perform the following BioTek protocol.
		  Start Kinetic	Runtime 18:00:00 (HH:MM:SS), Interval 0:04:00							
			- Linear shake: 0:05 (MM:SS), frequency: 360 cpm (6 mm)	
			- Read MGaptmer
			  Filter Set 1							
	    			Excitation: 610,  Emission: 650							
	    			Optics: Bottom,  Gain: 150													
			  Light Source: Xenon Flash,  Lamp Energy: High							
			  Read Speed: Normal,  Delay: 100 msec,  Measurements/Data Point: 10							
			  Read Height: 9.5 mm						
    			- Read deGFP														
			  Filter Set 1							
	    			Excitation: 485,  Emission: 515							
	    			Optics: Bottom,  Gain: 61													
			  Light Source: Xenon Flash,  Lamp Energy: High							
			  Read Speed: Normal,  Delay: 100 msec,  Measurements/Data Point: 10							
			  Read Height: 7 mm

Results:
	- deGFP expression strongly favors more GTP than ATP
	- mRNA expression cannot be used to predict deGFP yields or dynamics, apart from the general trend that sharp transcriptional peak corresponds to more expression
	
