## Supplementary material for "Metabolic perturbations to an *E. coli*-based cell-free system reveal a trade-off between transcription and translation": Raw Data, Tidy Data, DNA GenBank files, Figures, and Jupyter Notebooks for Figure Generation and Analysis: readme.txt

Description: We perform a salt calibration of the extract prepared by Alex Johnson, Miki Yun, and Manisha Kapasiawala in
July 2022 via the French Press and subsequent dialysis. We follow the lab's standard extract preparation protocol.
This extract had a protein concentration of roughly 30mg/ml using the Bradford assay.

Protocol: 
	0. Create an Echo picklist to titrate Mg and K at the following concentrations:
		- Mg (mM): 2, 4, 6, 8, 10, 12
		- K (mM): 40, 60, 80, 100, 120, 140
	1. Create a minimal master mix that will be added to all samples. 
	The composition is computed from the same Jupyter notebook used to create the Echo picklist.
	   (36 samples * 7.014uL/sample * 1.1 excess = 276uL)
		Component				Protocol Amount (uL)
		---------				--------------------
		Extract					130.68
		PEG (40%)				19.8
		Energy solution (14x)			28.28
		Amino acids (6mM)			99			
	2. We use the Echo to load whatever is not in the minimal master mix into our sample wells, as defined by
	   our Echo picklist. We make sure to leave one well space between wells with different samples.
		Source Well		Component			Volume (uL)
		-----------	-------------------------		-----------
		    L01		Mg-glutamate, 100mM			42	
		    L03		K-glutamate, 1000mM			57	    
		    L05		DNA, 228 ng/uL				26
		    L07		Water					65
		    L08		Water					65
	3. Load 7.02uL of minimal master mix into each sample.
	4. Perform the following BioTek protocol.
		  Start Kinetic	Runtime 18:00:00 (HH:MM:SS), Interval 0:04:00							
			- Linear shake: 0:05 (MM:SS), frequency: 360 cpm (6 mm)						
    			- Read deGFP														
			  Filter Set 1							
	    			Excitation: 485,  Emission: 515							
	    			Optics: Bottom,  Gain: 61													
			  Light Source: Xenon Flash,  Lamp Energy: High							
			  Read Speed: Normal,  Delay: 100 msec,  Measurements/Data Point: 10							
			  Read Height: 7 mm


Results: 10uM deGFP at 8mM Mg, 80mM K.
