## Supplementary material for "Metabolic perturbations to an *E. coli*-based cell-free system reveal a trade-off between transcription and translation": Raw Data, Tidy Data, DNA GenBank files, Figures, and Jupyter Notebooks for Figure Generation and Analysis: readme.txt

Description: In collaboration with William Poole and Zoila Jurado, I am doing experiments to probe cell extract metabolism.
In this experiment and subsequent experiments, we vary the amount of fuel we are using at the following levels: 5mM, 10mM,
15mM, 20mM, 30mM, and 45mM. We are effectively performing a panel of fuel vs Mg to understand the complex interplay between
fuel and magnesium. We use a Jupyter notebook to compute the final concentrations of these components in the TXTL 
reaction mixture. 
	We create the following solutions for our experiments: a minimal energy solution without 3-PGA and NTPs; ATP 
solution, GTP solution, NTP solution, 3-PGA solution (from Miki), pyruvate solution, succinate solution, and maltose
solution. We also create a malachite green solution to measure mRNA dynamics.
	In this experiment, we perform a 3-PGA vs Magnesium panel, albeit unlike the 20220513 expt, we use 5nM DNA 
template instead of 1nM and we use a dialyzed, French-pressed extract. Here we use a newly prepared batch of energy
solution. Here we also note a newly prepared NTP solution but do not use it yet.


Minimal energy solution (5.774mL/3.6):

	Component		Protocol Amount		Amount for 1604uL (/3.6)
	---------		---------------		-------------------------
	HEPES			3.6mL				1000uL
	tRNA			576uL				160uL
	CoA			576uL				160uL
	cAMP			170uL				47.22uL
	Folinic acid		288uL				80uL
	Spermidine		144uL				40uL
	NAD+			276uL				76.67uL
	H2O			144uL				40uL


GTP (760mM):

	Component		Protocol Amount		Actual amount (/4.237)
	---------		---------------		-------------------------
	GTP			   500mg			118mg
	H2O			   1245uL			294uL


CTP (621mM):

	Component		Protocol Amount		Actual amount (/1.667)
	---------		---------------		-------------------------
	CTP			   300mg			180mg
	H2O			   708uL			424.8uL


UTP (813mM):

	Component		Protocol Amount		Actual amount (/1.345)
	---------		---------------		-------------------------
	UTP			   300mg			223mg
	H2O			   480uL			357uL


NTP solution (500mM):

	Component		Protocol Amount		Actual amount (x2)
	---------		---------------		-------------------------
	ATP (540mM)			74uL			148uL
	GTP (760mM)			52.8uL			105.6uL
	UTP (813mM)			29.6uL			59.2uL
	CTP (621mM)			38.8uL			77.6uL
	KOH (15%)			60uL			120uL 	


3-PGA (1400mM):

	Component		Protocol Amount
	---------		---------------	
	3-PGA			    3g
	2M Tris			    5.04mL
	H2O			    3.156mL


Malachite Green Dye (62.5mM):

	Component		Protocol Amount
	---------		---------------	
	Malachite green		0.0929g
	H2O			1050uL


Malachite Green Dye (500uM):

	Component		Protocol Amount
	---------		---------------	
	62.5mM MG (above)	8uL
	H2O			992  uL


Pyruvate (1250mM, pH ~ 7.5):

	Component		Protocol Amount
	---------		---------------	
	Pyruvate		    0.77g
	2M Tris-HCl pH 7.4	    4.257mL
	Tris base (2M)		    350uL
	KOH (5M)		    100uL


Maltose (1400mM, pH 7.44):

	Component		Protocol Amount
	---------		---------------	
	Maltose			    2.52g
	2M Tris-HCl pH 7.4	    4.305mL
	Tris base (2M)		    90uL


Succinate (1400mM, pH 7.54):

	Component		Protocol Amount
	---------		---------------	
	Succinate		    1.89g
	2M Tris-HCl pH 7.4	    5mL
	HCl			    55uL


Experiment Protocol:
	1. Create a minimal master mix that will be added to all samples. 
	The composition is computed from the same Jupyter notebook used to create the Echo picklist.
	   (108 samples * 7.266uL/sample * 1.1 excess = 863 uL)
		Component				Protocol Amount (uL)
		---------				--------------------
		Extract					392 (15 tubes)
		PEG (40%)				59.40
		Minimal energy sol. (24.862 x)		47.78
		Amino acids				297
		K(3000mM)				31.68
		Malachite Green Dye			23.76
		NTPs (500mM)				11.51
	2. We use the Echo to load whatever is not in the minimal master mix into our sample wells, as defined by
	   our Echo picklist. We make sure to leave one well space between wells with different samples.
		Source Well		Component			Volume (uL)
		-----------	-------------------------		-----------
		    L10		3-PGA, diluted 3X to 467mM		65	
		    L11		3-PGA, diluted 3X to 467mM		30	
		    L13		Mg-glutamate, 100mM			65    
		    L14		Mg-glutamate, 100mM			35
		    L16		DNA,  228 ng/uL				65
		    L17		DNA,  228 ng/uL				36
		    L19		H2O					65
		    L20		H2O					65
		    L21		H2O					65
		    L22		H2O					65
	3. Load 7.27uL of minimal master mix into each sample.
	4. Perform the following BioTek protocol.
		  Start Kinetic	Runtime 18:00:00 (HH:MM:SS), Interval 0:03:00, 193 Reads							
			- Linear shake: 0:05 (MM:SS), frequency: 360 cpm (6 mm)	
			- Read MGaptmer
			  Filter Set 1							
	    			Excitation: 610,  Emission: 650							
	    			Optics: Bottom,  Gain: 150													
			  Light Source: Xenon Flash,  Lamp Energy: High							
			  Read Speed: Normal,  Delay: 100 msec,  Measurements/Data Point: 10							
			  Read Height: 9.5 mm							
    			- Read deGFP														
			  Filter Set 1							
	    			Excitation: 485,  Emission: 515							
	    			Optics: Bottom,  Gain: 61													
			  Light Source: Xenon Flash,  Lamp Energy: High							
			  Read Speed: Normal,  Delay: 100 msec,  Measurements/Data Point: 10							
			  Read Height: 7 mm		


Results:
- Same linear trend observed (i.e. optimal deGFP as 3-PGA increases with Mg).
- Same optimal TX at high Mg, low 3-PGA and optimal TL at low Mg, high 3-PGA.
