## Supplementary material for "Metabolic perturbations to an *E. coli*-based cell-free system reveal a trade-off between transcription and translation": Raw Data, Tidy Data, DNA GenBank files, Figures, and Jupyter Notebooks for Figure Generation and Analysis: readme.txt

Description: In collaboration with William Poole and Zoila Jurado, I am doing experiments to probe cell extract metabolism.
In this experiment and subsequent experiments, we vary the amount of fuel we are using at the following levels: 5mM, 10mM,
15mM, 20mM, 30mM, and 45mM. We are effectively performing a panel of fuel vs Mg to understand the complex interplay between
fuel and magnesium. We use a Jupyter notebook to compute the final concentrations of these components in the TXTL 
reaction mixture. 
	We create the following solutions for our experiments: a minimal energy solution without 3-PGA and NTPs; ATP 
solution, GTP solution, NTP solution, 3-PGA solution (from Miki), pyruvate solution, succinate solution, and maltose
solution. We also create a malachite green solution to measure mRNA dynamics.
	In this experiment, we try various conditions to collect a validation data set. We try the following experimental
conditions, all at 5nM DNA and 0mM Mg-glutamate: 
	- Mixed fuel: 3-PGA and maltose at a 2:1 ratio (as per Vincent's all E coli TXTL toolbox 2.0) in DAJ_MK_MY extract
		As before, we want 0, 5, 10, 15, 20, 30, 45mM fuel
	- Mixed fuel: pyruvate and succinate at a 1:1 ratio (adding a glycolysis component and tca cycle component 
		seems interesting) in AR_ZJ extract
		As before, we want 0, 5, 10, 15, 20, 30, 45mM fuel
	- Combination of extracts: AR_ZJ extract (sonicated, non-dialyzed) + DAJ_MK_MY extract (French pressed, dialyzed) 
		at the following ratios: 2:1, 1:2, 1:1. 
		As before, we want 0, 5, 10, 15, 20, 30, 45mM fuel (here we choose 3-PGA)
Here we use newly prepared batches of energy solution and NTP solution.


Minimal energy solution (5.774mL/7.33):

	Component		Protocol Amount		Amount for 788uL (/7.33)
	---------		---------------		-------------------------
	HEPES			3.6mL				491.13uL
	tRNA			576uL				78.58uL
	CoA			576uL				78.58uL
	cAMP			170uL				23.19uL
	Folinic acid		288uL				39.29uL
	Spermidine		144uL				19.65uL
	NAD+			276uL				37.65uL
	H2O			144uL				19.65uL


GTP (760mM):

	Component		Protocol Amount		Actual amount (/4.237)
	---------		---------------		-------------------------
	GTP			   500mg			118mg
	H2O			   1245uL			294uL


CTP (621mM):

	Component		Protocol Amount		Actual amount (/1.667)
	---------		---------------		-------------------------
	CTP			   300mg			180mg
	H2O			   708uL			424.8uL


UTP (813mM):

	Component		Protocol Amount		Actual amount (/1.345)
	---------		---------------		-------------------------
	UTP			   300mg			223mg
	H2O			   480uL			357uL


NTP solution (500mM):

	Component		Protocol Amount		Actual amount (x2)
	---------		---------------		-------------------------
	ATP (540mM)			74uL			148uL
	GTP (760mM)			52.8uL			105.6uL
	UTP (813mM)			29.6uL			59.2uL
	CTP (621mM)			38.8uL			77.6uL
	KOH (15%)			60uL			120uL 


3-PGA (1400mM):

	Component		Protocol Amount
	---------		---------------	
	3-PGA			    3g
	2M Tris			    5.04mL
	H2O			    3.156mL


Malachite Green Dye (62.5mM):

	Component		Protocol Amount
	---------		---------------	
	Malachite green		0.0929g
	H2O			1050uL


Malachite Green Dye (500uM):

	Component		Protocol Amount
	---------		---------------	
	62.5mM MG (above)	8uL
	H2O			992  uL


Pyruvate (1250mM, pH ~ 7.5):

	Component		Protocol Amount
	---------		---------------	
	Pyruvate		    0.77g
	2M Tris-HCl pH 7.4	    4.257mL
	Tris base (2M)		    350uL
	KOH (5M)		    100uL


Maltose (1400mM, pH 7.44):

	Component		Protocol Amount
	---------		---------------	
	Maltose			    2.52g
	2M Tris-HCl pH 7.4	    4.305mL
	Tris base (2M)		    90uL


Succinate (1400mM, pH 7.54):

	Component		Protocol Amount
	---------		---------------	
	Succinate		    1.89g
	2M Tris-HCl pH 7.4	    5mL
	HCl			    55uL


Experiment Protocol:
	1. Create a minimal master mix for each extract that will be added to the respective samples. 
	The composition is computed from the same Jupyter notebook used to create the Echo picklist.
	   (52.5samples * 7.26uL/sample * 1.1 excess = 419.3uL)
		Component				Protocol Amount (uL)
		---------				--------------------
		DAJ_MK_MY Extract			190
		PEG (40%)				28.9
		Minimal energy sol. (24.862 x)		23.2 
		Amino acids (6mM)			144.2
		K(3000mM)				15.39	
		Malachite Green Dye (500uM)		11.54	
		NTPs (500mM)				5.58	
	   (52.5 samples * 7.46uL/sample * 1.1 excess = 430.8uL)
		Component				Protocol Amount (uL)
		---------				--------------------
		AR_ZJ Extract				190 
		PEG (40%)				28.9
		Minimal energy sol. (24.862 x)		23.2 
		Amino acids (6mM)			144.3
		K(3000mM)				26.9	
		Malachite Green Dye (500uM)		11.54
		NTPs (500mM)				5.58
	2. Create diluted solutions of fuels.
		- 40uL H2O + 20uL 3-PGA
		- 20uL H2O + 10uL pyruvate/succinate/maltose
	3. We use the Echo to load whatever is not in the minimal master mix into our sample wells, as defined by
	   our Echo picklist. We make sure to leave one well space between wells with different samples.
		Source Well		Component			Volume (uL)
		-----------	-------------------------		-----------
		    A01		3-PGA, diluted 3X to 467mM		55	
		    A03		Maltose, diluted 3X to 467mM		27
		    A05		DNA, 397ng/ul				52
		    A07		Pyruvate, diluted 3X to 417mM		27
		    A09		Succinate, diluted 3X to 467mM		27
		    A11		H2O					65
		    A12		H2O					65
		    A13		H2O					65
		    A14		H2O					65
		    A15		H2O					65
	4. Load minimal master mix in the following manner:
		   Wells	     Extract		Amount (uL)
		-----------	   -----------	      ---------------
		  B2-B22	    DAJ_MK_MY		   7.26
		  B23-C21	      AR_ZJ		   7.46
		  C22-D20	    DAJ_MK_MY		   3.63
		  C22-D20	      AR_ZJ		   3.73
		  D21-E19	    DAJ_MK_MY		   4.84
		  D21-E19	      AR_ZJ		   2.49
		  E20-F18	    DAJ_MK_MY		   2.42
		  E20-F18	      AR_ZJ		   4.89
	4. Perform the following BioTek protocol.
		  Start Kinetic	Runtime 18:00:00 (HH:MM:SS), Interval 0:04:00							
			- Linear shake: 0:05 (MM:SS), frequency: 360 cpm (6 mm)	
			- Read MGaptmer
			  Filter Set 1							
	    			Excitation: 610,  Emission: 650							
	    			Optics: Bottom,  Gain: 150													
			  Light Source: Xenon Flash,  Lamp Energy: High							
			  Read Speed: Normal,  Delay: 100 msec,  Measurements/Data Point: 10							
			  Read Height: 9.5 mm						
    			- Read deGFP														
			  Filter Set 1							
	    			Excitation: 485,  Emission: 515							
	    			Optics: Bottom,  Gain: 61													
			  Light Source: Xenon Flash,  Lamp Energy: High							
			  Read Speed: Normal,  Delay: 100 msec,  Measurements/Data Point: 10							
			  Read Height: 7 mm

Results:
- See plots (too much going on to make any conclusion here)
			
