## Supplementary figures and images for "Metabolic perturbations to an *E. coli*-based cell-free system reveal a trade-off between transcription and translation"

### Fig0.png

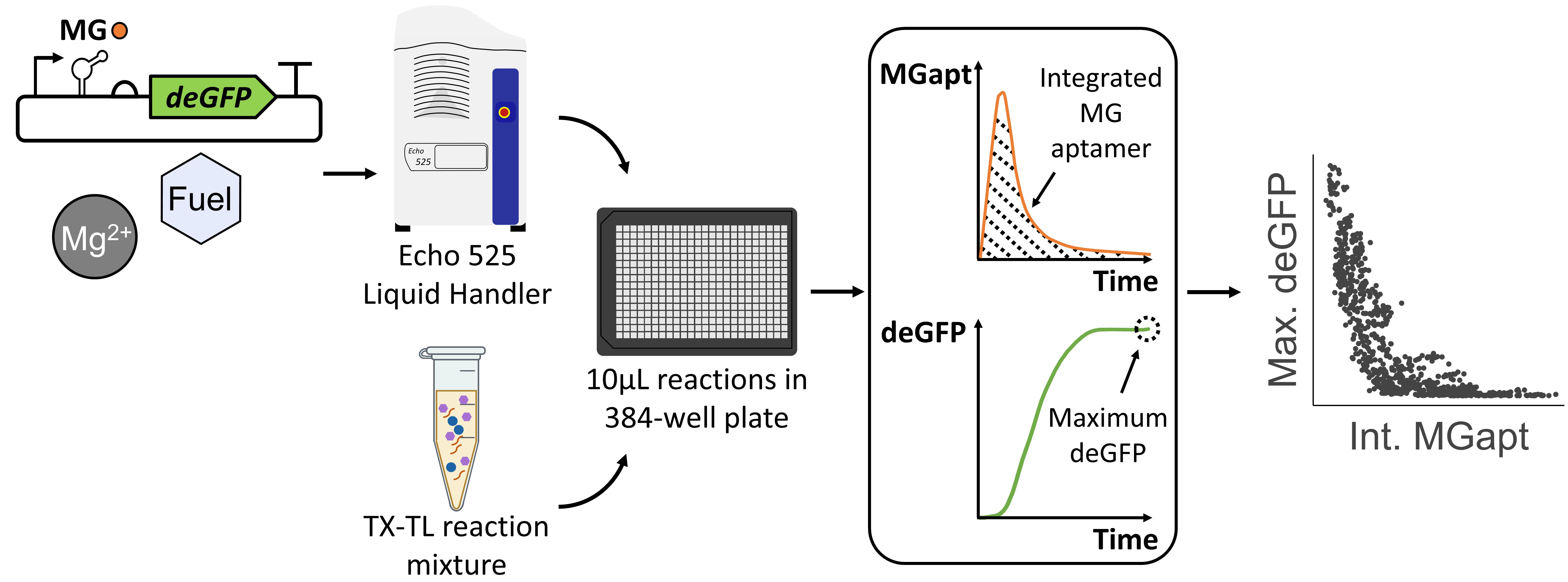

### Fig0_a.png

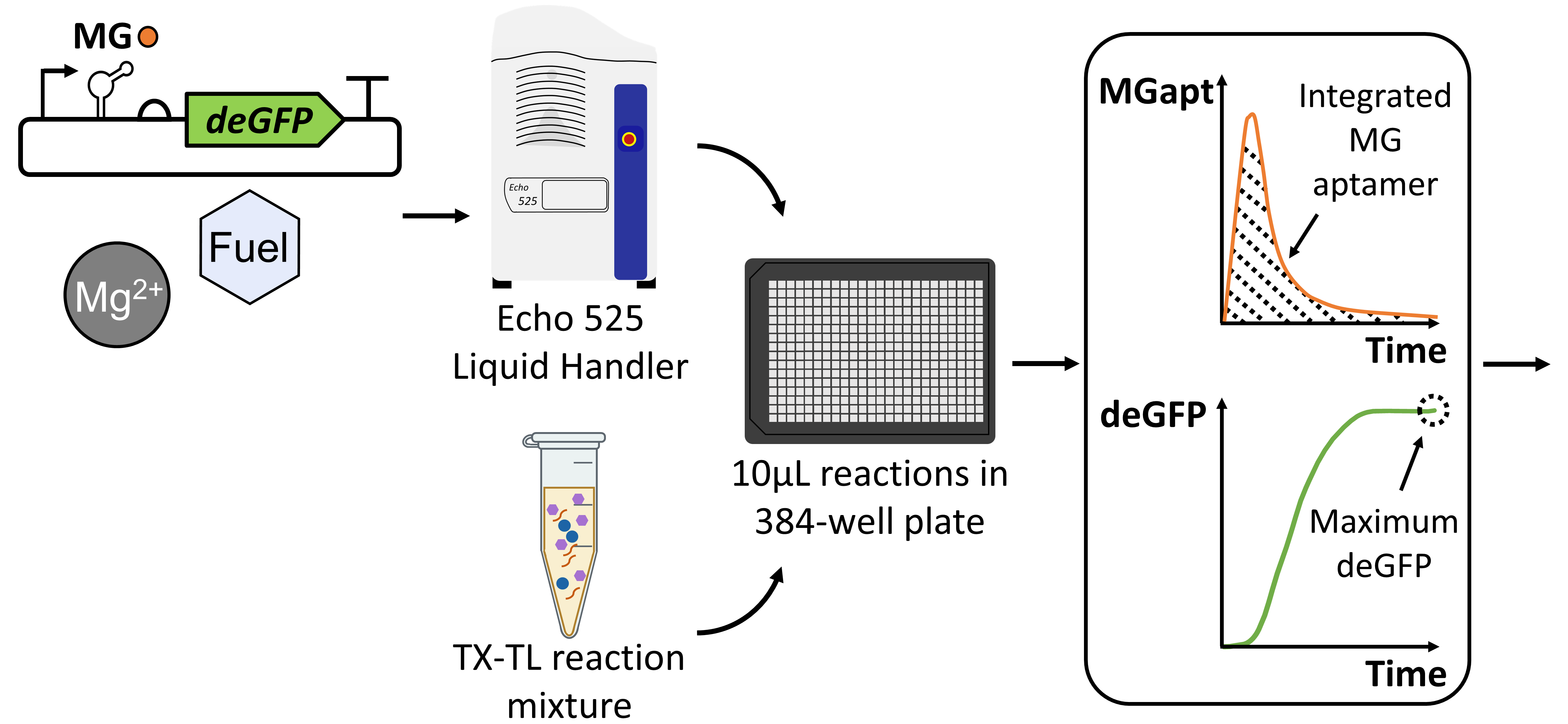

### Fig0_b.png

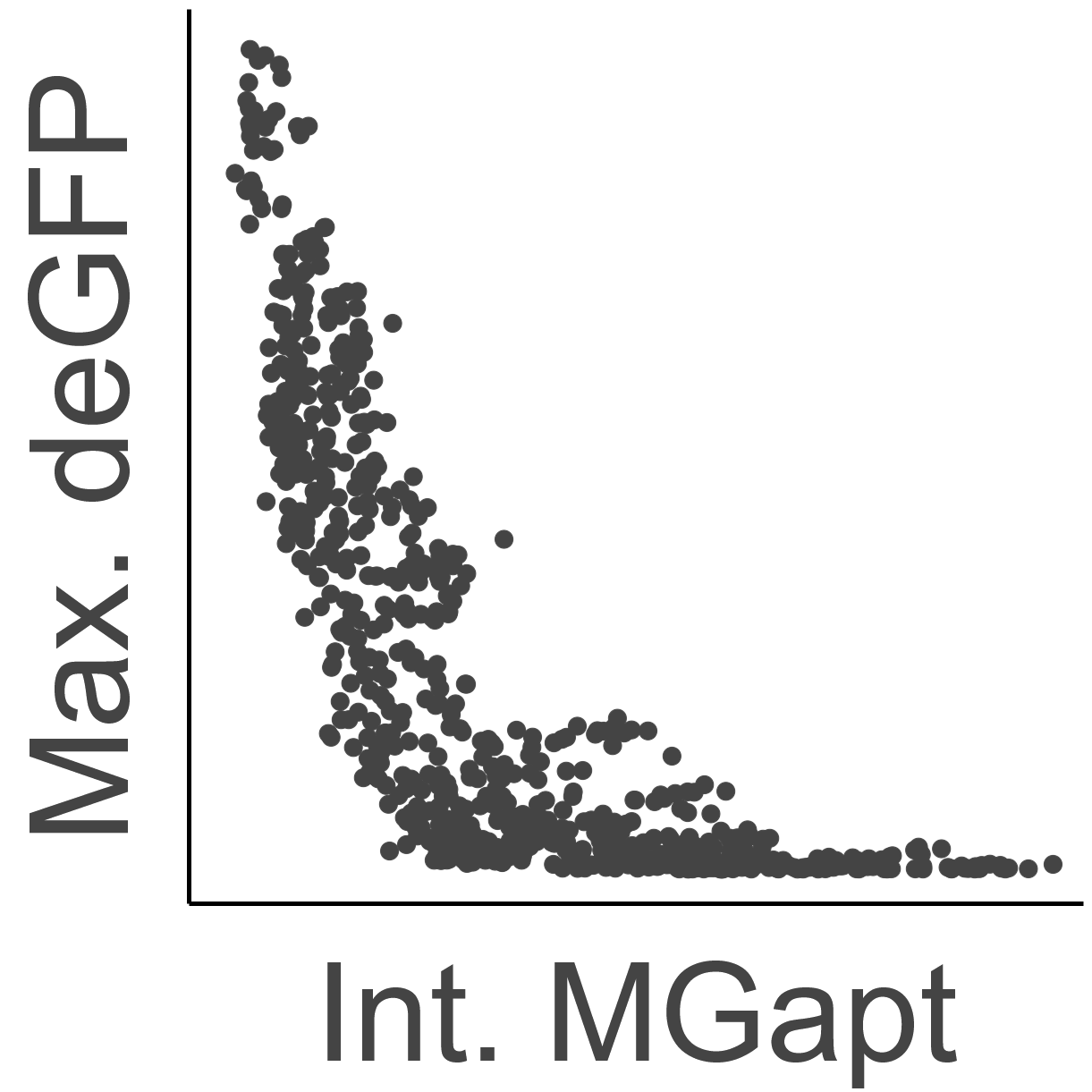

### Fig1.png

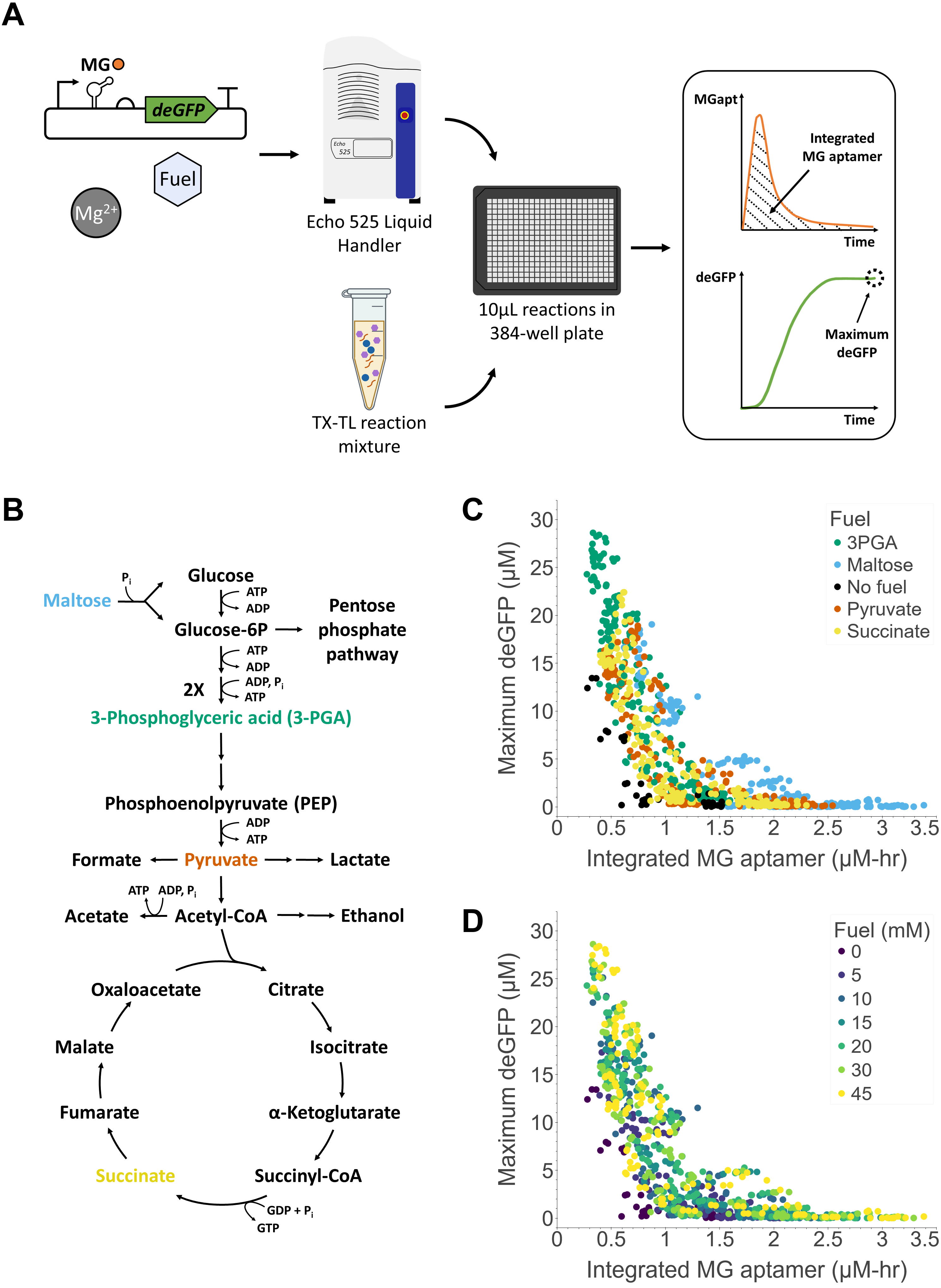

### Fig1_a.png

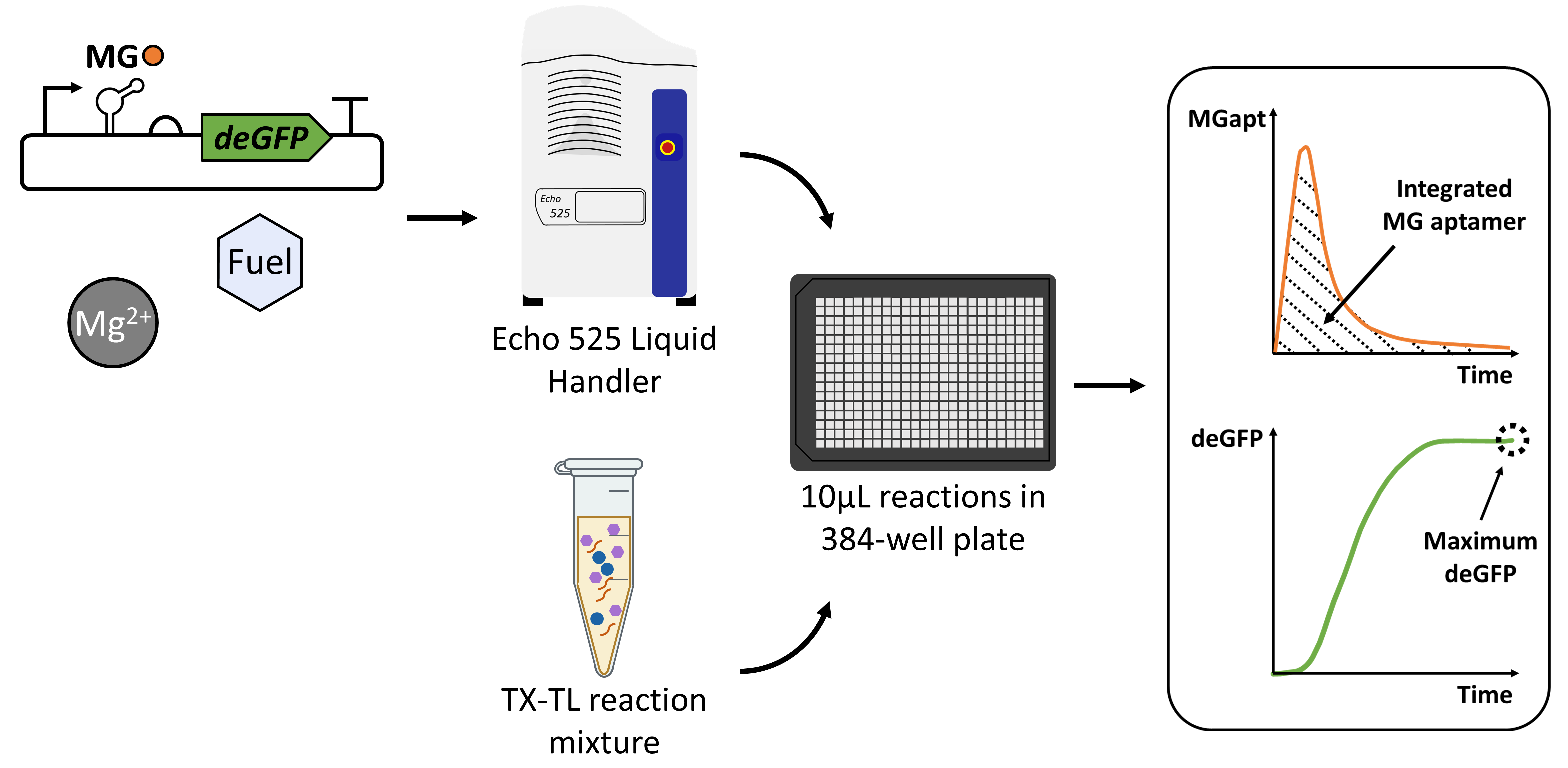

### Fig1_a.pptx

#### Slide 1
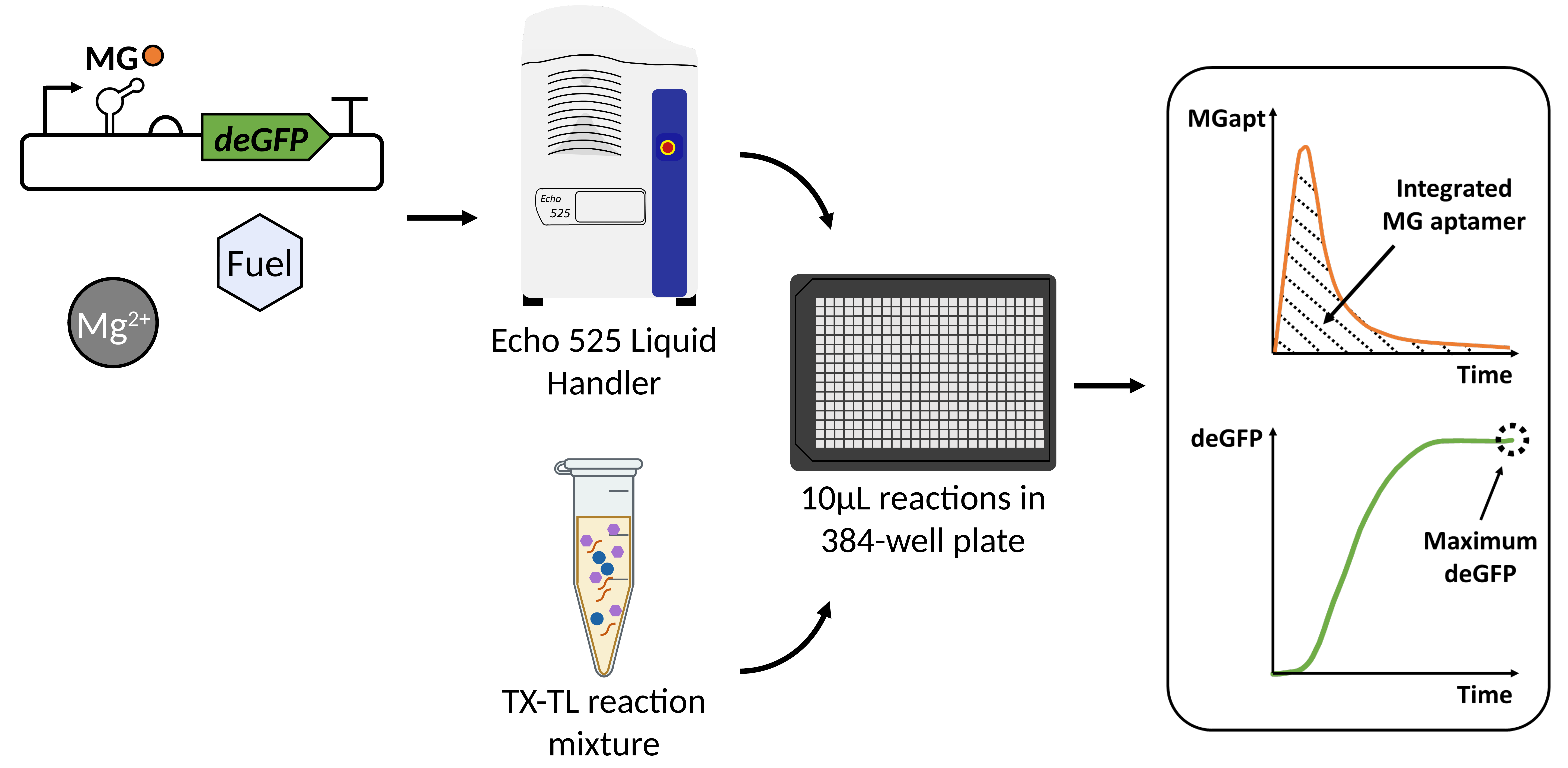

MG
deGFP
*
Fuel
Mg^2+^
Echo 525 Liquid Handler
TX-TL reaction mixture
10µL reactions in 384-well plate

### Fig1_b.png

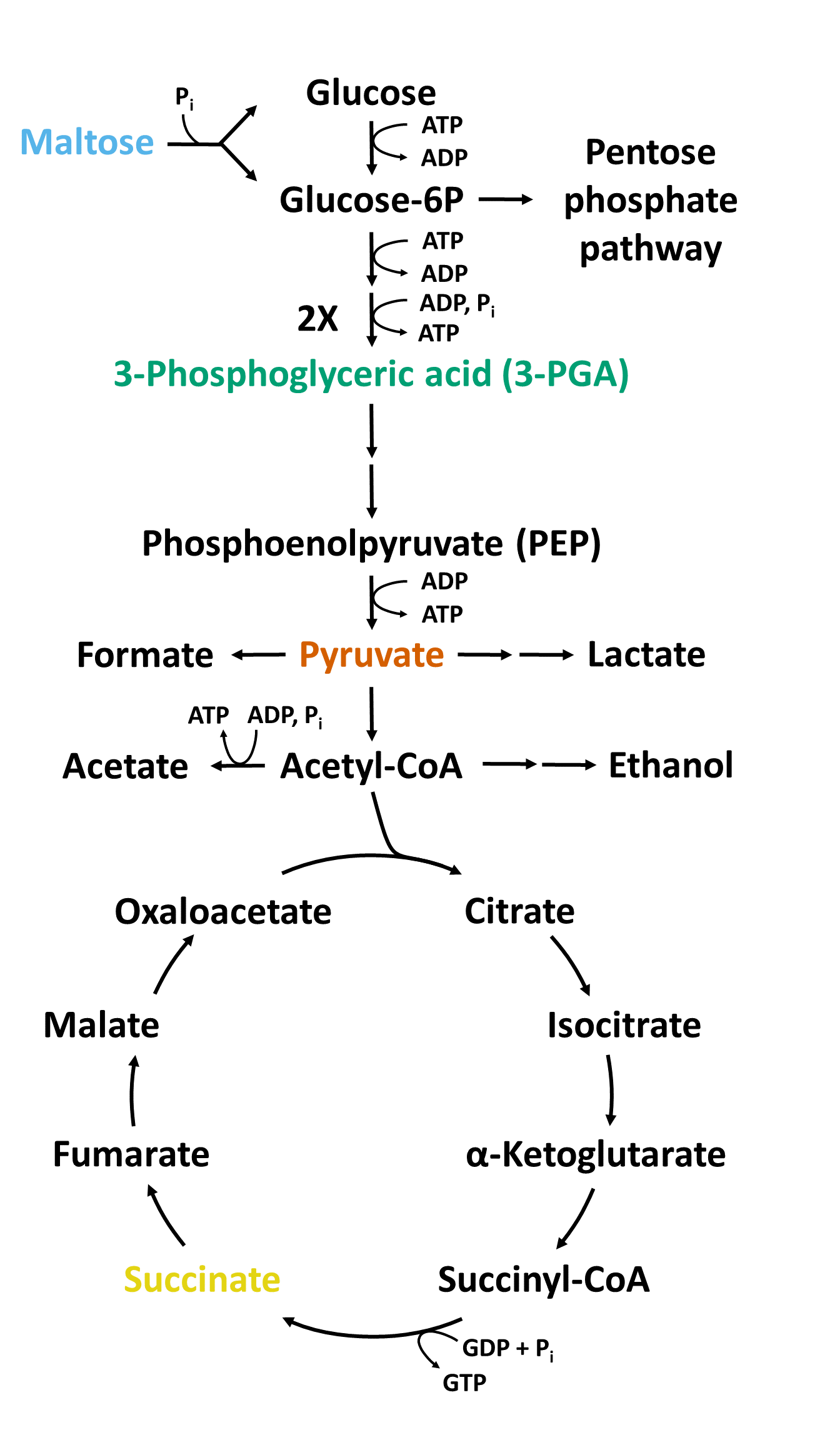

### Fig1_c.png

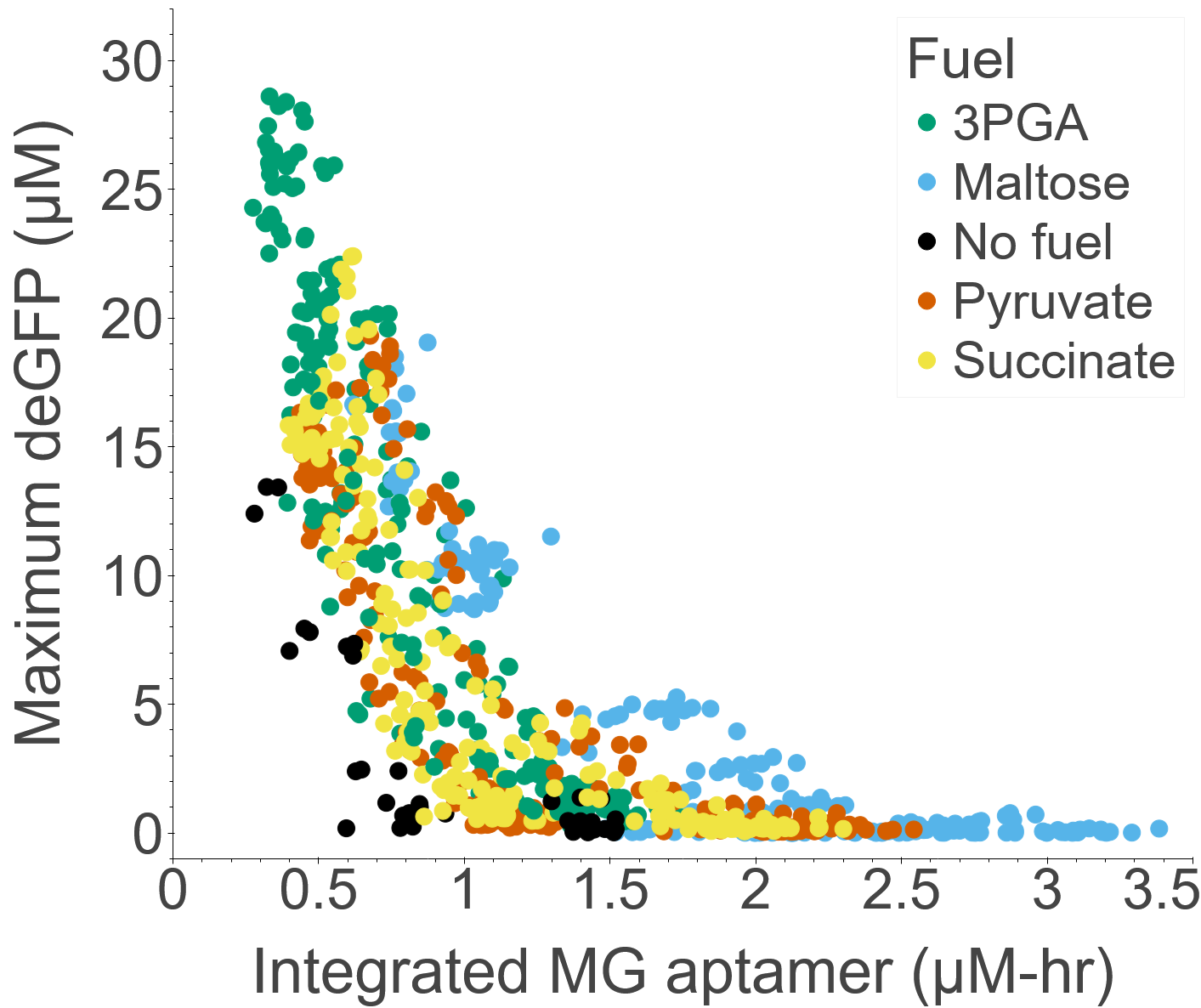

### Fig1_d.png

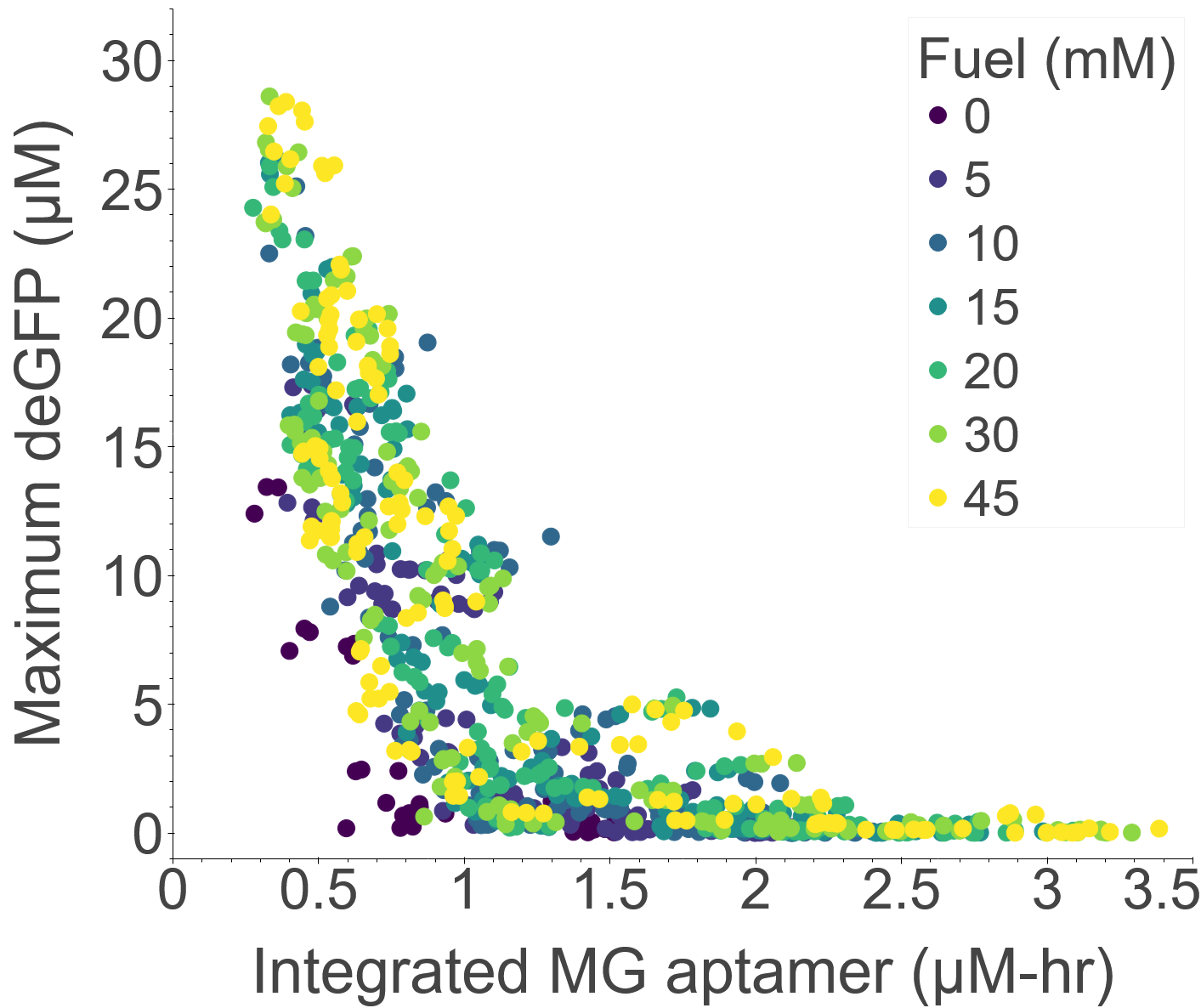

### Fig2.png

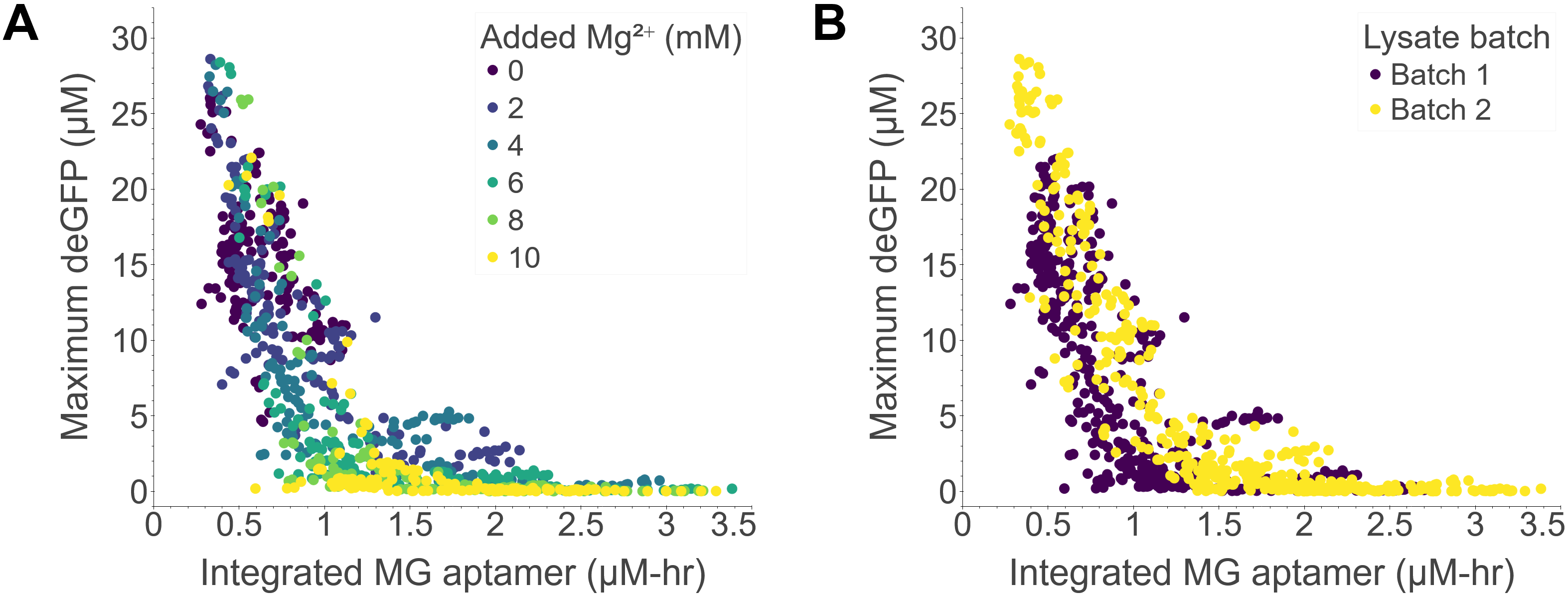

### Fig2_a.png

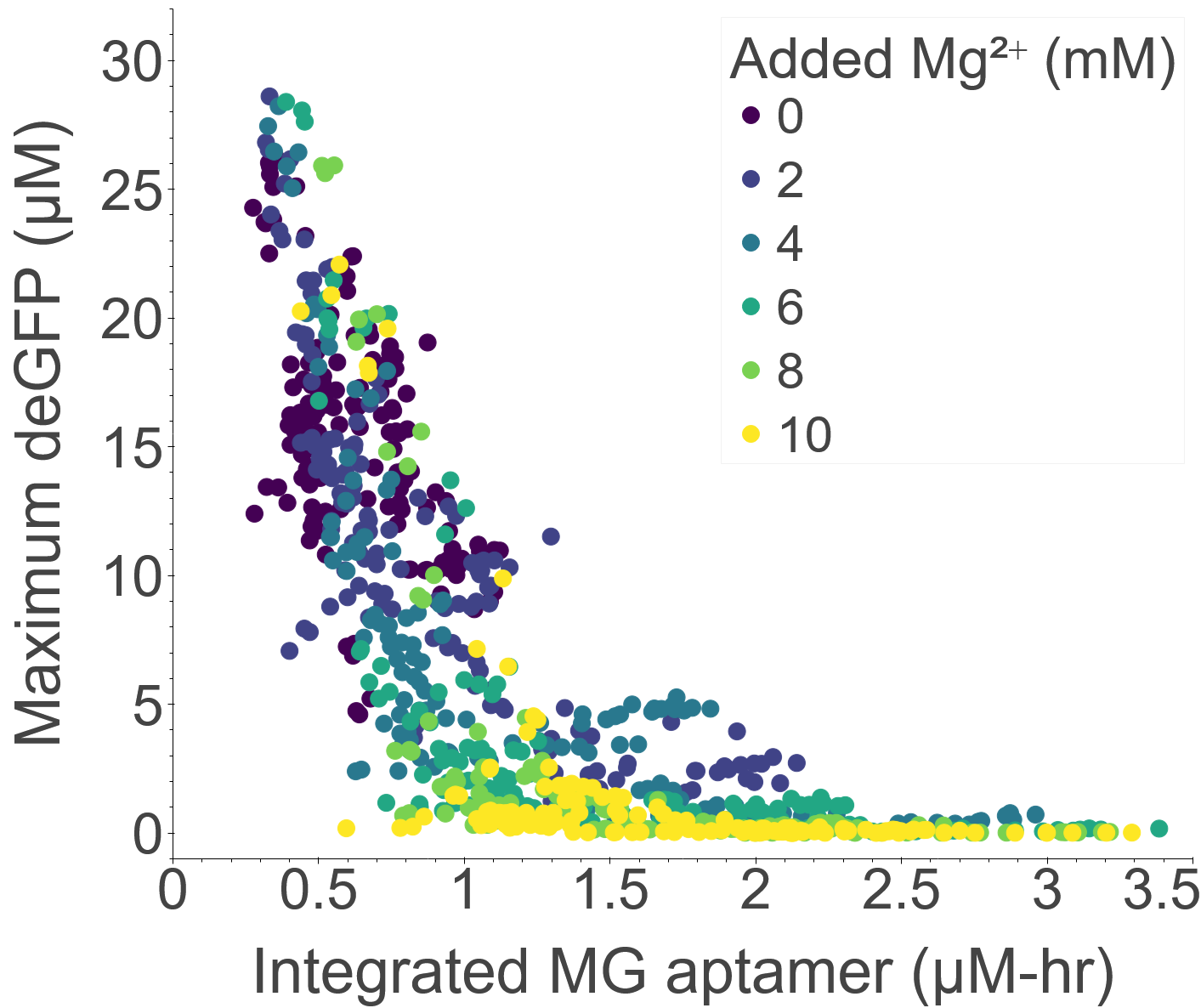

### Fig2_b.png

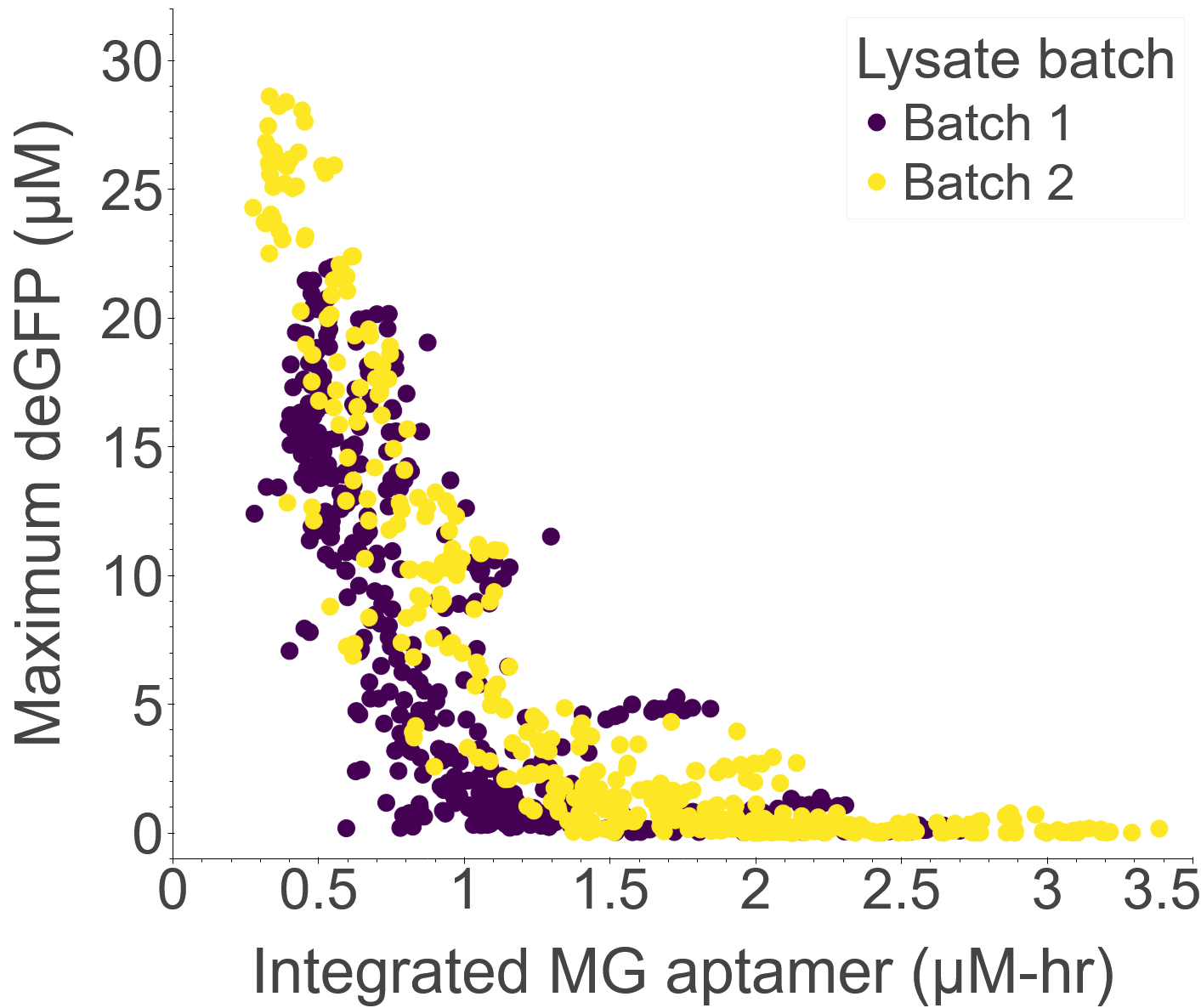

### Fig3.png

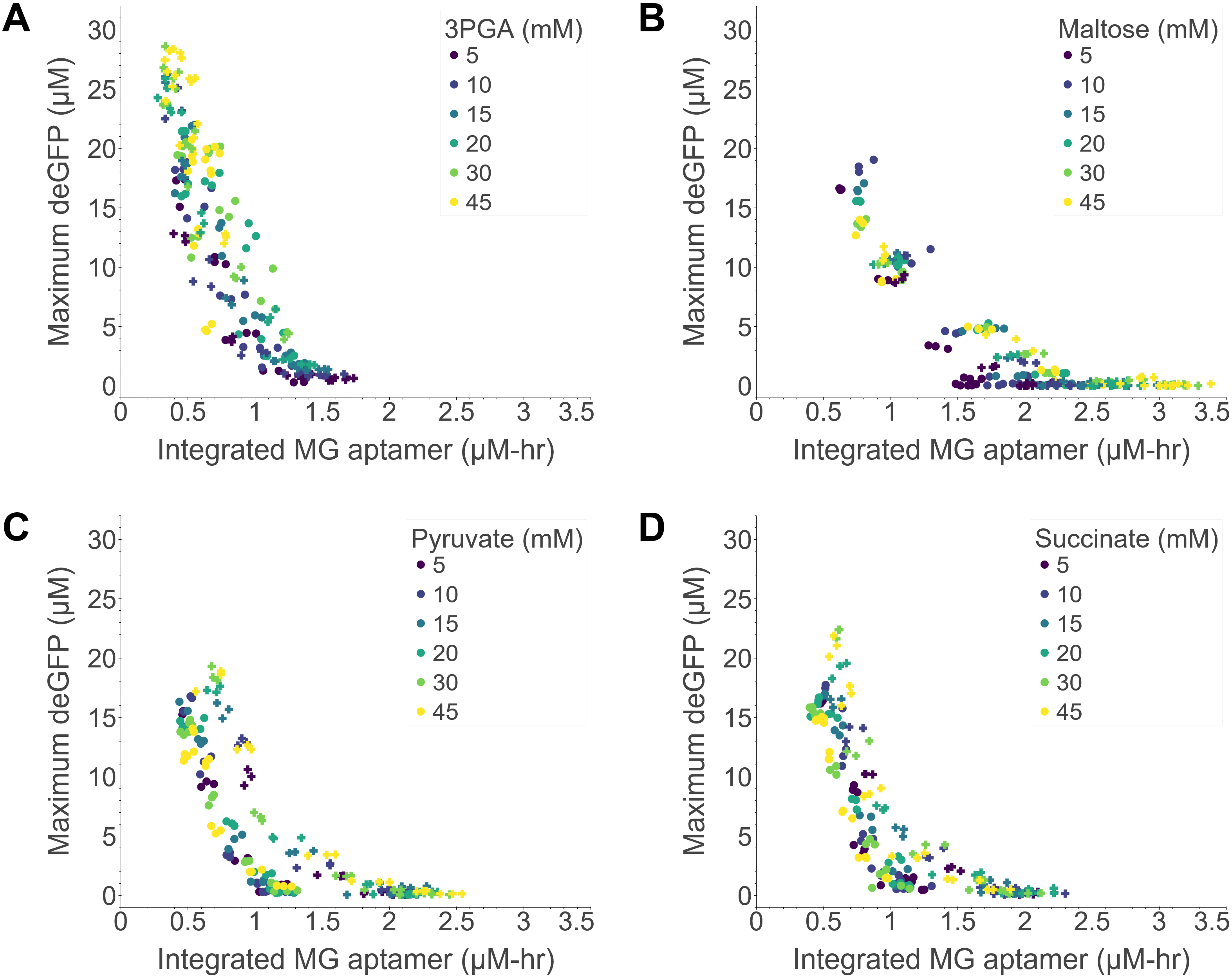

### Fig3_a.png

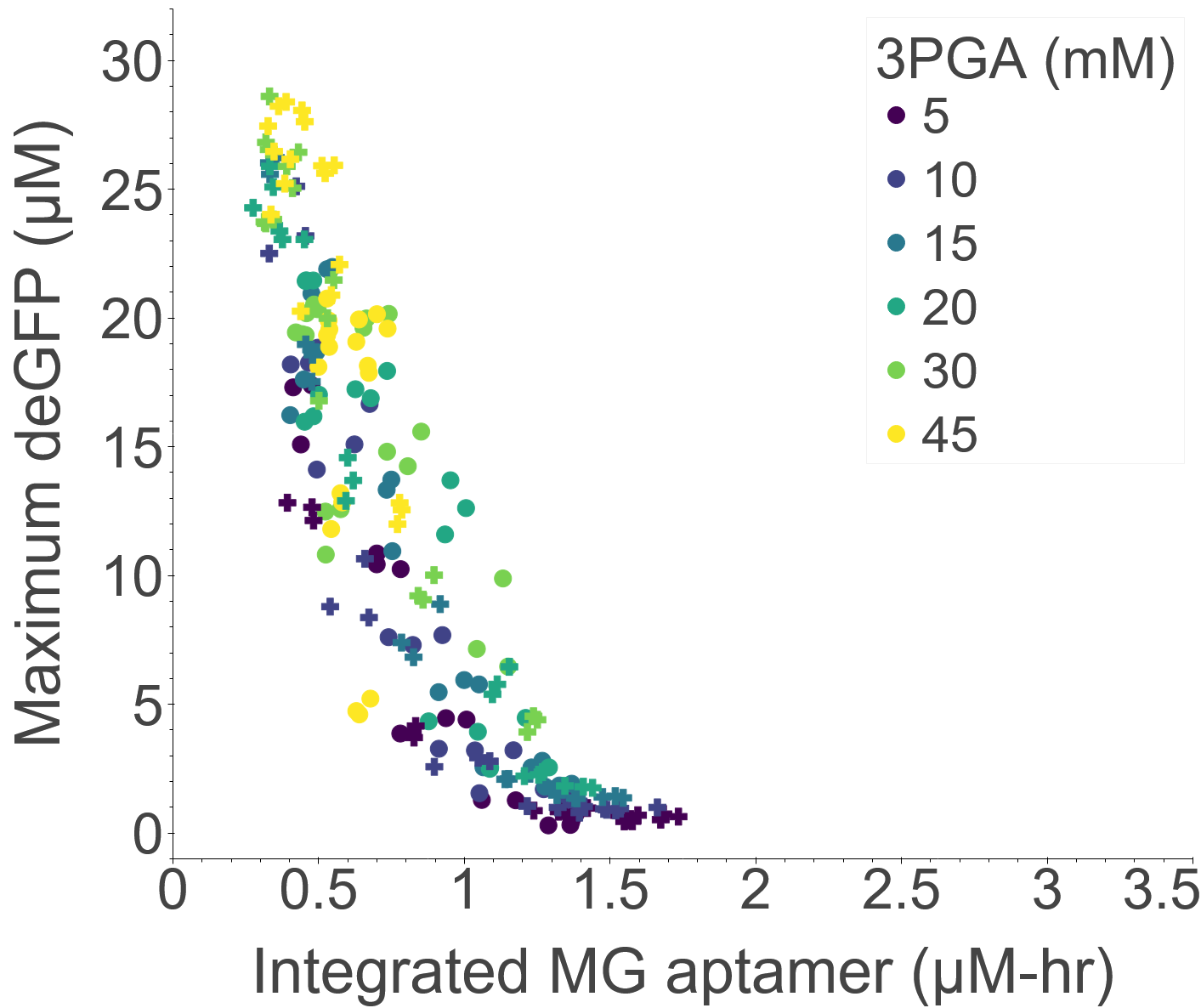

### Fig3_b.png

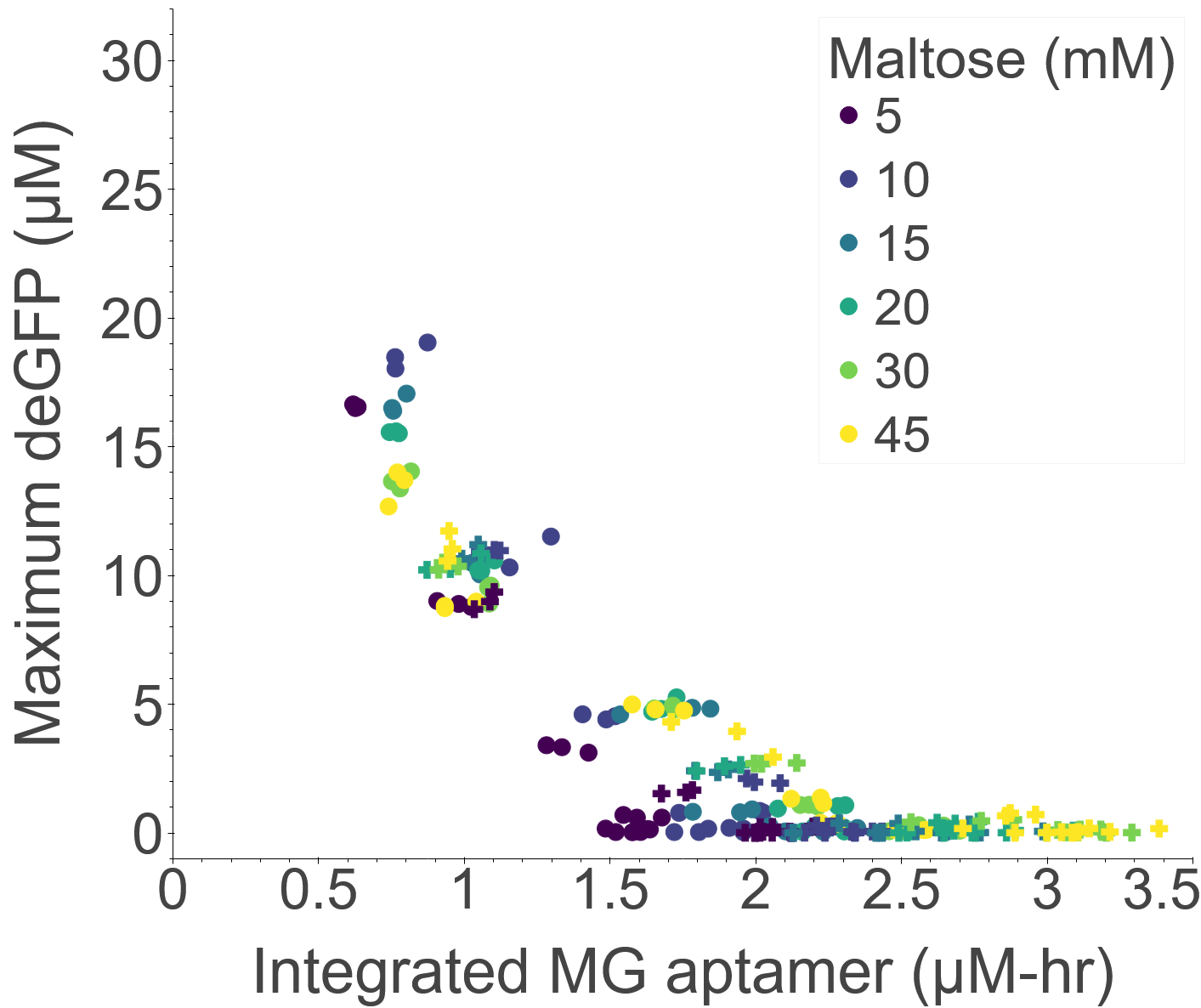

### Fig3_c.png

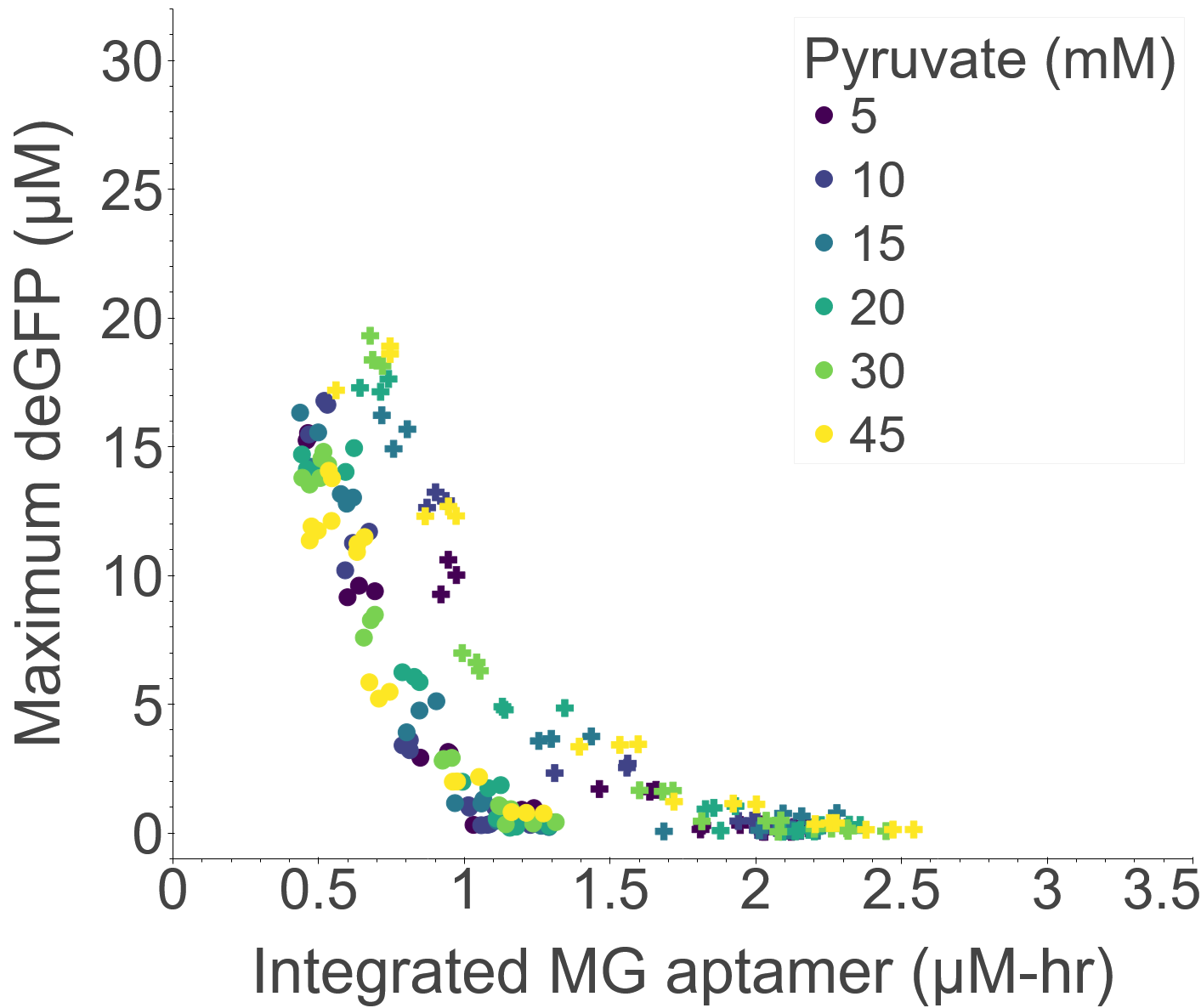

### Fig3_d.png

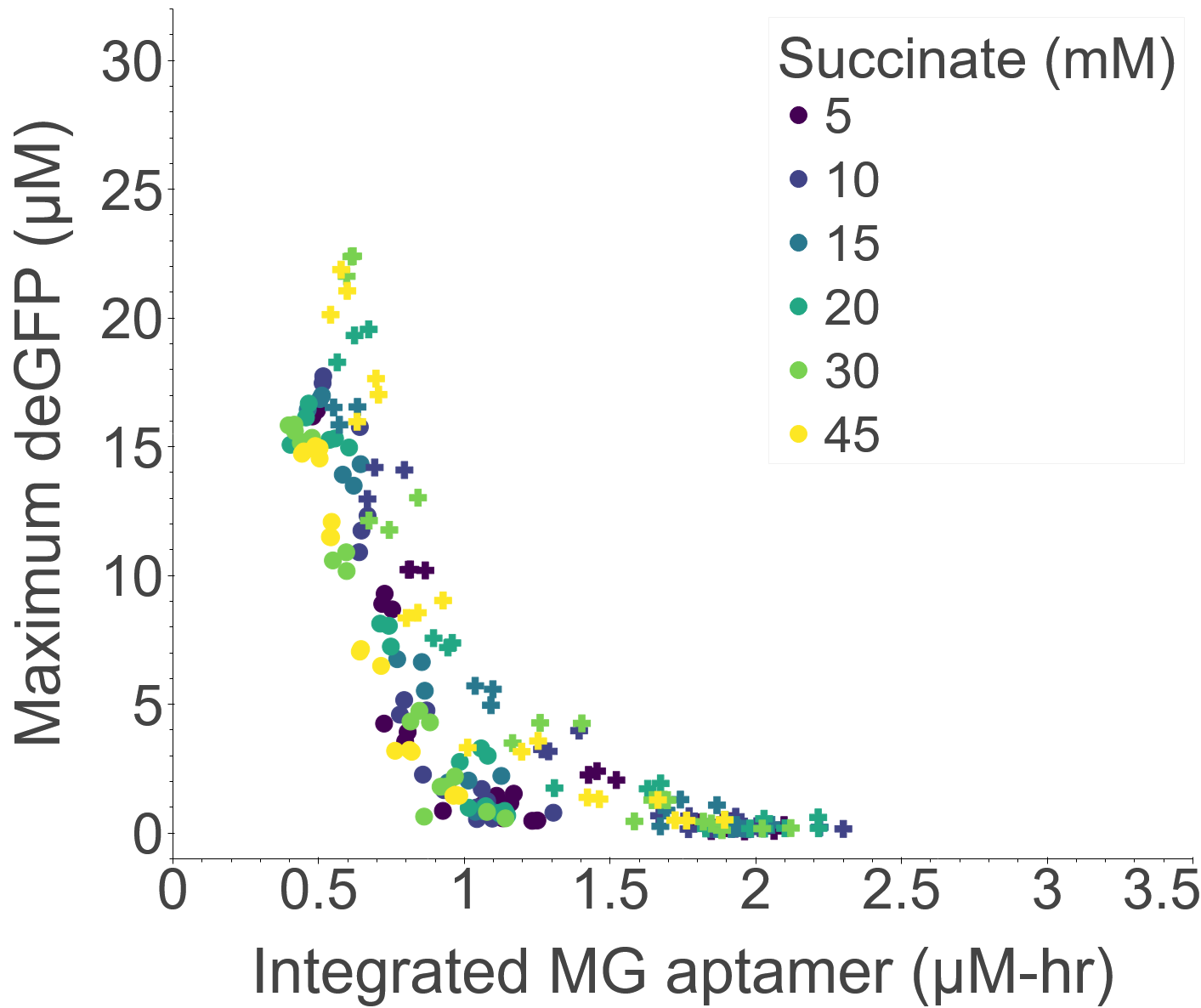

### Fig4.png

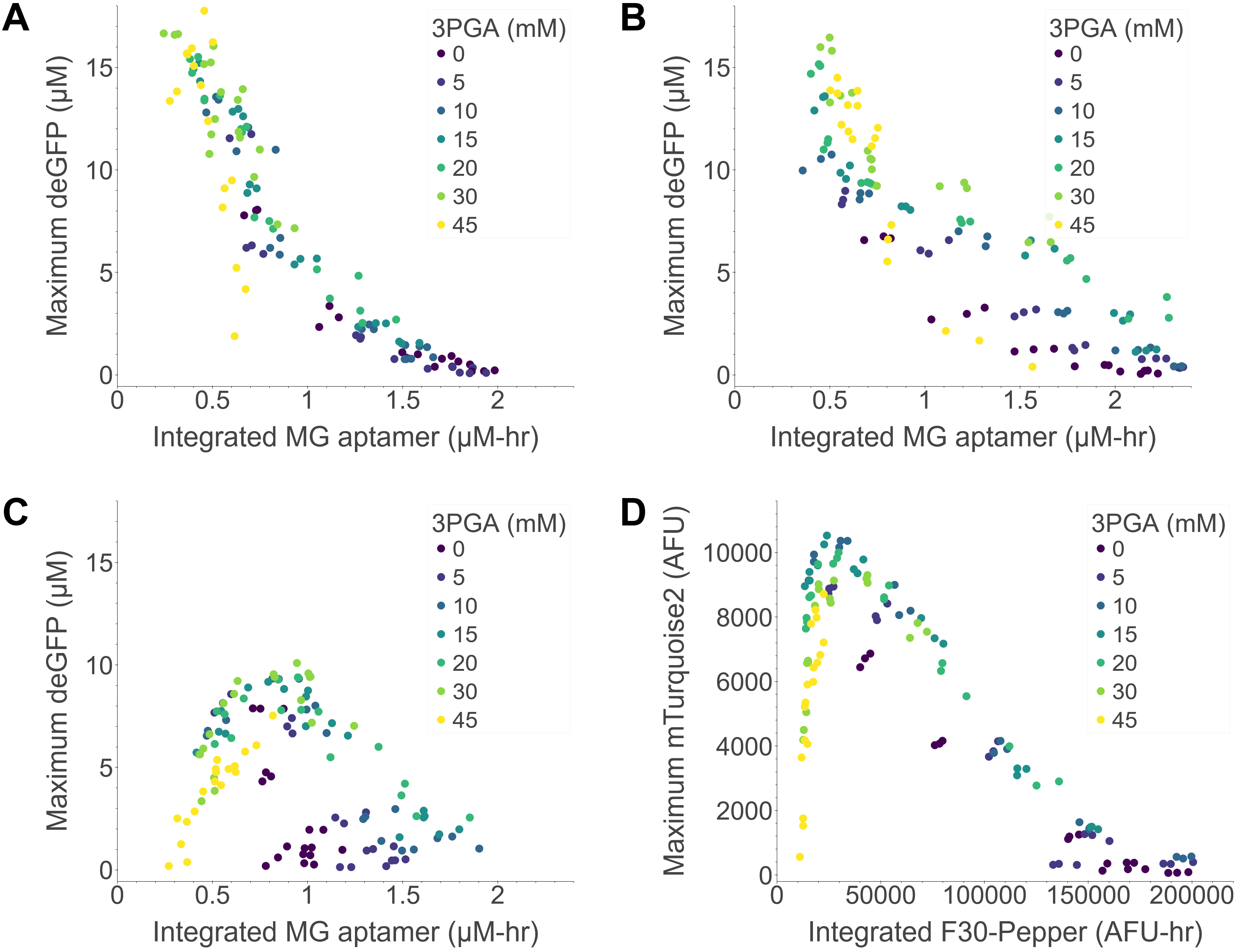

### Fig4_a.png

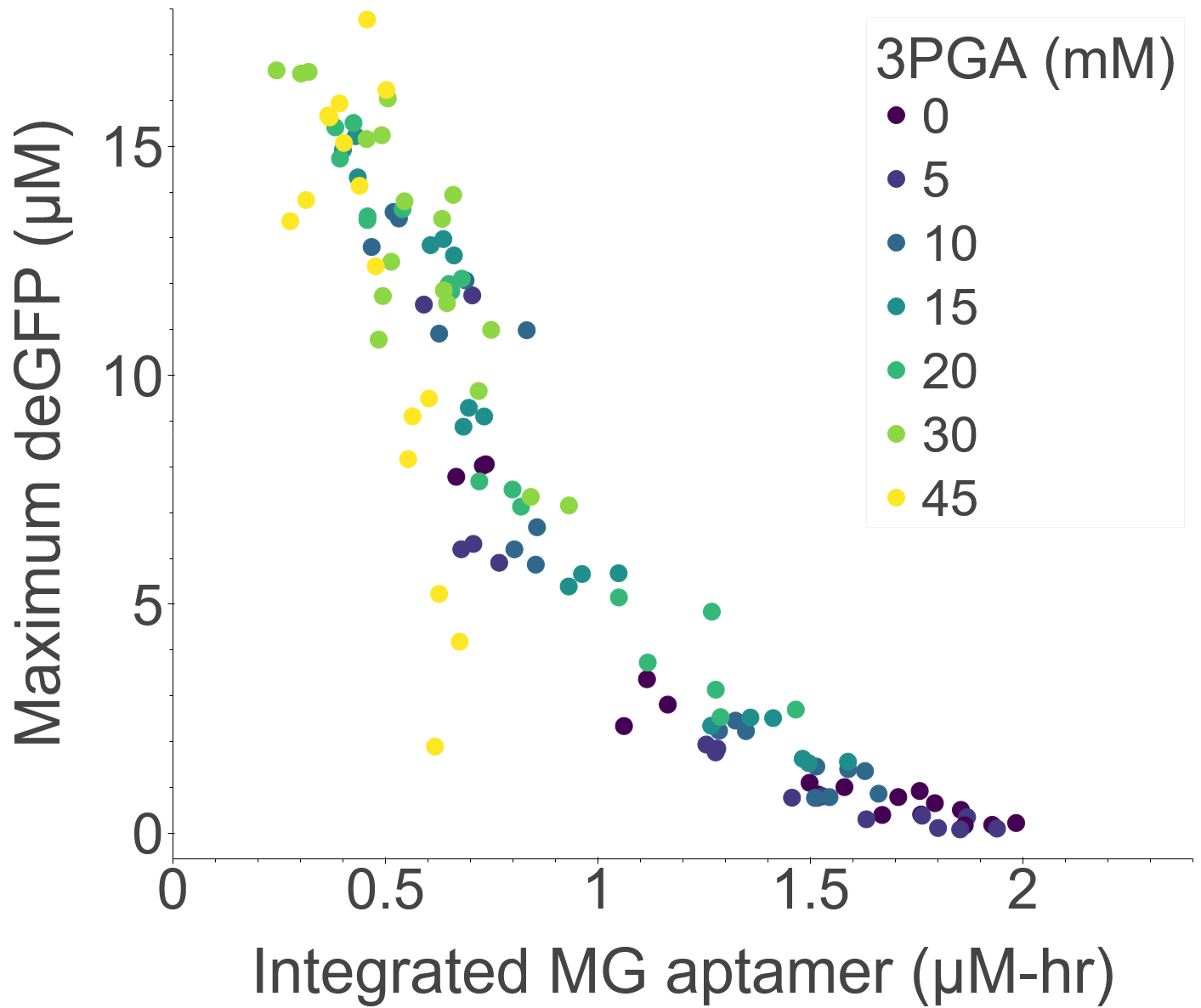

### Fig4_b.png

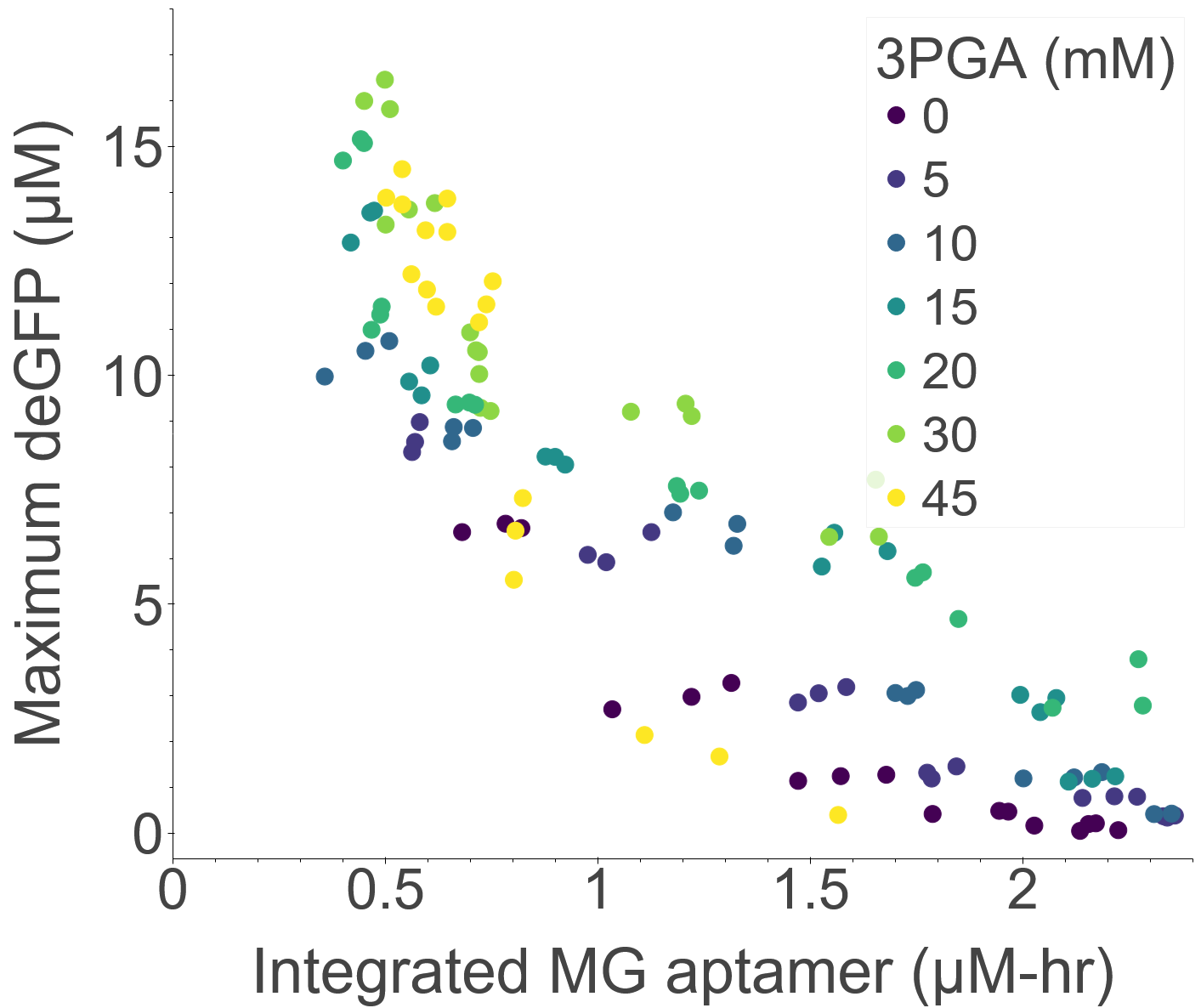

### Fig4_c.png

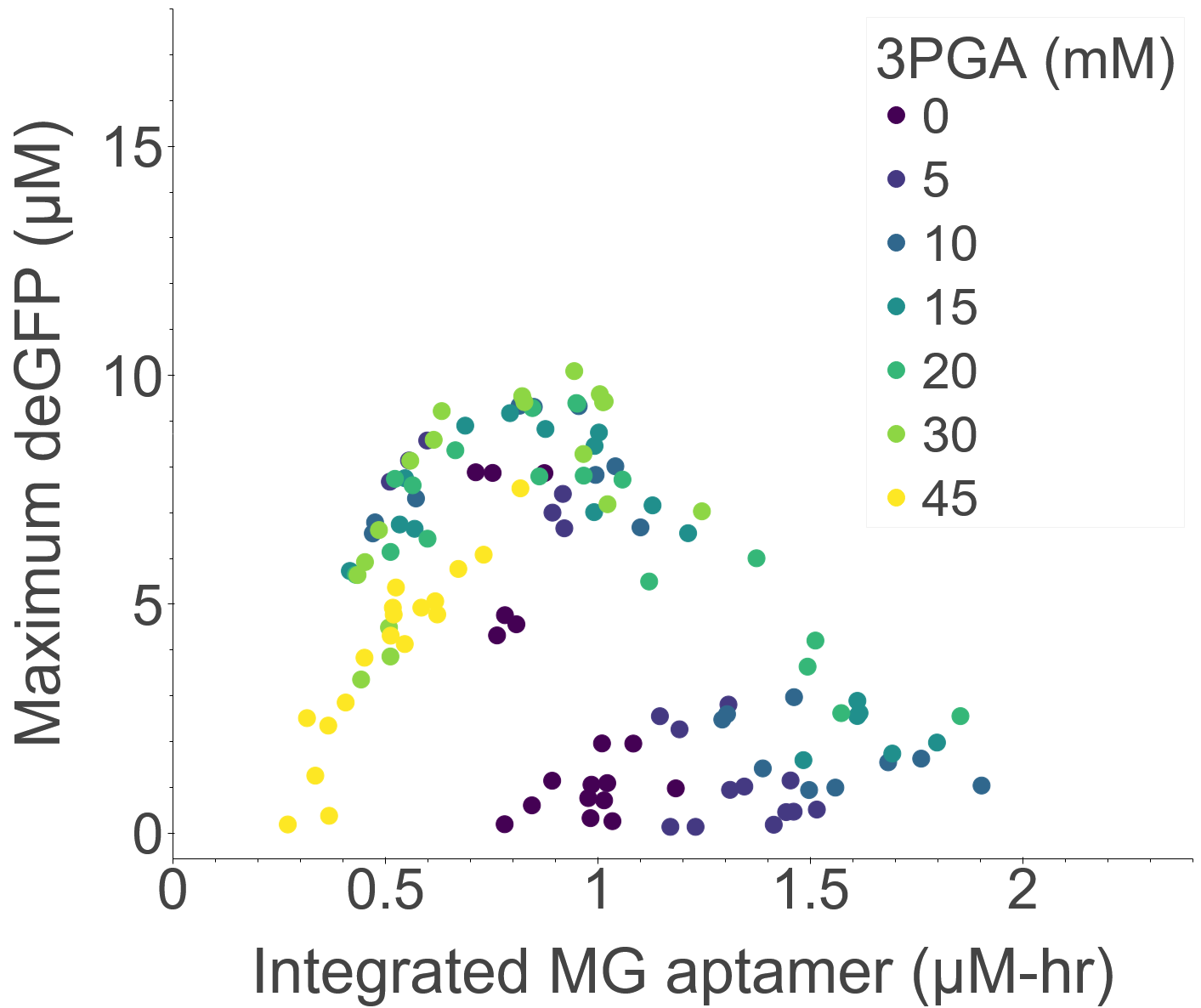

### Fig4_d.png

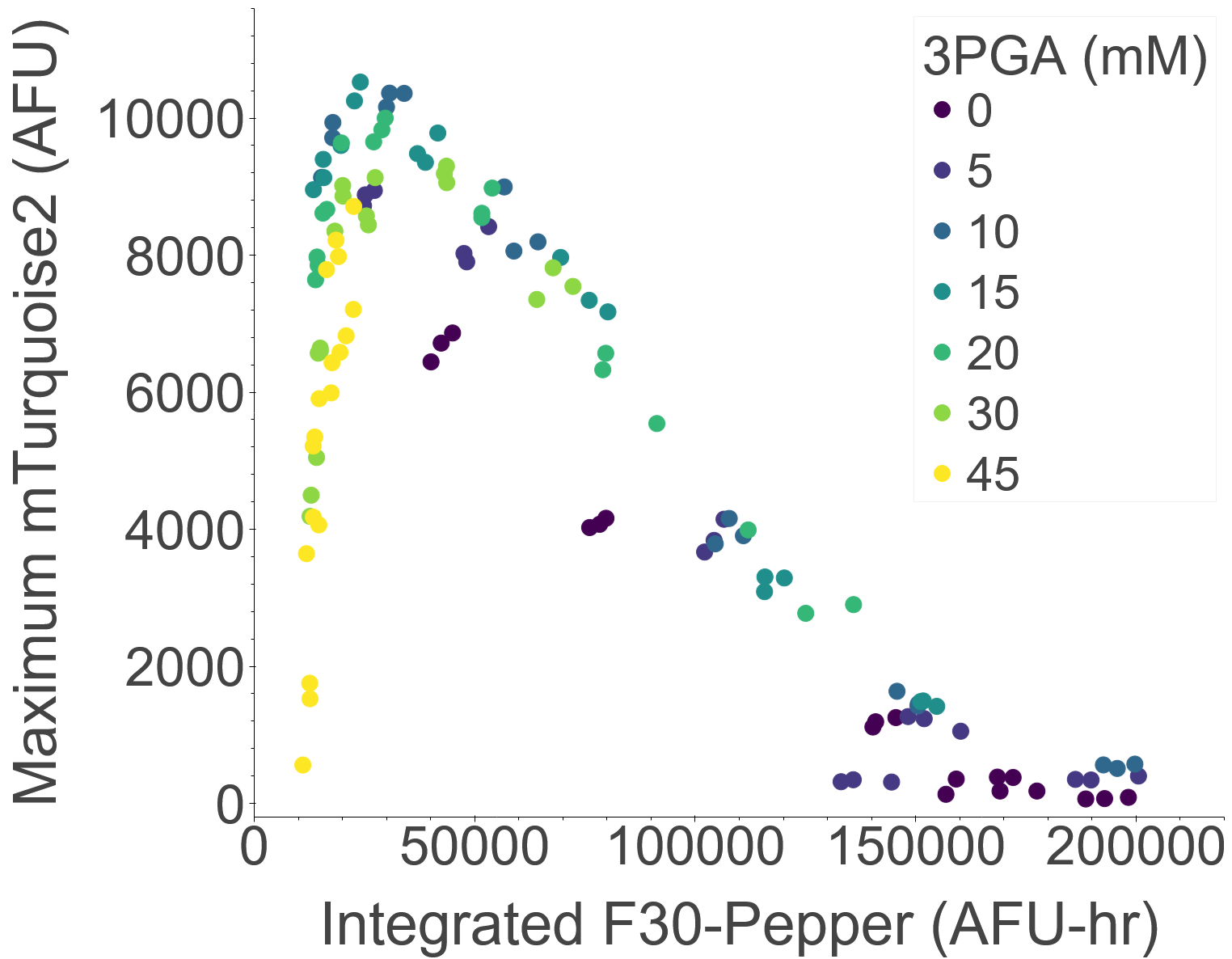

### Fig5.png

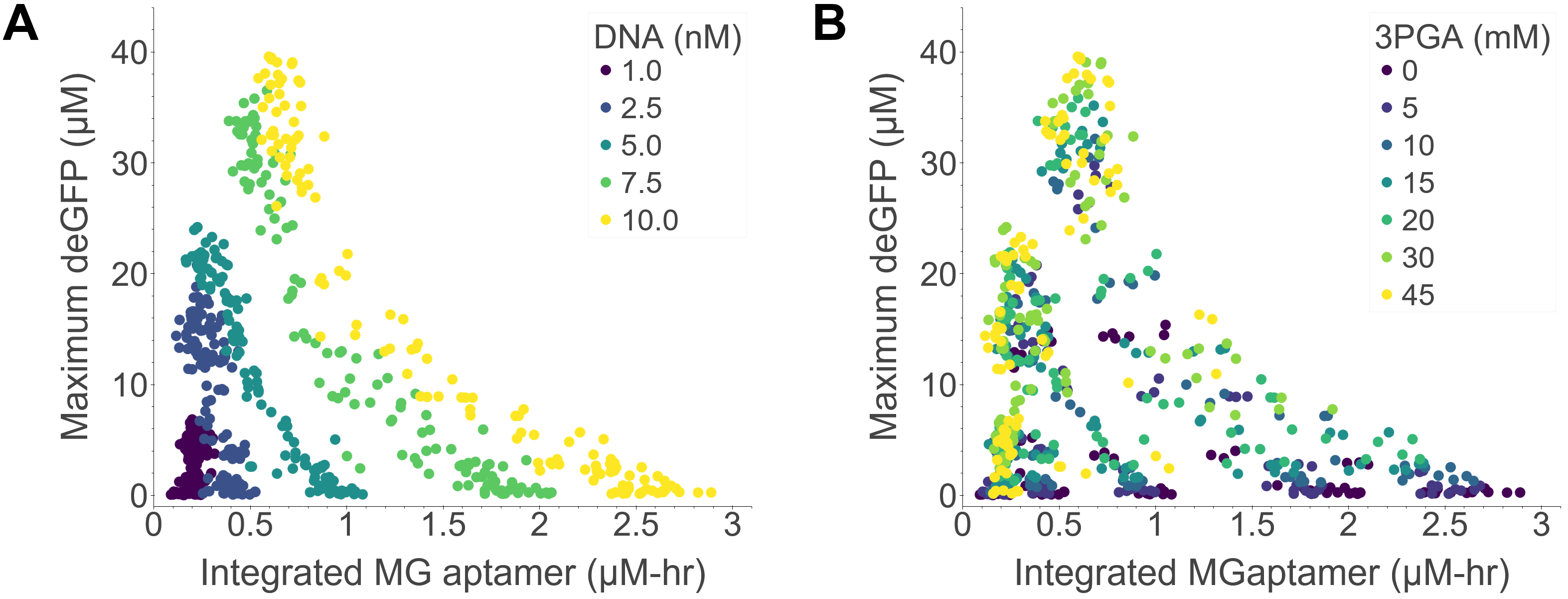

### Fig5_a.png

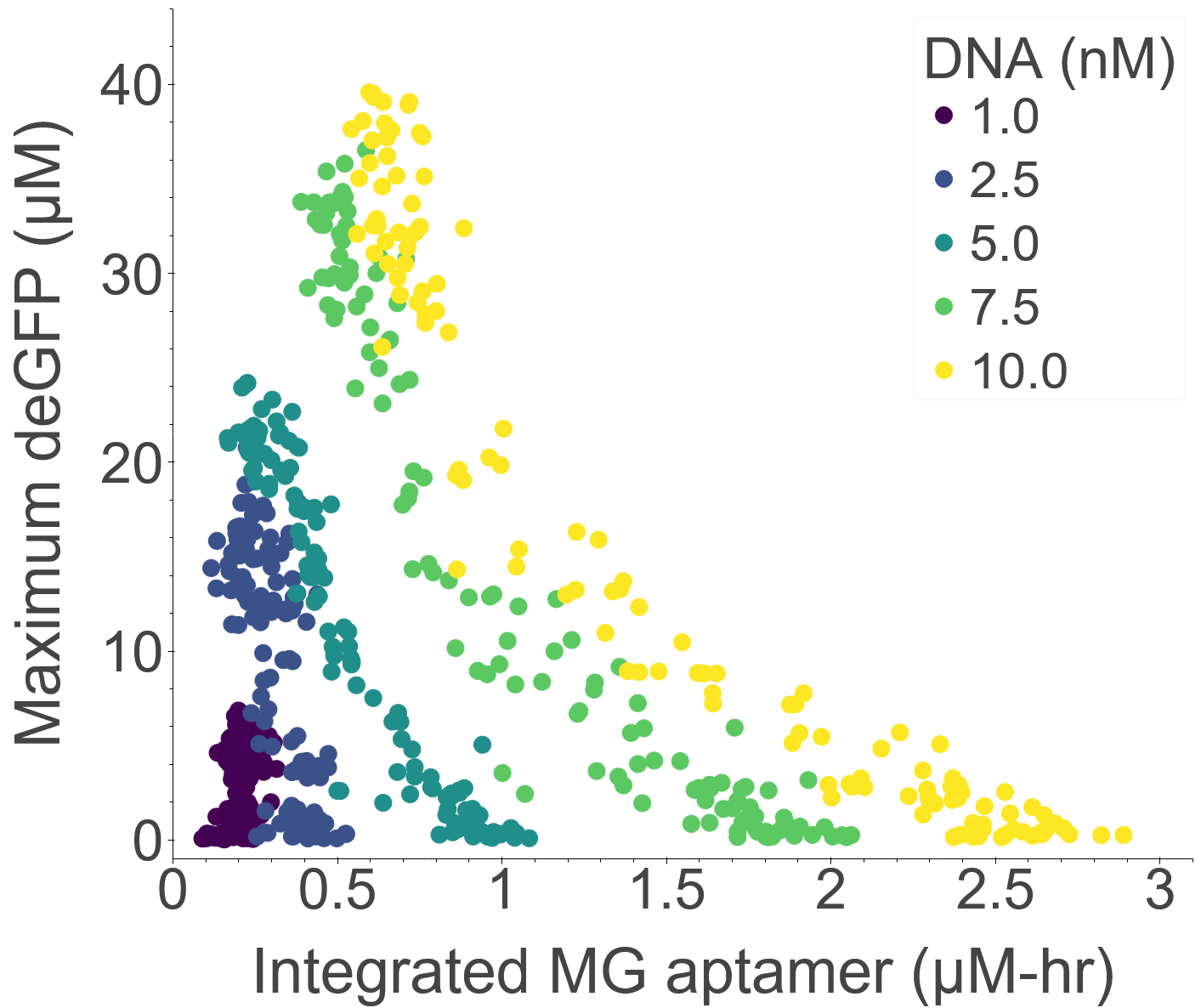

### Fig5_b.png

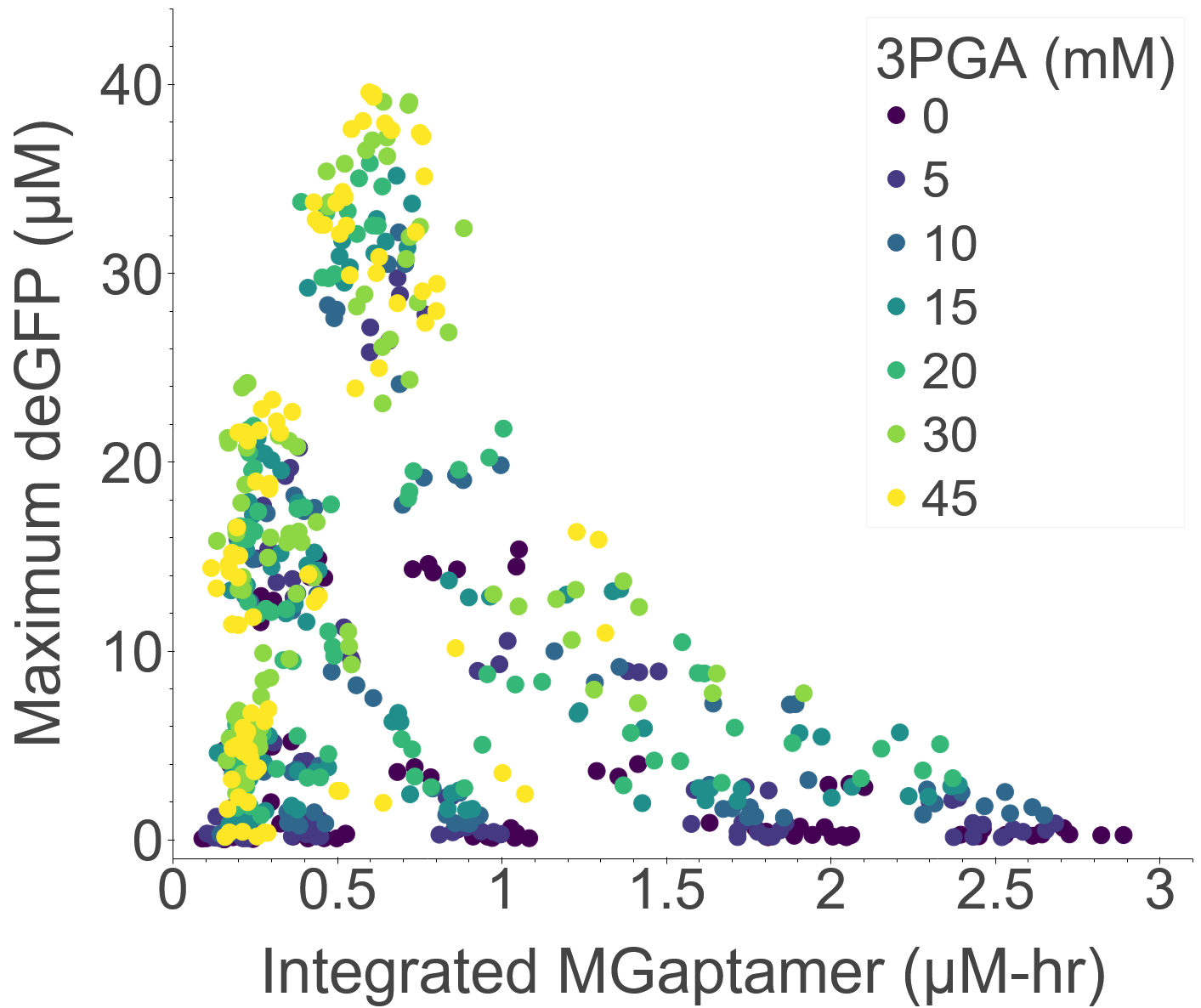

### Fig6.png

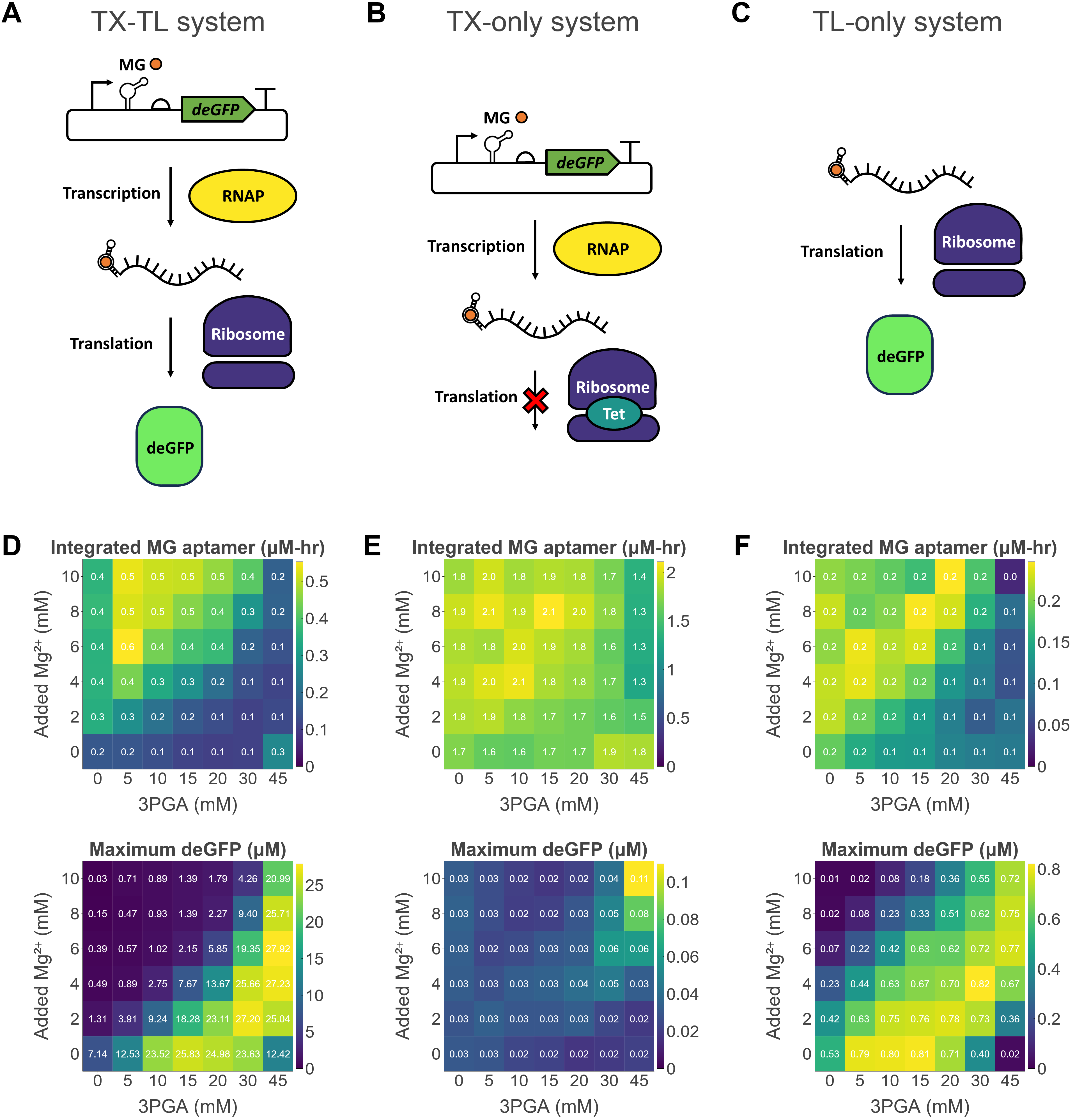

### Fig6_a.png

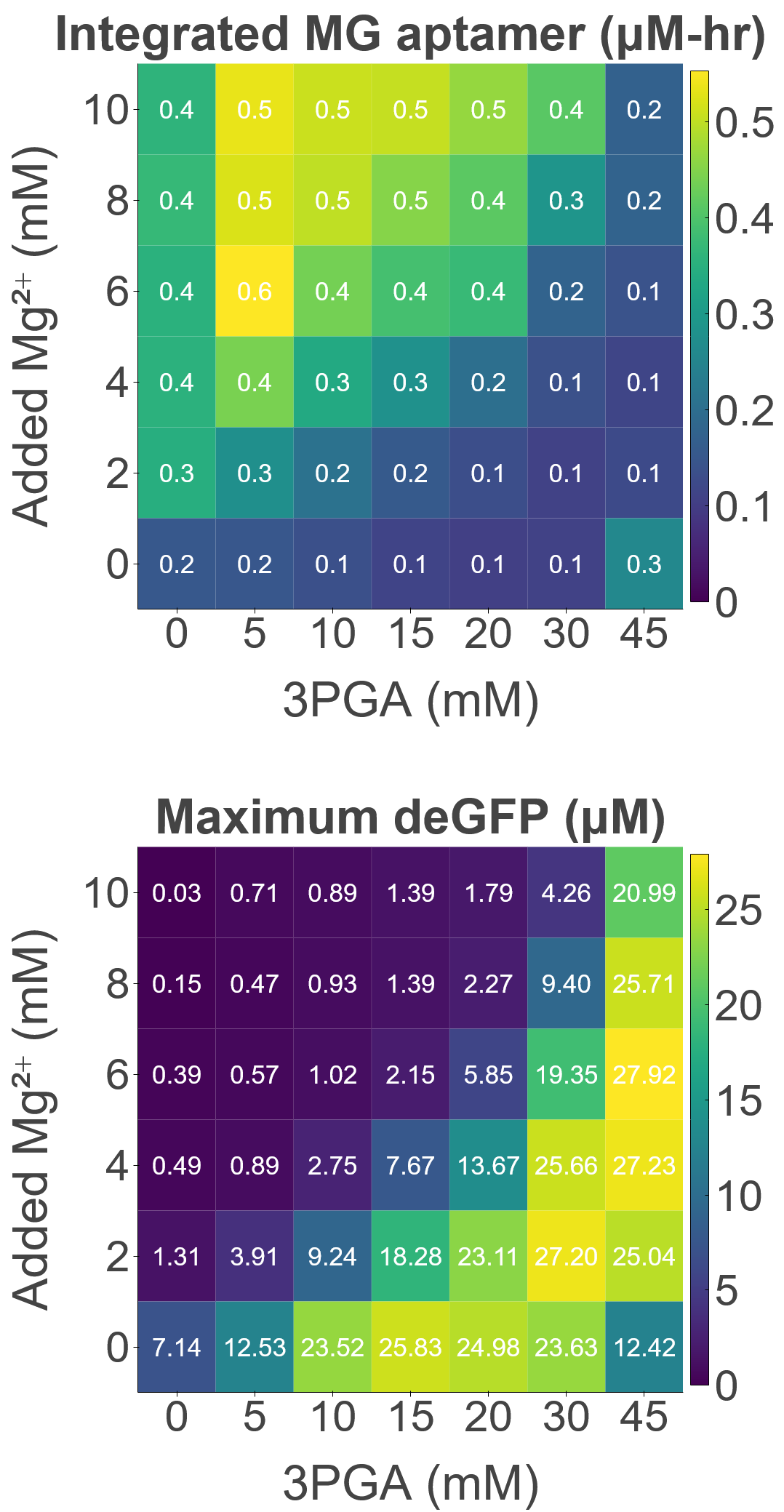

### Fig6_b.png

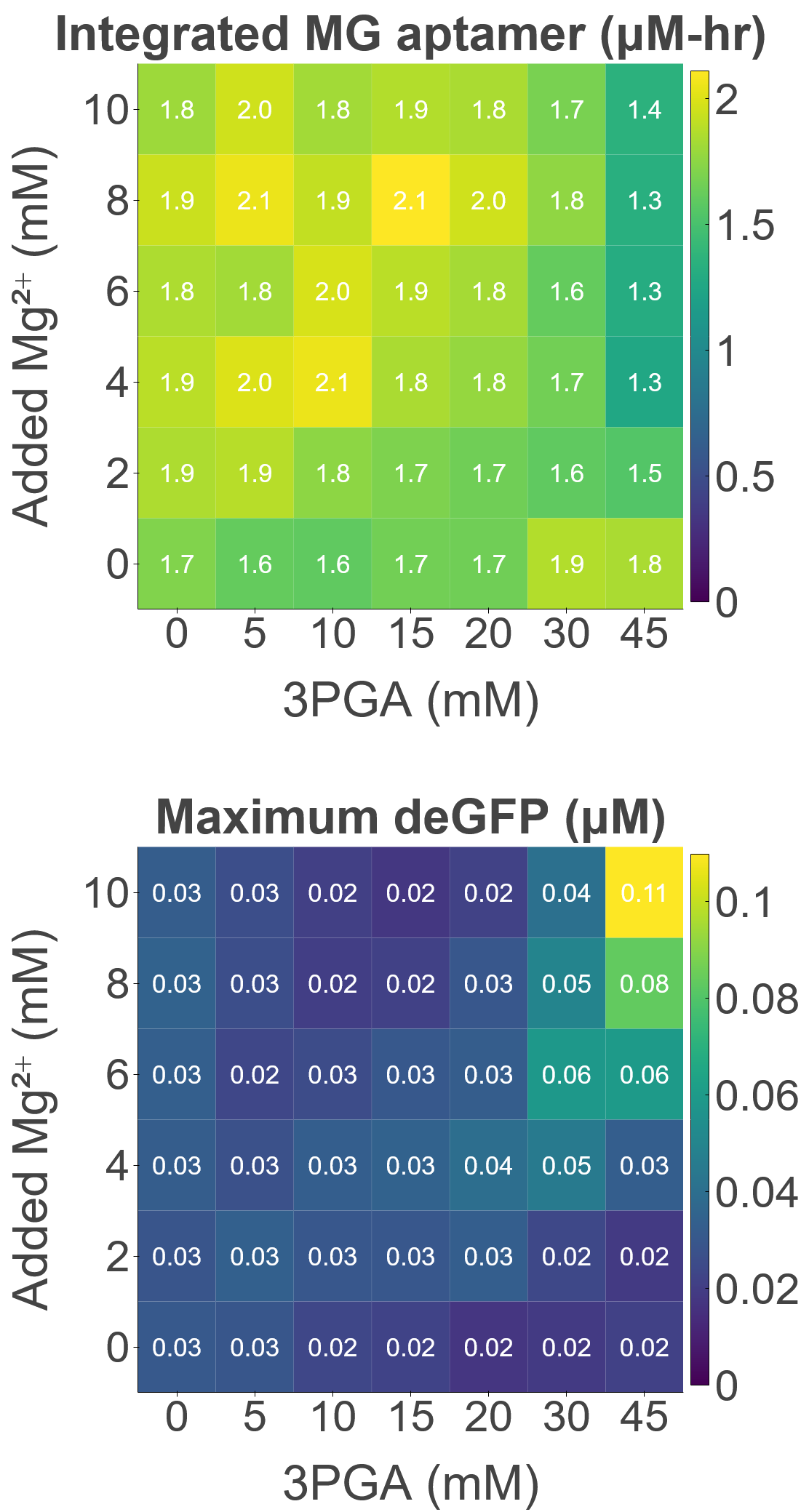

### Fig6_c.png

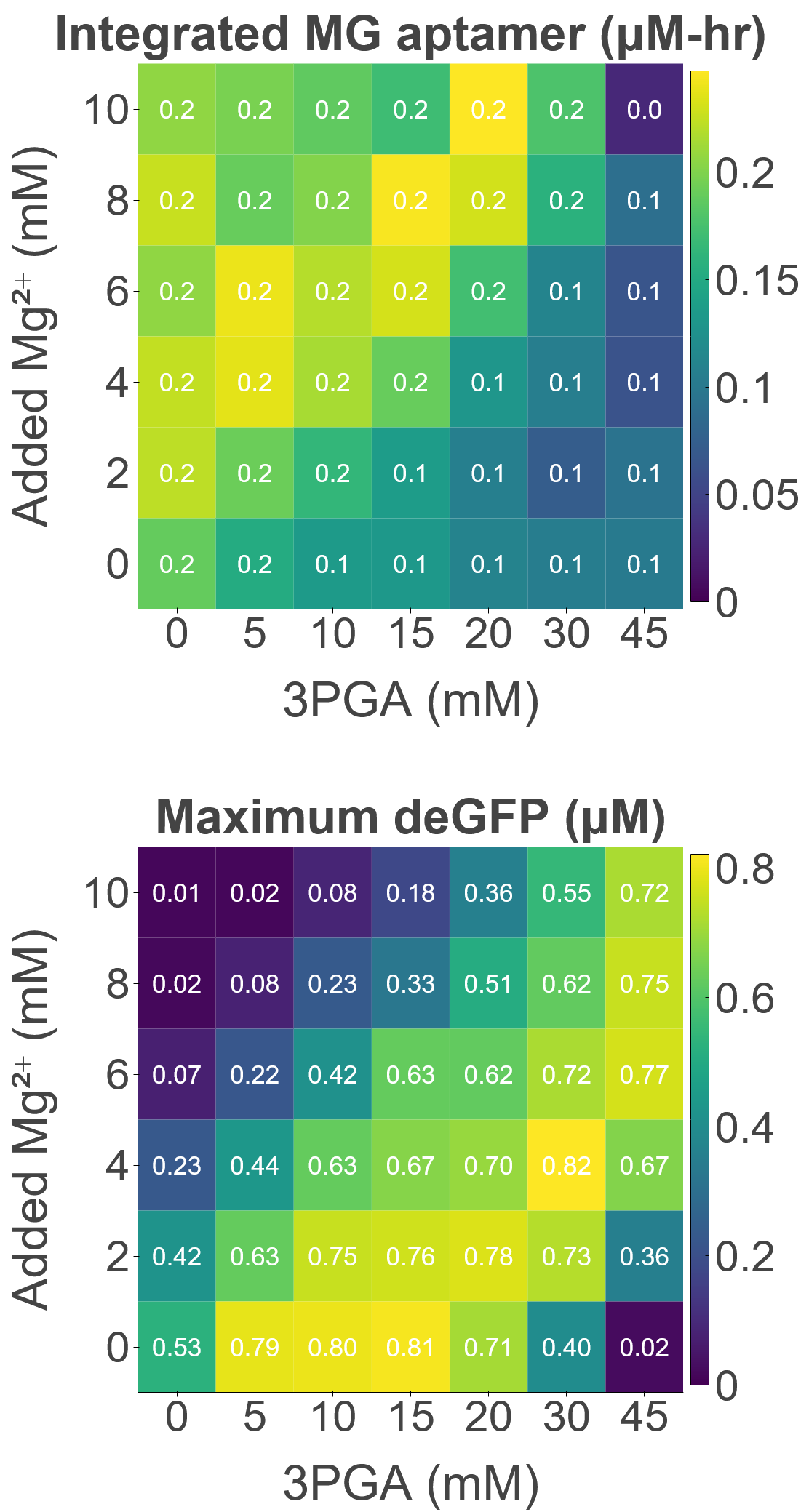

### Fig6_d.png

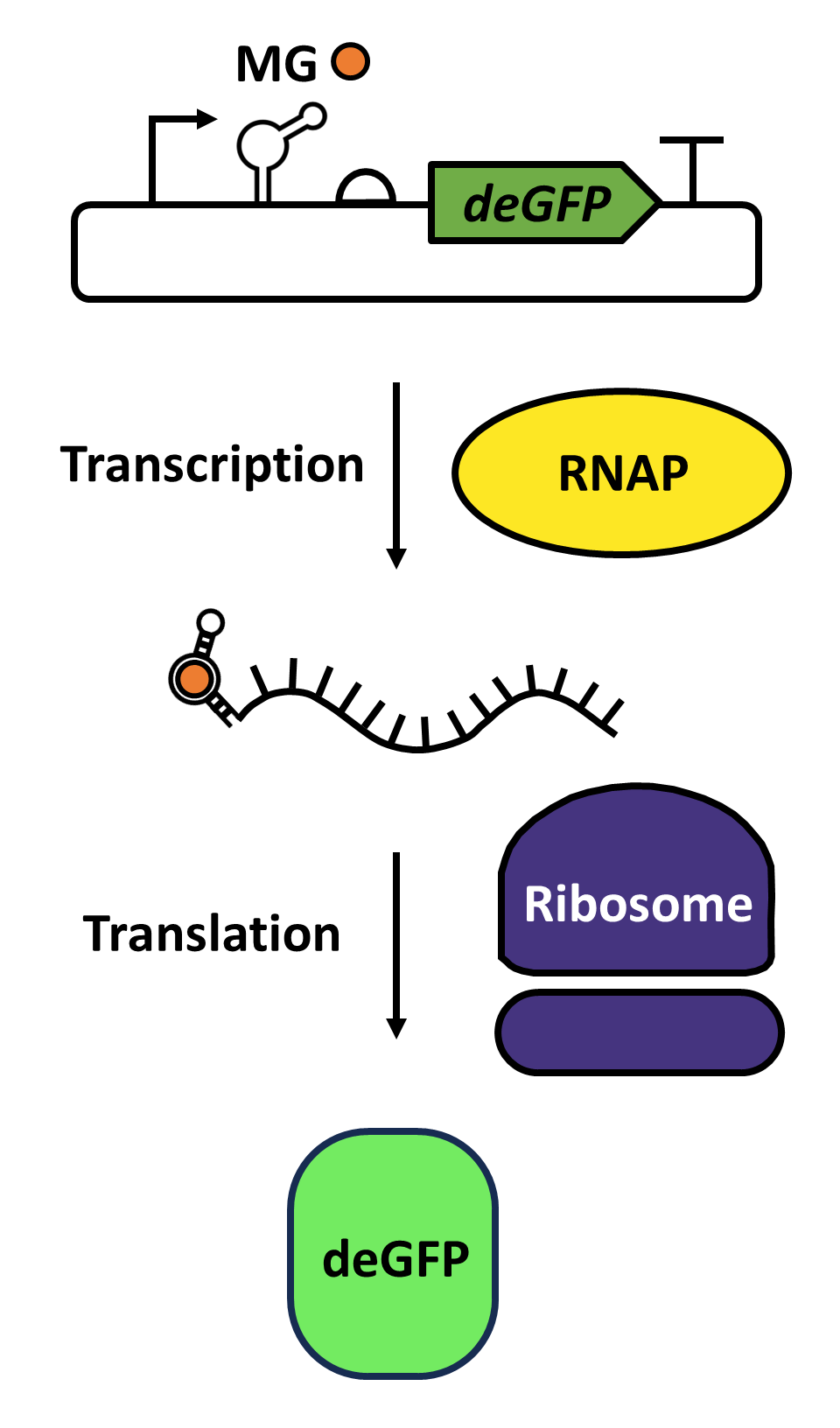

### Fig6_e.png

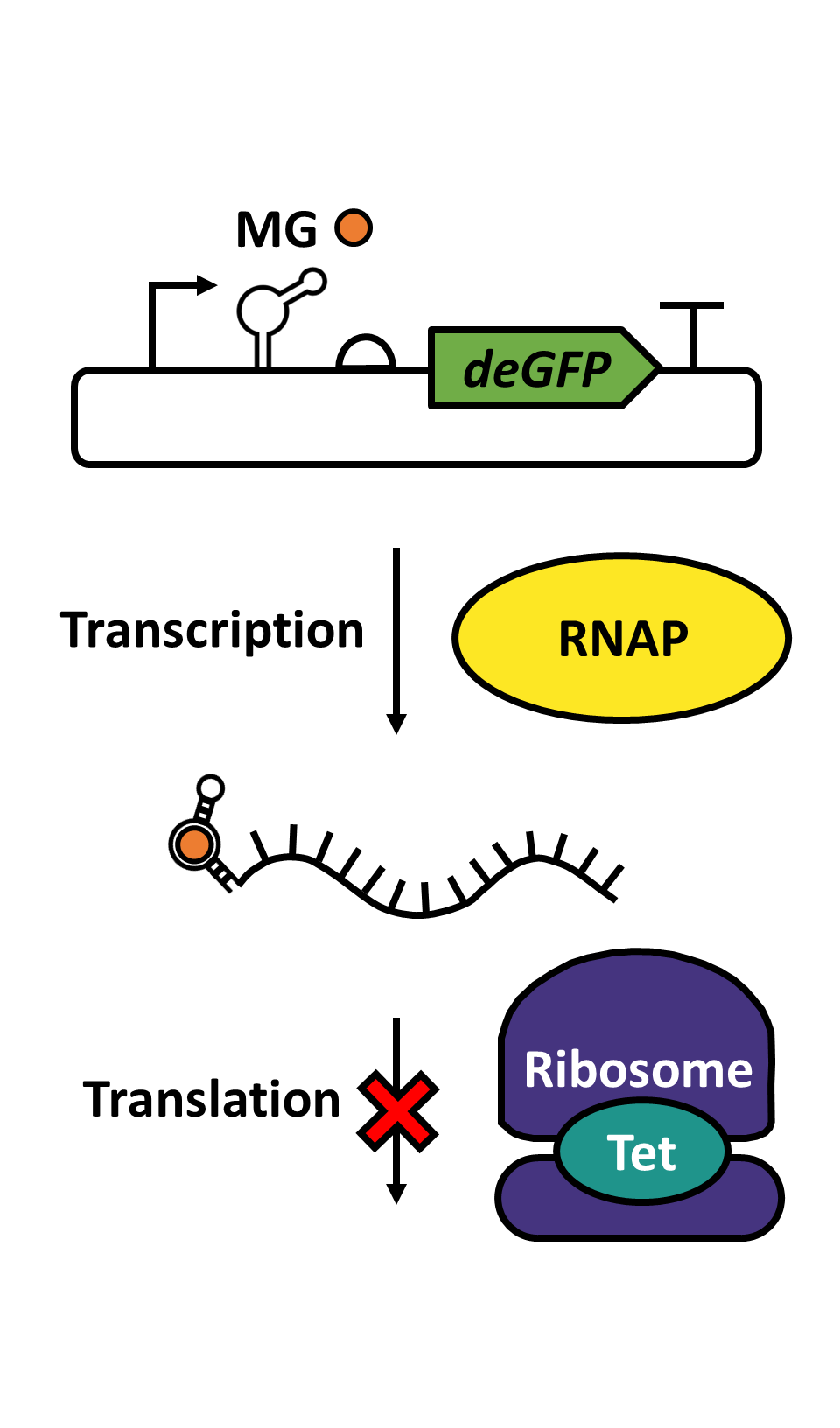

### Fig6_f.png

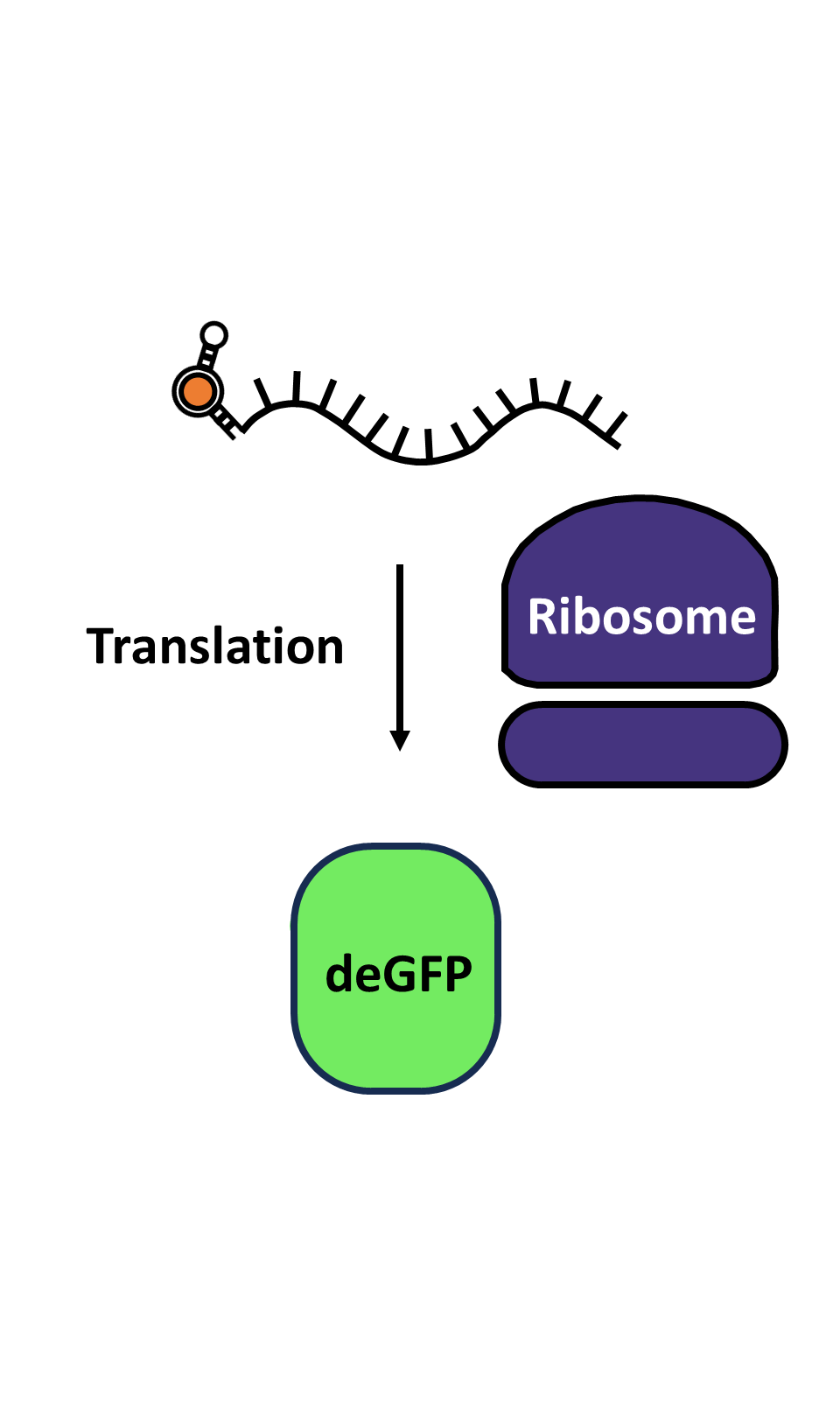

### Fig7.png

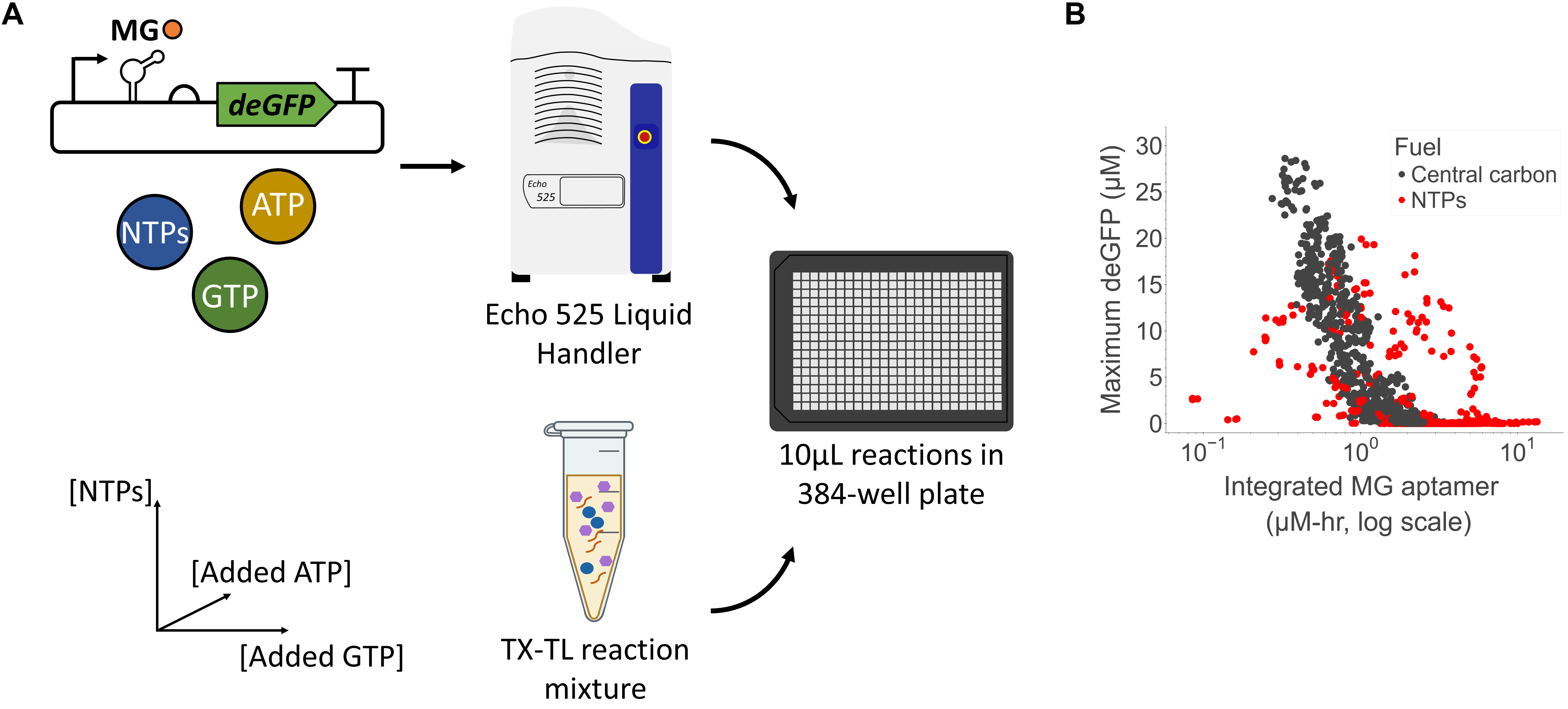

### Fig7_a.png

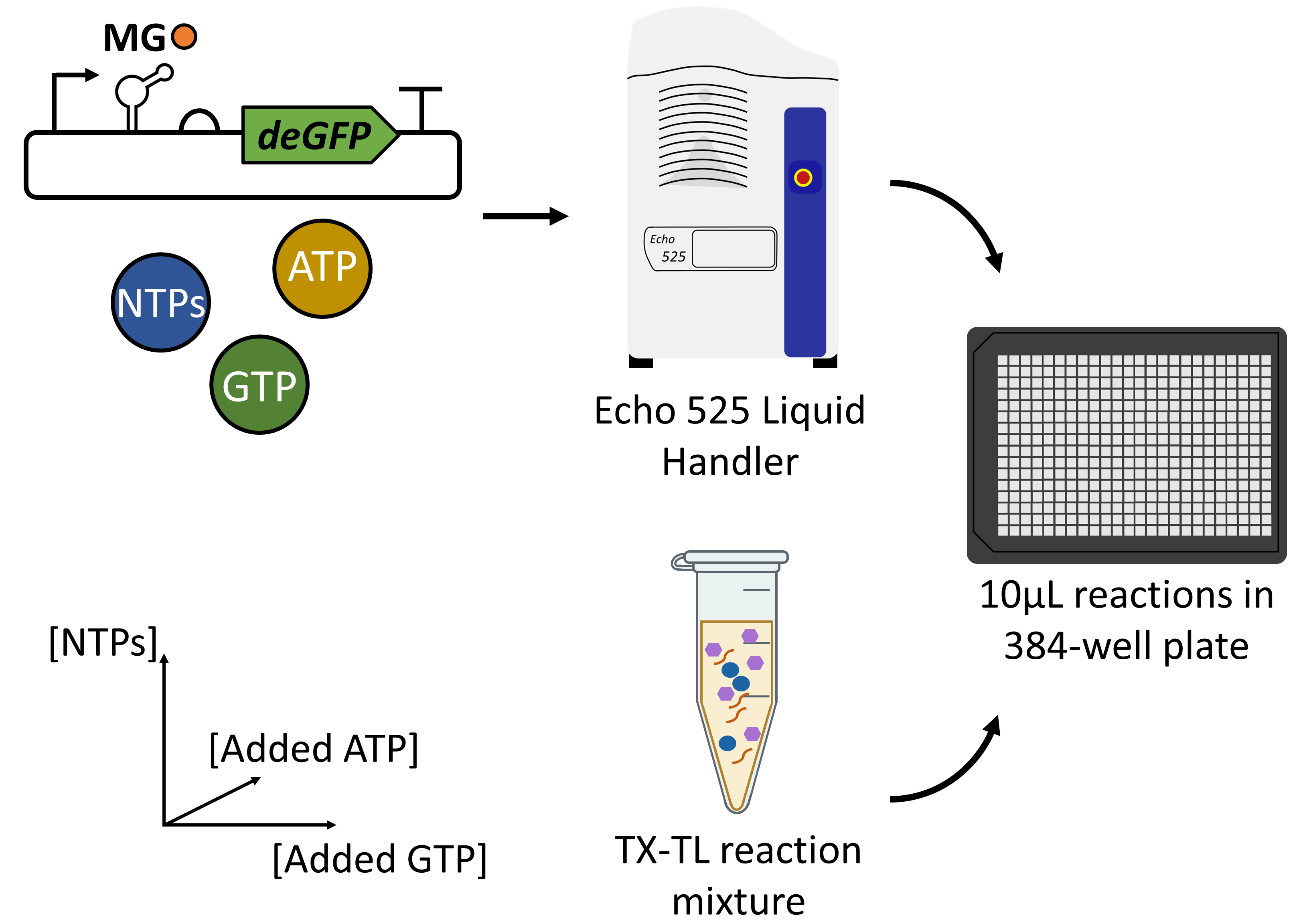

### Fig7_b.png

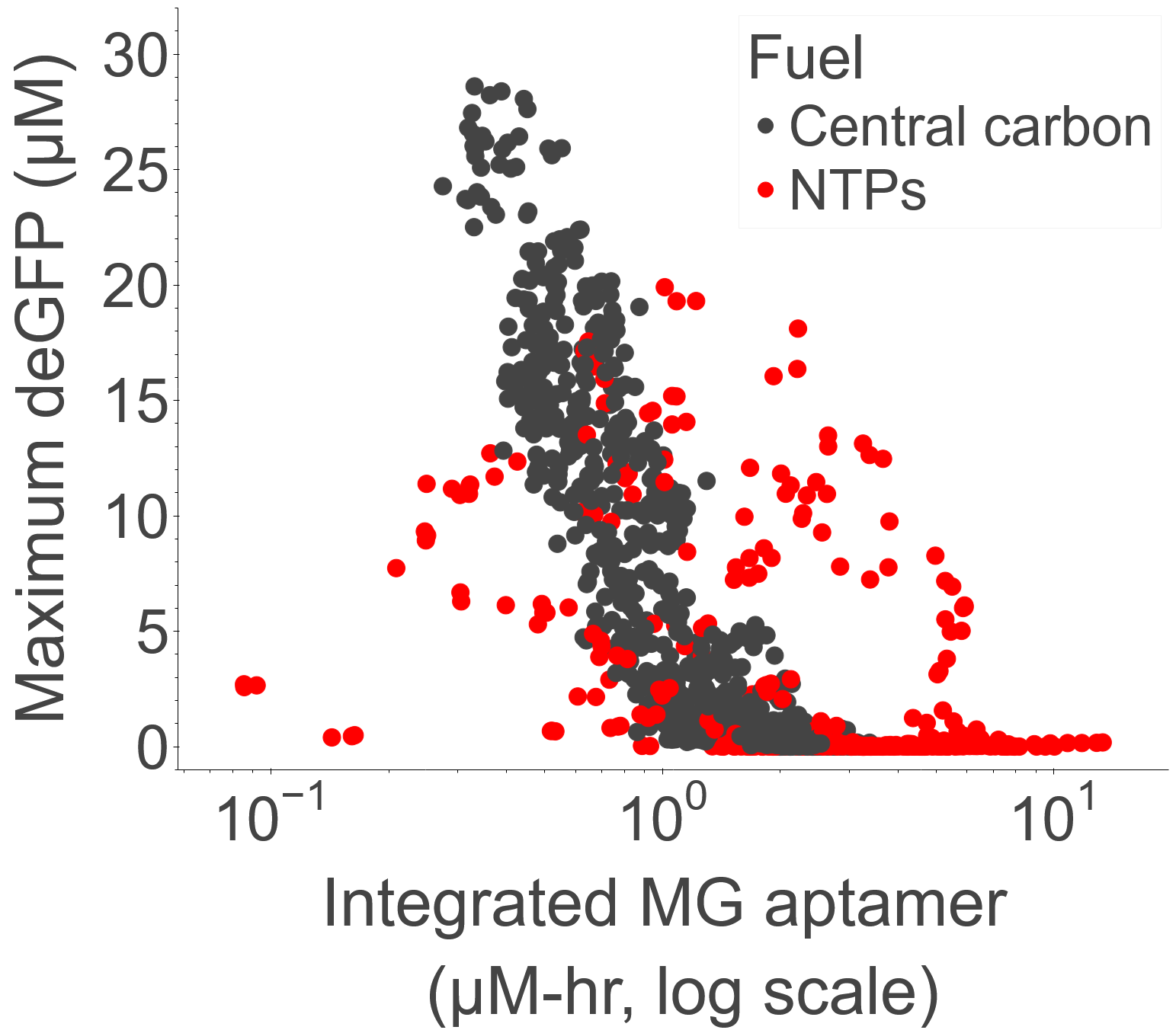

### Fig8.png

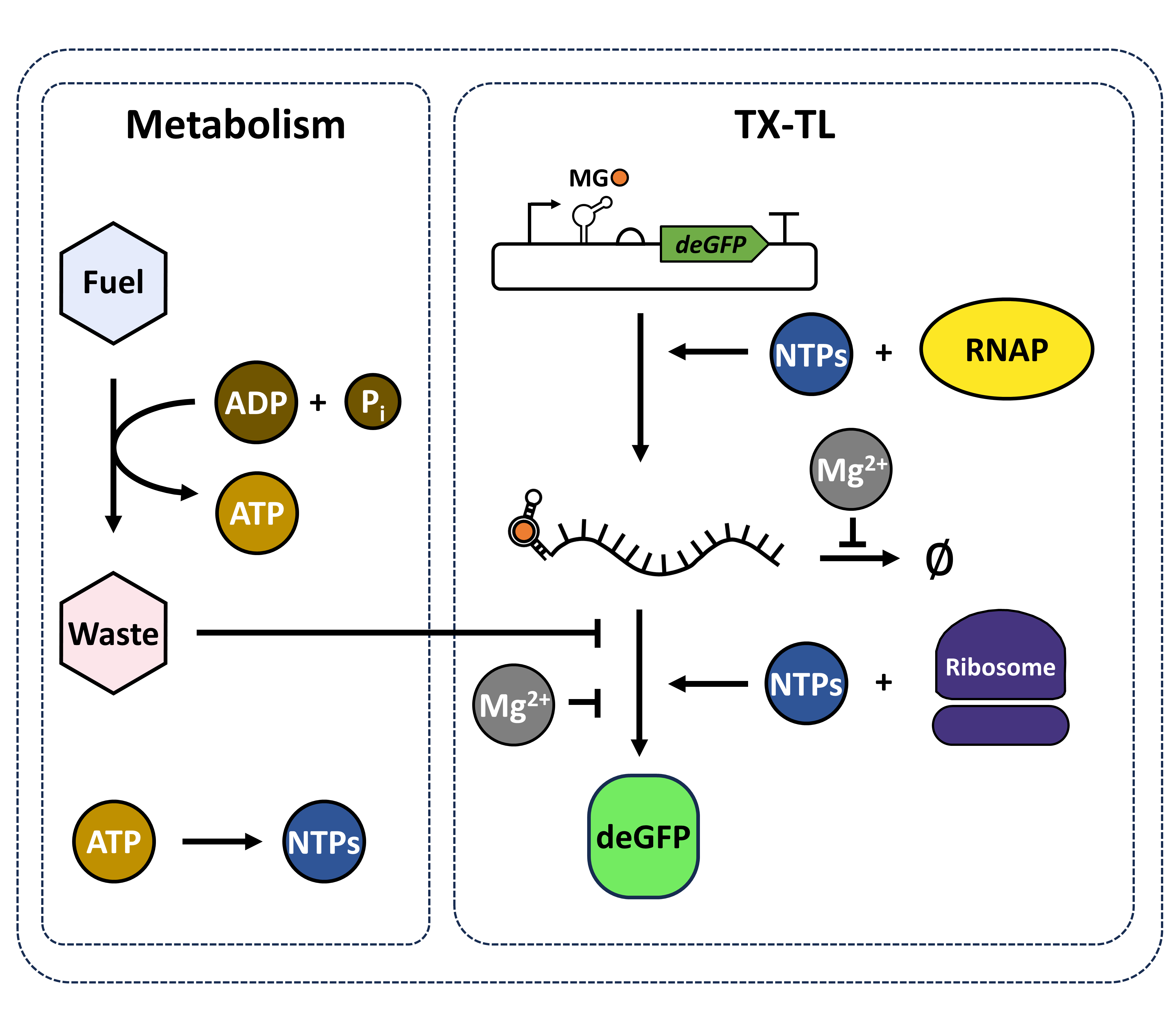

### Fig8.pptx

#### Slide 1
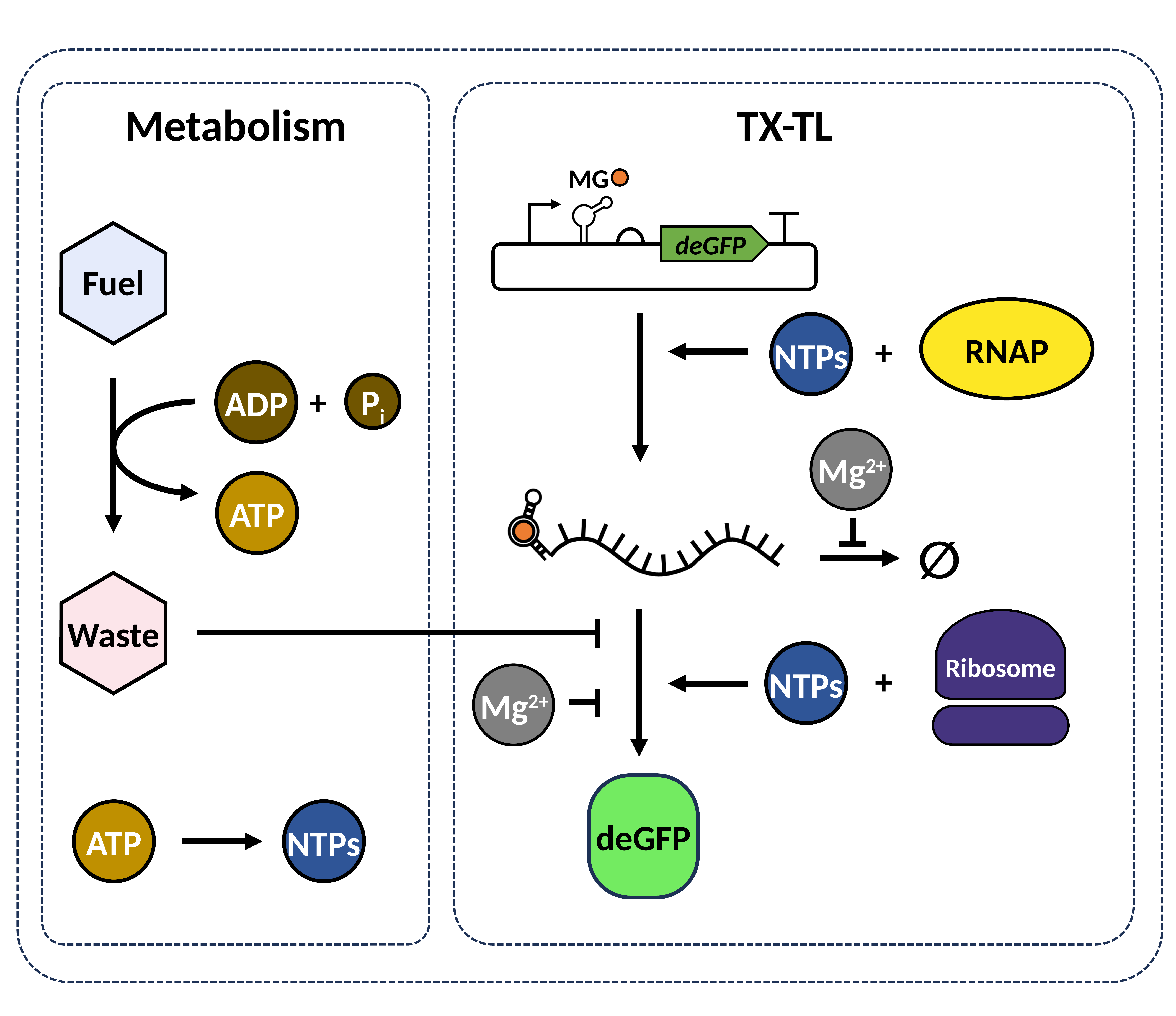

Metabolism
TX-TL
MG
deGFP
*
Fuel
RNAP
+
NTPs
+
P_i_
ADP
Mg^2+^
ATP
∅
*
Waste
Ribosome
+
NTPs
Mg^2+^
deGFP
ATP
NTPs

### FigS1.png

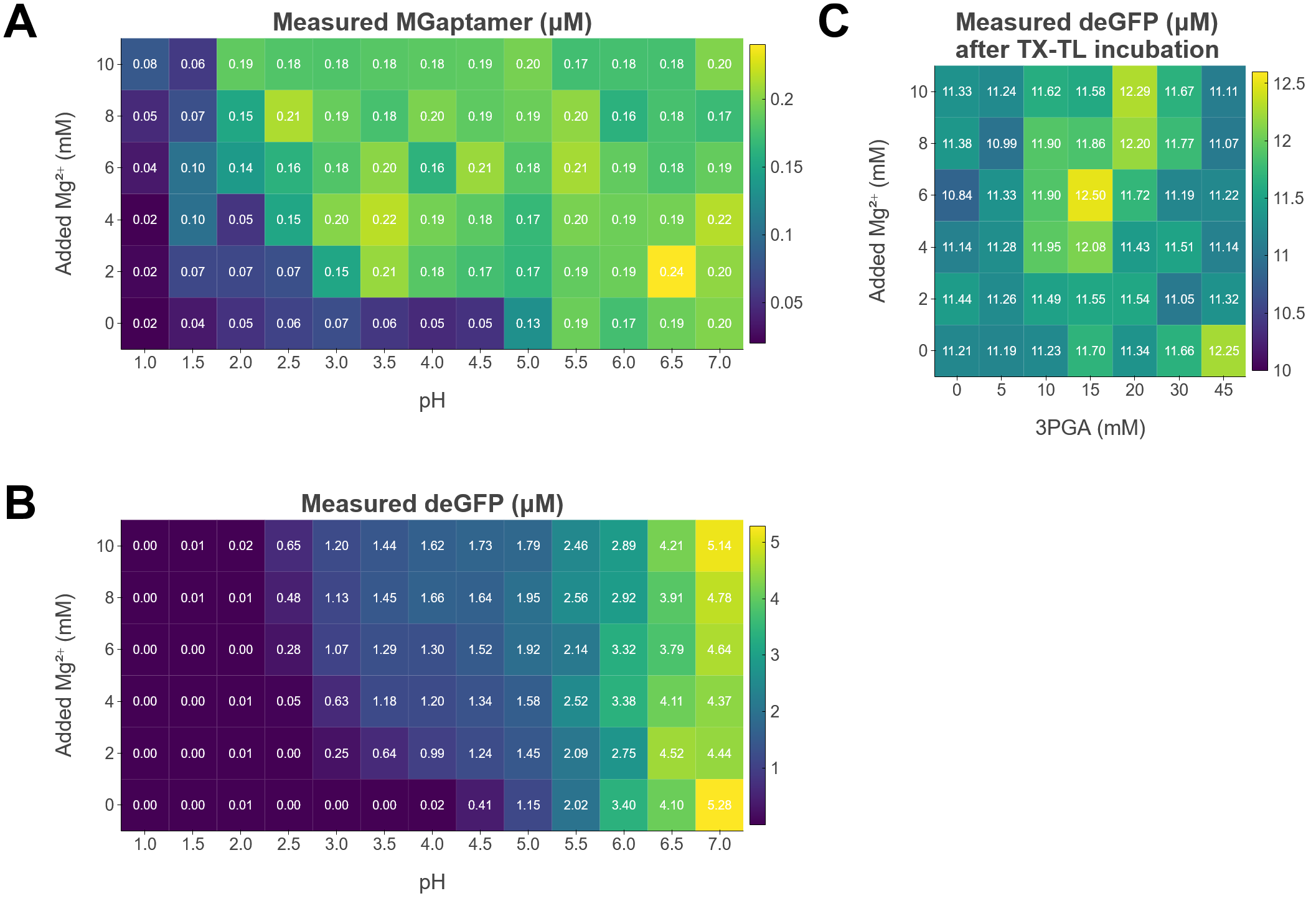

### FigS1_a.png

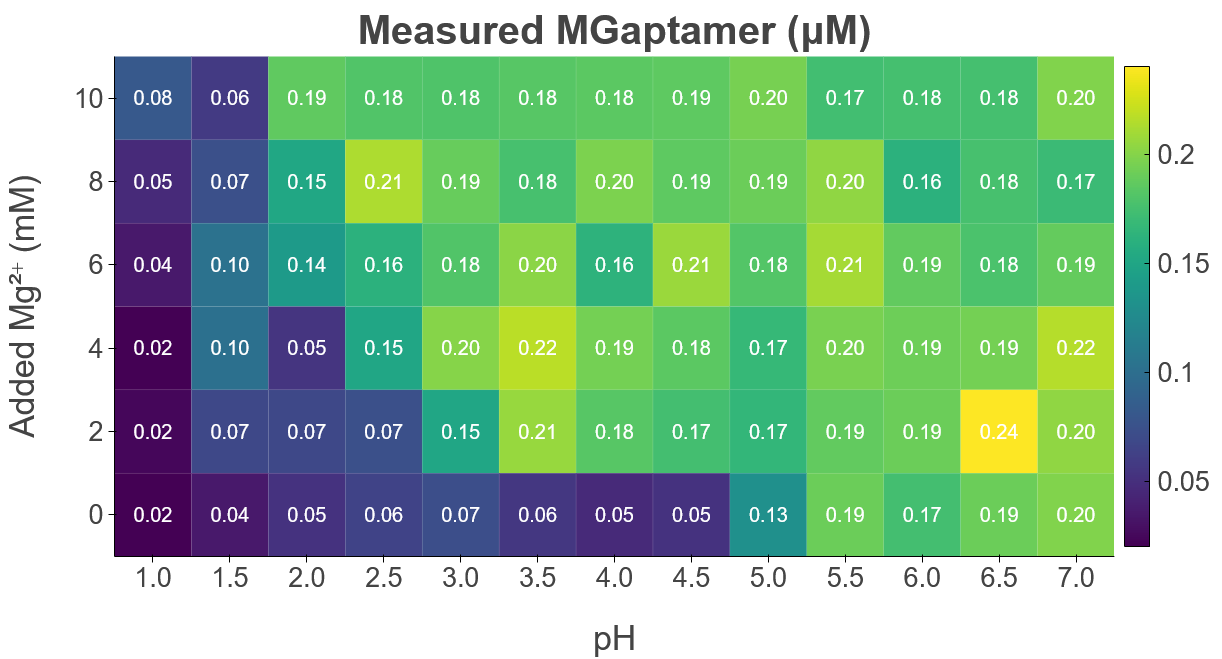

### FigS1_b.png

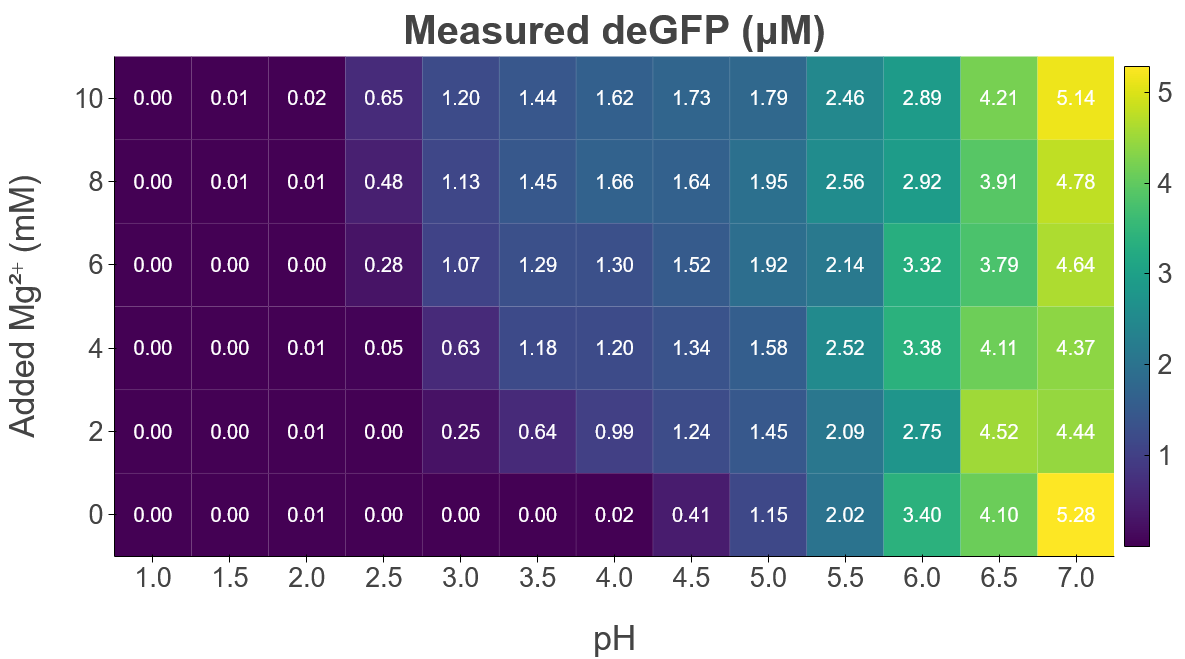

### FigS1_c.png

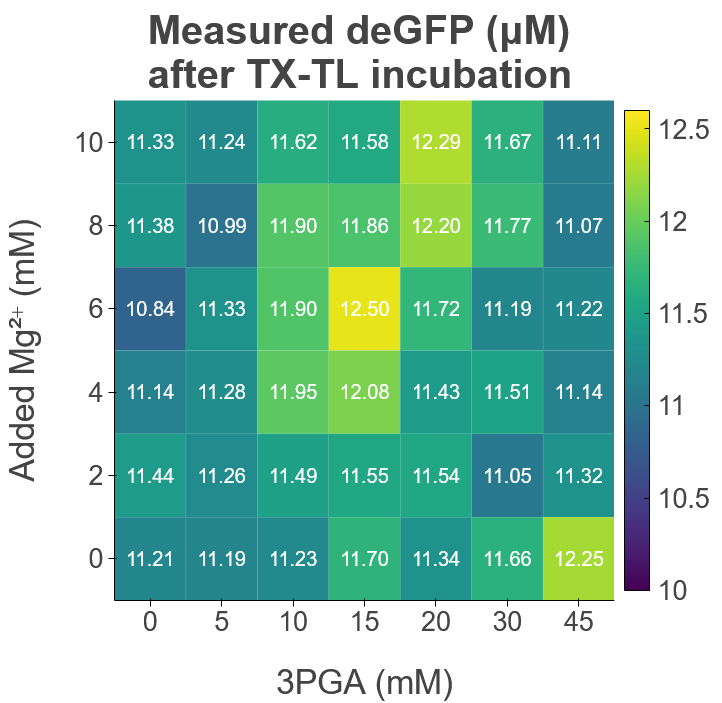

### FigS2.png

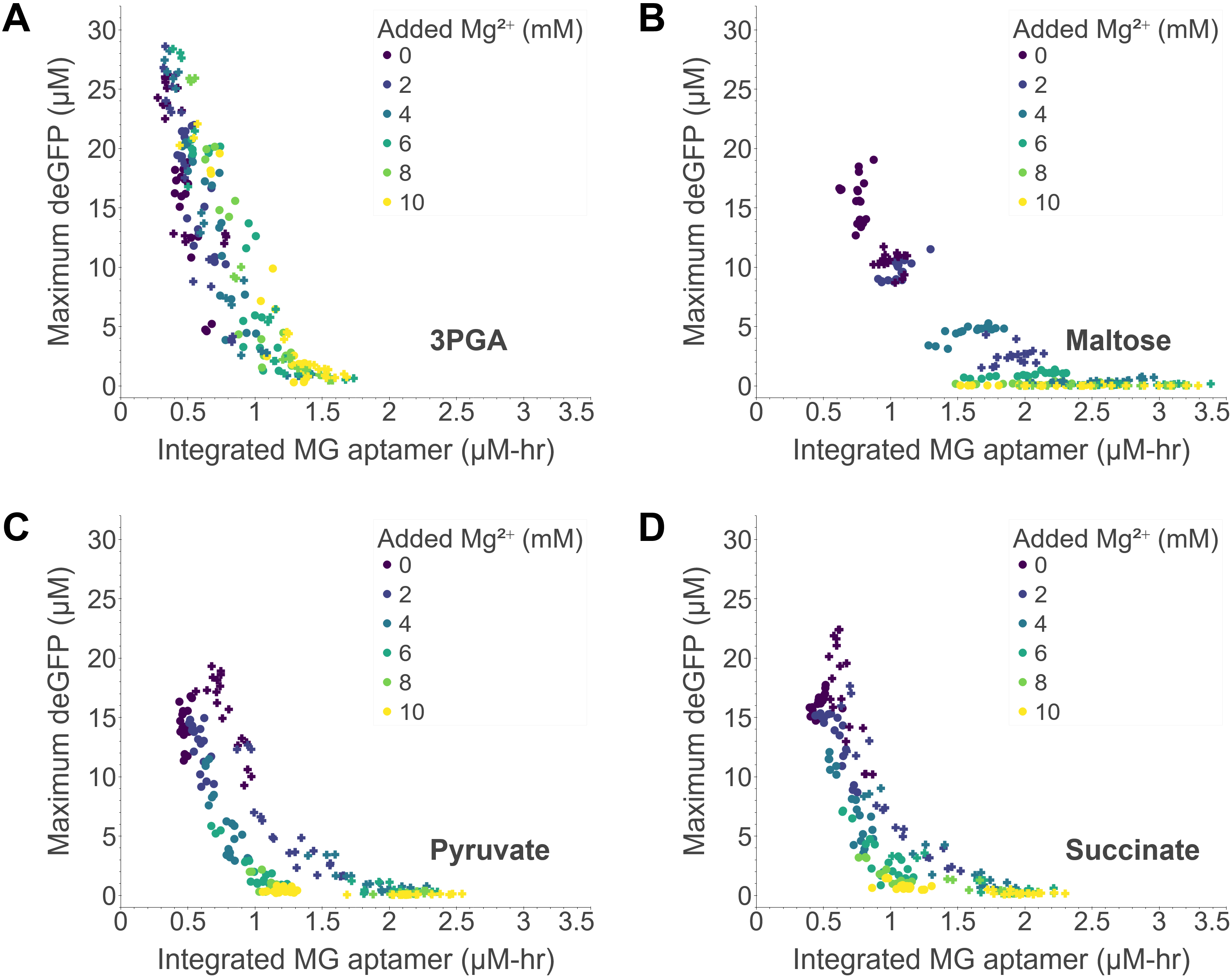

### FigS2_a.png

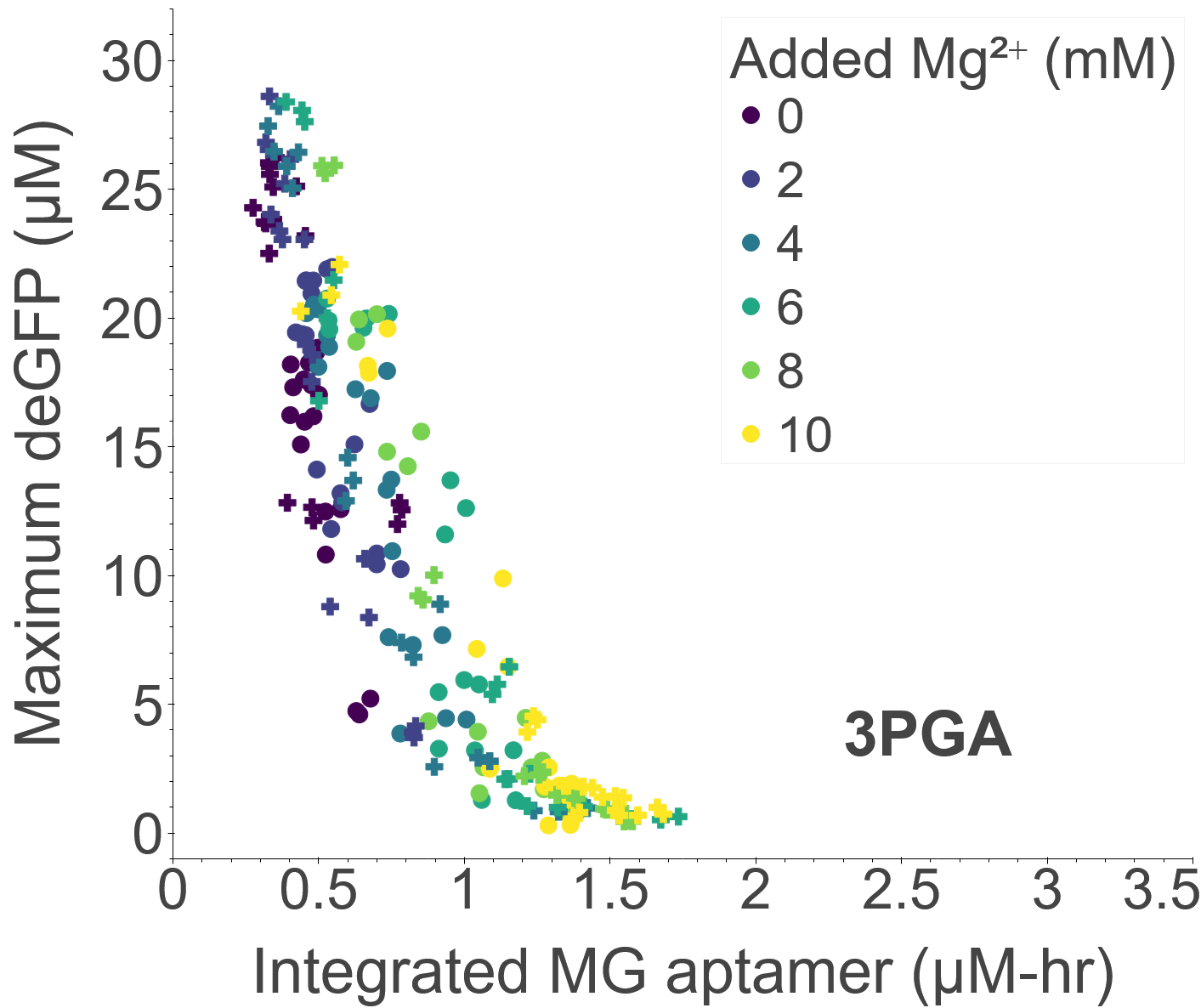

### FigS2_b.png

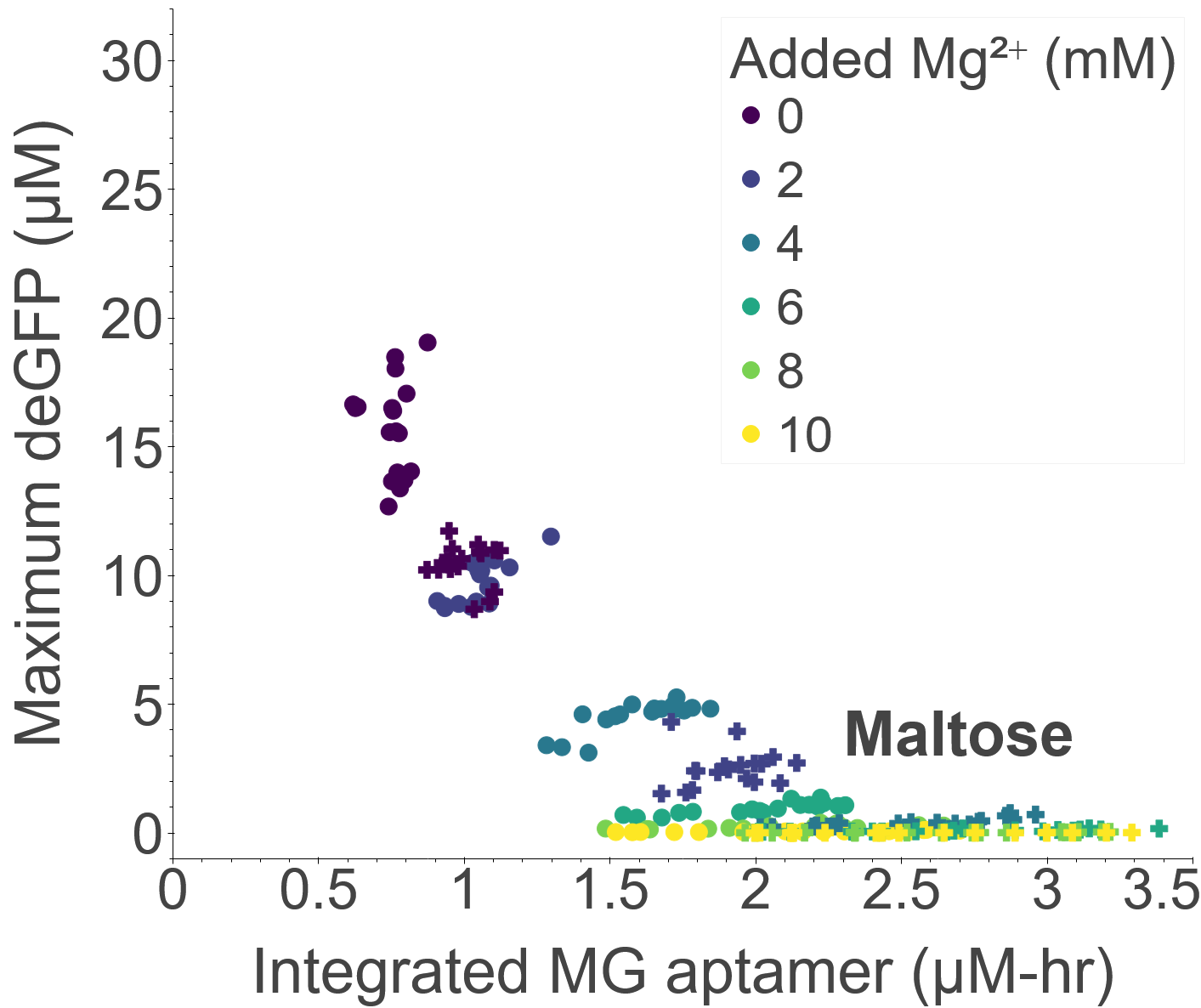

### FigS2_c.png

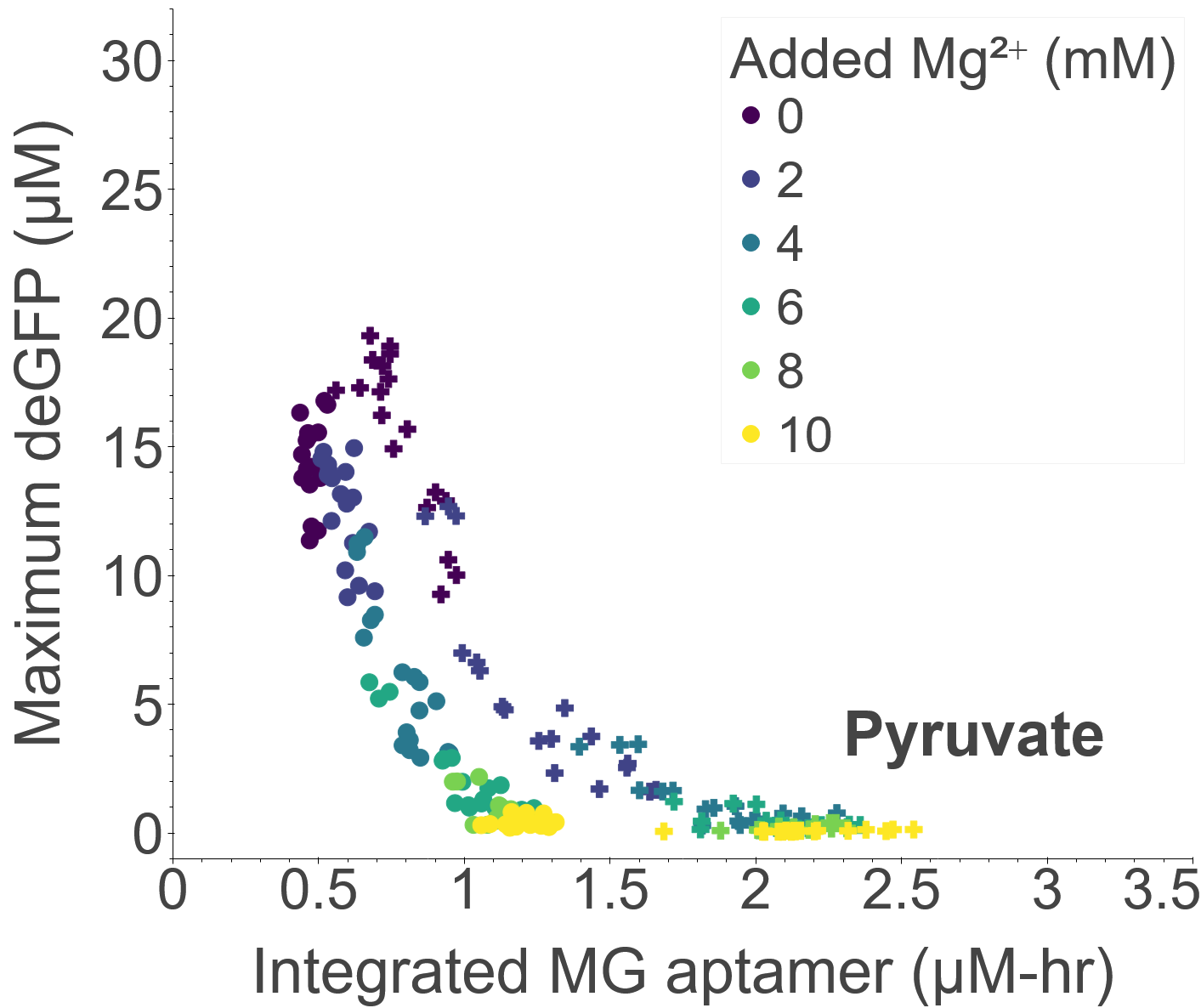

### FigS2_d.png

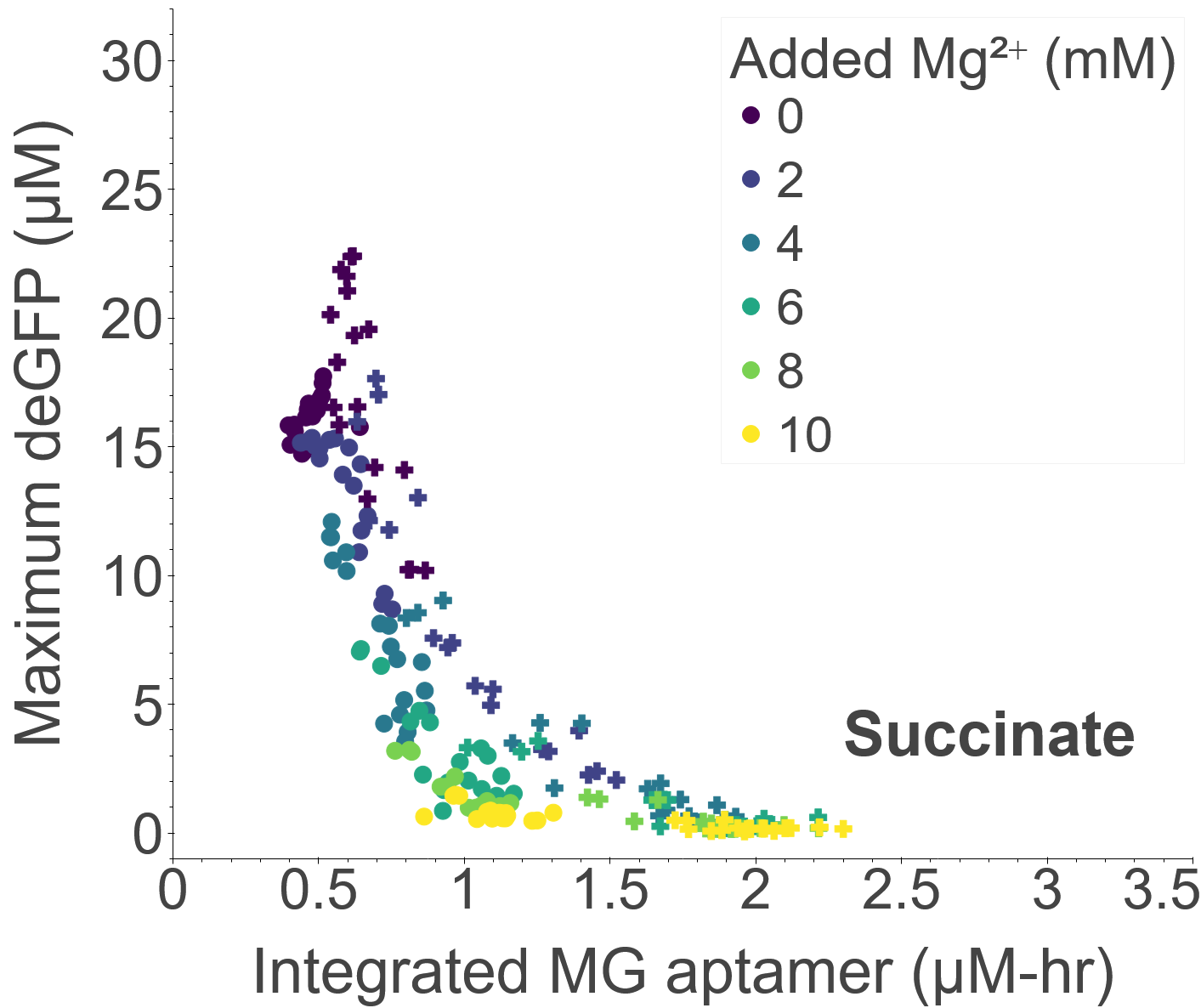

### FigS3.png

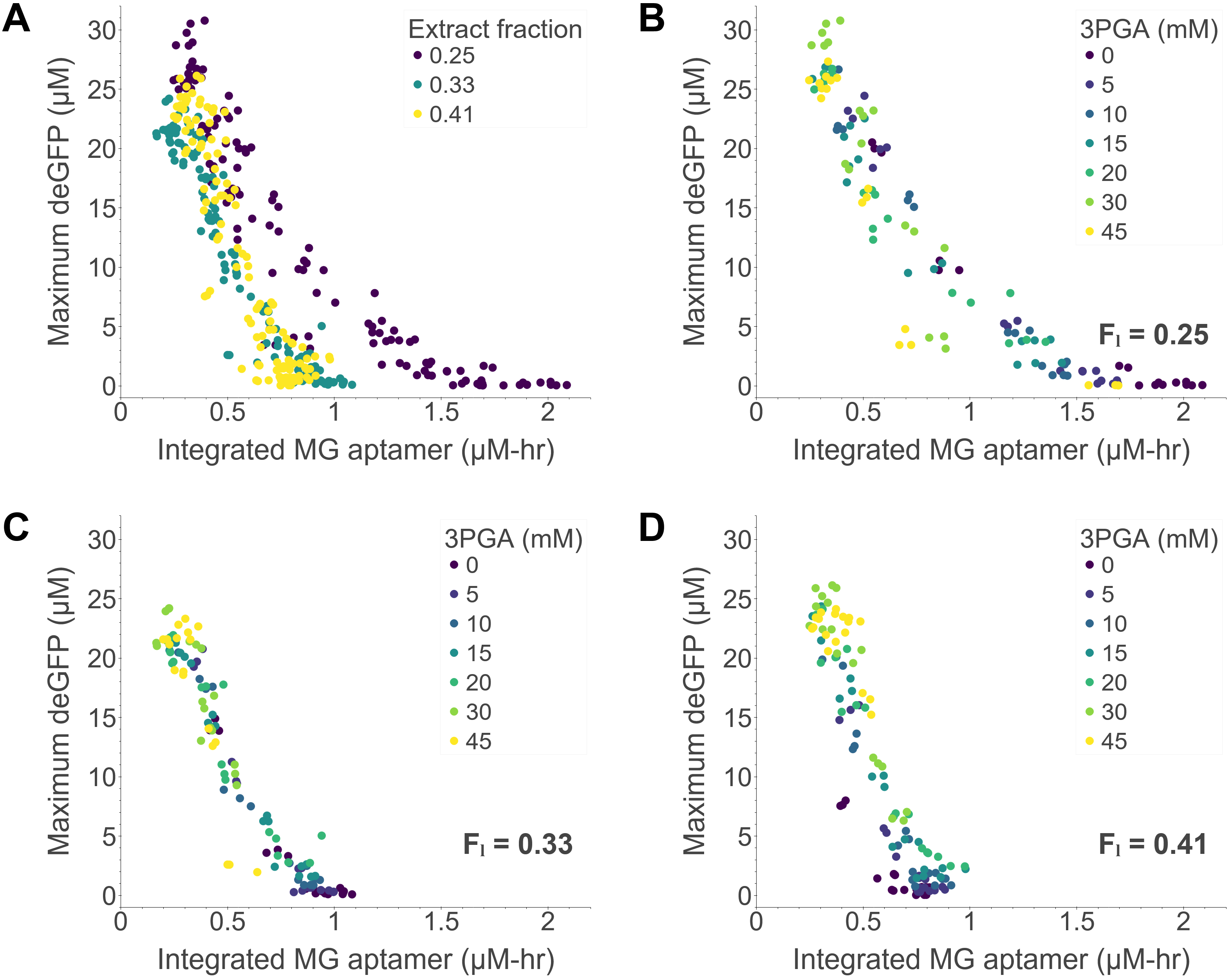

### FigS3_a.png

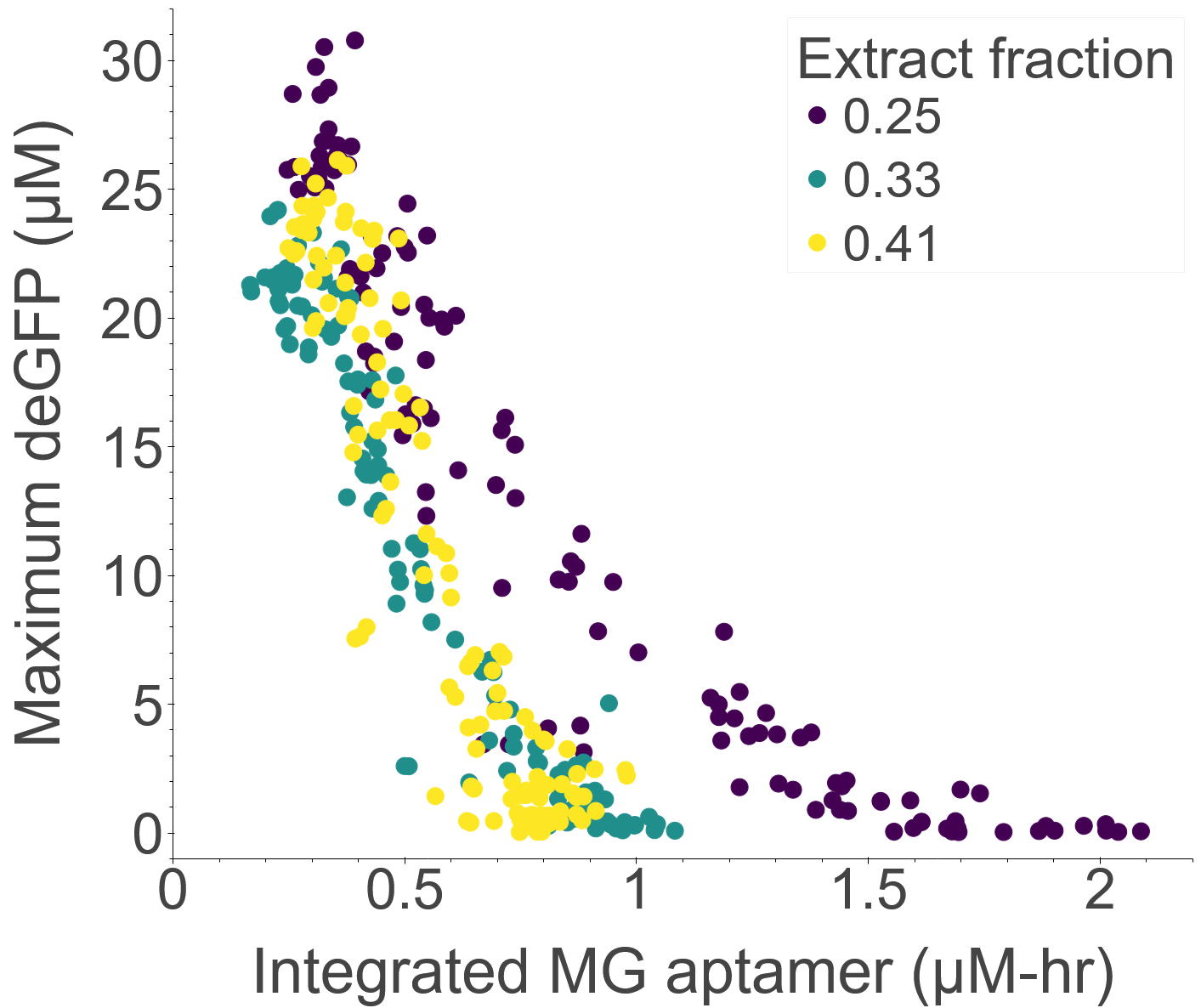

### FigS3_b.png

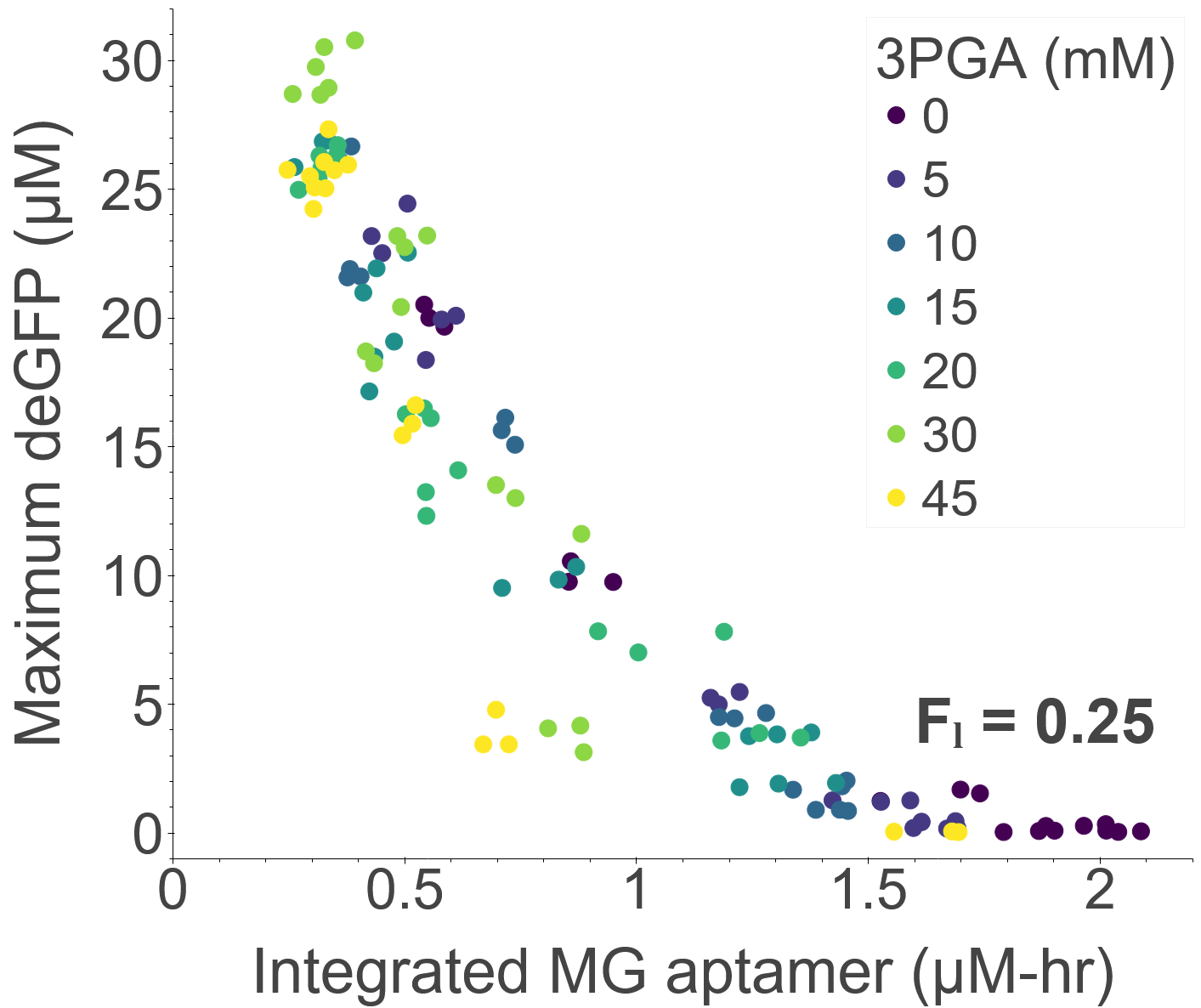

### FigS3_c.png

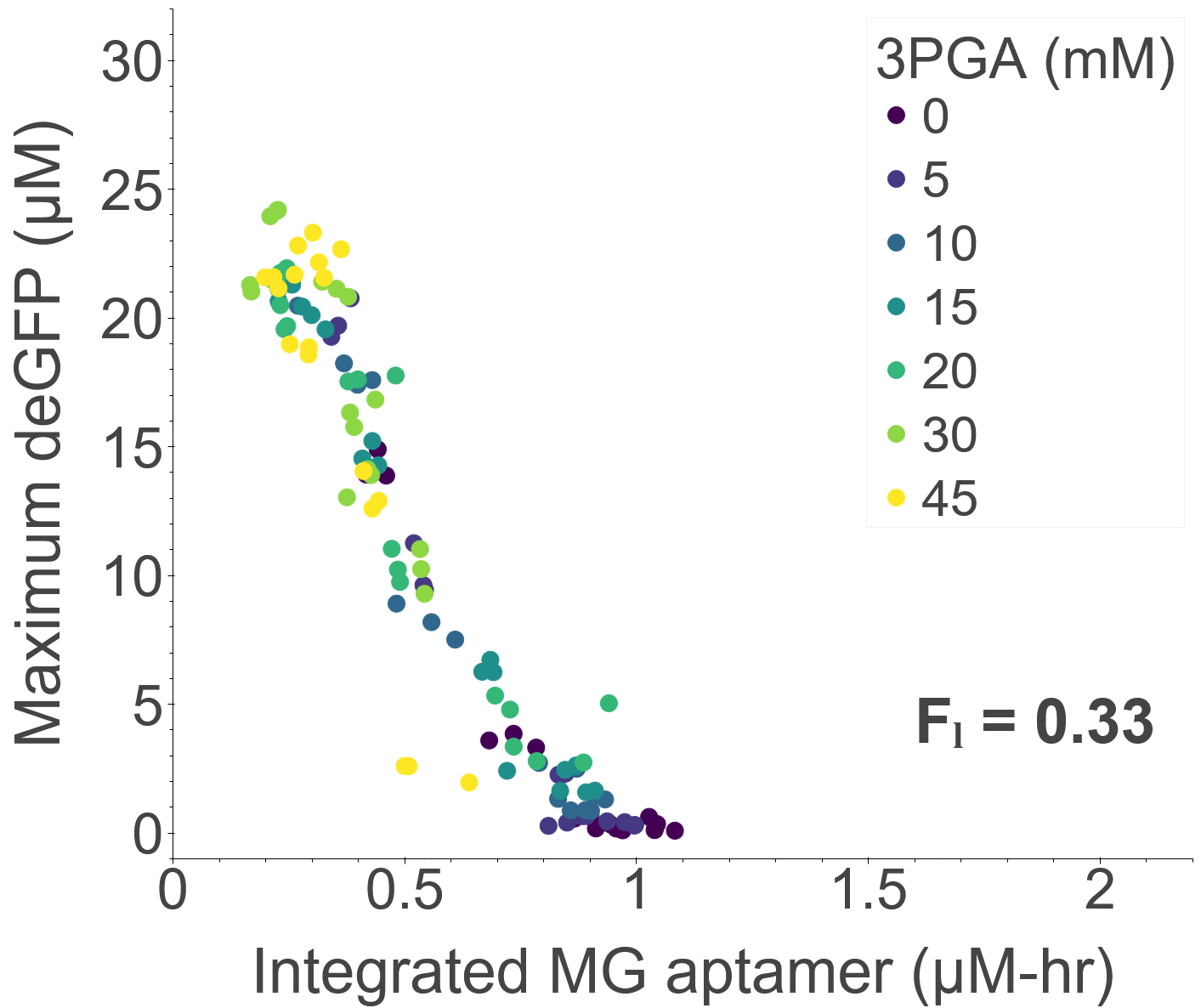

### FigS3_d.png

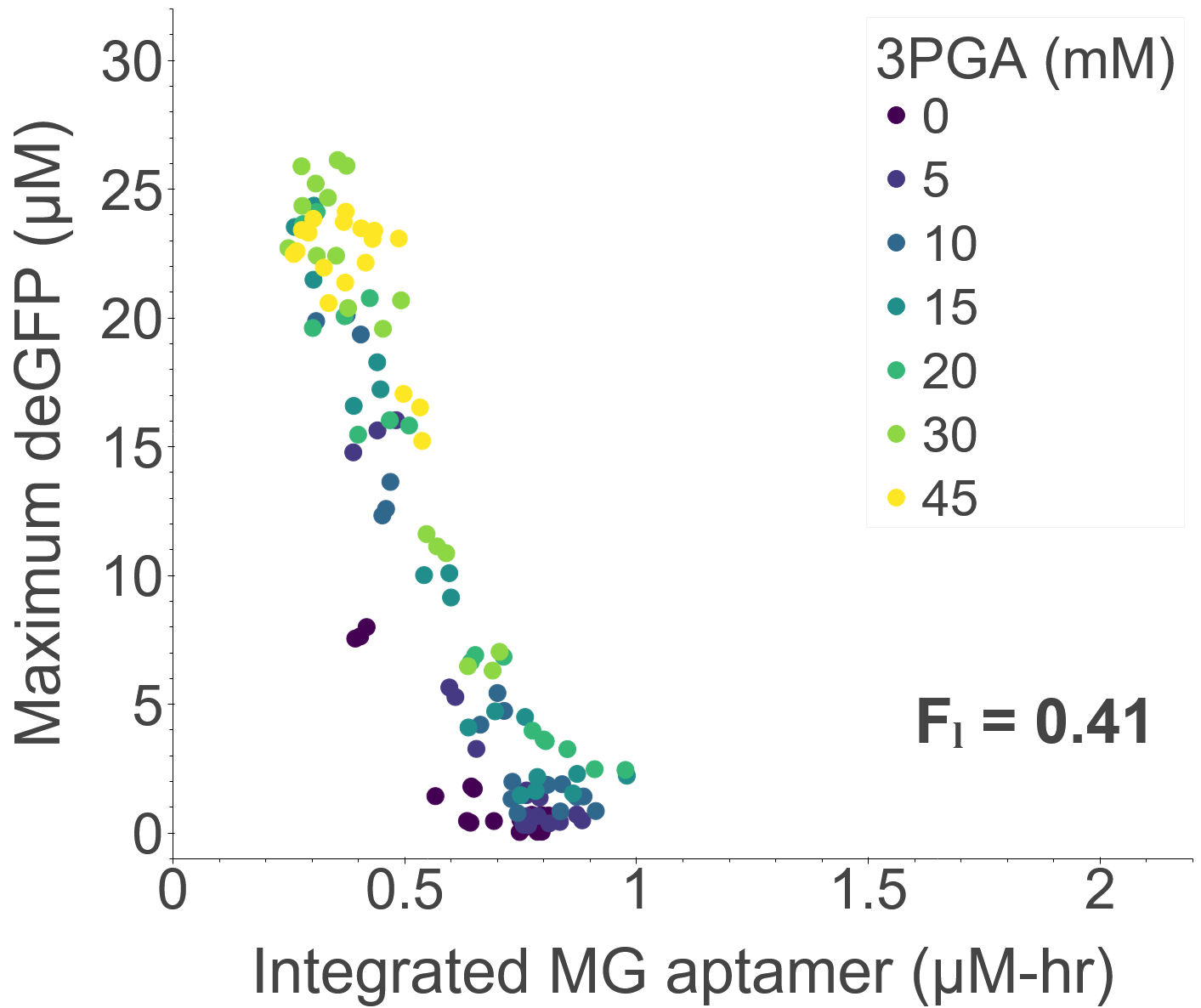

### FigS4.png

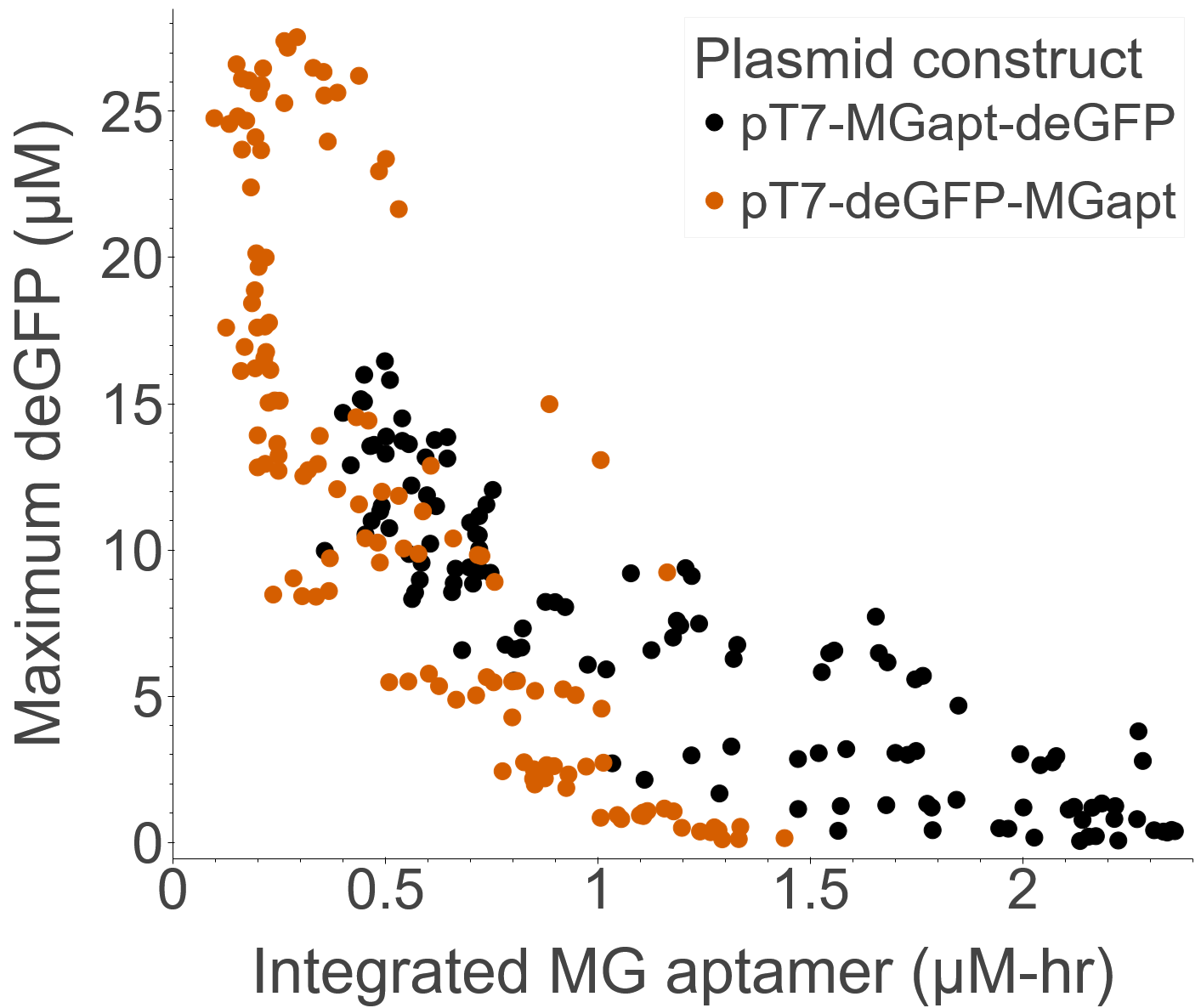

### FigS5.png

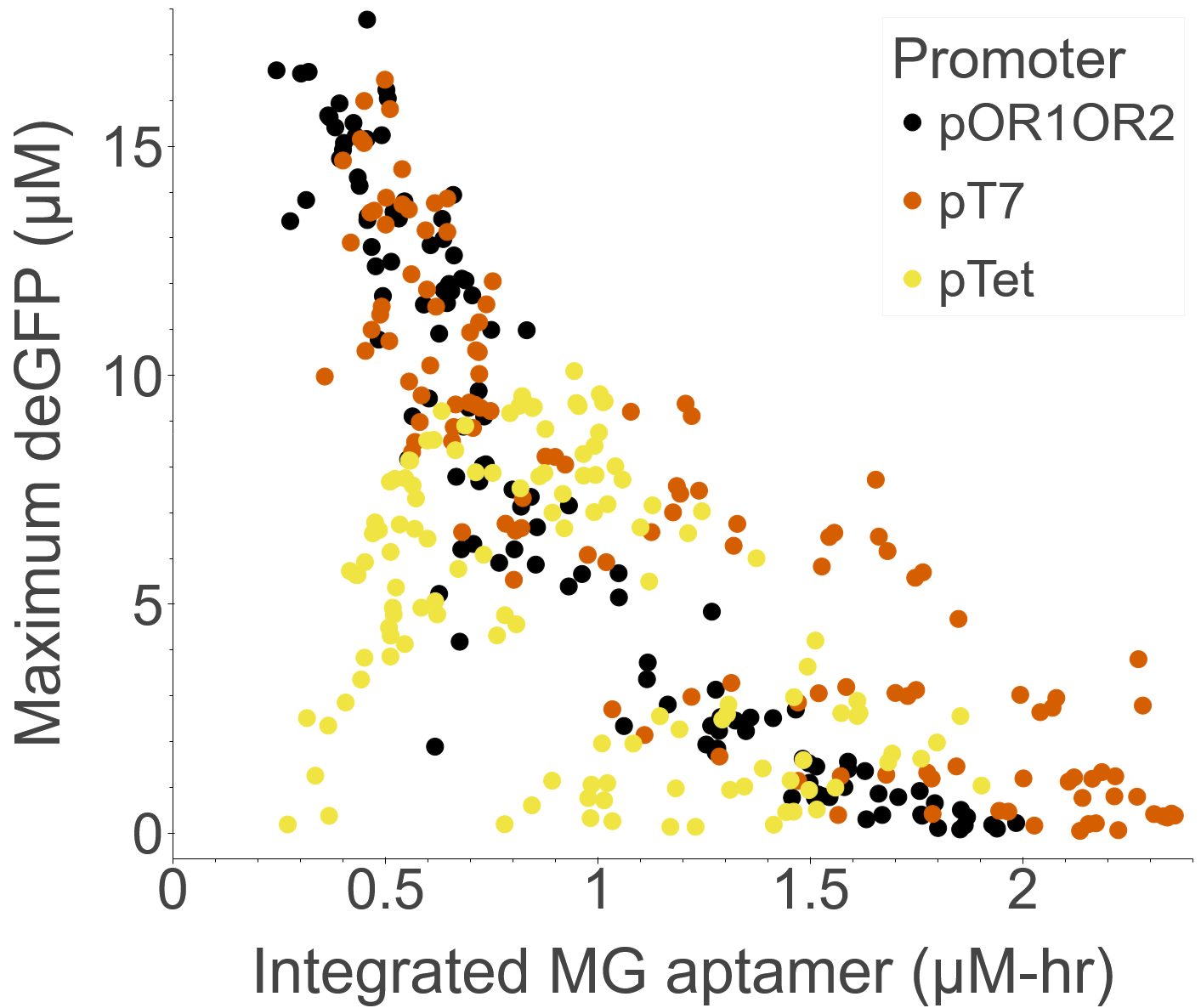

### FigS6.png

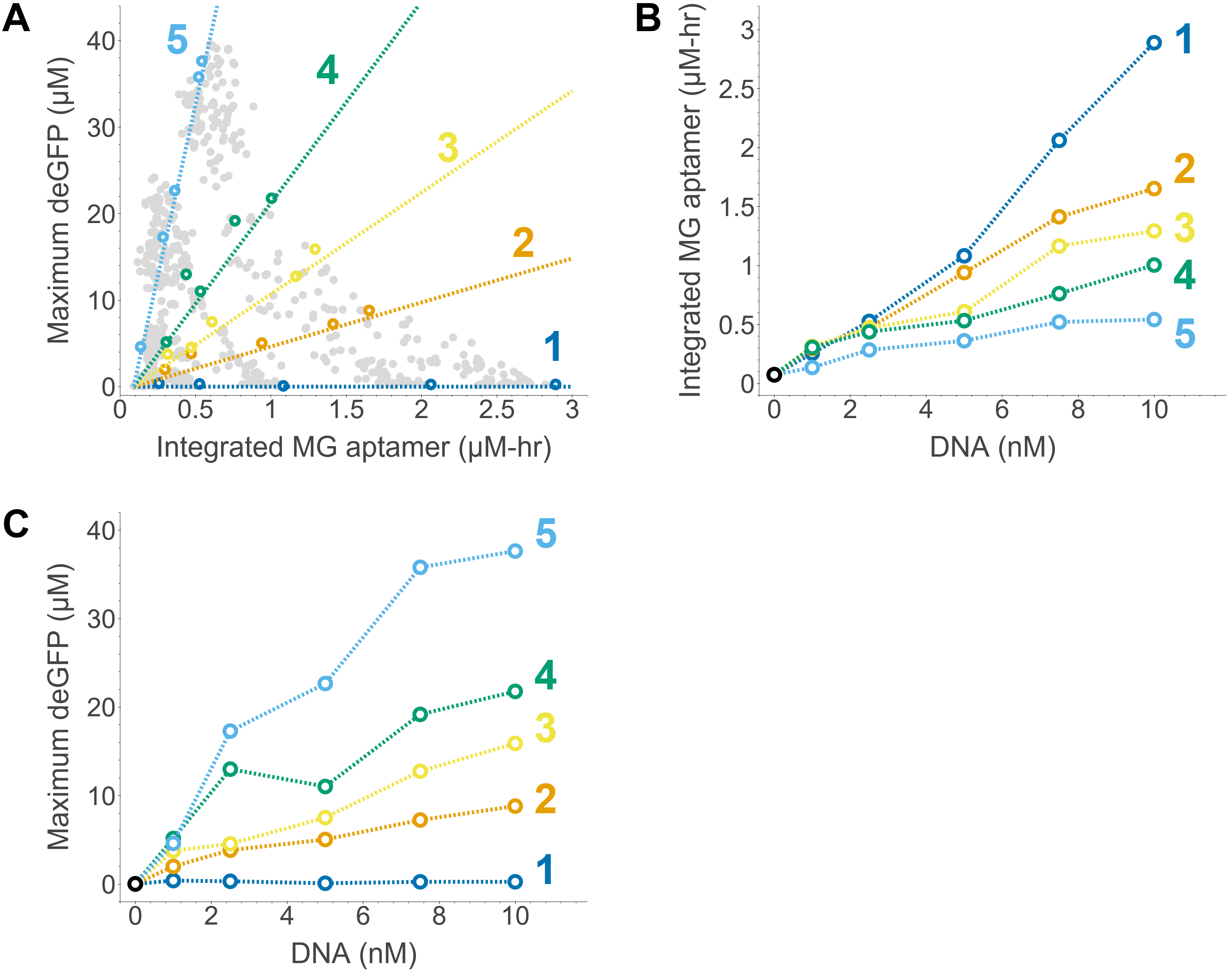

### FigS6_a.png

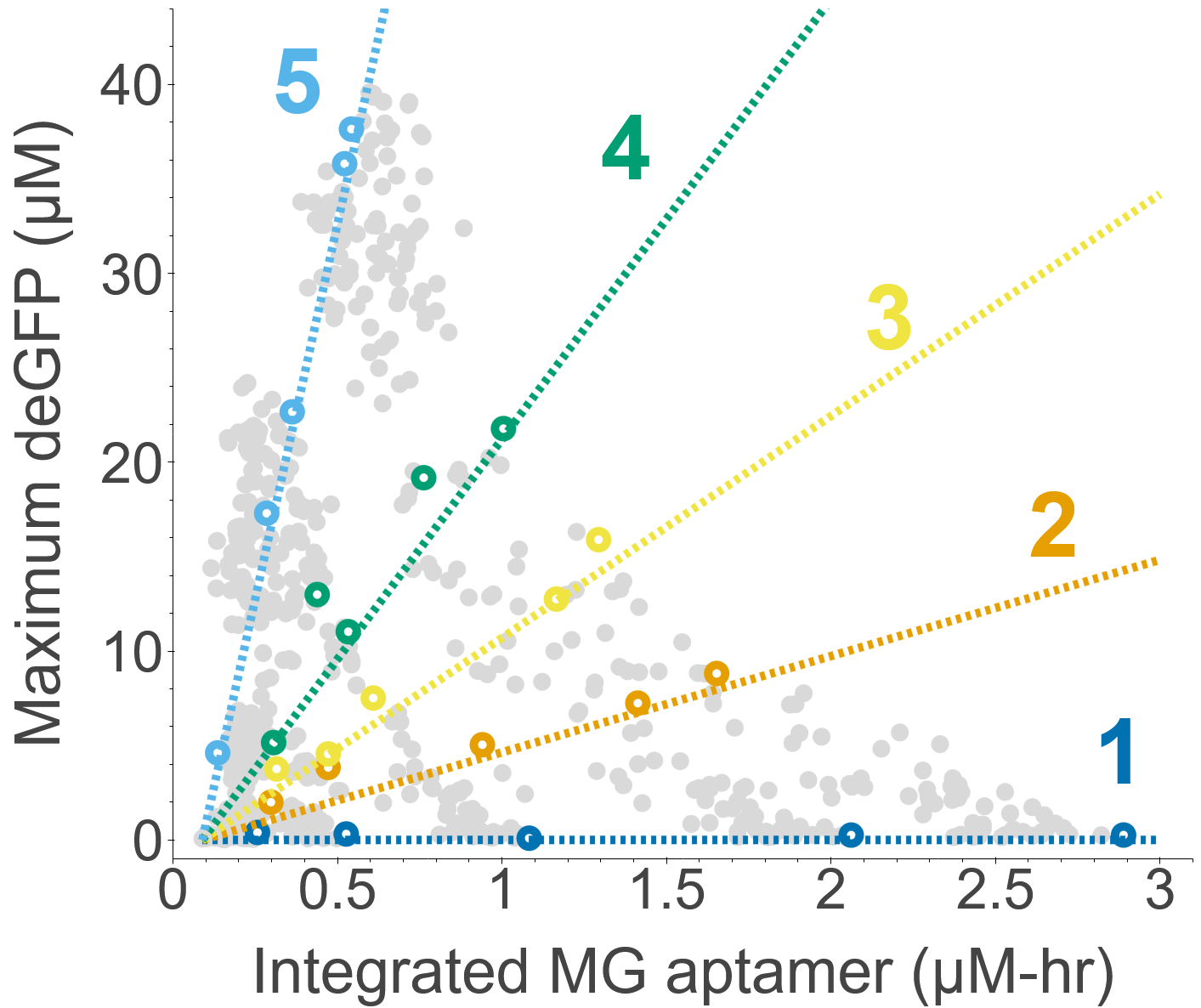

### FigS6_b.png

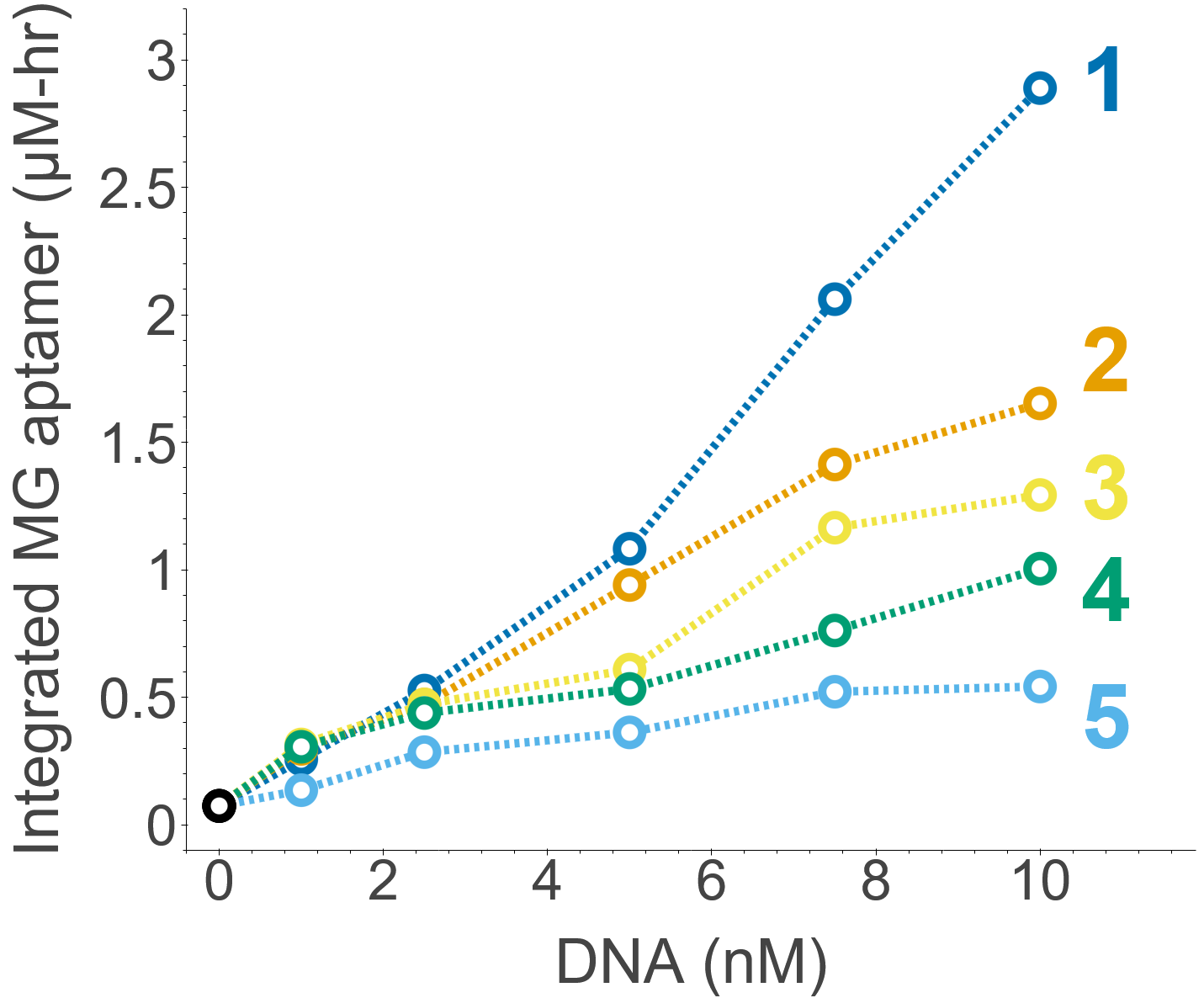

### FigS6_c.png

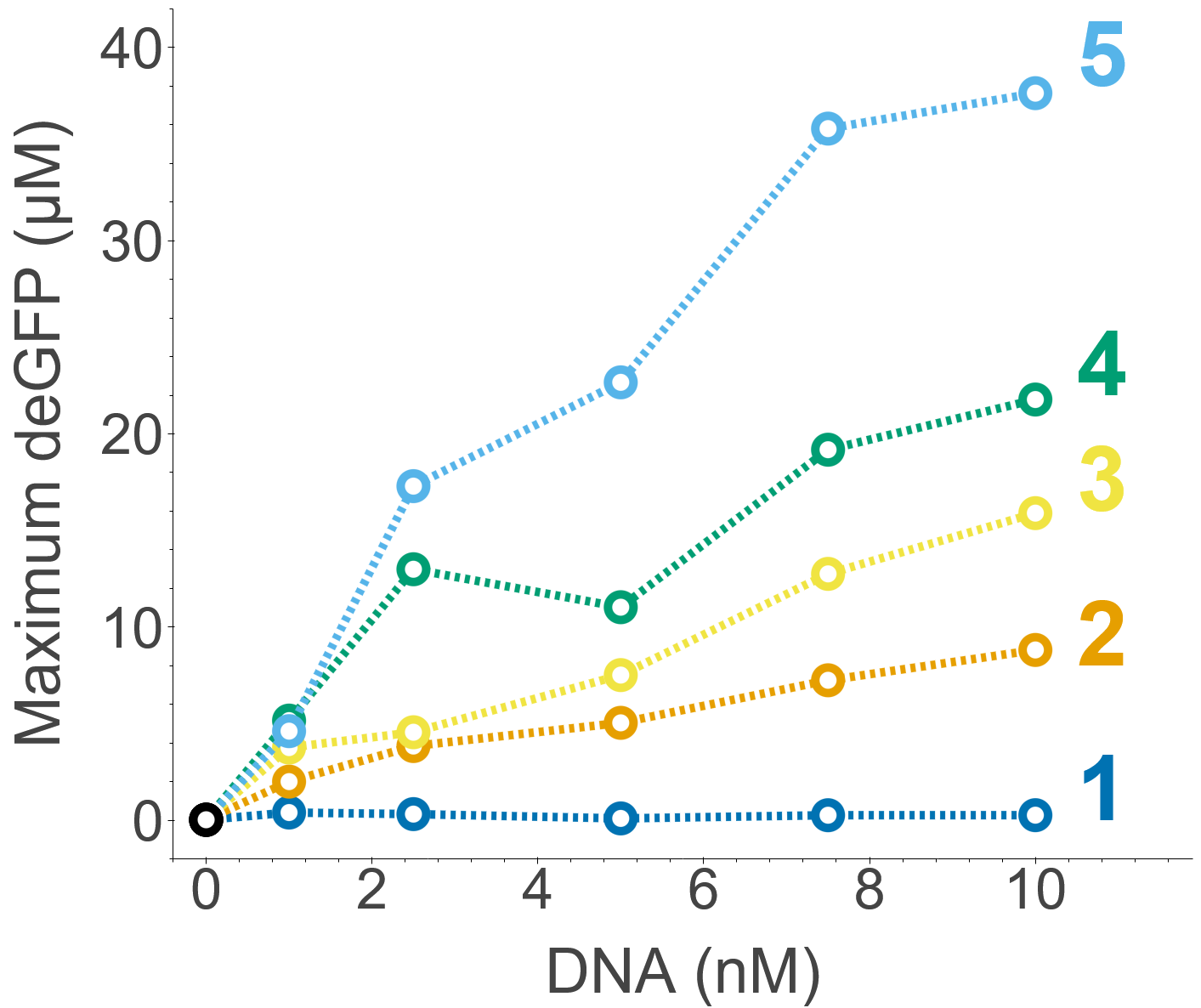

### FigS7.png

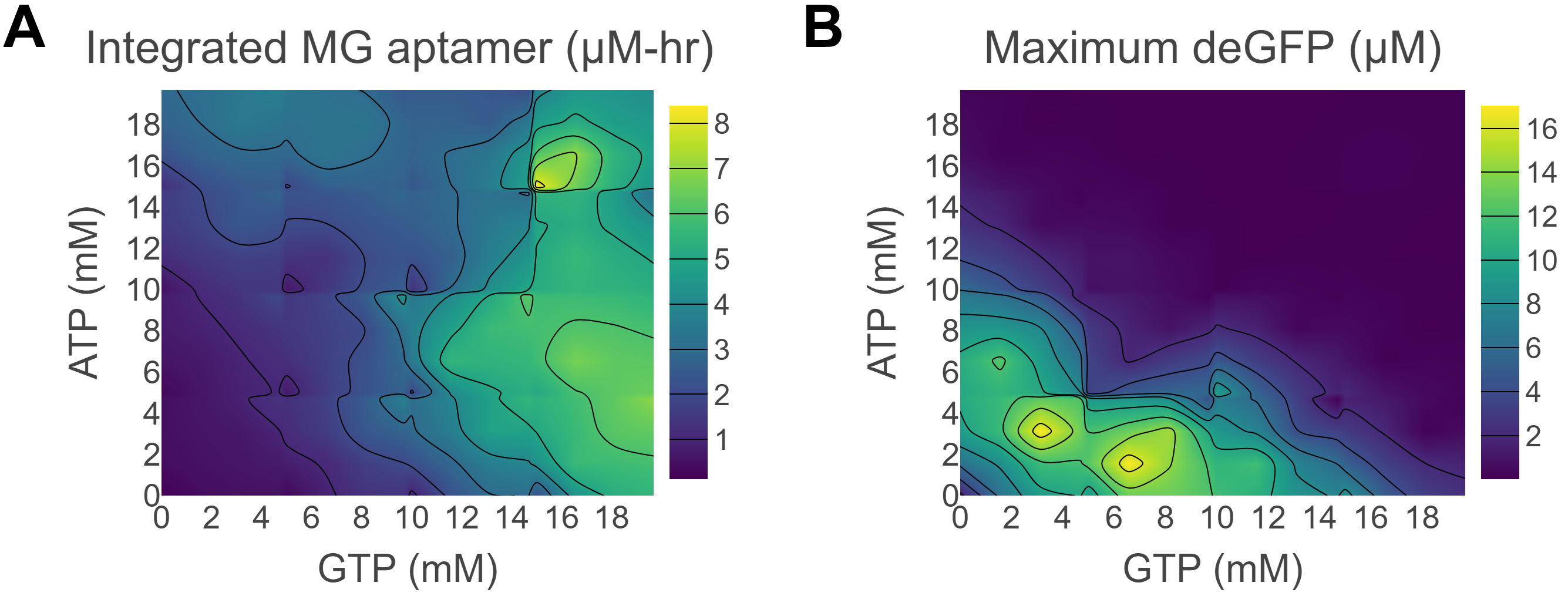

### FigS7_a.png

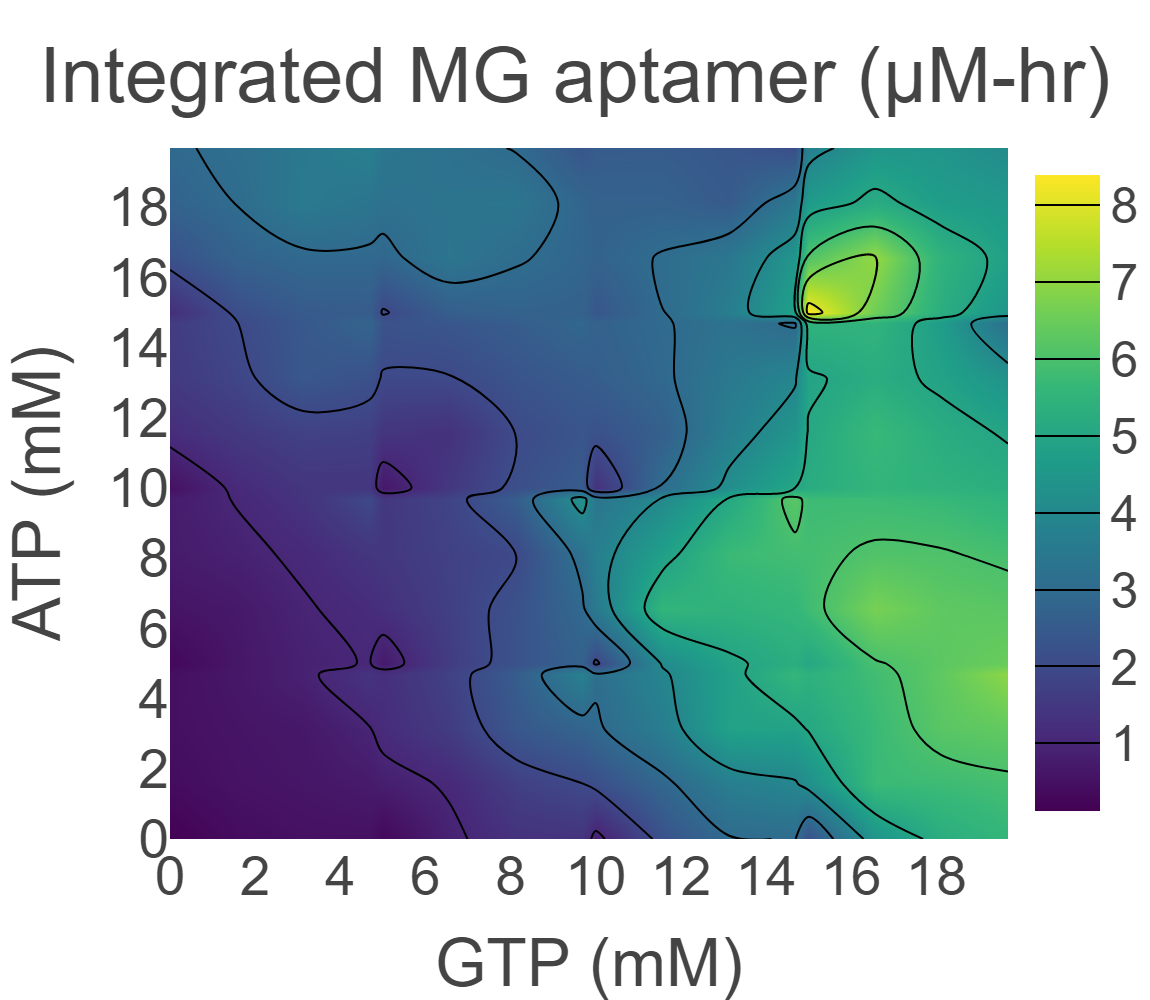

### FigS7_b.png

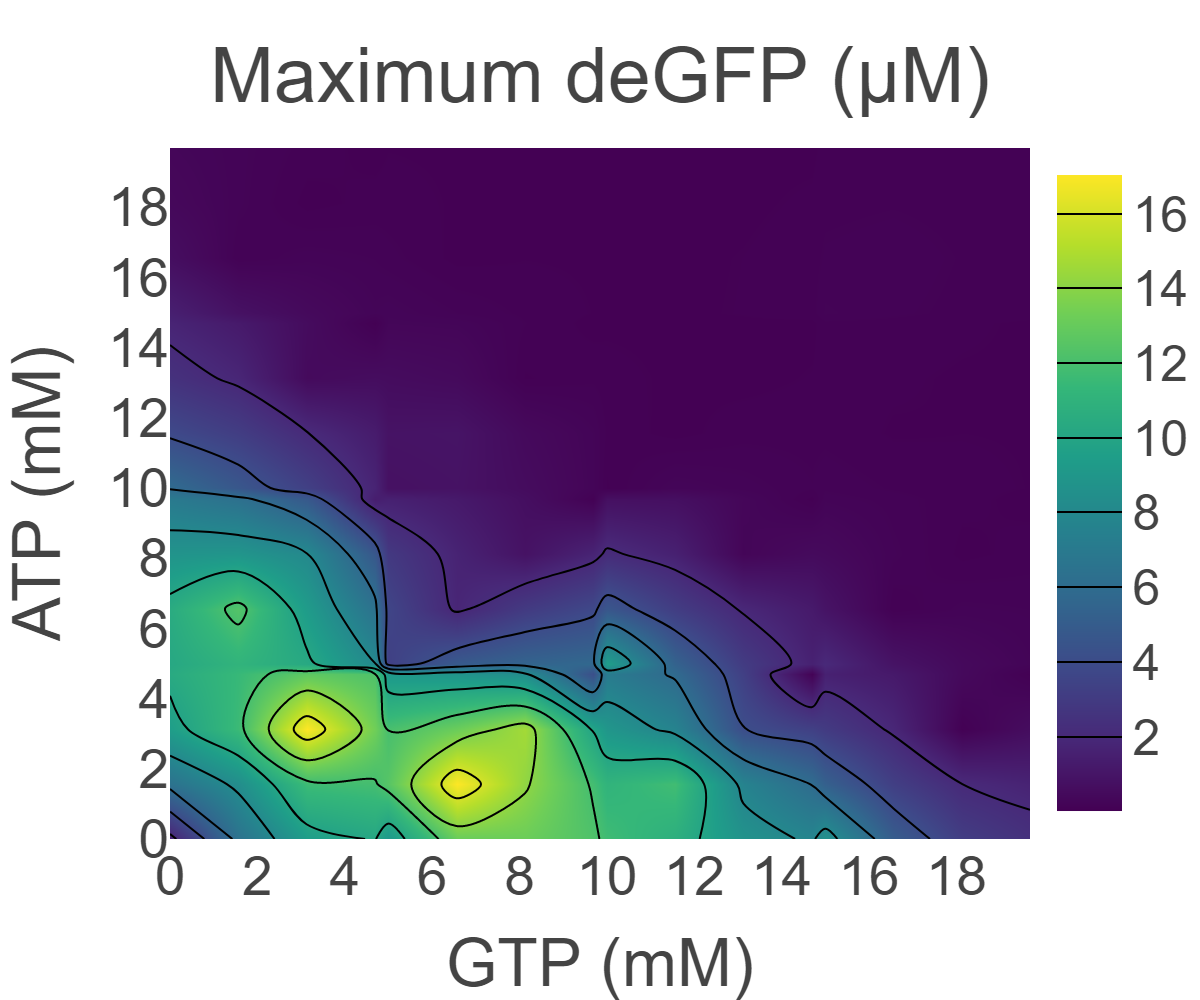

### FigS8.png

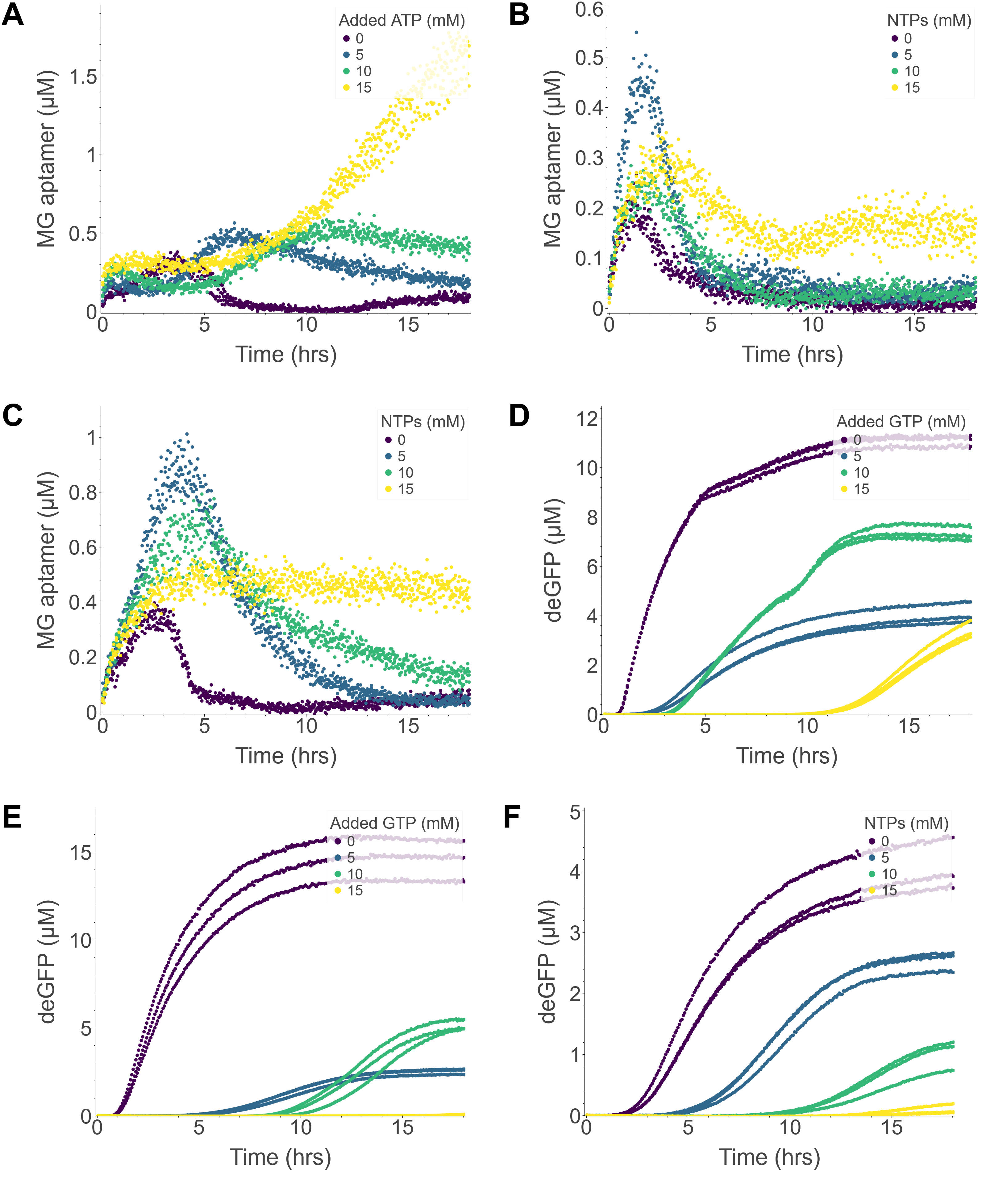

### FigS8_a.png

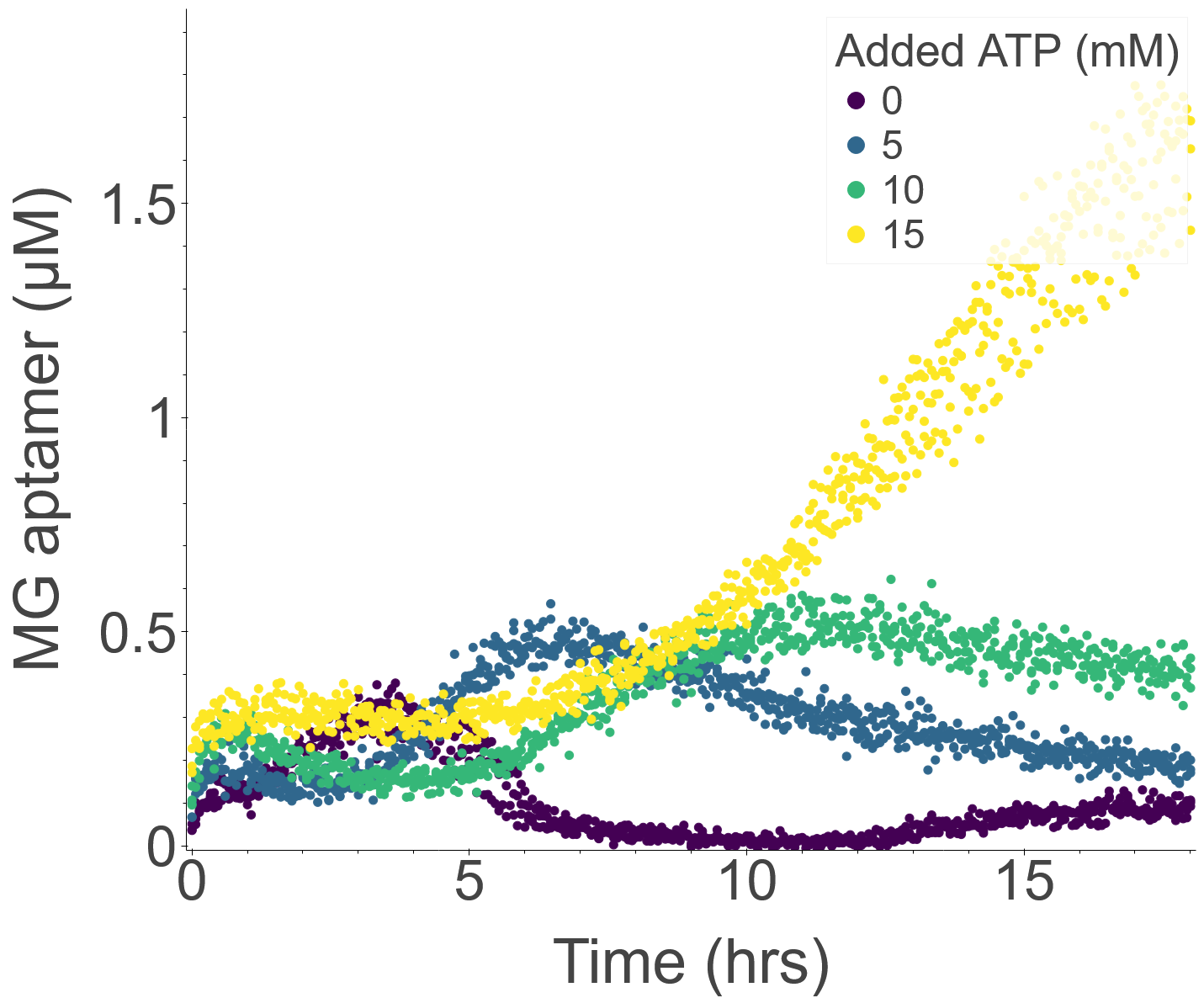

### FigS8_b.png

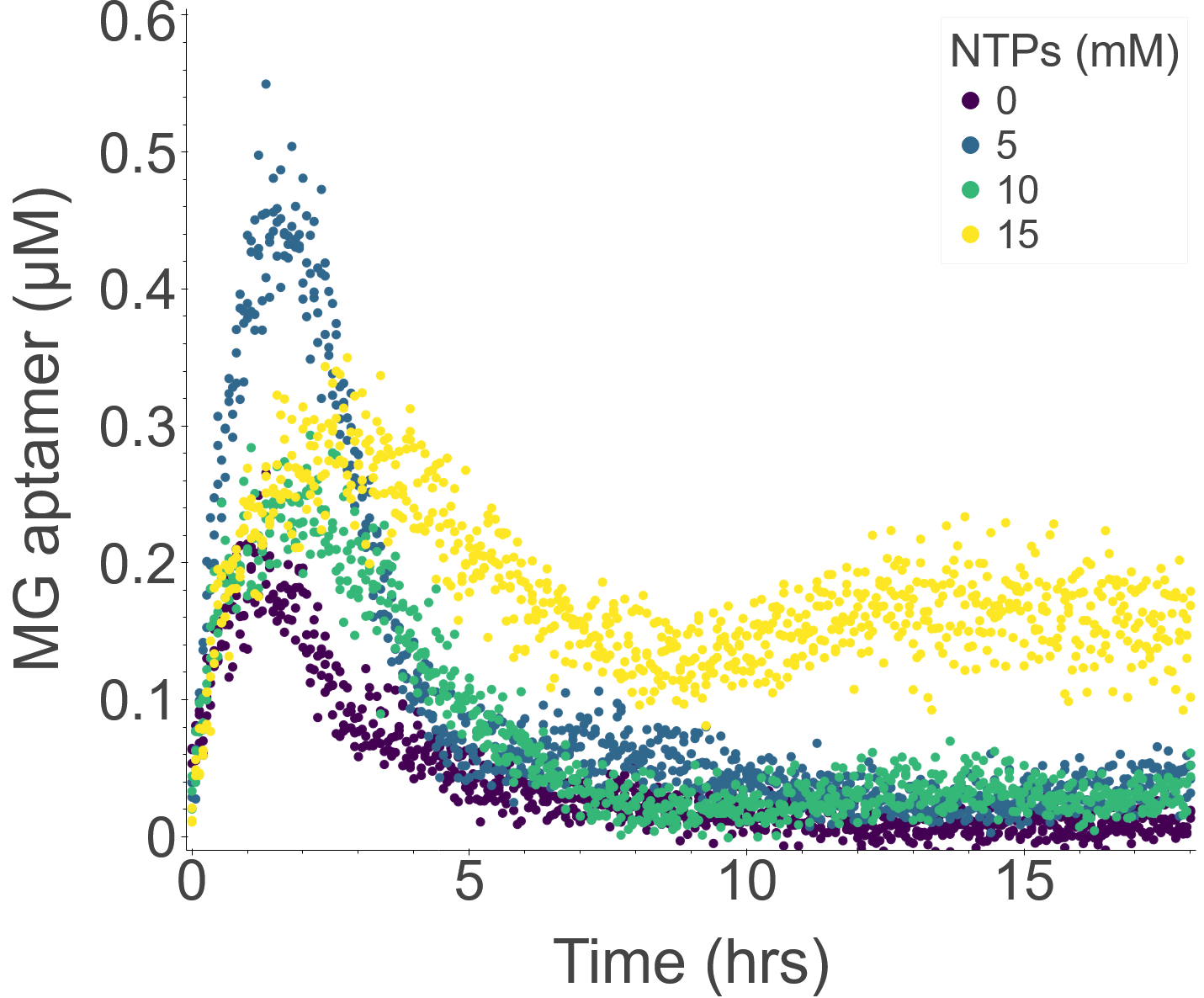

### FigS8_c.png

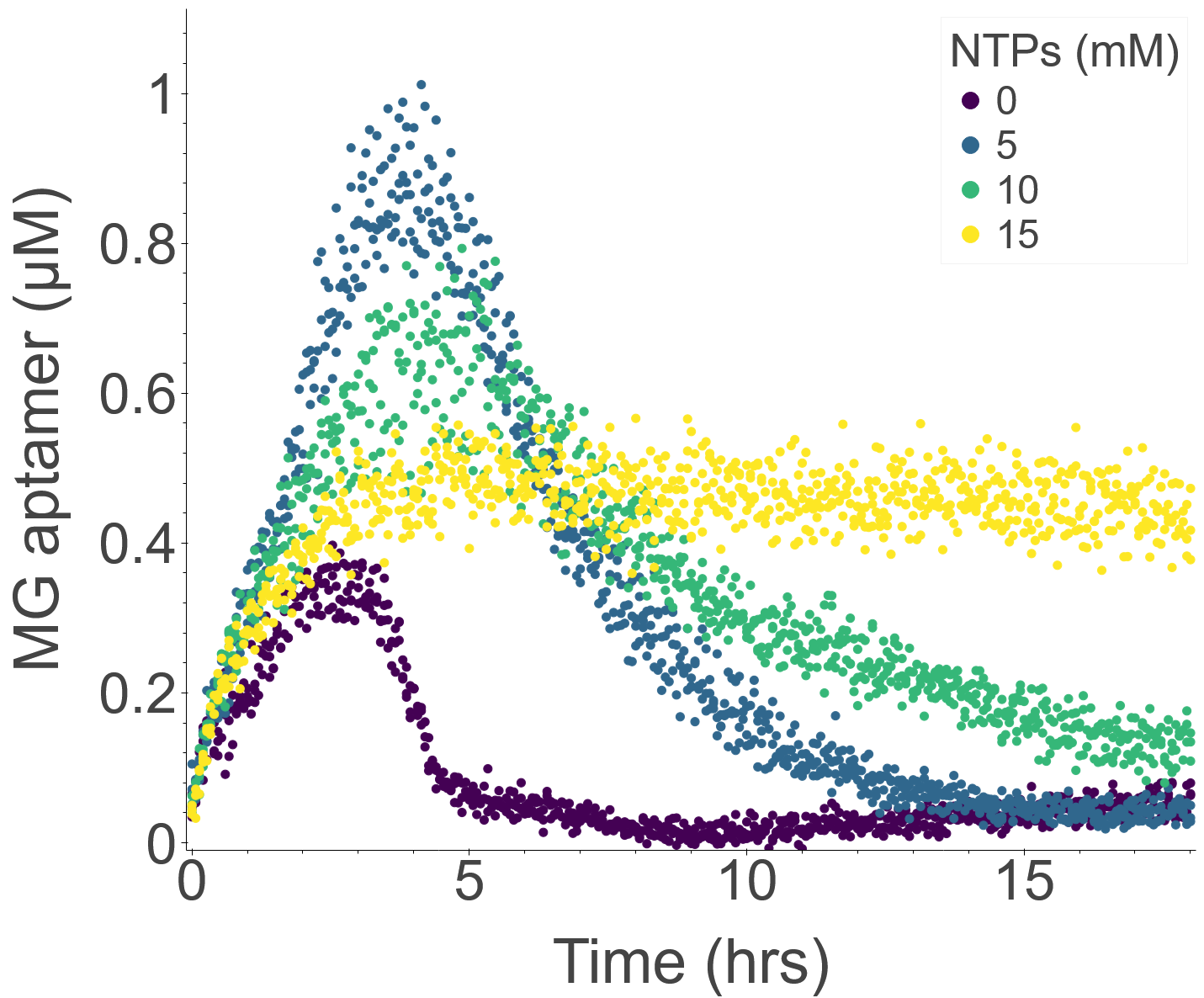

### FigS8_d.png

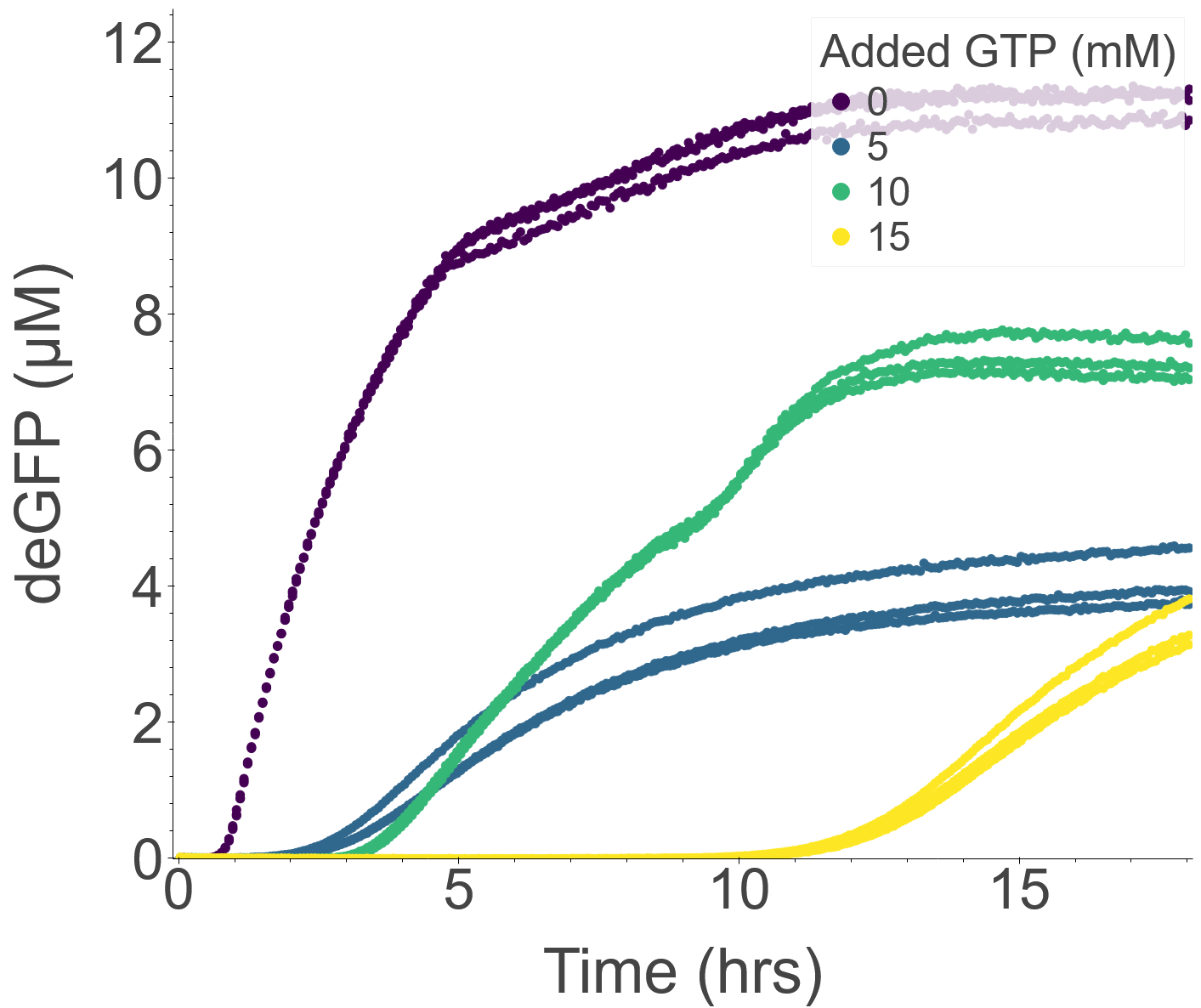

### FigS8_e.png

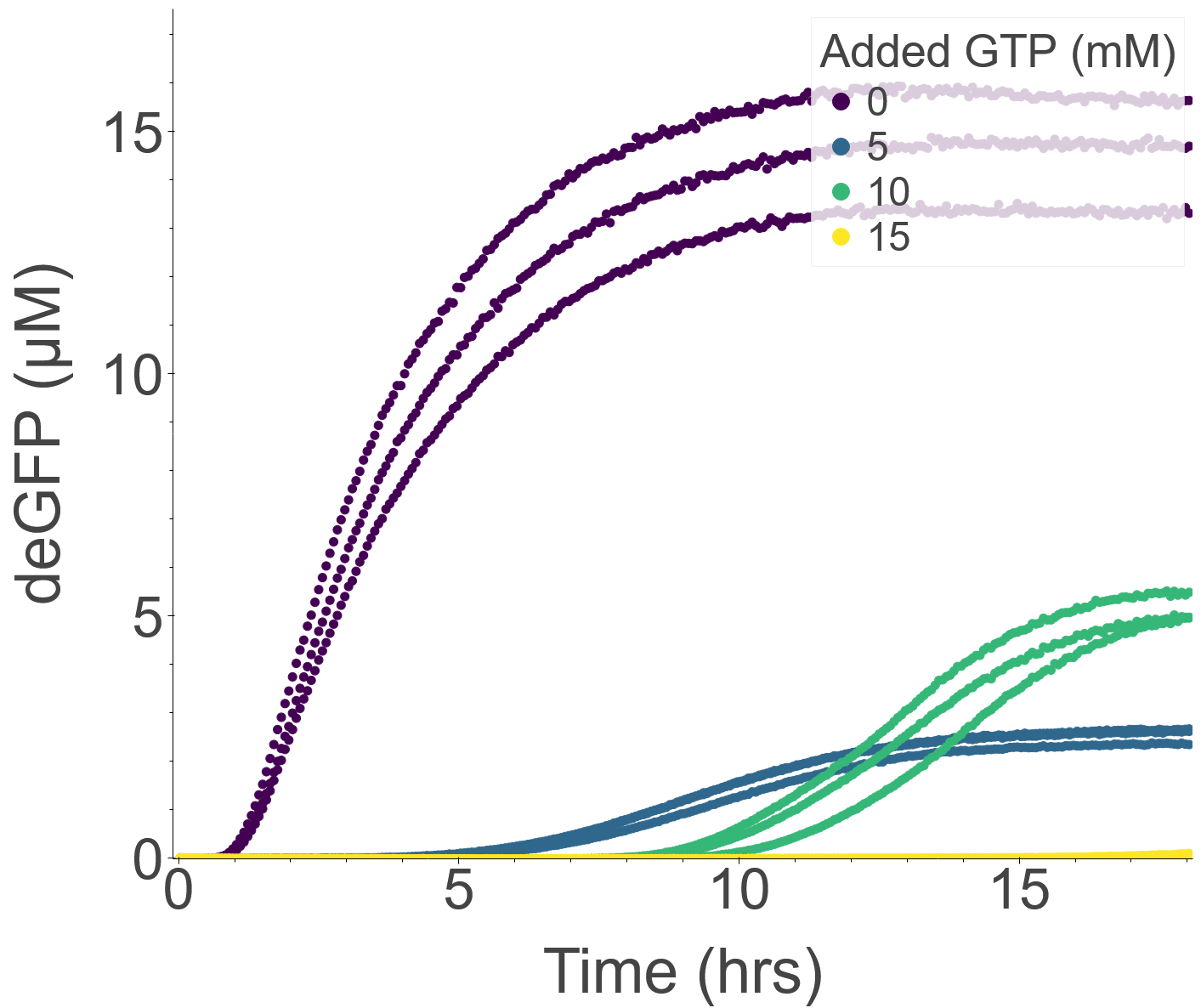

### FigS8_f.png

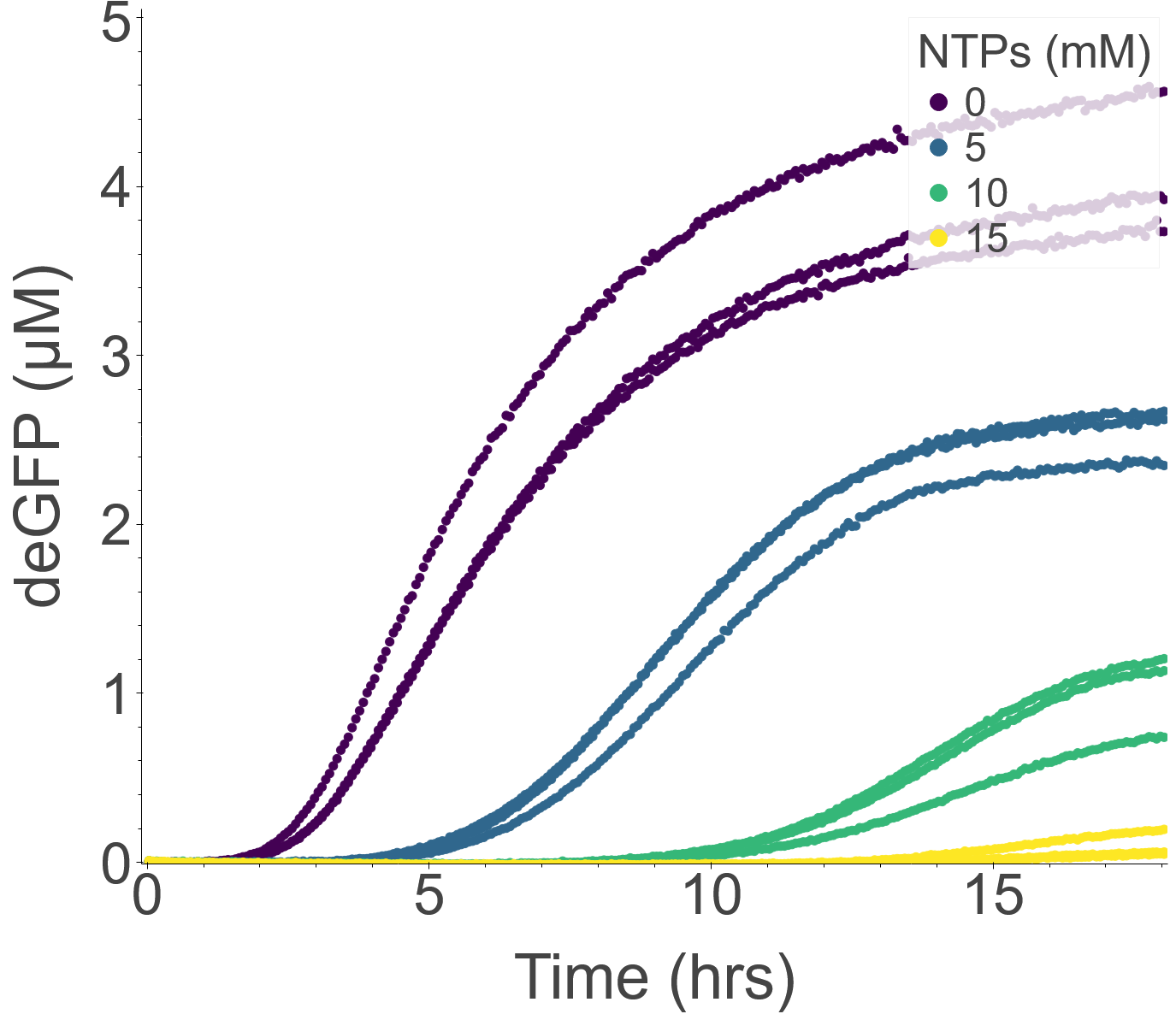

### FigS9.png

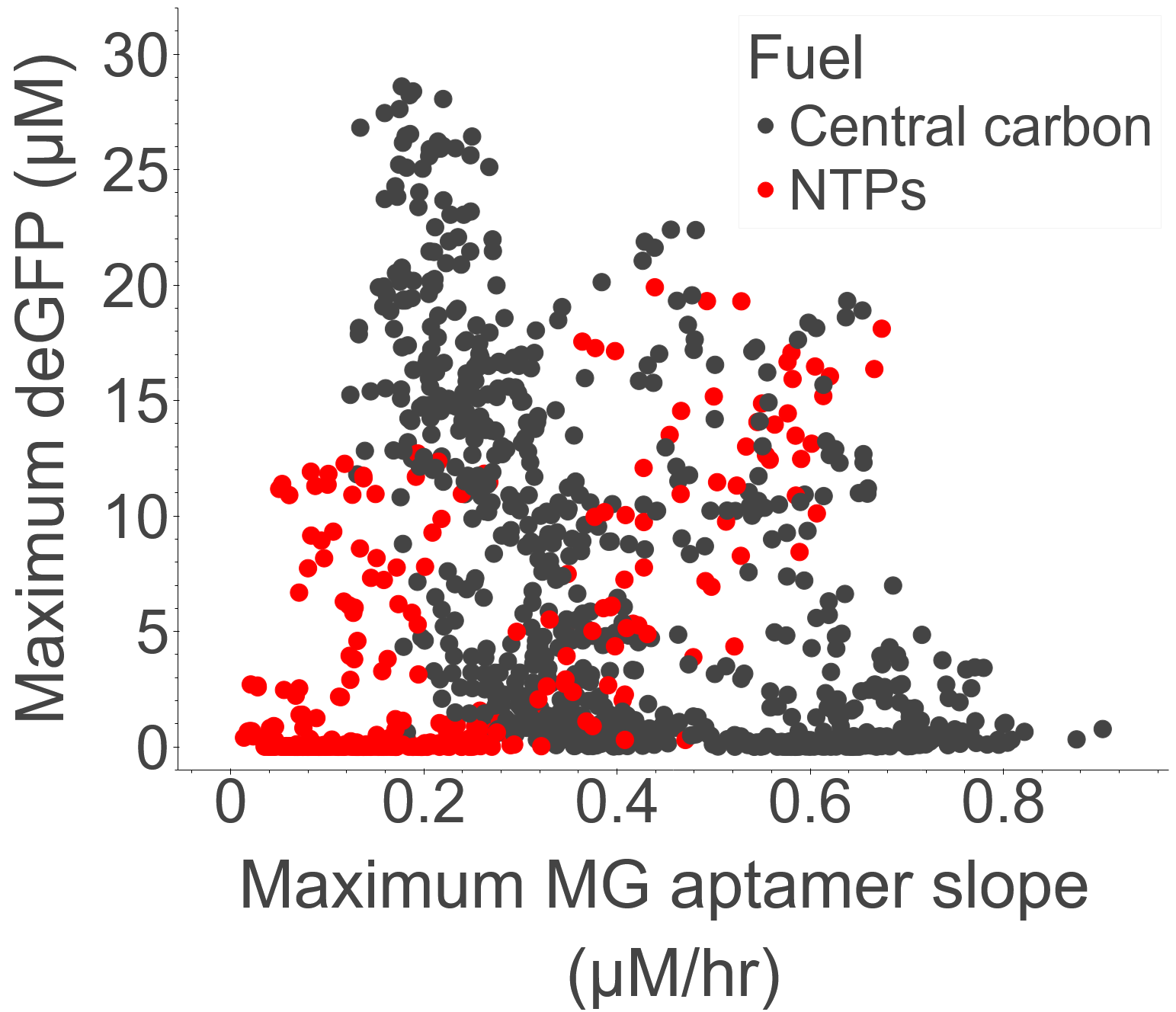

### FigS10.png

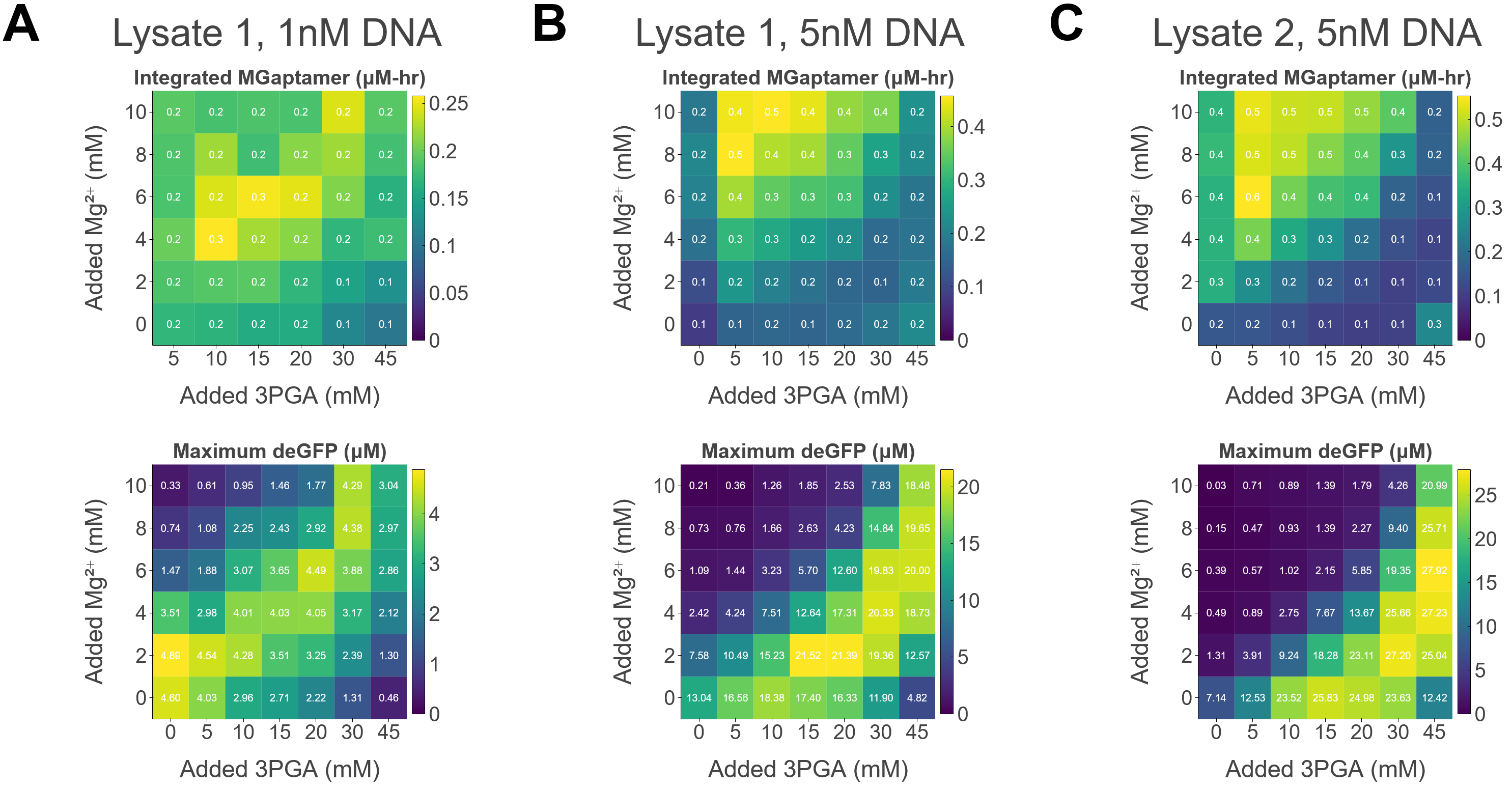

### FigS10_a.png

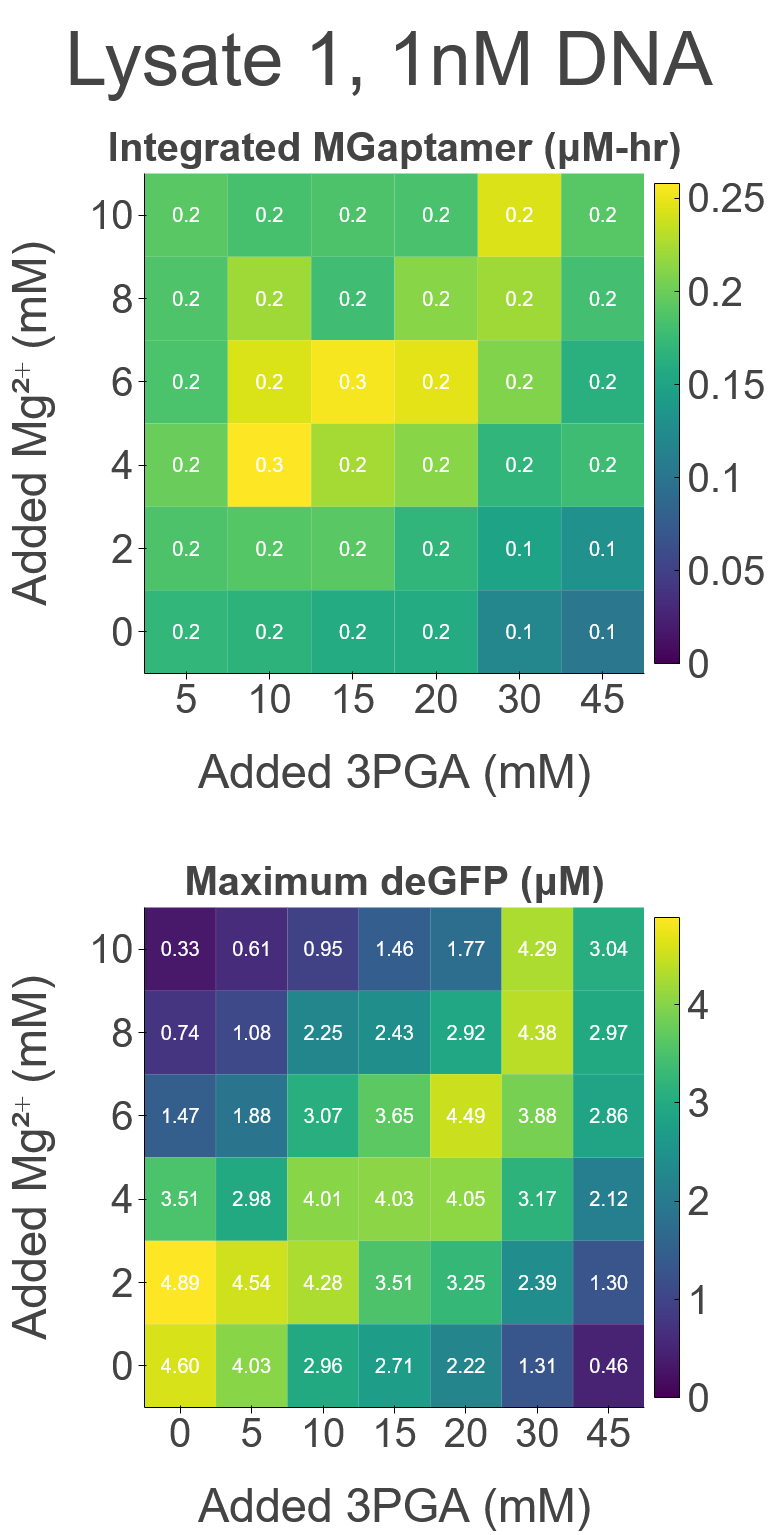

### FigS10_b.png

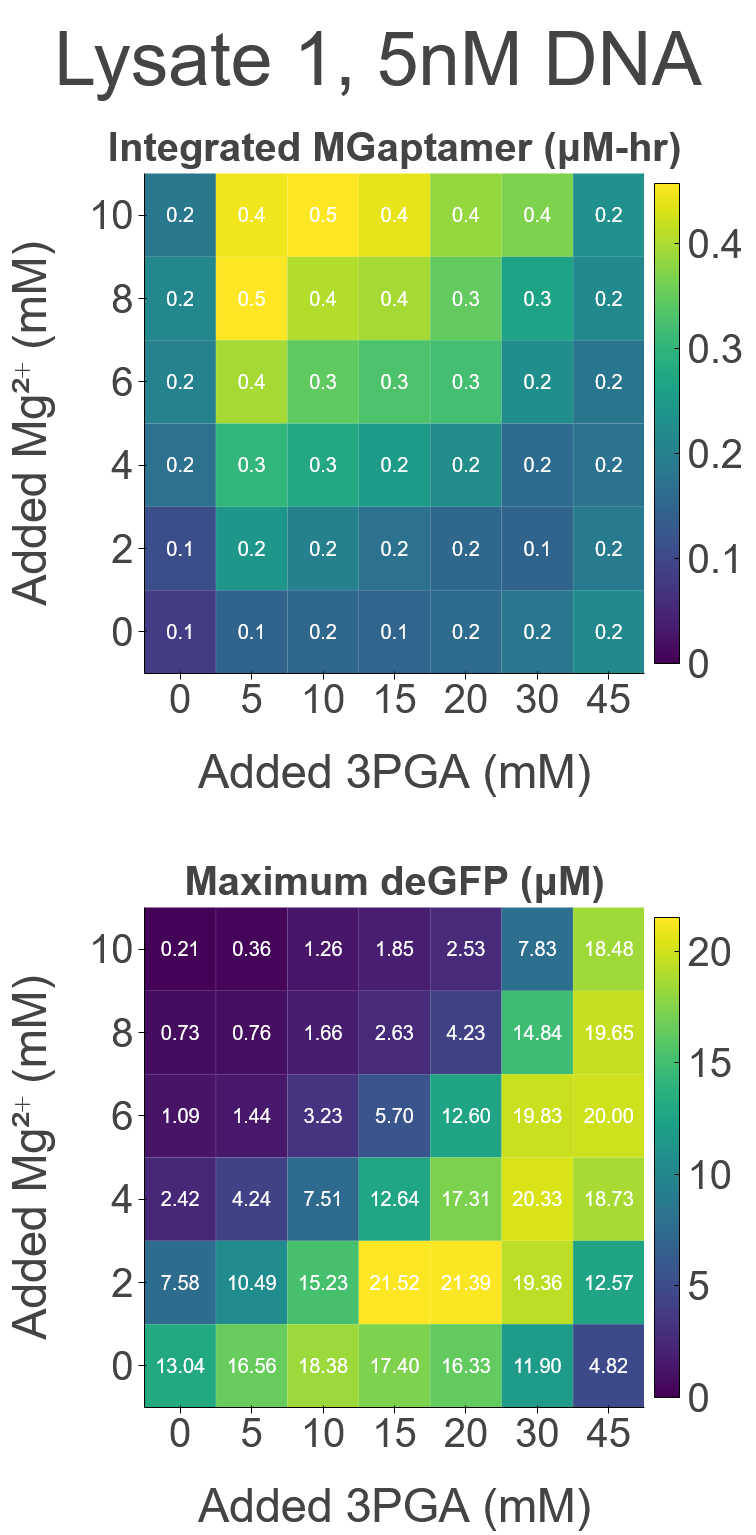

### FigS10_c.png

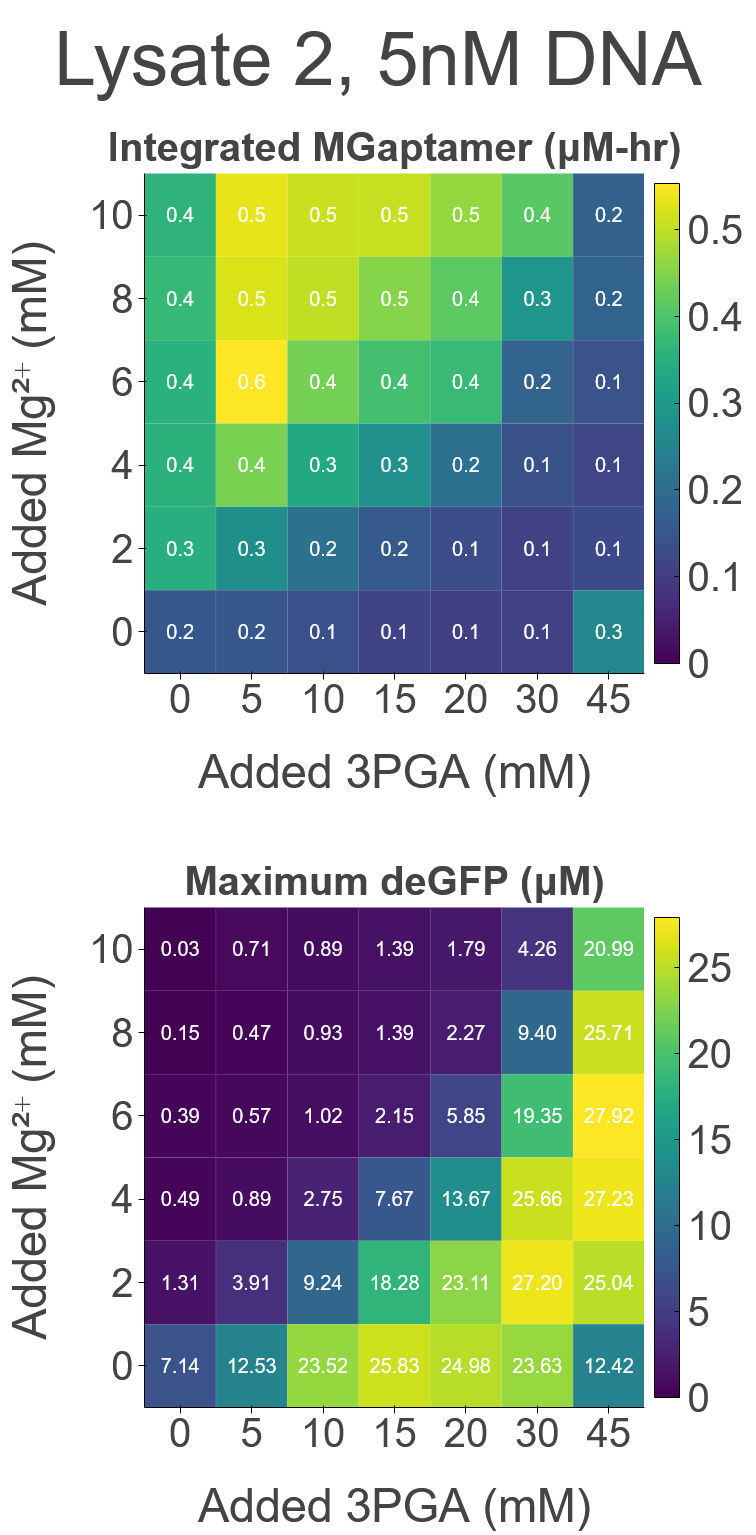

### FigS11.png

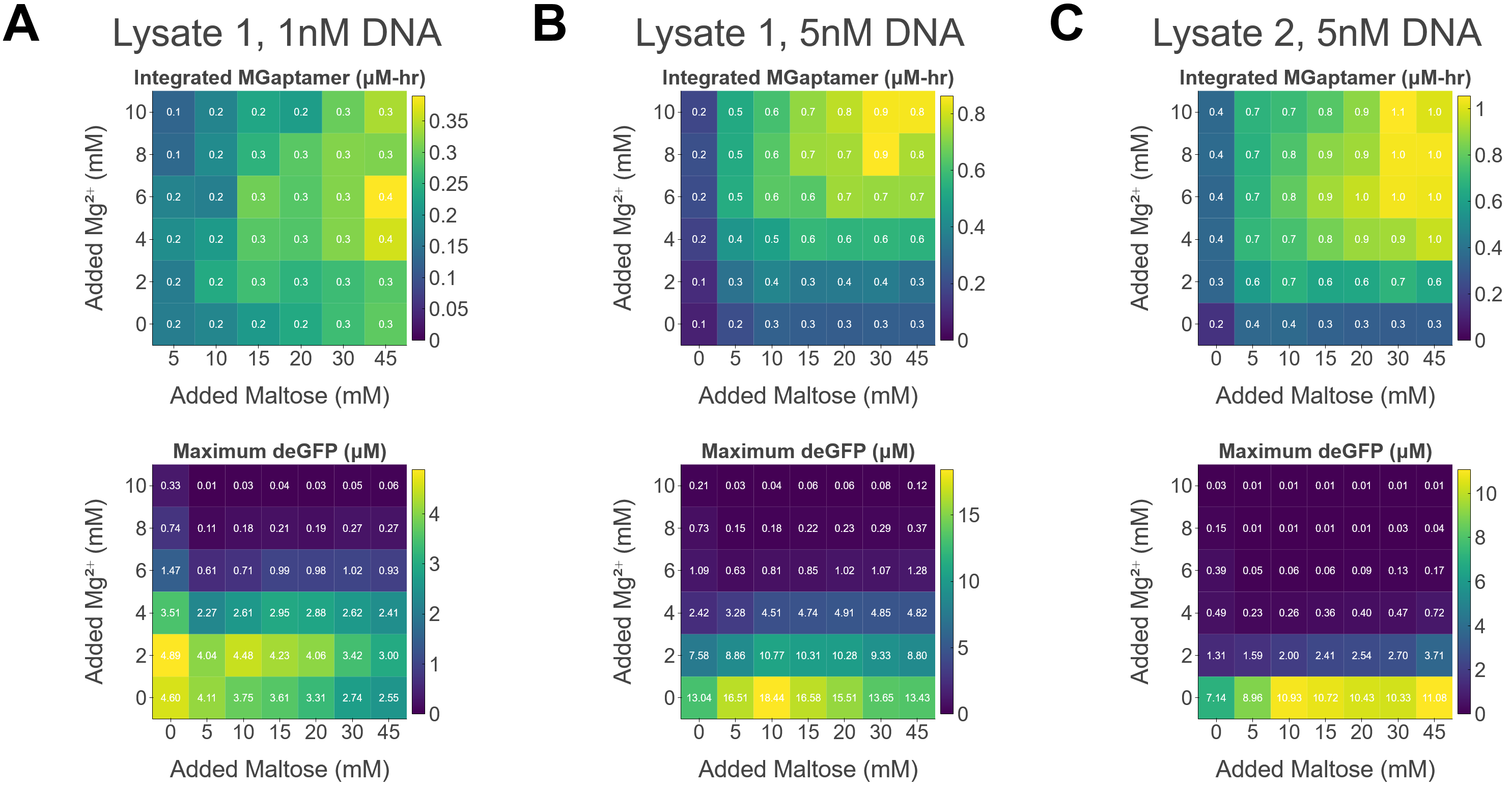

### FigS11_a.png

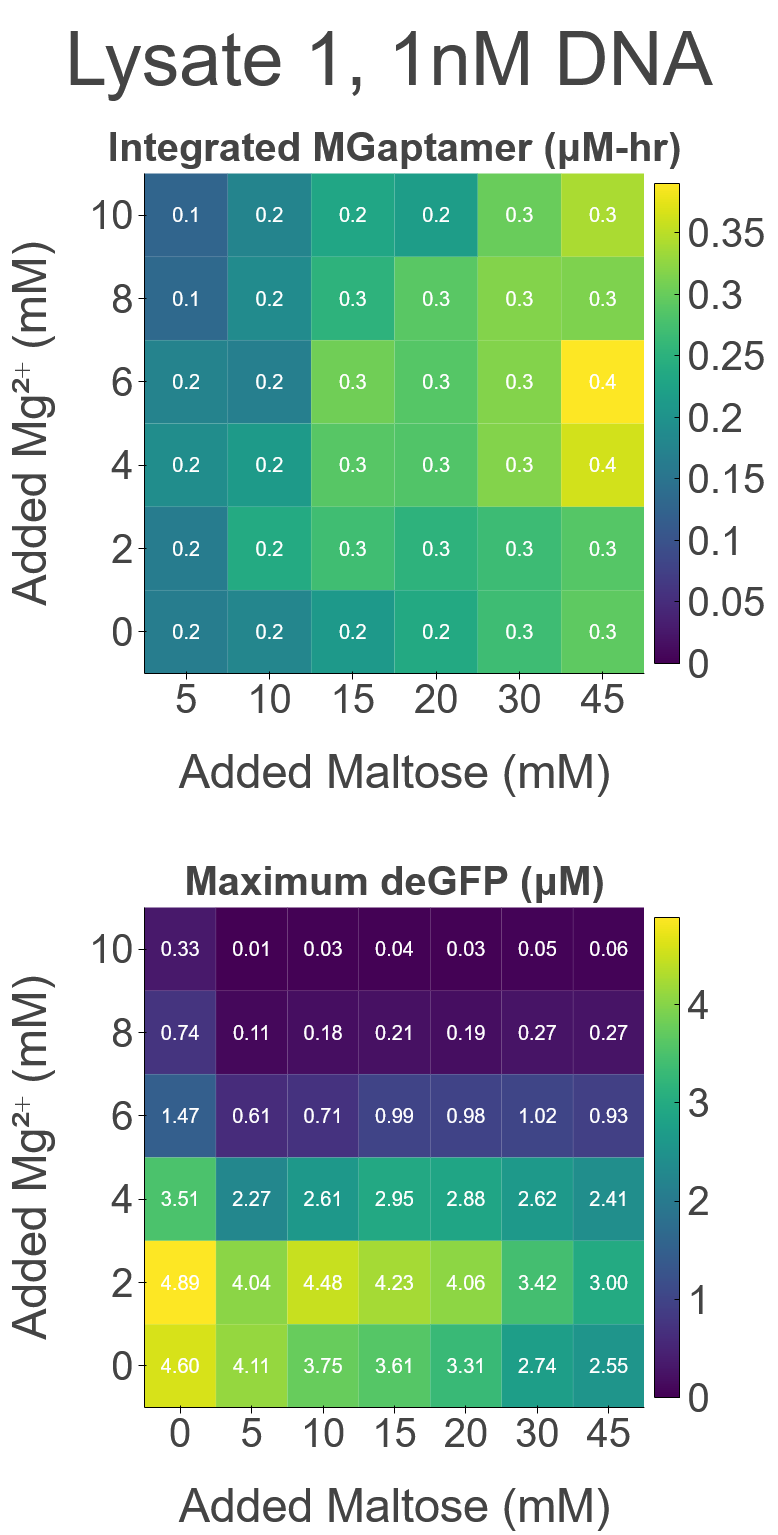

### FigS11_b.png

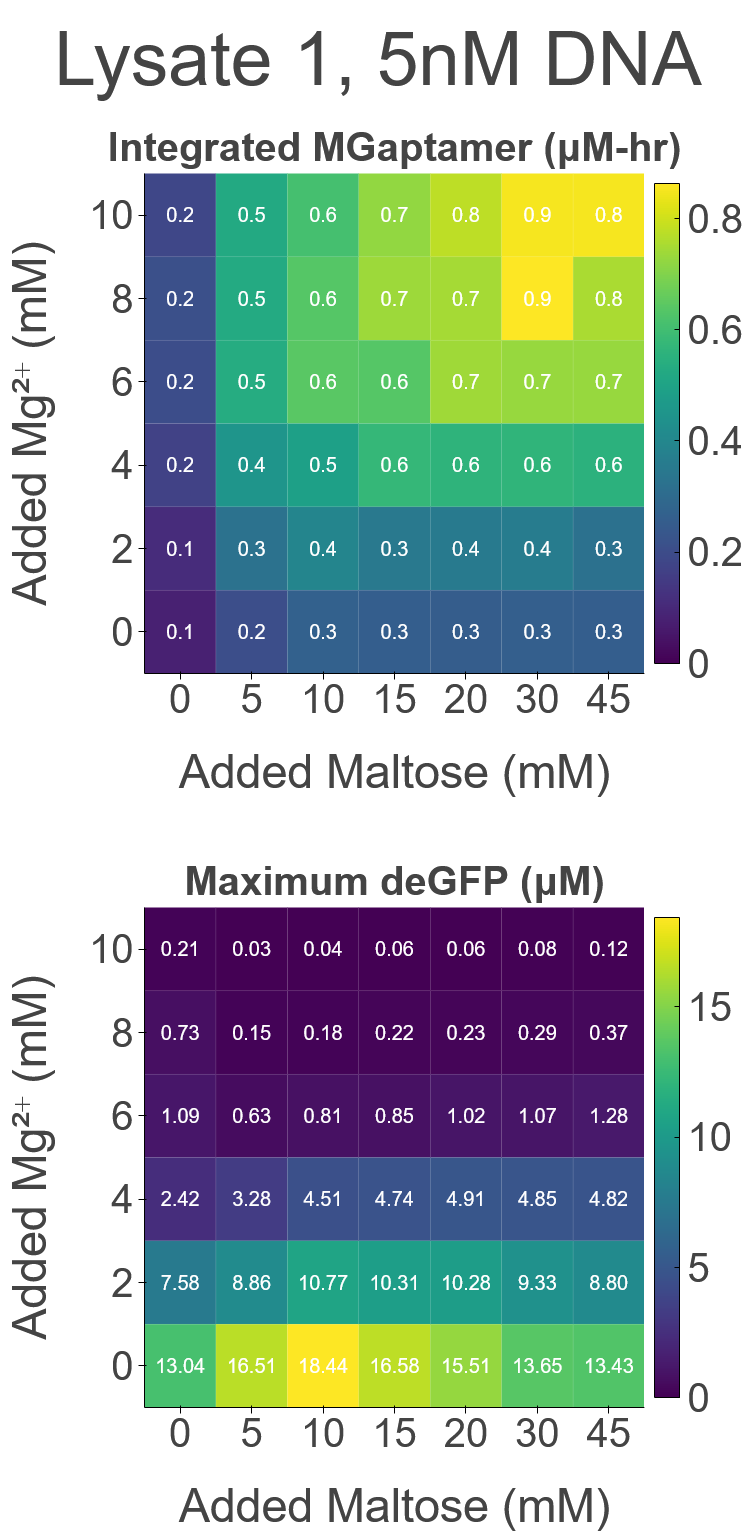

### FigS11_c.png

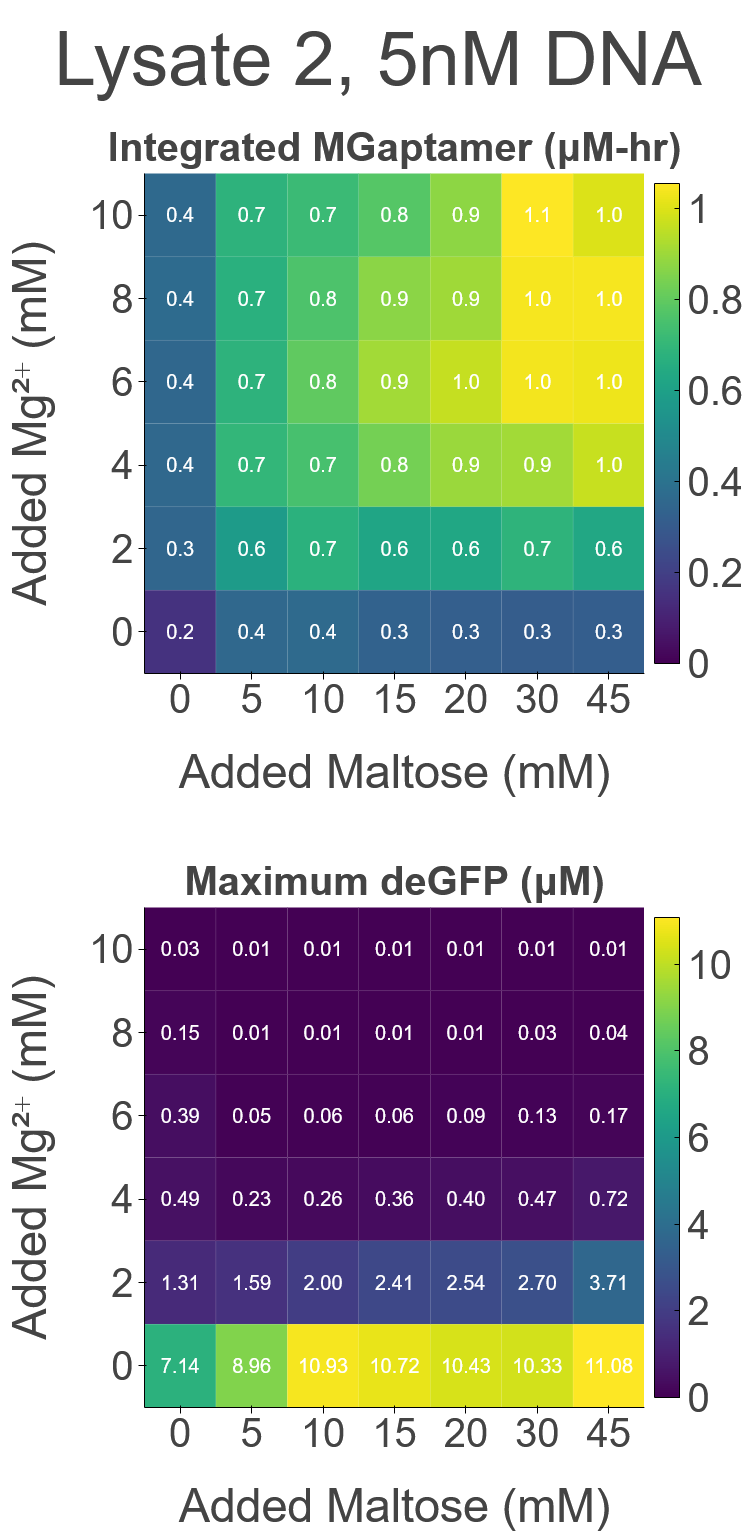

### FigS12.png

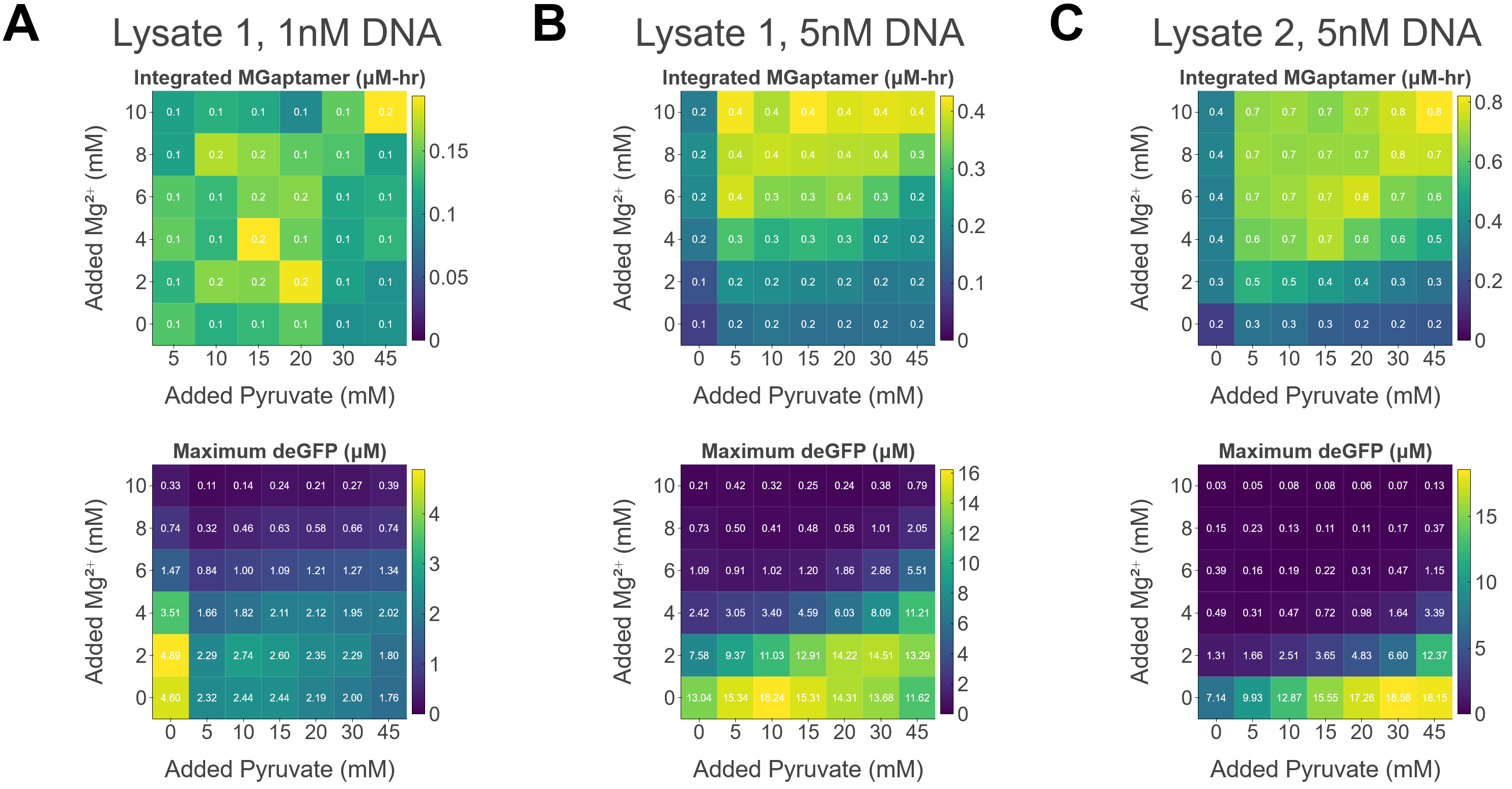

### FigS12_a.png

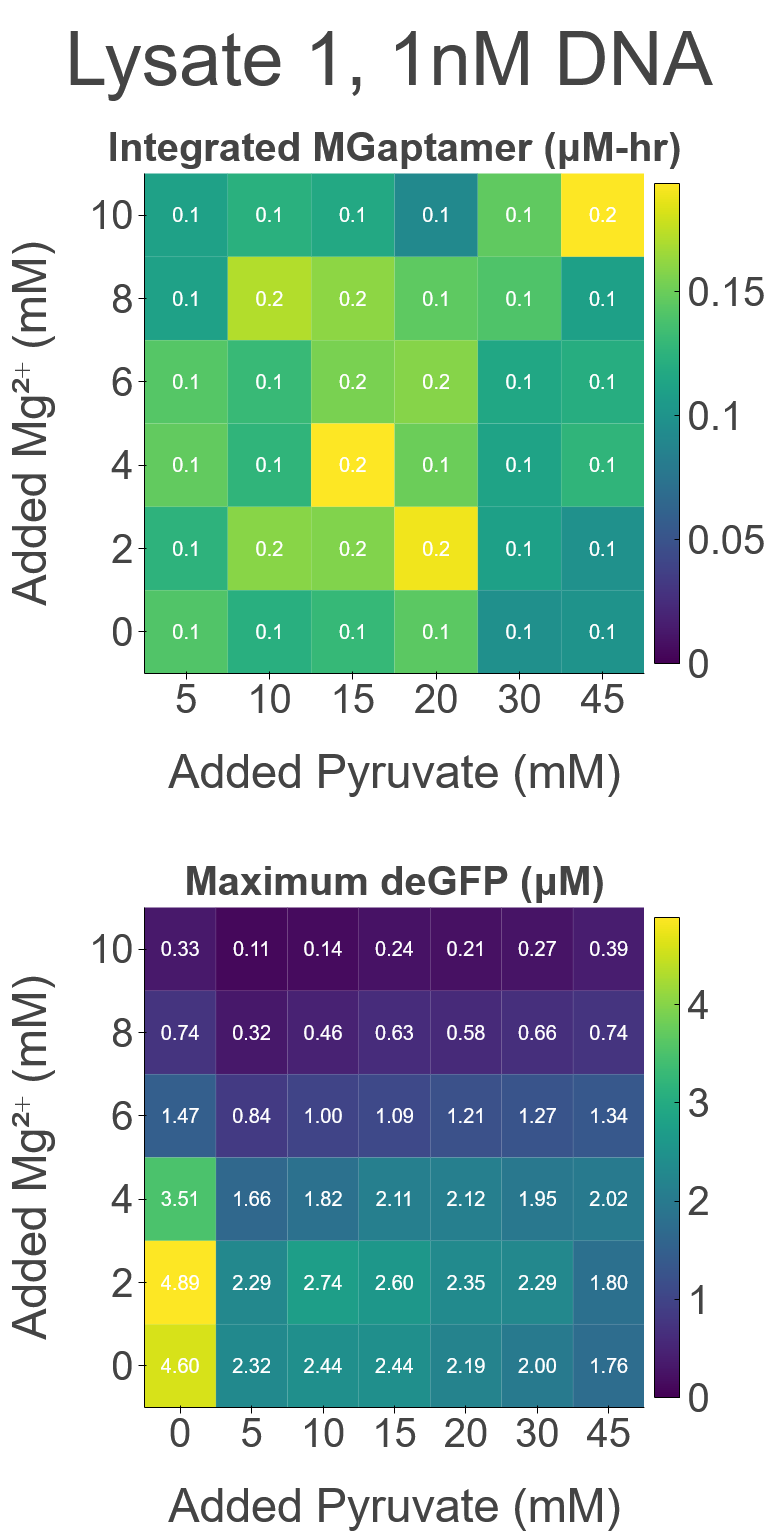

### FigS12_b.png

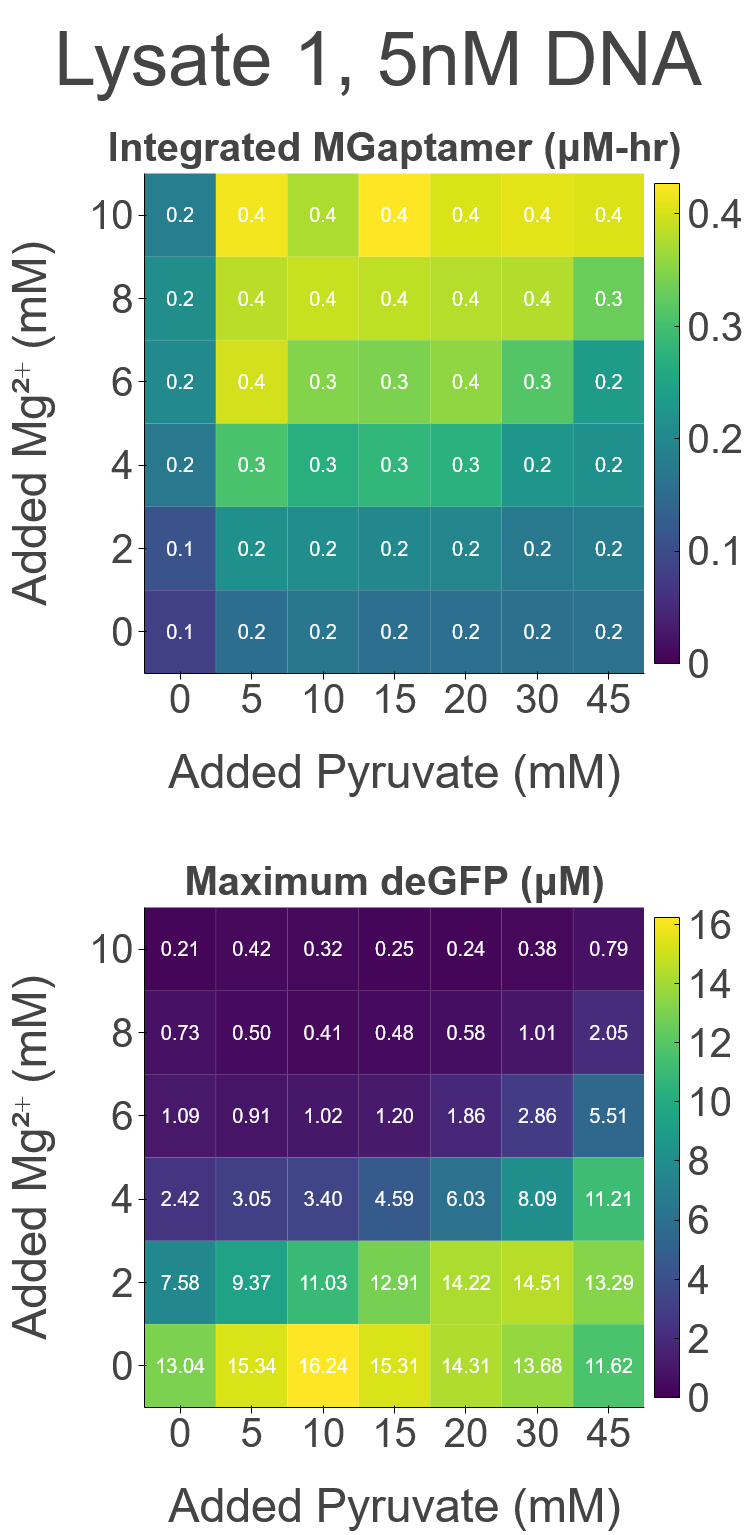

### FigS12_c.png

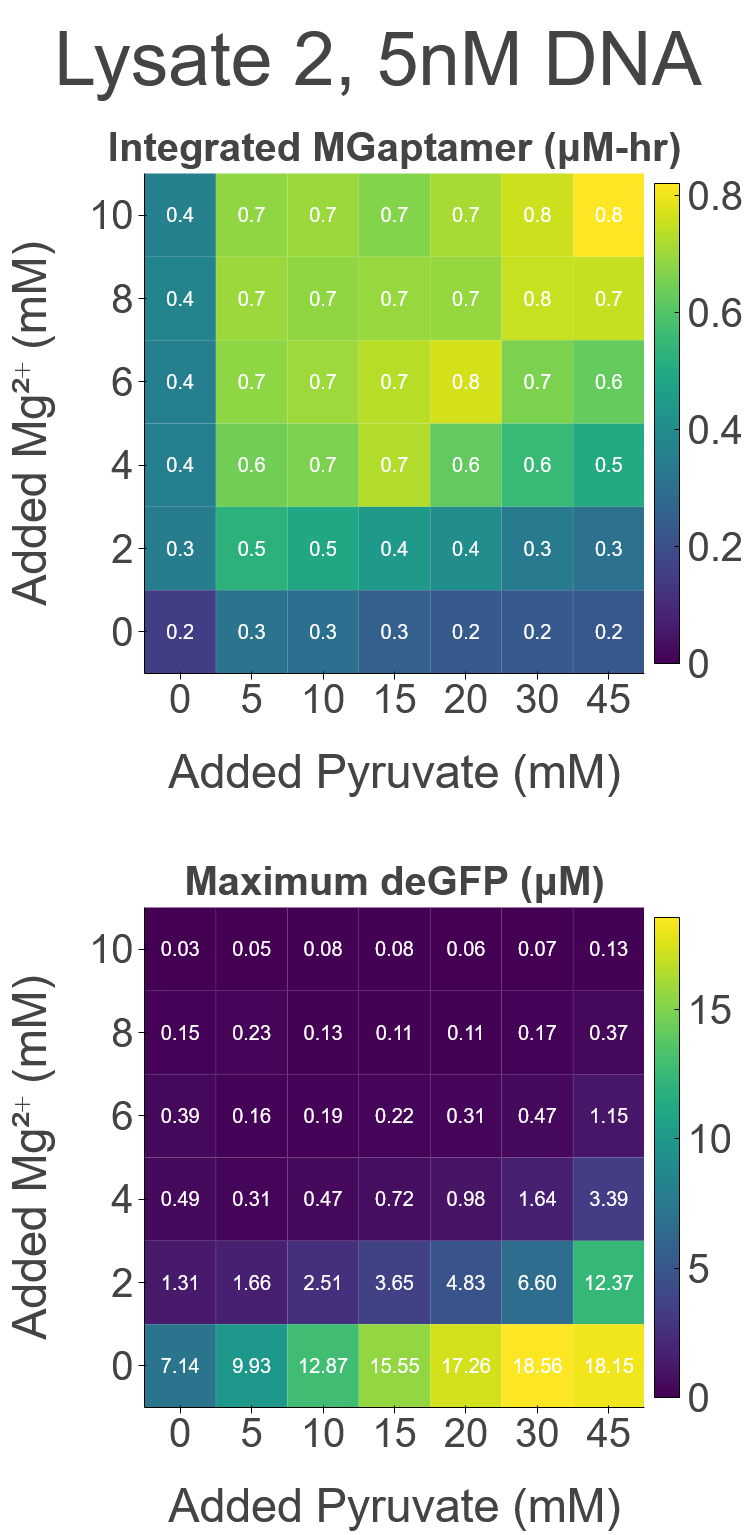

### FigS13.png

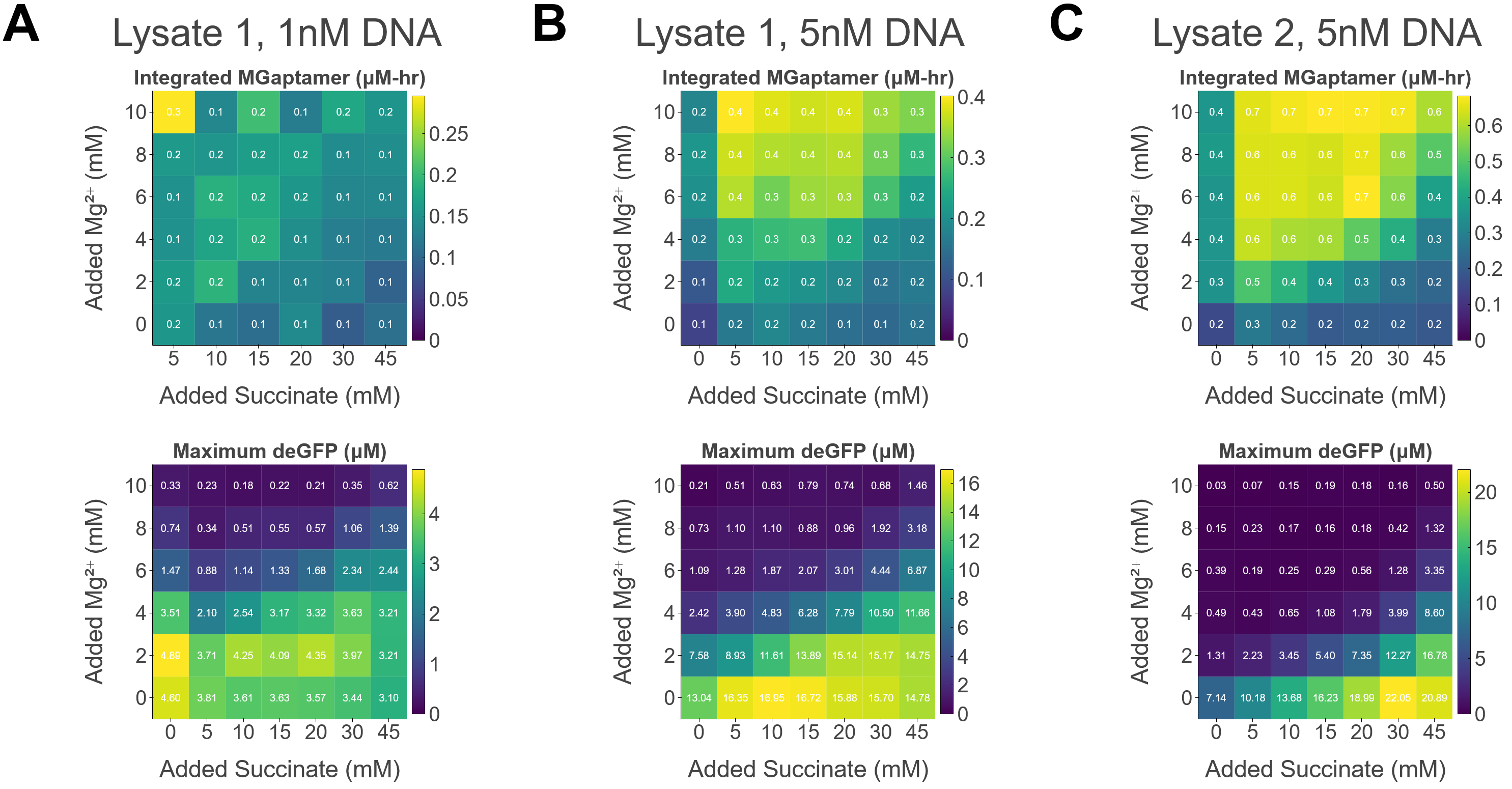

### FigS13_a.png

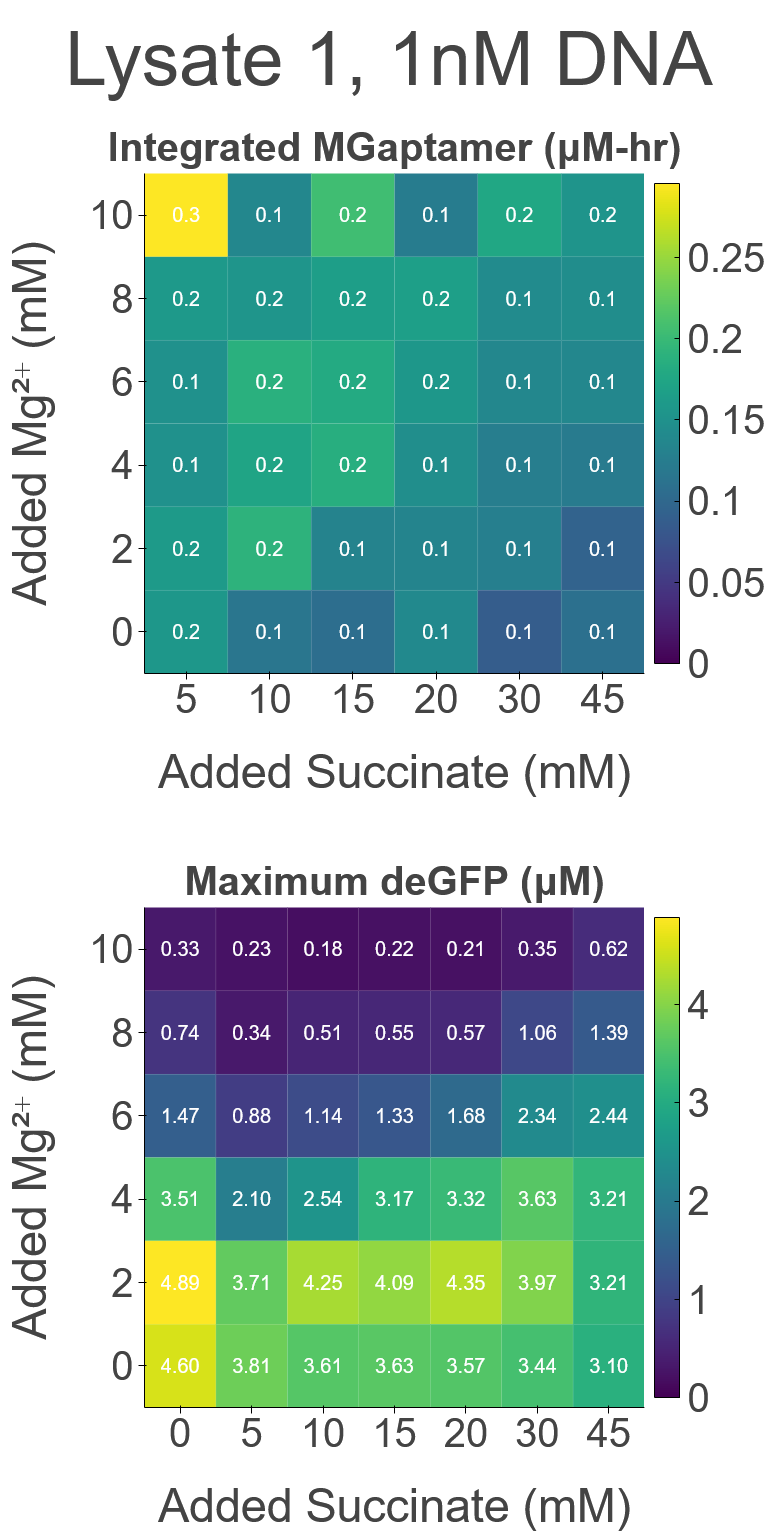

### FigS13_b.png

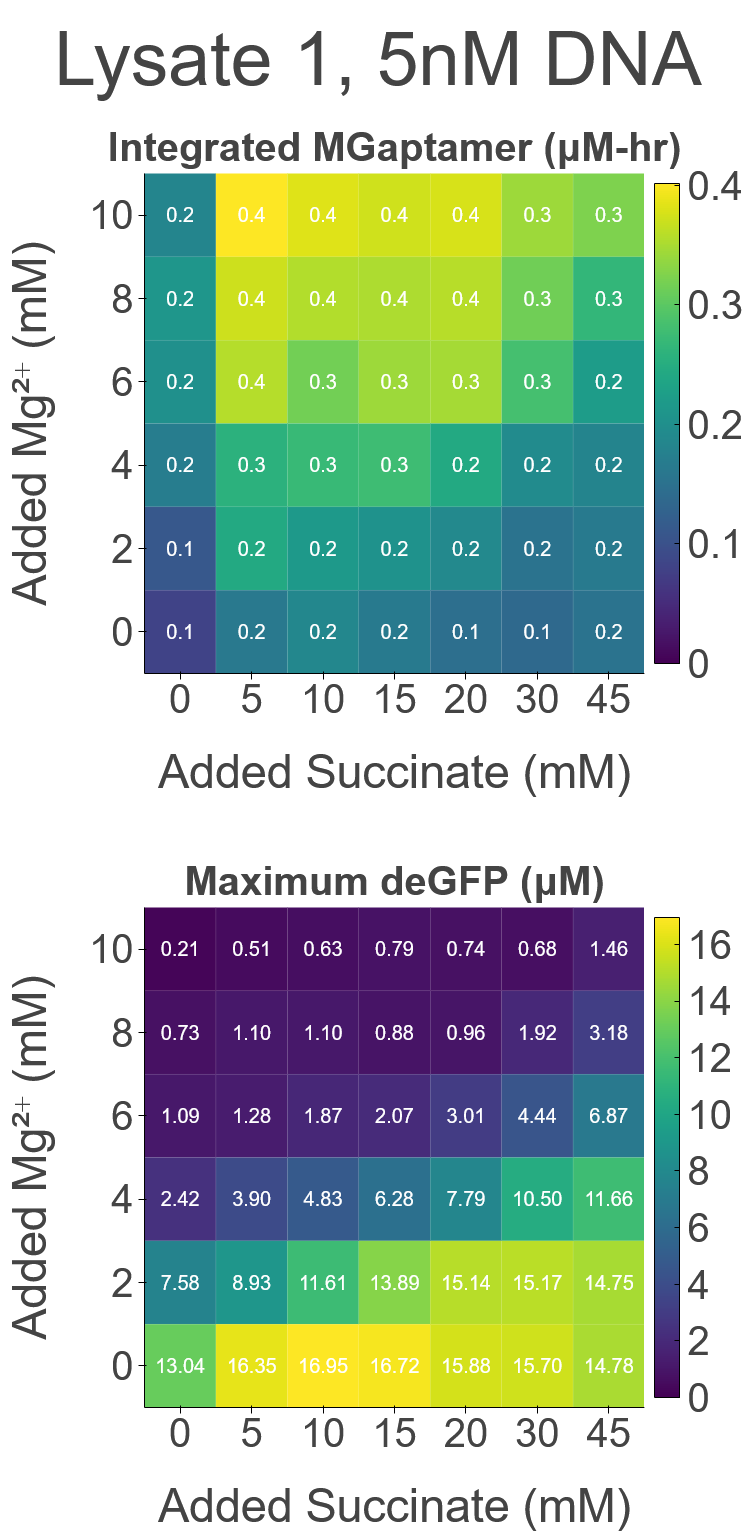

### FigS13_c.png

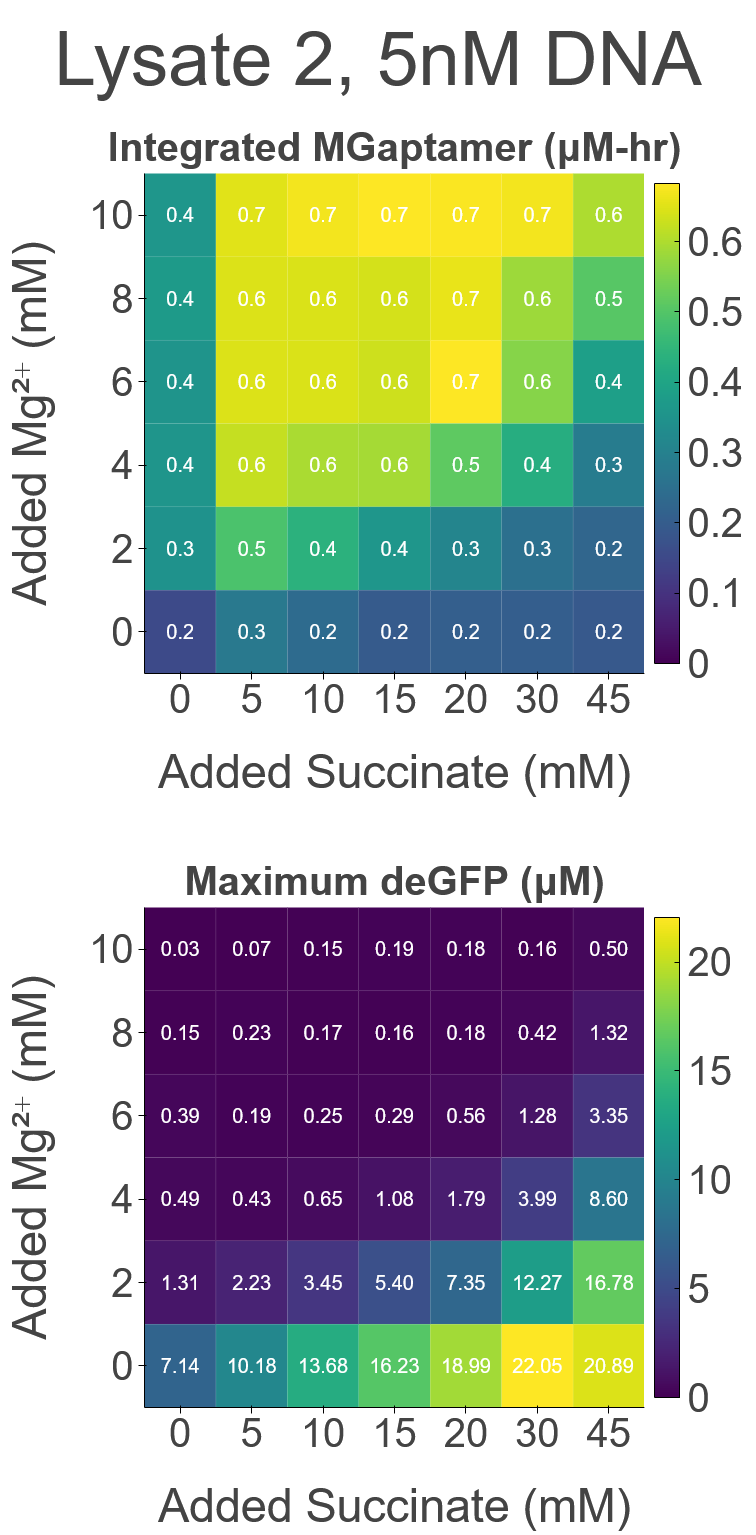

### FigS14.png

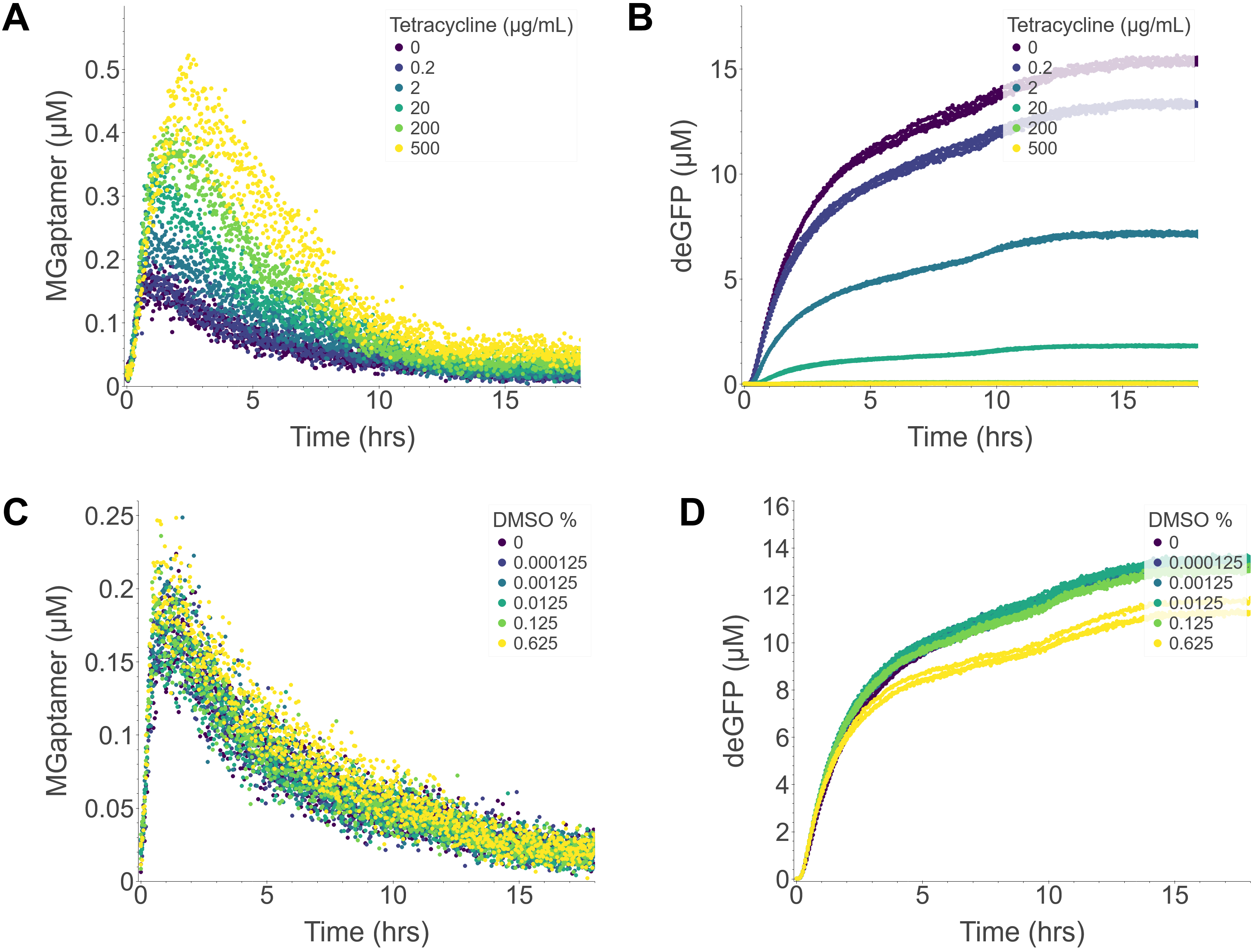

### FigS14_a.png

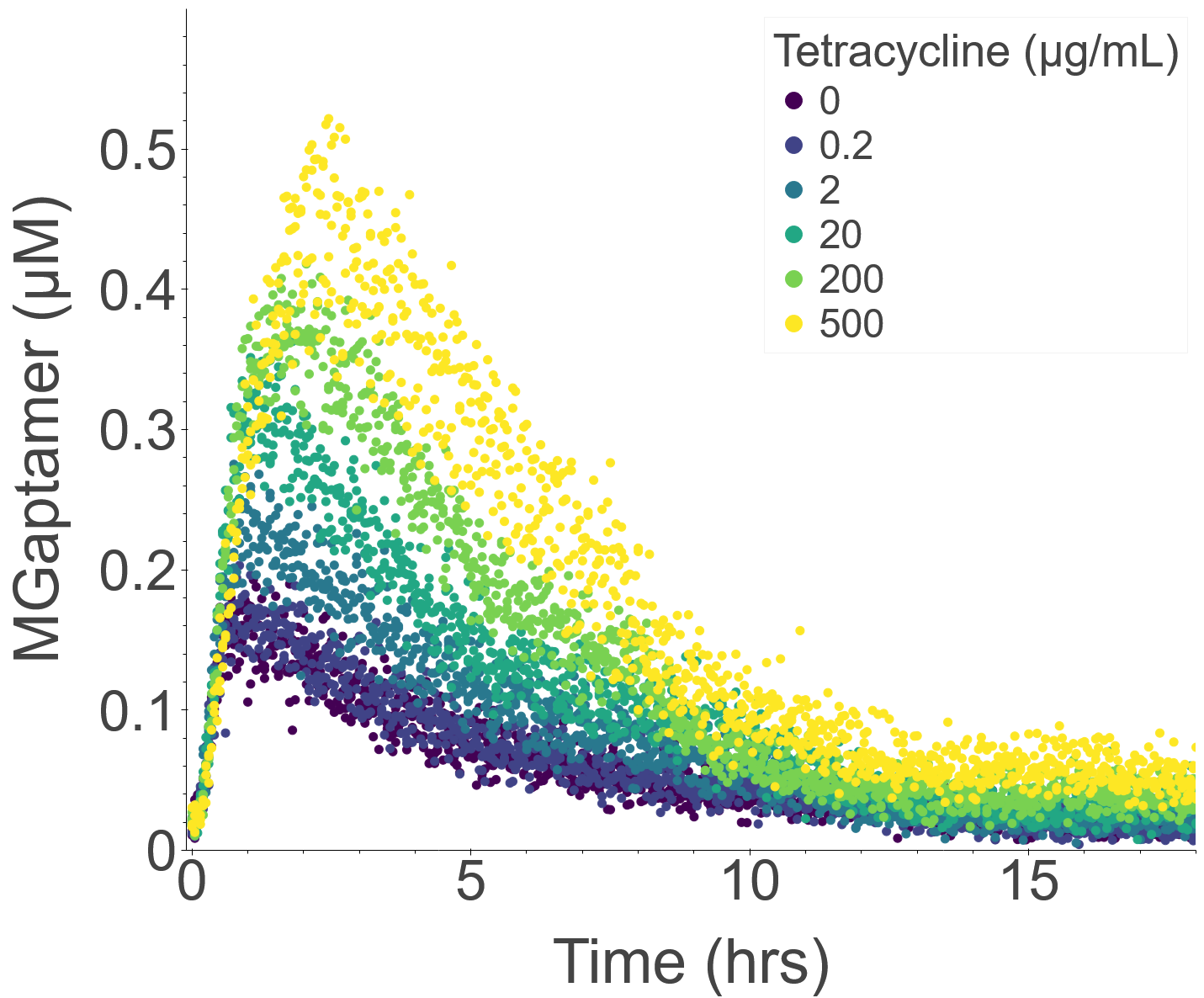

### FigS14_b.png

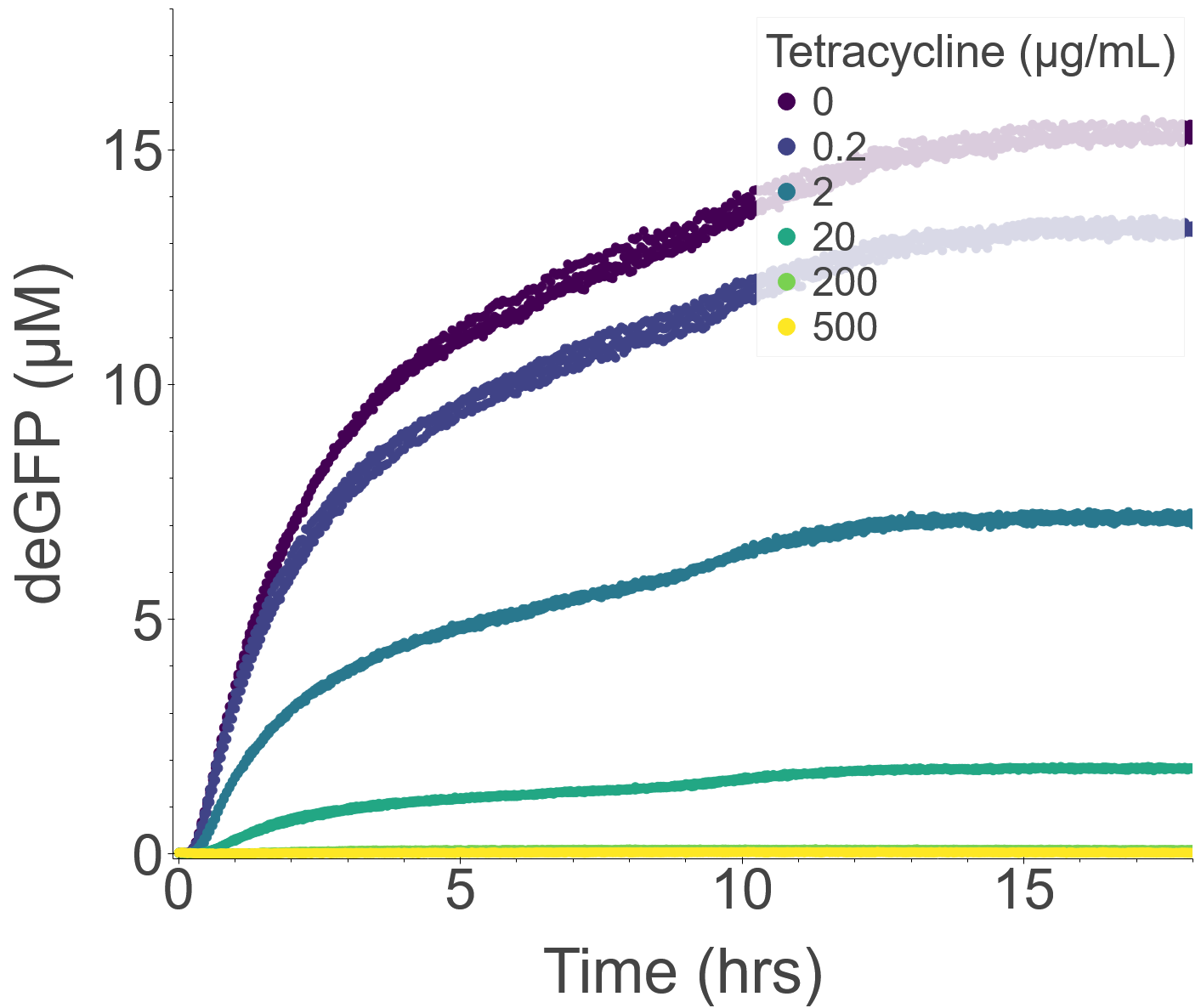

### FigS14_c.png

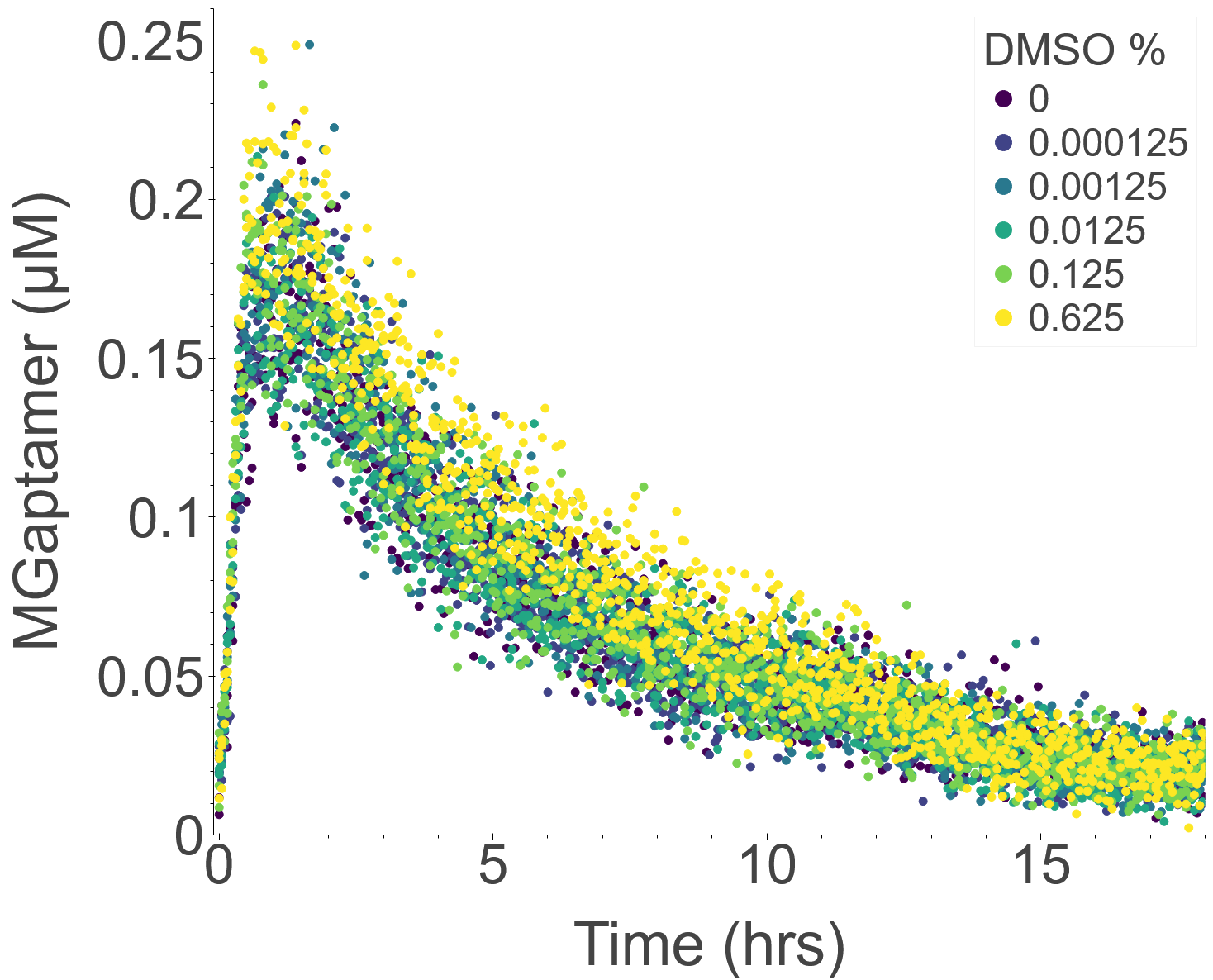

### FigS14_d.png

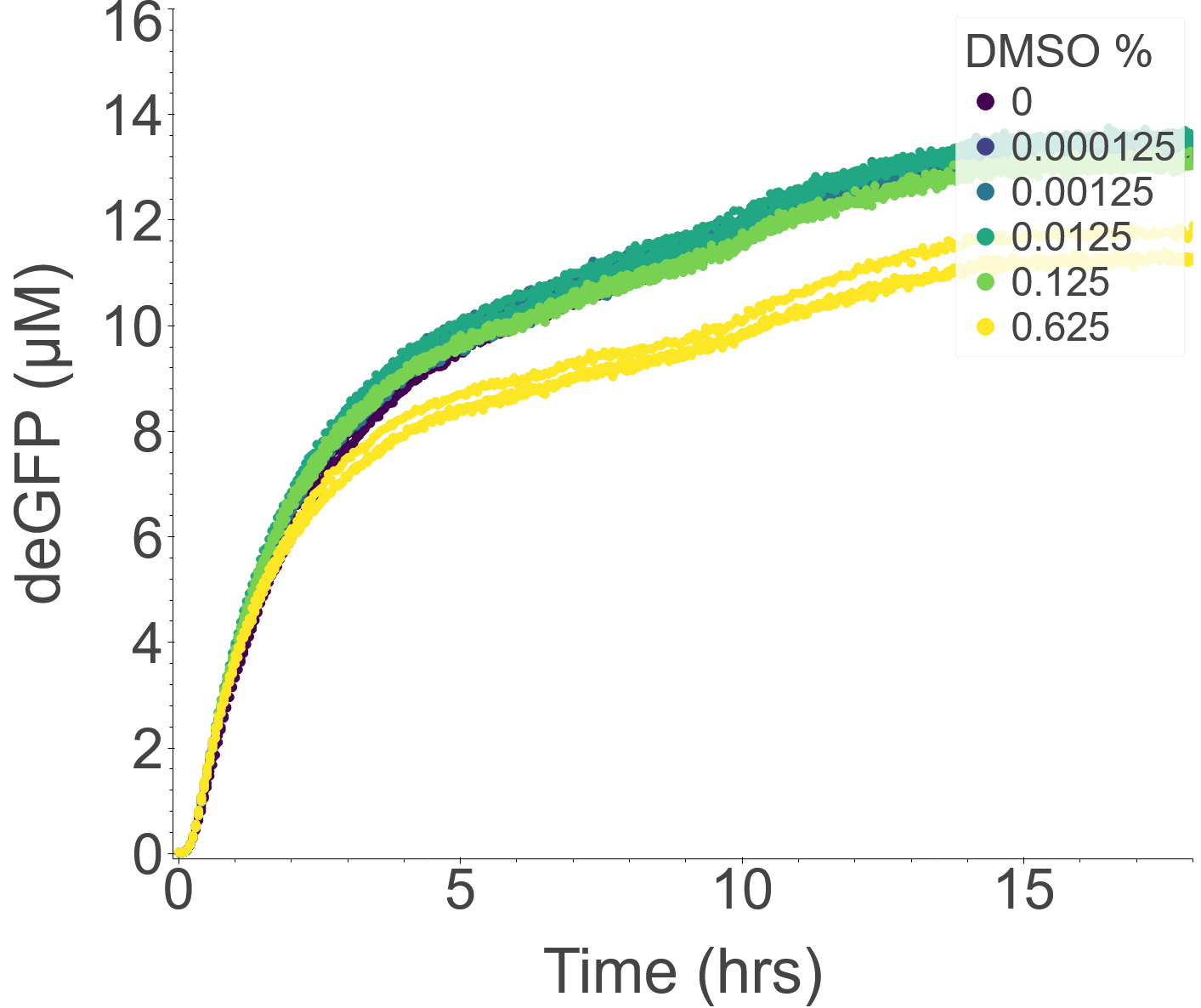

### FigS15.png

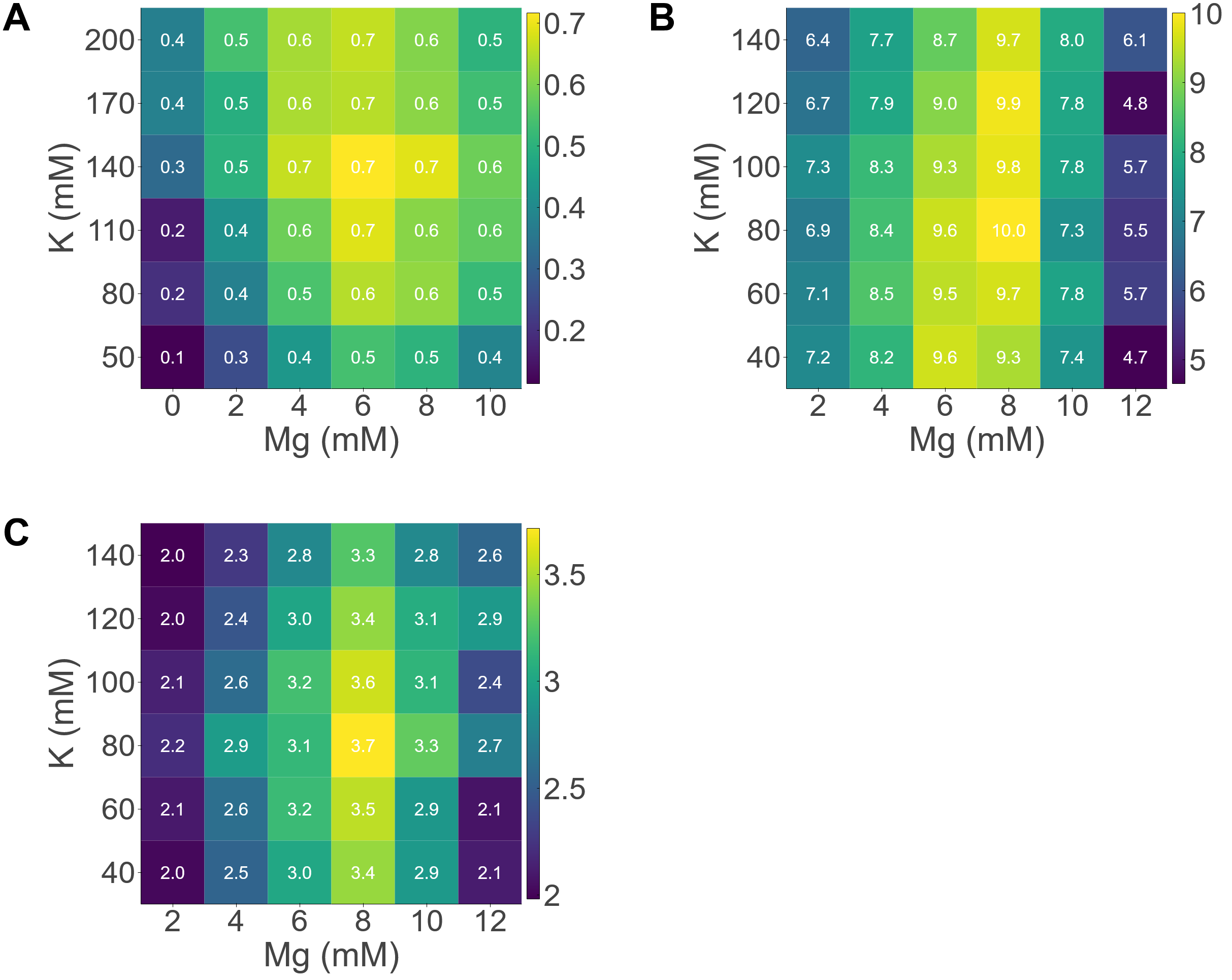

### FigS15_a.png

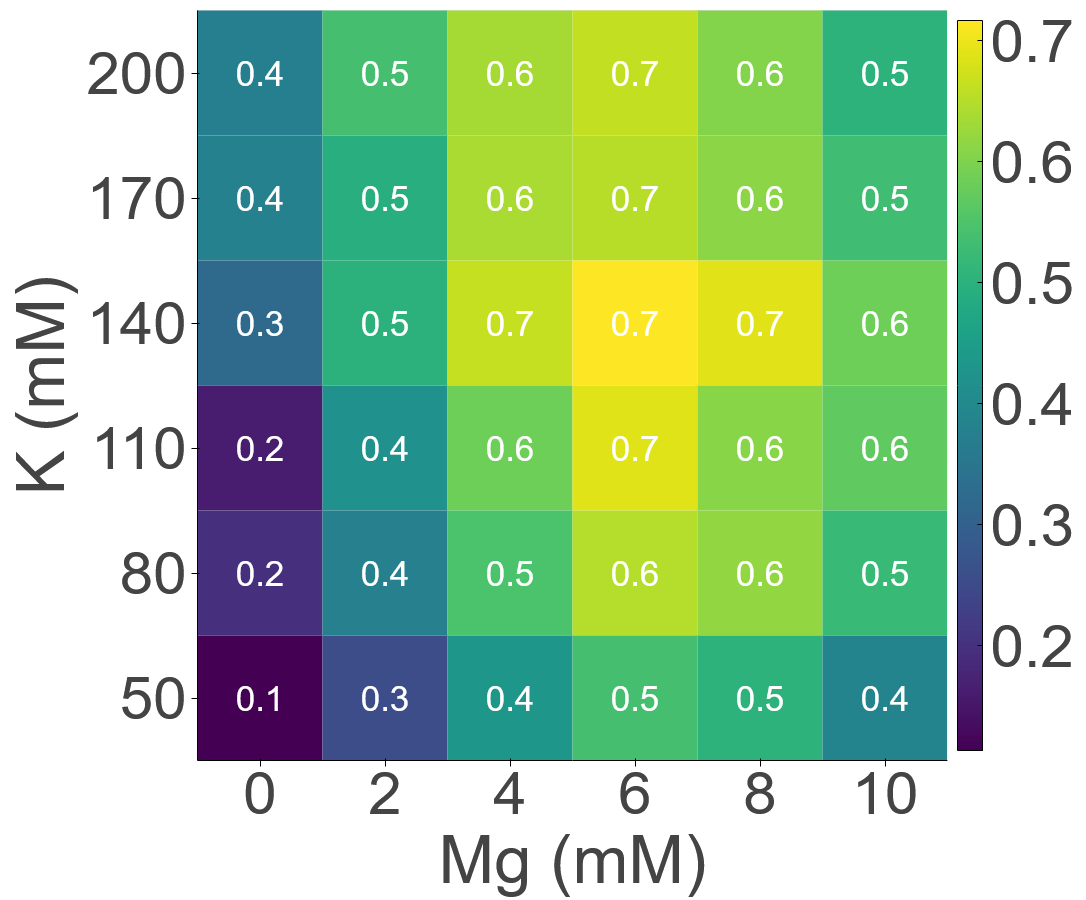

### FigS15_b.png

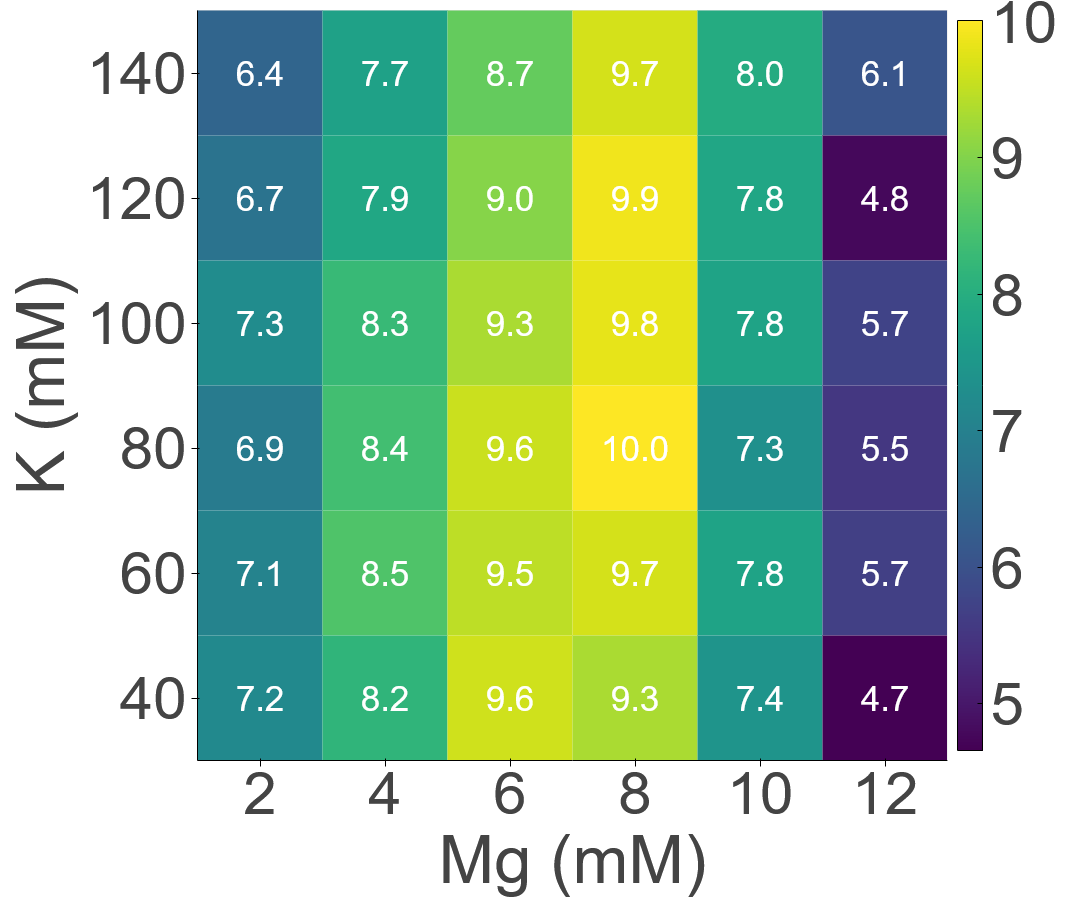

### FigS15_c.png

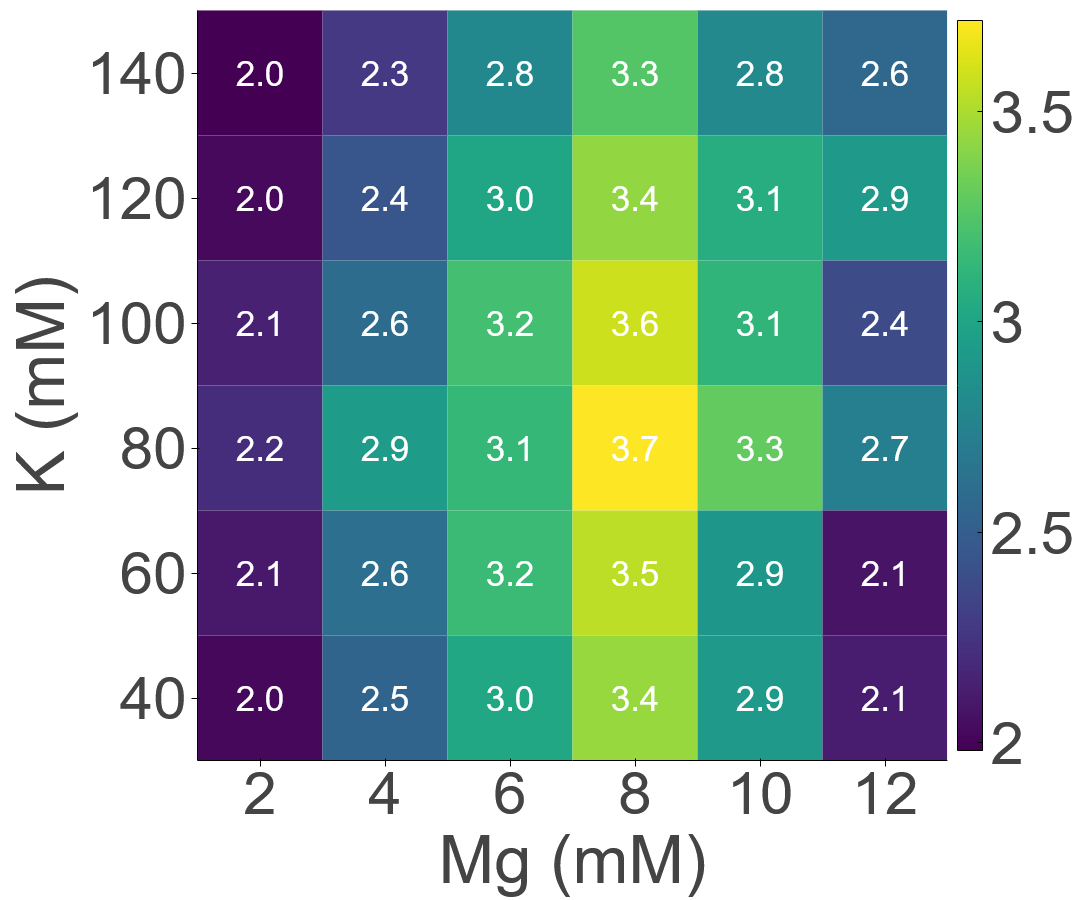
